# Antibiotics and the developing intestinal microbiome, metabolome and inflammatory environment: a randomized trial of preterm infants

**DOI:** 10.1101/2020.04.20.052142

**Authors:** Jordan T. Russell, J. Lauren Ruoss, Diomel de la Cruz, Nan Li, Catalina Bazacliu, Laura Patton, Kelley Lobean McKinley, Timothy J. Garrett, Richard A. Polin, Eric W. Triplett, Josef Neu

## Abstract

Antibiotic use in neonates can have detrimental effects on the developing gut microbiome, increasing the risk of morbidity. A majority of preterm neonates receive antibiotics after birth without clear evidence to guide this practice. Here microbiome, metabolomic, and immune marker results from the Routine Early Antibiotic use in SymptOmatic preterm Neonates (REASON) study are presented. The REASON study is the first trial to randomize symptomatic preterm neonates to receive or not receive antibiotics in the first 48 hours after birth. Using 16S rRNA sequencing of stool samples collected longitudinally for 91 neonates, the effect of such antibiotic use on microbiome diversity is assessed. The results illustrate that type of nutrition shapes the early infant gut microbiome. By integrating data for the gut microbiome, stool metabolites, stool immune markers, and inferred metabolic pathways, an association was discovered between *Veillonella* and the neurotransmitter gamma-aminobutyric acid (GABA). These results suggest early antibiotic use may impact the gut-brain axis with the potential for consequences in early life development, a finding that needs to be validated in a larger cohort.

## Main

Premature infants are particularly susceptible to infections secondary to increased need for invasive procedures and immaturity of the immune system, skin, and gastrointestinal tract^1–3^. Increasingly, there is growing concern that risk factors for mortality may originate from underlying pathologies that could also be responsible for premature birth^4^. Symptoms of prematurity are difficult to discern from symptoms of infection which, compounded by the increased risk of infection, have led to most premature infants being exposed to antibiotics early in life^5–7^. Despite high mortality rates, the incidence of culture positive early onset sepsis (EOS) is relatively low, between 0.2-0.6%^8^. In the absence of a positive culture, a majority of preterm infants receive antibiotics immediately after birth based on maternal risk factors (e.g. intra-amniotic infection) or laboratory abnormalities (e.g. elevated serum C-reactive protein (CRP)) because of the risk of mortality^8^. Given the low incidence of culture-positive EOS in this population, it is possible that such high rates of antibiotic use are unnecessary and may increase morbidity in these infants^9^. Other morbidities in the neonatal intensive care unit (NICU) such as necrotizing enterocolitis (NEC) and late onset sepsis (LOS) also have high mortality rates and have been associated with prolonged antibiotic exposure.^10,11^. Nevertheless, antibiotics remain the most commonly prescribed medication in the NICU^12,13^.

The gut microbiome comprises a highly volatile community structure early in life^14^. Microbial colonization is influenced as early as birth by mode of delivery, and perhaps even in the uterine environment by maternal factors^15,16^. Not surprisingly, antibiotic use has been shown to also change the composition of the preterm gut community^17–20^. Furthermore, antibiotic use early in life has increasingly been associated with adverse outcomes both short- and long-term^21,22^. One possible consequence is the disruption of the gut-brain axis (GBA), which involves bi-directional transmission of bio-molecular signals between the gut microbiota and the nervous system^23^. Aberrations in the GBA have been associated with altered immune homeostasis, as well as psychiatric, behavioral and metabolic conditions in adulthood^24^. It is therefore imperative to determine if such high rates of antibiotic use in preterm infants is necessary, as it could have lifelong consequences on future health.

Randomized clinical trials have the advantage of controlling for many of the numerous covariates that could interfere with answering whether preemptive antibiotic use in preterm infants affects outcomes. The Routine Early Antibiotic use in SymptOmatic preterm Neonates (REASON) study is the first to randomize symptomatic premature infants to either receive or not receive antibiotics soon after birth. Previously reported results from this study demonstrate the feasibility of such a trial and that withholding antibiotics did not lead to a significant increase in neonatal mortality or morbidity^25^. By employing a multi-omic approach, this cohort also provides the unique opportunity to understand how antibiotic intervention perturbs the early life gut microbiome, metabolome, and inflammatory environment in ways that may be consequential to health and development.

## Results

### Cohort and study description

Ninety-one of the total 98 enrolled infants had stool samples collected. Seven infants had no samples due to early mortality. Eligible infants were enrolled into groups based on previously described enrollment criteria25: group A – antibiotics indicated (n=28), group B – antibiotics not indicated (n=11), and group C – eligible for randomization (n=52). Twenty-six infants were from group C1 (antibiotics in first 48 hours) and 14 infants were from group C2 and did not receive antibiotics 48 hours after birth. For 12 infants (46%) randomized to group C2, antibiotics were prescribed in the first 48 hours after birth upon clinical assessment, and these infants were placed in a separate analysis group C2Bailed. One infant in group B were changed (bailed) to receive antibiotics within 48 hours after birth and was excluded from this analysis. Therefore, there are a total of 90 infants with stool samples analyzed across 5 enrollment groups, 2 of which did not receive antibiotics within 48 hours after birth (groups B and C2). Neither sex (p=0.352) nor mode of delivery (p=0.227) were significantly different between groups using the chi-square test. Both weight (p-value=0.005) and gestational age (GA) (p-value=0.002) were significantly different between groups overall, with group A infants on average with lower GA and at lower birth weights. Neither birth weight nor GA were significantly different between the randomized subgroups (C1, C2, C2Bailed) by the Kruskal-Wallis test (p>0.05). A summary of the infants in this analysis is provided (Table 1). Likewise, a summary of the types of antibiotics and the number of times antibiotics were prescribed by group (Supplementary Table S1). A full description of enrollment has been described previously^25^.

**Table 1.** Summary of infant enrollment, covariates, and samples. Summary of the number of enrolled infants per group used in this analysis and the number of infants changed from group C2 to C2Bailed. Enrollment groups are also summarized by infant sex (male::female), mode of delivery (vaginal::caesarean), gestational ages, maternal antibiotic exposure (yes::no) and birth weight ranges in grams. The number of infants and number of stool samples used in the metabolomics and immune marker analyses are listed.

|  | Group A | Group B | Group C1 | Group C2 | Group C2Bailed |
| --- | --- | --- | --- | --- | --- |
| Total infants | 28 | 11 | 26 | 14 | 12 |
| Sex (M::F) | 14::14 | 4::7 | 11::15 | 7::7 | 9::3 |
| Delivery mode (V::C) | 13::15 | 7::4 | 10::16 | 6::8 | 2::10 |
| Gestational age range (median) | 24 – 32 (28) | 29 – 32 (32) | 25 – 32 (29) | 23 – 32 (29) | 24 – 32 (28) |
| Maternal antibiotic exposure (Yes::No) | 20::8 | 6::5 | 20::6 | 8::6 | 11::1 |
| Birth weight range in grams (median) | 695 – 2132 (1015) | 1100 – 2770 (1820) | 525 – 2425 (1240) | 630 – 2116 (1223) | 605 – 1667 (888) |
| Samples post-normalization | 232 | 42 | 171 | 99 | 98 |
| Average number of samples/infant (median, 1st quartile, 3rd quartile) | 8.3 (9, 5, 11) | 3.8 (3, 2, 5) | 6.6 (6, 4, 8.75) | 7.1 (6, 4.5, 10) | 8.2 (9.5, 4.25, 11.25) |
| Number of infants with metabolomic samples | 4 | 0 | 2 | 2 | 2 |
| Number of metabolomic samples (samples per infant) | 32 (8, 10, 5, 9) | 0 | 17 (12, 5) | 16 (6, 10) | 25 (14, 11) |
| Number of infants with immune marker samples | 7 | 0 | 5 | 3 | 3 |
| Number of immune marker samples (samples per infant) | 40 (7, 7, 1, 3, 5, 7, 10) | 0 | 22 (12, 2, 2, 5, 1) | 21 (5, 10, 6) | 26 (13, 11, 2) |

Six hundred ninety-three stool samples were collected longitudinally for 91 of the total 98 enrolled infants. Stool data were not available for 7 infants due to early mortality. Sequencing data for 16S rRNA were obtained for 656 of those samples. After rarefying to an even sequencing depth of 10,000 reads per sample, 642 samples remained. Since GA is significantly different between groups, we chose to focus on corrected GA between weeks 28 to 39 because there were not enough samples among groups at younger and older timepoints. Therefore, 522 stool samples remained within this corrected GA window (Supplementary Table S2). The aim for this analysis is to test the effects of randomization to antibiotics vs. no antibiotics on the developing gut microbiome, metabolome and inflammatory environment using high-throughput 16S rRNA sequencing, quantitative PCR (qPCR), metabolomics, pathway inference, immune marker analysis and open-source statistical tools.

### Antibiotic use and trends in early gut microbiome diversity development

Using amplicon sequencing variants (ASVs), the richness and Shannon alpha diversity were not significantly different between enrollment groups at each corrected GA timepoint using the Kruskal-Wallis test (Fig. 1A, B). Furthermore, the number of copies of 16S rRNA were not significantly different between groups at any timepoint (Fig. 1C). Interestingly, group B infants who did not receive antibiotics and were typically older and had an increasing trend in copies of 16S rRNA over time, but the same trend was weaker for group C2 infants who also did not receive antibiotics but were typically younger. Using a linear mixed-effects model (LME) through Qiime2^26–28^, neither richness nor Shannon diversity changed significantly over the time frame of corrected GA between 28 – 39 weeks (p=0.407, p=0.861, respectively) (Fig. 1D, E). Notably, groups C1 and C2Bailed had significant positive trends in richness over time (p=0.019, p=0.002, respectively). Groups B and C2 had negative trends in richness that were not significant (p>0.05). All groups had positive trends in Shannon diversity development over time. However none were significant (p>0.05). Surprisingly, there was no significant difference in diversity between groups C1 (or similarly, C2Bailed) and C2, which are the informative groups for comparing effects of antibiotics or no antibiotics 48 hours after birth. Certain considerations need to be made when comparing infants by enrollment group. For example, group A infants had significantly lower GA and birth weights, particularly compared to group B infants who on average had the highest GAs and birth weights, and the shortest stays in the NICU. Comparison between group B and the other enrollment groups is limited in scope because of shorter stays, i.e. fewer longitudinal samples. Furthermore, although the randomized groups C1 and C2 had similar number of enrolled infants in the beginning, nearly half of group C2 infants were bailed to receive antibiotics 48 hours after birth. Therefore, the power to compare the randomized groups by antibiotic use with 48 hours after birth is limited.

**Figure 1.**
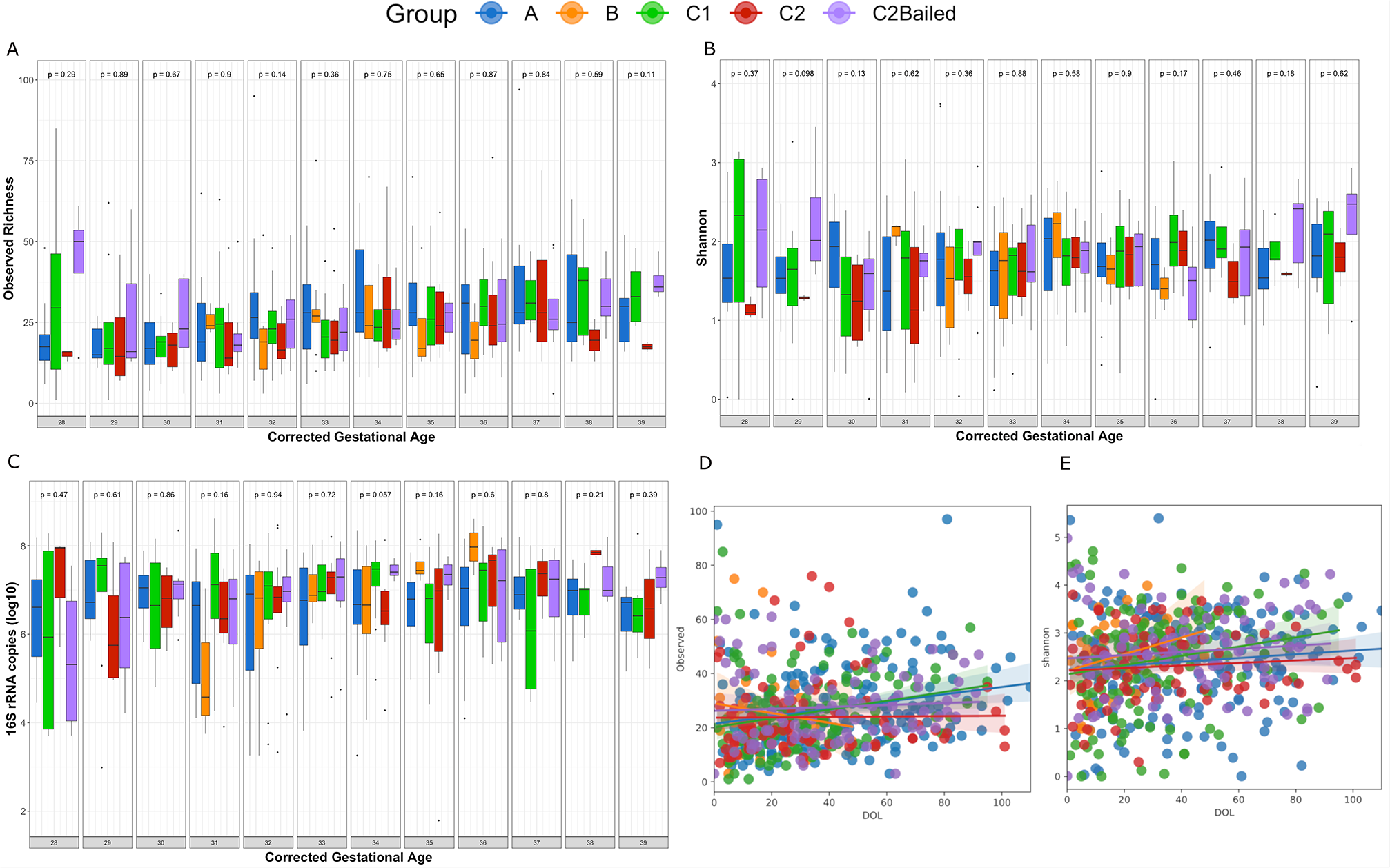
Antibiotic use 48 hours after birth does not significantly affect alpha diversity development. Boxplots displaying **(A)** the observed ASV richness **(B)** the Shannon diversity and **(C)** log_10_-transformed copies of 16S rRNA by enrollment group across corrected gestational ages between weeks 28 and 39. P-values were calculated at each corrected GA timepoint between enrollment groups using the non-parametric Kruskal-Wallis test. Linear mixed-effects modeling of the **(D)** observed ASV richness and **(E)** Shannon diversity over time between enrollment groups. Time scale on the x-axis is days of life (DOL) for corrected GA weeks 28 – 39. Greyed areas around each regression line represent 95% confidence intervals upper and lower around the coefficients.

Similar to alpha diversity, there were no apparent changes in overall bacterial community structure between groups when assessing beta-diversity over time and at each timepoint. Both the Bray-Curtis and Jaccard distance indices were used to assess community structure, taking into account quantitative ASV abundance and qualitative ASV presence/absence information, respectively. Principle coordinates analysis (PCoA) did not reveal any immediately obvious clustering differences between groups for either metric (Fig. 2), which may suggest little or no persistent effect of antibiotic use beyond 48 hours after birth. Beta dispersion was not significantly different between groups using the Bray-Curtis metric (ANOVA; Df = 4, Sum of Squares = 0.011, Mean Squares = 0.003, F = 1.088, p = 0.362) but was significant using Jaccard (ANOVA: Df = 4, Sum of Squares = 0.048, Mean Squares = 0.012, F = 9.574, p = 1.79E-07), specifically group A versus all other groups (TukeyHSD, A vs. B: p = 0.010, A vs. C1: p-value = 0.0003, A vs. C2: p = 0.004, A vs. C2Bailed: p=0.00001). This might suggest differences in dispersion heterogeneity (i.e. greater spread in variance) between group A infants and infants in other groups, which could be explained by group A infants often receiving antibiotics beyond 48 hours after birth. However, when the non-parametric permutational analysis of variance (PERMANOVA) test was applied to each timepoint across groups, there were no significant differences in Bray-Curtis or Jaccard distances among all groups at any given corrected GA timepoint (Fig. 2).

**Figure 2.**
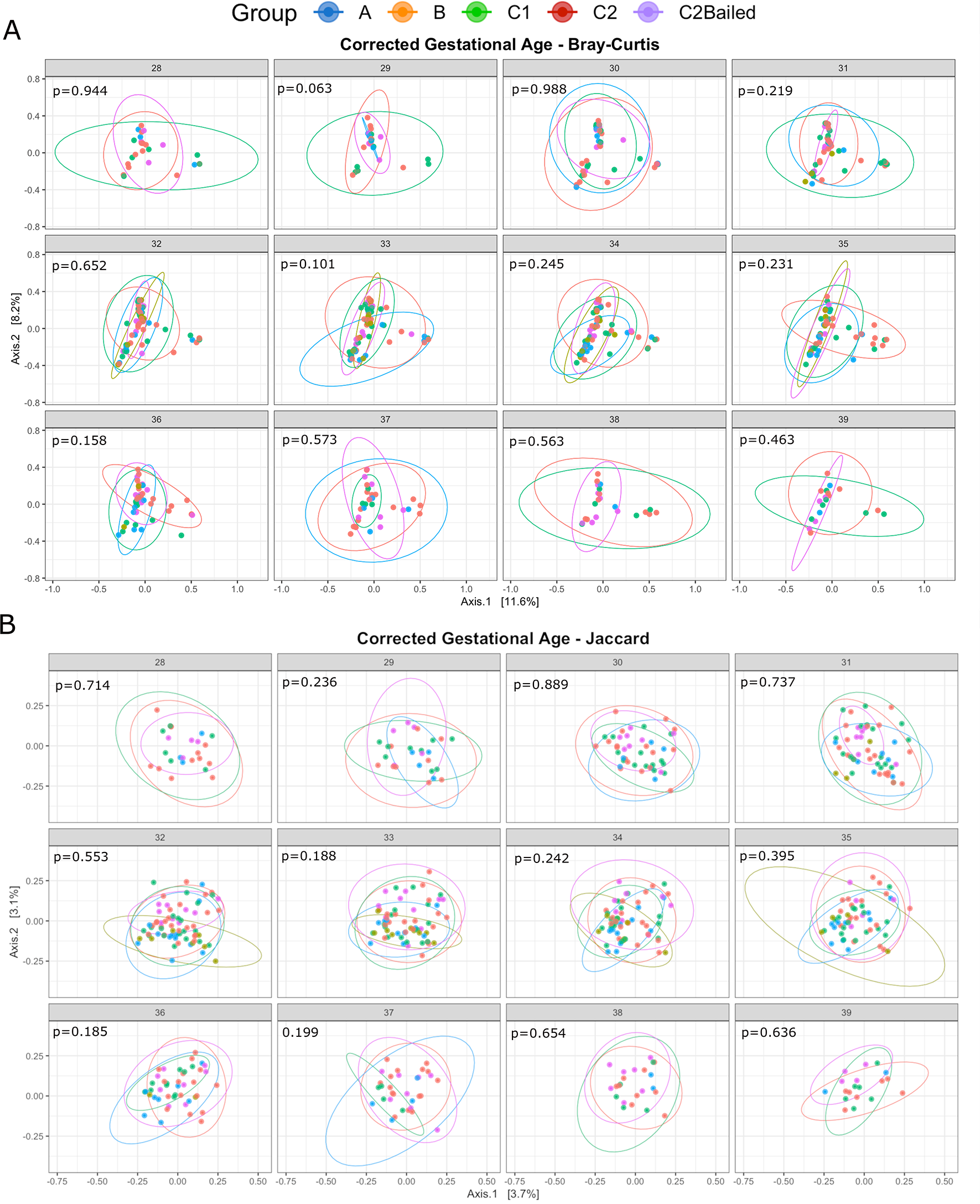
Antibiotic use explains little effect on beta diversity. PCoA ordination of stool samples using the **(A)** abundance-based Bray-Curtis and **(B)** presence/absence-based Jaccard distance metric among enrollment groups of corrected GA between 28-39. Ellipses are calculated based on a 95% confidence interval of a multivariate t-distribution.

### Feeding patterns drive changes in gut diversity and bacterial load

For preterm infants, diet generally consists of mother’s breast milk (MBM), pasteurized donor breast milk (DBM), formula, or some combination of these sources. Some infants also experienced periods of no enteral feeding (NPO: nil per os). To investigate effects of feeding while still considering effects from antibiotic use, feeding types were compared within each respective analysis group. In addition, for purposes of comparing feeding types at each corrected GA timepoint, feeding types with only a single sample at each timepoint (n=1) were removed. This reduced the total number of stool samples from 522 to 461. The number of samples in each group at each timepoint, and also by feeding type, is summarized in Supplementary Table S3. Using the calculated alpha diversity metrics described previously, feeding type was significantly different in bacterial richness only at corrected GA week 32 in group A infants (Kruskal-Wallis, p= 0.0069), where samples collected during feeding with all or partial mother’s milk tended to have higher bacterial richness (Fig. 3A). Furthermore, Shannon diversity was significantly lower in infants not fed orally at corrected GA week 36 in group A infants (p=0.042) (Fig. 3B). The log_10_-transformed number of 16S rRNA copies were not significant at any timepoint in any group, although notably feeding with MBM alone typically had higher 16S rRNA copies compared with formula in all groups except for group A, which might be explained by continued antibiotic use beyond 48 hours (Fig. 3C).

**Figure 3.**
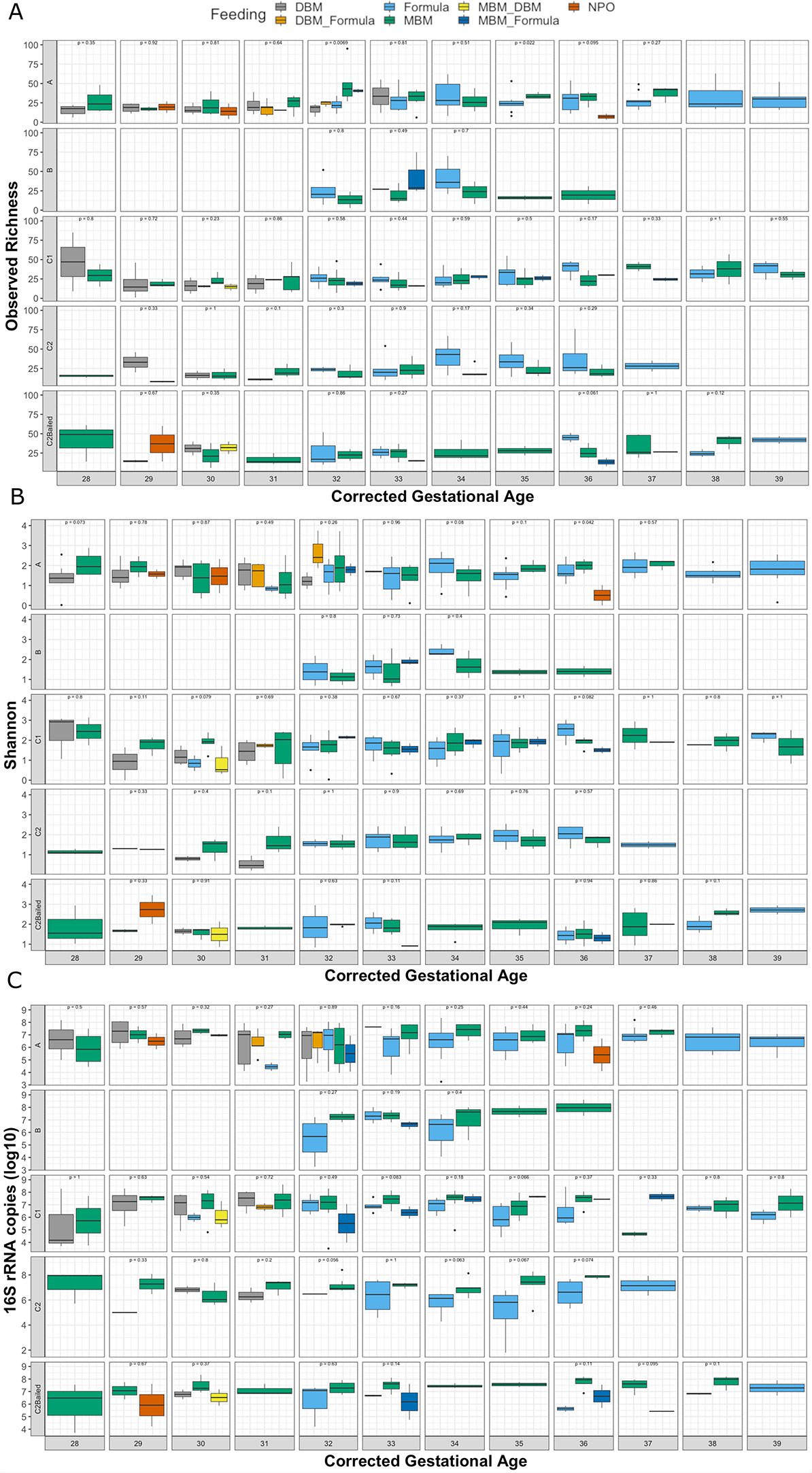
Effects of feeding patterns on the gut microbiome are transient over time. Effect of various feeding patterns on the **(A)** observed ASV richness **(B)** Shannon diversity and **(C)** log_10_-transformed copies of 16S rRNA by enrollment group over corrected GA from weeks 28 – 39. Only feeding patterns that have at least 2 samples at each timepoint were kept. Statistical comparison of feeding patterns at each corrected GA timepoint was performed using the non-parametric Kruskal-Wallis test.

Beta diversity between feeding types within groups was likewise only significant at few specific timepoints. For group A infants, Bray-Curtis distances between formula and MBM feeding were significantly different at corrected GA week 34 using PERMANOVA (R2=0.093, p=0.030). For group C1, Bray-Curtis distances were different between formula, MBM and MBM+formula at corrected GA week 36 (R_2_=0.370, p=0.005). Also in group C1, Jaccard distances were significantly different between formula, MBM and MBM+formula at corrected GA week 34 (R_2_=0.141, p=0.024), corrected GA week 35 (R_2_=0.158, p=0.011) and corrected GA week 36 (R_2_=0.296, p=0.007).

Applying LME modelling by feeding type according to analysis group allows the observation of feeding effect over time, focusing again on the 12 weeks of corrected GA after removal of feeding type singletons at each timepoint (Fig. 4). Perhaps not surprisingly, among all groups, periods of NPO led to a lower trend in Shannon diversity over time (p=0.003) (Fig. 4G). While bacterial richness appeared to trend lower, the trend was not significant (p=0.341) (Fig. 4A). For group A infants, which received antibiotics in 48 hours after birth and often beyond, MBM was associated with a slight increase in richness (p=0.009) (Fig. 4B), and periods of NPO led to a lower trend in Shannon diversity (p=0.031) (Fig. 4H). in Group B infants, who never received antibiotics and tended to be older, larger and healthier, all feeding types including formula (p=0.004), MBM (p<0.001) and MBM+formula (p=0.018) led to increasing trends in Shannon diversity (Fig. 4I). However, no significant trends could be identified in richness (Fig. 4C). It is difficult to evaluate group B infants due to lower enrollment size, shorter NICU stays, and fewer samples overall. For the randomized infants that received antibiotics 48 hours after birth, MBM and formula were associated with positive trends in richness (C1 and C2Bailed MBM: p<0.001; C1 formula: p=0.004; C2Bailed formula: p=0.016) (Figs. 4D, 4F). Only group C2Bailed infants saw increased trends in Shannon diversity for feeding MBM (MBM only: p=0.018; MBM+DBM: p=0.004; MBM_formula: p=0.002) (Fig. 4L). Finally, group C2 infants randomized to not receive antibiotics 48 hours after birth saw a lower trend in both richness and Shannon diversity during feeding with DBM (Figs. 4E, 4K). However, neither trend was significant, likely due to the few numbers of samples. The power for detecting trends in group C2 is likely hampered because half of the infants randomized were bailed within 48 hours after birth.

**Figure 4.**
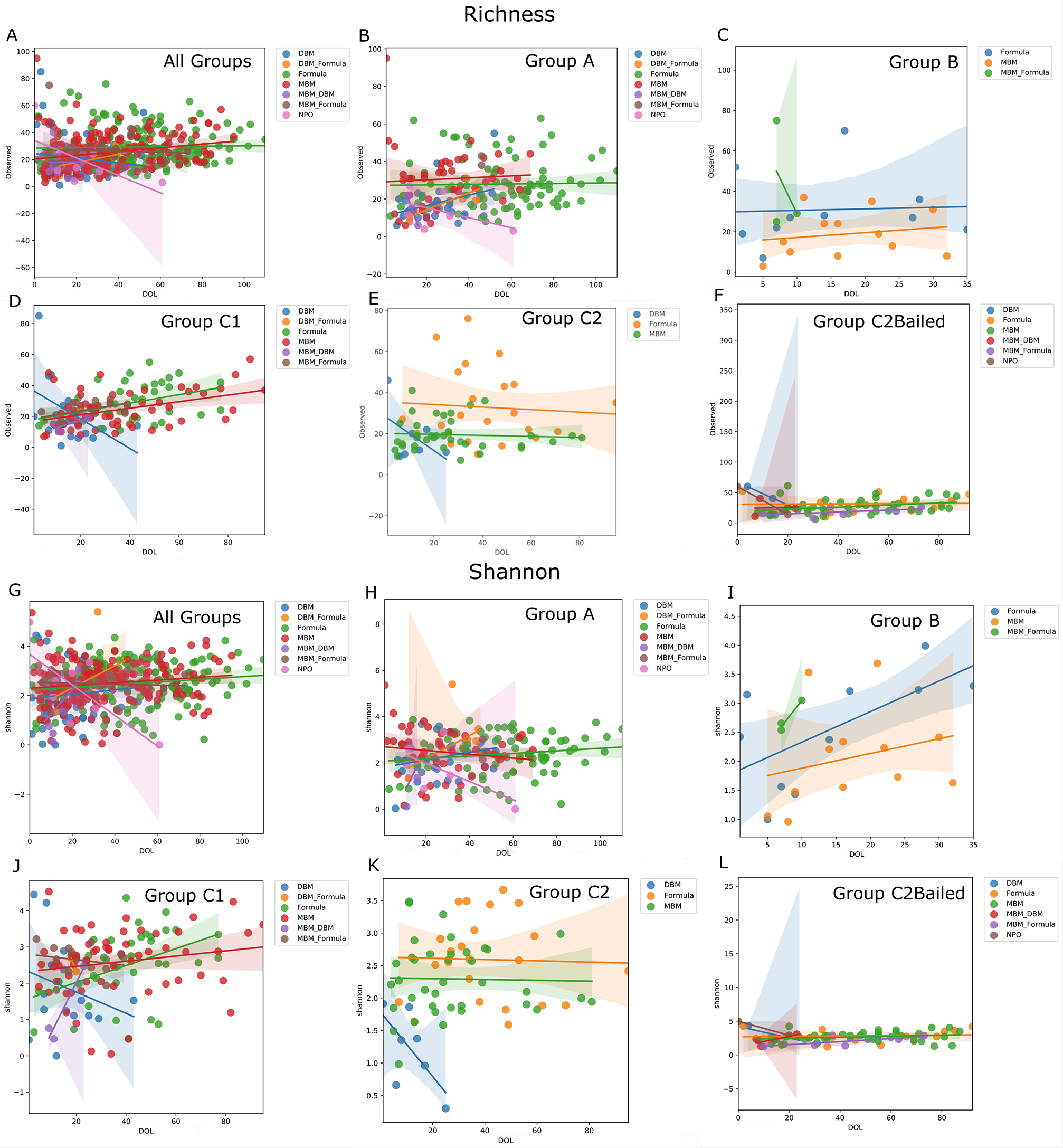
Linear mixed effects modelling identifies feeding effects by group over time. Linear mixed-effects results plotted as observed bacterial richness over time by group and feeding types **(A – F)** and the Shannon diversity over time by group and feeding types **(G – L)**. The number of samples used in this analysis by group and by feeding type are listed in Supplementary Table S3.

### Gut microbial community development is highly variable and unique to each infant

Although the infants in this study were analyzed among 5 groups, each infant’s stay in the NICU is highly personalized, by type, frequency, number or length of antibiotic use, type and length of feeding patterns, and adverse clinical events over time. To aid in visual identification of patterns throughout the NICU course, detailed charts were created for each infant that depict both clinical and laboratory data over time (Days Post Birth) integrated into a single graphic per infant (Fig. 5 and Supplementary Figure S1). This includes results from the 16S rRNA analysis as pie charts for each stool sample, color coded by bacterial taxonomy and sized based on the log_10_-transformed number of 16S rRNA copies per gram of stool, as well as adverse clinical events coded by a single letter code (Fig. 5). With these visualizations, patterns are more easily observed between antibiotic treatments, feeding types, and the gut microbiome. Typically, the microbiome composition after birth is homogenous in composition and diversifies, as well as increases in size over time, as might be expected. One interesting association involved the administration of the anti-fungal fluconazole and a resultant lowering of bacterial load and diversity. However, fluconazole was often administered in conjunction with antibiotics and was administered frequently to group A infants.

**Figure 5.**
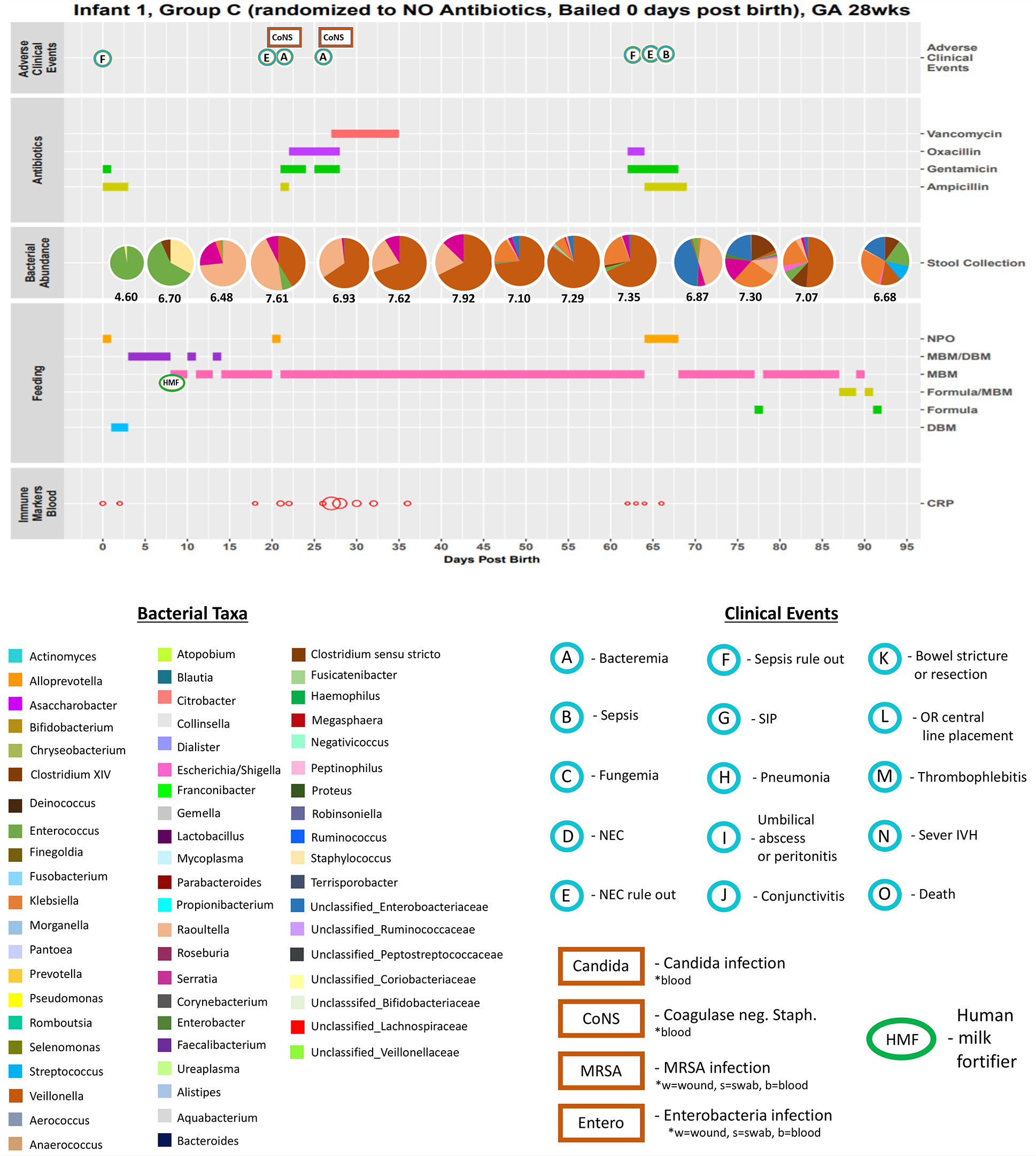
Integration of clinical and laboratory data gives detailed view of infant stay in NICU. Extensive clinical and laboratory data, when combined, provide a detailed summary of each infant’s stay in the NICU. Data included in each chart from top to bottom include: the infant ID, group assignment, antibiotic change status (bail), gestational age, any adverse clinical events (which are further described in Supplementary Figure S1), the type and duration of antibiotic use (if any), the copy-number corrected absolute composition of each weekly stool sample and it’s log_10_-scale number of bacterial 16S rRNA copies, the type and duration of each feeding including administration of human milk fortifier, the relative levels of C-reactive protein measured from blood, and relative concentrations of measured stool immune markers (for infants where these measurements were performed). DBM: donor breast milk, MBM: mother’s breast milk, NPO: no enteral nutrition, CRP: C-reactive protein, EGF: epidermal growth factor. Listed below is a key for the color-coded bacterial taxa used in the stool 16S rRNA copy-number corrected composition pie chart for each infant chart. A key for the bacterial color codes, adverse clinical events (including infections by body site) and the administration of human milk fortifier for each infant chart is given below.

The more subtle effects of antibiotic use and feeding patterns become visually apparent by taking this individualized approach. To illustrate, infant 5 was exclusively fed mother’s milk from day 14 through day 71 post birth. At day 31, antibiotics vancomycin and piperacillin were administered (Supplementary Figure S1). During treatment, *Veillonella* was entirely removed from the stool microbiome, falling from 3.23E+06 cells per gram (73%) from the pre-treatment time point to an undetectable level while antibiotics were used. At day 47, 9 days after antibiotic treatment ceased, *Veillonella* again dominated the stool at 3.83E+06 cells per gram (80%), almost a complete replacement of levels before treatment. Also, the proportions of the other 2 genera found in the stool, *Escherichia* and an unclassified Enterobacteriaceae spp., were nearly identical after treatment and continuation of mother’s milk as before treatment (pre-treatment: *Escherichia* – 21%, Enterobacteriaceae spp. – 4.7%; post-treatment: *Escherichia* – 15%, Enterobacteriaceae spp. – 3.3%). Thus, either that mother’s milk effectively restored the stool microbiome to its pre-treatment state or this effect occurred due to removal of antibiotic selective pressure, or both. A similar effect can be seen in infant 12 between 25 and 50 DOL where the microbiome is restored post-antibiotics. In this case, the restoration is observed with MBM, DBM, and formula. In some cases, antibiotic use had no effect on the microbiome composition (e.g. infant 42, 84), suggesting the presence of resistance mechanisms in the dominant gut microbes (in these 2 cases, members of Enterobacteriaceae). In fact, Enterobacteriaceae presence followed administration of ampicillin and gentamicin, the 2 most commonly prescribed antibiotics immediately after birth. This occurred in 24 of the 91 infants. Other times antibiotic use appears to dramatically and irreversibly change microbiome composition (e.g. infant 25).

### Bacterial genera correlate with stool metabolites and inferred metabolic pathways

In addition to 16S rRNA profiling, 90 stool samples from 10 infants were analyzed for metabolomic profiling (Table 1). Four of the 5 groups were included in these samples for comparison (group B samples not included). Peak height responses were recorded for 454 identifiable metabolites. To determine if gut bacteria were associated with relative concentrations of metabolites in stool, the top 10 most abundant bacterial genera associated with identified metabolites were determined. Repeated measures correlation values were plotted using a heatmap, which indicated numerous significant, positive and negative, associations between bacteria and metabolites (Fig. 6A). Interestingly, *Veillonella* were positively associated with the neurotransmitter 4-aminobutanoate (GABA) (R = 0.27, p = 0.013) and *Veillonella* counts were significantly different between groups A and C2 (p =0.0475), C1 and C2 (p=0.029), and C2 and C2Bailed (p=0.042) using the Wilcoxon paired test (Fig. 6C). Also, *Veillonella* counts were not significantly different between samples of infants that received antibiotics, i.e. A and C1 (p=0.57) or C1 and C2Bailed (p=0.17). GABA peak height responses followed similar trends as *Veillonella* counts, that is, responses were significantly different between groups that received and did not receive antibiotics (A vs. C2, C1 vs. C2, C2 vs. C2Bailed) but not between groups that both received antibiotics (A vs. C1, C1 vs. C2Bailed) (Fig. 6B).

**Figure 6.**
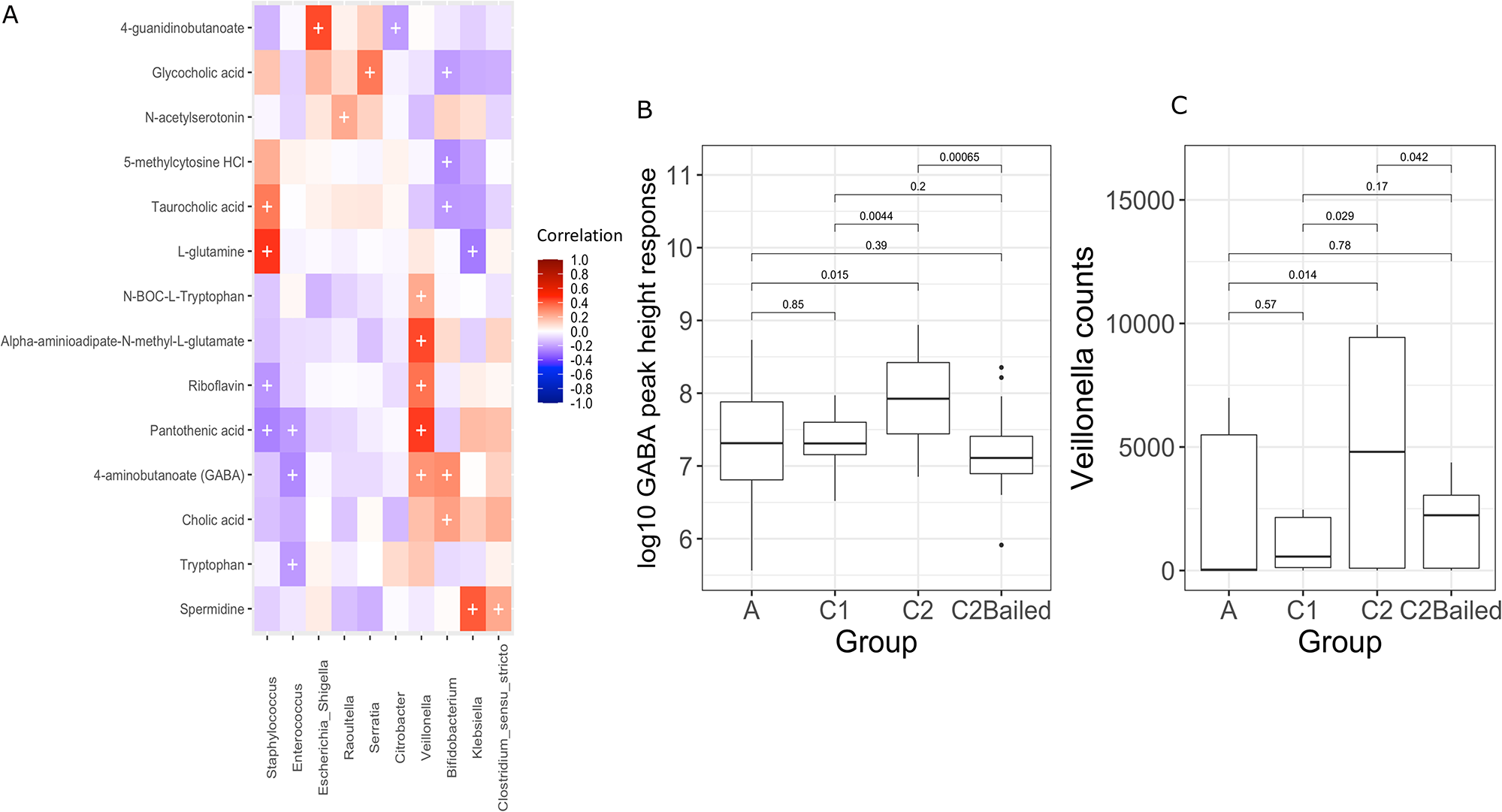
Metabolites in stool correlate with abundance of bacterial genera. **(A)** Heatmap of repeated measures correlation coefficients between peak response heights of identified metabolites in stool and the top 10 bacterial genera from the same samples (n=90 stool samples). Significant correlations are indicated by a ‘+’ with FDR-corrected p-values < 0.05. **(B)** Boxplot comparing the peak response heights for 4-aminobutyric acid (GABA) between enrollment groups. Statistical comparisons were made using the Wilcoxon test. **(C)** Boxplot comparing the number of rarefied *Veillonella* counts between the enrollment groups. Statistical comparisons were made using the Wilcoxon test. A summary of the number of infants and samples by group for metabolomics is given in Table 1.

Furthermore, using PICRUSt2, functional pathway abundances were inferred based on the rarefied 16S rRNA data29. The *Veillonella* counts of predicted pathways were strongly correlated with biosynthesis of the GABA precursor L-glutamate (R = 0.88, p = 3.02E-27) (Supplementary Figure S2). Thus, it may be that *Veillonella* could be at least partially responsible for GABA neurotransmitter production and that this function is negatively impacted by antibiotic use early in life. Alternatively, *Veillonella* may instead be involved in biosynthesis and export of L-glutamate in the gut, which is then converted to GABA by the host glutamate decarboxylase. However, PICRUSt2 results are based on inferred pathways from reference genomes closely related to the 16S rRNA data used here and are, at best, predictions in the absence of functional data specific to this cohort.

A negative correlation between *Bifidobacterium* counts and glycocholic acid was observed (R = −0.39, p = 0.0098), which was also impacted by antibiotic use between groups. In addition, bifidobacteria were negatively associated with other conjugated bile acids including taurocholic (R = −0.22, p = 0.045) and glycocholic acids (R = −0.21, p = 0.048), but positively associated with deconjugated cholic (R = 0.25, p = 0.027) (Fig. 6A). Thus, gut microbiota affected by antibiotic use may be responsible for modification of neuroactive metabolites (i.e. deconjugated bile salts) in addition to production of neurotransmitters.

### Immune markers in stool correlate with bacterial abundance

Antibiotic use was examined for its correlation with inflammatory marker levels in stool. These levels were also correlated with gut bacterial abundances. Twelve immune markers were measured in 110 stool samples across 18 of the first enrolled infants. A summary of immune marker data samples including infants per group and number of samples per infant is given in Table 1. Ten bacterial genera had at least one significant correlation with an immune marker (p < 0.05) (Fig. 7A). Significant correlations between the bacterial genera and stool immune markers were classified as either inflammatory or anti-inflammatory based on the known function of the marker (Fig. 7B). Interestingly, *Enterococcus* counts were negatively correlated with levels of TNF-alpha and macrophage inflammatory protein 1-alpha (MIP1α). *Citrobacter* were positively correlated with MIP1α and IL6 (R = 0.21, p = 3.74E-05), and were significantly higher in group C1 compared to group C2 (p=7.7E-07) and group C2 compared to C2Bailed (p=0.00022) by the Wilcoxon test (Fig. 7C). Lastly, counts of *Escherichia*/*Shigella* were significantly negatively correlated with levels of epidermal growth factor (EGF), which was the strongest correlation within the dataset. *Escherichia*/*Shigella* counts were highest among group A samples, but not significantly higher compared to other groups (Wilcoxon, p>0.05).

**Figure 7.**
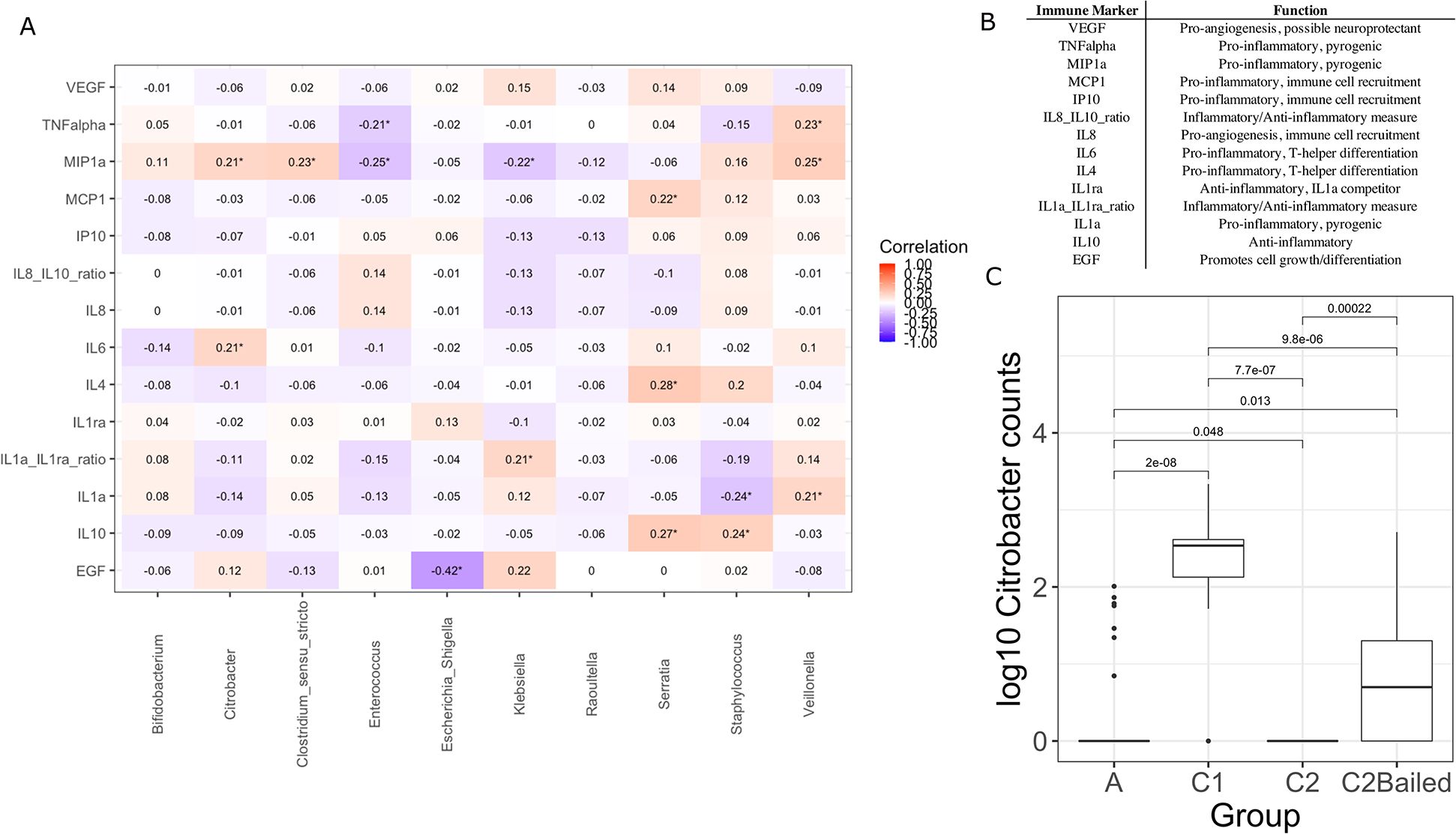
Stool immune marker levels show modest correlation with gut microbiota. **(A**) Heatmap of repeated measures correlation coefficients between immune markers measured from stool and the most abundant bacterial genera from the same samples (n=110 stool samples). Only the bacterial genera with at least one significant correlation with an immune marker are displayed (10 genera). Significant correlations are marked with an ‘*’ by the coefficient, with FDR-adjusted p-values <0.05. **(B)** Table listing the immune markers used for correlation analysis and their commonly known general functions. **(C)** Comparison of log_10_-transformed number of *Citrobacter* counts by enrollment group and their significance by the Wilcoxon test. A summary of the number of infants and samples by group for immune marker analysis is given in Table 1.

## Discussion

There is an urgent need for evidence supporting or refuting the widespread practice of routine antibiotic use after birth in symptomatic preterm neonates. The REASON study represents a significant step as it is the first randomized controlled trial to test the feasibility of randomizing symptomatic preterm infants to antibiotics versus no antibiotics, evaluating the effect of antibiotic treatment on the developing gut microbiome, metabolome, and inflammatory environment. Our results expand upon previous reports that early routine antibiotic use leads to alterations in the early life gut microbiome, even after discontinuation of antibiotics^17,30,31^. The results presented here suggest that antibiotic use 48 hours after birth did not tend to have a lasting effect on the development of gut microbiome diversity over time, and that the gut microbiota diversity was recoverable. However, use of antibiotics extending beyond 48 hours after birth often did have significant impacts on the microbiome over time, as evidenced in group A infants compared to the other enrollment groups. The power to detect significant associations in this study was hampered however, mainly because many of the infants randomized to not receive antibiotics were changed to antibiotic administration. Furthermore, there were few infants who were enrolled in group B (an important antibiotic-free control group) and those enrolled in group B had few samples due to short stays in the NICU. A larger multi-center randomized study is needed to validate and expand upon the extended effect of antibiotics on the developing gut microbiome.

Our results support the notion that feeding types likely also have a significant influence on gut microbiome richness and diversity, though in this case only at specific timepoints^32–34^. Exclusive or partial feeding with mother’s own milk appeared to have higher bacterial load compared to formula and NPO, though not significantly. This observation is backed by previous evidence that breast milk harbors maternal-originating bacteria, as well as nutritional components (prebiotics) that support bacterial proliferation in the intestinal tract^35,36^.

Interestingly, formula-fed infants had comparable levels of richness and diversity as mother’s milk. This supports the idea that mother’s milk drives early colonization of a limited set of dominant microbes through nutrient and antimicrobial-mediated selection^37–39^. Feeding trends over time were able to be assessed for the main feeding types such as MBM, DBM and formula however again the ability to detect meaningful results for rarer feeding types (particularly combinations of sources) and group B infants was hampered by small sample size and will require a larger cohort. Integrating detailed and personalized records of clinical and laboratory data led us to identify overlooked patterns in the data. One such peculiar pattern was that stool samples taken during administration of the anti-fungal fluconazole had lowered copies of 16S rRNA, suggesting lower bacterial load. A previous study reported that fluconazole, though not inherently bactericidal, increased the bactericidal activity of neutrophils^40^. Immune marker data were collected for 18 of the first enrolled infants, and one or more of those markers, such as calprotectin which is secreted by neutrophils, may help explain this pattern^41^. However, only 3 of the 18 infants that had immune marker data received fluconazole. Therefore more data are needed to test this hypothesis. Interestingly, counts of *Enterococcus* were negatively correlated with levels of pro-inflammatory markers such as TNFα and MIP1α, an odd finding considering Enterococci have been associated with risk for infection in preterm neonates^42,43^. On the other hand, *Citrobacter* counts were associated with increased levels of the macrophage chemokine MIP1α and counts were significantly higher in infants randomized to receive or were bailed to receive antibiotics. Increased levels of MIP1α are likely related to recruitment of intestinal macrophages leading to a heightened inflammatory environment, suggesting that antibiotic use may select for bacteria which lead to intestinal inflammation^44^. Finally, *Escherichia*/*Shigella* counts had a relatively strong negative correlation with EGF levels. Previous studies have found that reduced concentrations of maternally derived EGF in mice correlated with *E. coli* gut translocation, and that supplementation with EGF protected the gut from colonization by enteropathogenic *E. coli* in a young rabbit model^45,46^. Perhaps such factors as EGF concentration could be important in ameliorating the effect of antibiotics on pathogen colonization in the preterm gut. Further work including a larger sample size will be needed to understand how changes in the preterm infant gut caused by routine antibiotics impacts the gut inflammatory environment.

Neurological development can be impaired in infants born very prematurely compared to their full-term counterparts; a trend that extends into delayed cognitive and behavioral development through childhood^47–49^. Could routine early antibiotic use, or prolonged antibiotic use, in preterm neonates play a role in this association? Intestinal microbes produce a plethora of metabolites and bio-active compounds that can be absorbed by the host^50^. Some of these compounds have direct neurologic implications including neurotransmitters such as GABA, which is reduced in preterm infants, is critical for early brain development, and possesses immunomodulatory properties^51,52^. Antibiotic use was negatively affected the abundance of *Veillonella* and that *Veillonella* were positively correlated with GABA concentrations in the gut. Furthermore, *Veillonella* correlated strongly with the L-glutamine biosynthesis pathway, the precursor to GABA production. Aside from production of neurotransmitters, negative correlations were identified between *Bifidobacterium* abundance and concentrations of conjugated bile acids, particularly glyco- and taurocholic acid. Conjugated bile acids were also significantly different based on antibiotic use. Bifidobacteria, which were more abundant in infants that did not receive antibiotics, are known to deconjugate bile acids to primary forms including cholic acid^53,54^. Cholic acid can passively diffuse into the brain where it blocks signaling in the GABAA receptor^55^. Bifidobacteria may therefore be essential in regulating GABA signaling in the developing brain. These are significant findings, for they suggest routine antibiotic use could be disrupting processes involved in the gut-brain axis and immunomodulatory pathways critical for neonatal and future childhood development.

Evidence-based antibiotic use to prevent infection in preterm neonates is critical in preventing unnecessary treatment that may be doing more harm than good. Overuse of antibiotics can change the developmental trajectory of the infant gut microbiome during a time of critical establishment and interaction. However, antibiotics remain a critical treatment for a population at greater risk for infection, and there naturally exists a delicate balance between when antibiotics are truly necessary for treatment or not. Given the potential for extensive crosstalk between gut microbiota and the host, changes in microbiome composition could have both short- and long-term effects on outcomes and overall health and development. Future randomized studies with greater infant enrollment will be crucial in our understanding of the effects current neonatal practice has on health which will allow for the reevaluation of practices. Such trials will need to expand on the findings from this pilot study from a multi-omic standpoint to identify direct links between antibiotic-induced dysbiosis and health outcomes.

## Materials and Methods

### Experimental design, enrollment, and clinical sample and data collection

The REASON study was conducted from January 2017 - January 2019 at the University of Florida and was approved by the institutional review board (IRB201501045). This study is funded by the NIH (R21HD088005). A detailed description of the study design including enrollment, group selection, randomization, and collection of clinical samples and data including outcomes has been previously described^25^. Briefly, 98 premature infants were enrolled in the study and placed into one of three groups according to previously described criteria: group A with indication for antibiotic use, group B without indication for antibiotic use, and group C eligible for randomization to antibiotics (C1) or no antibiotics (C2) in the first 48 hours after birth. Infants not receiving antibiotics in the first 48 hours after birth (group B, C2) could be changed to receive antibiotics at any time at the medical team’s discretion. Clinical samples relevant to this analysis include weekly fecal collection starting with meconium when possible (all stored at −80°C) and results of bacterial and fungal cultures (blood, urine, sputum, and cerebrospinal fluid - when available) and laboratory measurements of CRP, white blood cell count and immature to neutrophil ratio. Clinical metadata from the mothers such as antepartum antibiotic use, type, duration, and proximity to delivery were recorded. Pertinent clinical metadata from the infants include group placement, antibiotic use status, antibiotics and antifungal use including type and duration throughout NICU course, feeding type and duration, GA at birth, sex, mode of delivery and any serious adverse events (SAEs) including NEC, late onset sepsis, spontaneous intestinal perforations, bronchopulmonary dysplasia, and death.

### Stool DNA extraction, 16S rRNA PCR and Sequencing Analysis

DNA extraction and 16S rRNA barcoded PCR was carried out exactly as described previously^56^. Approximately 60 gigabases of nucleotide sequencing data was generated across 5 Illumina Miseq flowcells for stool samples collected from 91 (of the 98 total) study participants where samples were collected (ICBR, Gainesville, FL, USA). The resulting sequencing reads were merged, demultiplexed, trimmed, filtered for quality and processed into amplicon sequencing variants (ASVs) as previously described with no alterations in method^56^. Briefly, sequences were processed to ASVs using the DADA2 package in R (https://www.R-project.org) and assigned taxonomy using the SILVA_v132 training dataset^57–61^. Samples were rarefied to 10,000 reads per sample, leaving 642 of the total 656 individual longitudinal stool samples for analysis.

### Total bacterial quantification by universal 16S qPCR

Total bacterial load per gram of stool was determined by universal 16S rRNA qPCR using the same primer set used for amplicon sequencing (341F and 806R). QPCR assays were performed on a QuantStudio 3 system (Applied Biosystems, Life Technologies, USA). The reaction mixture contained 12.5 μl PowerUp SYBR Green 2X Master Mix (Applied Biosystems), 1 μl each of forward (341F) and reverse (806R) primer (10 μM), 1 μl of DNA template, 0.1 μg/μl BSA and brought to a final volume of 25 μl with nuclease free water. Cycling conditions were identical to those of the endpoint PCR used for sequencing. However, with a total of 40 cycles and replacing the final elongation step with a melt curve. Each sample reaction was performed in triplicate and these values were averaged for each sample copy calculation. A standard curve was generated for copy quantification using known concentrations of the expected PCR product amplified from a similar stool sample. Copies of 16S rRNA per gram of stool was calculated by multiplying the average copy number per replicate reaction (i.e. 1 μl DNA template) by the total DNA extraction volume (75 μl) and dividing this value by the mass of stool extracted in grams.

### Absolute bacterial abundance by copy number correction

Absolute bacterial abundance was calculated on a per gram of stool basis by correcting the relative sequencing abundance by the variable number of copies of the 16S rRNA gene in each observed organism. This correction was done using the “Estimate” tool provided as part of the rrnDB copy number database^62^. Briefly, after rarefying each sample to an even sequencing depth, the ASV sequences were submitted through the rrnDB online portal where they were classified down to the genus level using the RDP classifier version 2.12 and copy number adjusted using rrnDB copy number data version 5.6^62,63^. The copy number adjusted relative abundance for each observed taxon was multiplied by the total number of 16S rRNA copies obtained by qPCR, resulting in the absolute abundance of each taxon per gram of stool.

### Fecal inflammatory markers

Inflammatory markers were analyzed using a combination of multiplex technologies using the Bio-Rad Bio-Plex platform (Bio-Rad, California, USA). The markers evaluated include common markers of intestinal inflammation including calprotectin and S100A12, in addition to other markers such as IL-6, TNF, IL-10 and other cytokines and chemokines that may play a role in inflammatory or anti-inflammatory processes. The data were analyzed using direct comparisons of all infant groups using ANOVA and subsequent individual comparisons. Fecal calprotectin and S100A12 levels were measured using the fCal ELISA kit from BUHLMANN Laboratories AG (Schonenbuch, Switzerland) and the Inflamark S100A12 kit from Cisbio Bioassays (Codolet, France), respectively, according to the manufacturer’s instructions. Samples were then analyzed for the presence of both pro-inflammatory and anti-inflammatory cytokines/chemokines using Multiplex Human Cytokine Magnetic kit, Milliplex MAP Kit (Millipore, Billerica, MA, USA). Twelve cytokines/chemokines, including EGF, IL-10, IL-1RA, IL-B, IL-4, IL-6, IL-8, IP-10, MCP-1, MIP-1a, TNFα, and VEGF were analyzed according to the manufacturer’s instructions. Plates were read using the MILLIPLEX Analyzer 3.1 xPONENT™ System (Luminex 200). Cytokine concentrations were determined using BeadView software (Millipore, Billerica, MA, USA).

### Metabolomics

The infant stool samples were suspended in 400 μl 5 mM ammonium acetate. Homogenization was done 3 times for 30 seconds each time using a cell disruptor. Protein concentrations of the homogenates were measured using Qubit Protein Assay. Samples were normalized to 500 μg/ml protein at 25 μl for extraction. Each normalized sample was spiked with 5 μl of internal standards solution. Extraction of metabolites was performed by protein precipitation by adding 200 μl of extraction solution consisting of 8:1:1 acetonitrile: methanol: acetone to each sample. Samples were mixed thoroughly, incubated at 4°C to allow protein precipitation, and centrifuged at 20,000 x g to pellet the protein. 190 μl supernatant was transferred into clean tube and dried using nitrogen. Samples were reconstituted with 25 μl of reconstitution solution consisting of injection standards, mixed, and incubated at 4° C for 10-15 min. Samples were centrifuged at 20000 x g. Supernatants were transferred into LC-vials. Global metabolomics profiling was performed as previously described using a Thermo Q-Exactive Orbitrap mass spectrometer with Dionex UHPLC and autosampler^64^. Briefly, samples were analyzed in positive and negative heated electrospray ionization with a mass resolution of 35,000 at m/z 200 as separate injections. Separation was achieved on an ACE 18-pfp 100 × 2.1 mm, 2 μm column with mobile phase A as 0.1% formic acid in water and mobile phase B as acetonitrile. The flow rate was 350 μl/min with a column temperature of 25°C. 4 μl was injected for negative ions and 2 μl for positive ions.

Data from positive and negative ions modes were processed separately. LC-MS files were first converted to open-format files (i.e. mzXML) using MSConvert from Proteowizard^65^. Mzmine was used to identify features, deisotope, align features and perform gap filling to fill in any features that may have been missed in the first alignment algorithm^66^. Features were matched with SECIM internal compound database to identify metabolites. All adducts and complexes were identified and removed from the data set prior to statistical analysis.

### Statistical Analysis

The ASV and taxonomy tables resulting from DADA2 were manipulated using the phyloseq R package v1.30.0^67^. Inferred metabolic pathway abundances were determined from the rarefied 16S rRNA data using PICRUSt2^29^. Alpha diversity measures, including the observed number of ASVs and the Shannon diversity index were calculated using the microbiome R package v1.8.0 (https://bioconductor.org/packages/devel/bioc/html/microbiome.html). Box plots (including statistical testing where applicable) were generated using the ggpubr R package v0.2.4 (https://github.com/kassambara/ggpubr), which serves as a wrapper for ggplot2^68^. The linear mixed-effects modeling and associated plots were done using the longitudinal plugin “q2-longitudinal” offered in Qiime2 v2019.4^26–28^. The biomformat R package (https://biom-format.org) was used to convert data in phyloseq format to BIOM format for import into Qiime2^69^. Bray-Curtis and Jaccard distance dissimilarities were calculated using the vegan R package v2.5.6 (https://github.com/vegandevs/vegan) and PCoA plots were made using ggplot2 v3.3.0^68^. Individual infant charts were also generated using ggplot2. Non-parametric statistical tests including the Wilcoxon and Kruskal-Wallis tests were used for pairwise and overall comparisons of 3 or more factors, respectively^70,71^. The permutational analysis of variance (PERMANOVA) test was used in the vegan package to compare overall microbiome dissimilarities between antibiotic use, feeding type, and enrollment groups. P-values were adjusted for false discovery rate (FDR) via the Benjamin-Hochberg method^72^. Repeated measures correlation values (for non-independent repeated samples for multiple subjects) were calculated using the rmcorr R package^73^.

## Data availability

The demultiplexed 16S rRNA sequencing data generated in this study is deposited in the NCBI Sequence Read Archive (SRA) under BioProject PRJNA515272.

## Trial Registration

This project is registered at clinicaltrials.gov under the name “Antibiotic ‘Dysbiosis’ in Preterm Infants” with trial number NCT02784821.

## Acknowledgements

This study was funded by the NIH under grant R21HD088005. The funder did not have a role in the design of the study nor how the study was carried out including sample collection, data analysis and interpretation or manuscript submission. We gratefully acknowledge the University of Florida Data Safety Monitoring Board; Susmita Datta, PhD - Biostatistician and Epidemiologist at University of Florida; Michael Cotton, M.D. - Neonatologist at Duke University School of Medicine; William Benitz, M.D. - Neonatologist at Lucile Packard Children's Hospital, Stanford; and Robert Lawrence, M.D. - Pediatric Infectious Disease specialist at University of Florida

## Author contributions

J.N. designed and oversaw the study. J.L.R. aided in implementation in the NICU, review of clinical events for the cohort, preparation for the DSMB and IRB reviews, and analyzed clinical data. J.N, J.L.R., D.C., and C.B. carried out patient enrollment/consent/group allocation, provided care for the infants in the NICU during the study. J.L.R, C.B. L.P. recorded clinical data. N.L. maintained/distributed the samples and performed the stool immune marker assays. T.J.G performed the stool metabolomics assays and analysis. K.L.M. assisted with stool DNA extraction, 16S PCR/qPCR, and figure generation. J.T.R. performed the stool microbiome analysis, data integration, figure generation and wrote most of the manuscript. E.W.T. helped with manuscript writing and designed the data figures for each infant which were then prepared by K.L.M. R.A.P. and E.W.T. assisted with data interpretation. All authors reviewed the manuscript before submission.

## Ethics Declarations Competing Interests

Dr. Josef Neu is the principal investigator of a study with Infant Bacterial Therapeutics and on the Scientific Advisory Boards of Medela and Astarte. No other authors have conflicts of interest to disclose.

## References

1. Israel, E. J. Neonatal necrotizing enterocolitis, a disease of the immature intestinal mucosal barrier. Acta. Paediatr. Suppl. 396, 27–32 (1994).

2. Sadeghi, K. et al. Immaturity of Infection Control in Preterm and Term Newborns Is Associated with Impaired Toll-Like Receptor Signaling. J. Infect. Dis. 195, 296–302 (2007).

3. Hunter, C. J., Upperman, J. S., Ford, H. R. & Camerini, V. Understanding the Susceptibility of the Premature Infant to Necrotizing Enterocolitis (NEC). Pediatr. Res. 63, 117–123 (2008).

4. Basso, O. & Wilcox, A. Mortality risk among preterm babies: Immaturity vs. underlying pathology. Epidemiology 21, 521–527 (2010).

5. Tripathi, N., Cotten, C. M. & Smith, P. B. Antibiotic use and misuse in the neonatal intensive care unit. Clin. Perinatol. 39, 61–68 (2012).

6. Schulman, J. et al. Neonatal intensive care unit antibiotic use. Pediatrics 135, 826–833 (2015).

7. Greenberg, R. G. et al. Prolonged duration of early antibiotic therapy in extremely premature infants. Pediatr. Res. 85, 994–1000 (2019).

8. Puopolo, K. M., Benitz, W. E., Zaoutis, T. E., Newborn, C. on F. A. & Diseases, C. on I. Management of Neonates Born at ≤34 6/7 Weeks’ Gestation With Suspected or Proven Early-Onset Bacterial Sepsis. Pediatrics 142, 1006 (2018).

9. Fajardo, C., Alshaikh, B. & Harabor, A. Prolonged use of antibiotics after birth is associated with increased morbidity in preterm infants with negative cultures. J. Matern.-Fetal Neonatal Med. 32, 4060–4066 (2019).

10. Samuels, N., van de Graaf, R. A., de Jonge, R. C. J., Reiss, I. K. M. & Vermeulen, M. J. Risk factors for necrotizing enterocolitis in neonates: a systematic review of prognostic studies. BMC Pediatr. 17, 105 (2017).

11. Alganabi, M., Lee, C., Bindi, E., Li, B. & Pierro, A. Recent advances in understanding necrotizing enterocolitis. F1000Res. 8, 107 (2019).

12. Clark, R. H., Bloom, B. T., Spitzer, A. R. & Gerstmann, D. R. Reported Medication Use in the Neonatal Intensive Care Unit: Data From a Large National Data Set. Pediatrics 117, 1979–1987 (2006).

13. Hsieh, E. M. et al. Medication use in the neonatal intensive care unit. Am. J. Perinatol. 31, 811–821 (2014).

14. Derrien, M., Alvarez, A.-S. & Vos, W. M. de. The Gut Microbiota in the First Decade of Life. Trends Microbiol. 27, 997–1010 (2019).

15. Shao, Y. et al. Stunted microbiota and opportunistic pathogen colonization in caesarean-section birth. Nature 574, 117–121 (2019).

16. Stinson, L. F., Boyce, M. C., Payne, M. S. & Keelan, J. A. The Not-so-Sterile Womb: Evidence That the Human Fetus Is Exposed to Bacteria Prior to Birth. Front. Microbiol. 10, 1124 (2019).

17. Dardas, M. et al. The impact of postnatal antibiotics on the preterm intestinal microbiome. Pediatr. Res. 76, 150–158 (2014).

18. Gasparrini, A. J. et al. Antibiotic perturbation of the preterm infant gut microbiome and resistome. Gut Microbes 7, 443–449 (2016).

19. Sun, X., Zhuan, C., Xiao, J., Yao, E. & Chen, L. Impact of postnatal exposure to antibiotics on intestinal microbiome in preterm infants. Chinese Journal of Perinatal Medicine 21, 458–464 (2018).

20. Zou, Z.-H. et al. Prenatal and postnatal antibiotic exposure influences the gut microbiota of preterm infants in neonatal intensive care units. Ann. Clin. Microbiol. Antimicrob. 17, 9 (2018).

21. Cantey, J. B., Pyle, A. K., Wozniak, P. S., Hynan, L. S. & Sánchez, P. J. Early Antibiotic Exposure and Adverse Outcomes in Preterm, Very Low Birth Weight Infants. J. Pediatr. 203, 62–67 (2018).

22. Langdon, A., Crook, N. & Dantas, G. The effects of antibiotics on the microbiome throughout development and alternative approaches for therapeutic modulation. Genome Med. 8, 39 (2016).

23. Sherman, M. P., Zaghouani, H. & Niklas, V. Gut microbiota, the immune system, and diet influence the neonatal gut–brain axis. Pediatr. Res. 77, 127–135 (2015).

24. Collins, S. M., Surette, M. & Bercik, P. The interplay between the intestinal microbiota and the brain. Nat. Rev. Microbiol. 10, 735–742 (2012).

25. Ruoss, J. L., et al. Routine Early Antibiotic use in SymptOmatic preterm Neonates (REASON): a prospective randomized controlled trial. medRxiv 2020.04.17.20069617 (2020) doi:10.1101/2020.04.17.20069617

26. Lindstrom, M. J. & Bates, D. M. Newton—Raphson and EM Algorithms for Linear Mixed-Effects Models for Repeated-Measures Data. J. Am. Stat. Assoc. 83, 1014–1022 (1988).

27. Bokulich, N. A. et al. q2-longitudinal: Longitudinal and Paired-Sample Analyses of Microbiome Data. mSystems 3, e00219–18 (2018).

28. Bolyen, E. et al. Reproducible, interactive, scalable and extensible microbiome data science using QIIME 2. Nat. Biotechnol. 37, 852–857 (2019).

29. Douglas, G. M. et al. PICRUSt2: An improved and customizable approach for metagenome inference. bioRxiv 672295 (2020) doi:10.1101/672295.

30. Tanaka, S. et al. Influence of antibiotic exposure in the early postnatal period on the development of intestinal microbiota. FEMS Immunol. Med. Mic. 56, 80–87 (2009).

31. Greenwood, C. et al. Early Empiric Antibiotic Use in Preterm Infants Is Associated with Lower Bacterial Diversity and Higher Relative Abundance of Enterobacter. J. Pediatr. 165, 23–29 (2014).

32. Cong, X. et al. Gut Microbiome Developmental Patterns in Early Life of Preterm Infants: Impacts of Feeding and Gender. PLoS One 11, e0152751 (2016).

33. Cong, X. et al. Influence of Infant Feeding Type on Gut Microbiome Development in Hospitalized Preterm Infants. Nurs. Res. 66, 123–133 (2017).

34. Parra-Llorca, A. et al. Preterm Gut Microbiome Depending on Feeding Type: Significance of Donor Human Milk. Front. Microbiol. 9, 1376 (2018).

35. Gregory, K. E. et al. Influence of maternal breast milk ingestion on acquisition of the intestinal microbiome in preterm infants. Microbiome 4, 68 (2016).

36. Kordy, K. et al. Contributions to human breast milk microbiome and enteromammary transfer of Bifidobacterium breve. PLoS ONE 15, e0219633 (2020).

37. Pannaraj, P. S. et al. Association Between Breast Milk Bacterial Communities and Establishment and Development of the Infant Gut Microbiome. JAMA Pediatr. 171, 647–654 (2017).

38. Forbes, J. D. et al. Association of Exposure to Formula in the Hospital and Subsequent Infant Feeding Practices With Gut Microbiota and Risk of Overweight in the First Year of Life. JAMA Pediatr. 172, e181161 (2018).

39. Mueller, E. & Blaser, M. Breast milk, formula, the microbiome and overweight. Nat. Rev. Endocrinol. 14, 510–511 (2018).

40. Zervos, E. E., Bass, S. S., Robson, M. C. & Rosemurgy, A. S. Fluconazole increases bactericidal activity of neutrophils. J. Trauma 41, 10–14 (1996).

41. Rougé, C. et al. Fecal Calprotectin Excretion in Preterm Infants during the Neonatal Period. PLoS One 5, e11083 (2010).

42. Coggins, S. A., Wynn, J. L. & Weitkamp, J.-H. Infectious Causes of Necrotizing Enterocolitis. Clin. Perinatol. 42, 133–154 (2015).

43. Dobson, S. R. M. & Baker, C. J. Enterococcal Sepsis in Neonates: Features by Age at Onset and Occurrence of Focal Infection. Pediatrics 85, 165–171 (1990).

44. O’Grady, N. P. et al. Detection of Macrophage Inflammatory Protein (MIP)-1α and MIP-β during Experimental Endotoxemia and Human Sepsis. J. Infect. 179, 136–141 (1999).

45. Knoop, K. A., et al. Maternal activation of the EDFR prevents translocation of gut-residing pathogenic *Escherichia coli* in a model of late-onset neonatal sepsis. PNAS 117, 7941–7949 (2020).

46. Buret, A., Olson, M. E., Gall, D. G., & Hardin, J. A. Effects of Orally Administered Epidermal Growth Factor on Enteropathogenic *Escherichia coli* Infection in Rabbits. Infect. Immunol. 66, 4917–4923 (1998).

47. Bhutta, A. T., Cleves, M. A., Casey, P. H., Cradock, M. M. & Anand, K. J. S. Cognitive and Behavioral Outcomes of School-Aged Children Who Were Born Preterm: A Meta-analysis. JAMA 288, 728–737 (2002).

48. Counsell, S. J. & Boardman, J. P. Differential brain growth in the infant born preterm: Current knowledge and future developments from brain imaging. Semin. Fetal. Neonat. M. 10, 403–410 (2005).

49. Marlow, N. & Samara, M. Neurologic and Developmental Disability at Six Years of Age after Extremely Preterm Birth. N. Engl. J. Med. 352, 9–19 (2005).

50. Sharon, G. et al. Specialized Metabolites from the Microbiome in Health and Disease. Cell Metab. 20, 719–730 (2014).

51. Kwon, S. H. et al. GABA, Resting State Connectivity and the Developing Brain. Neonatology 106, 149–155 (2014).

52. Jin, Z., Mendu, S. K. & Birnir, B. GABA is an effective immunomodulatory molecule. Amino Acids 45, 87–94 (2013).

53. Grill, J. P., Manginot-Dürr, C., Schneider, F. & Ballongue, J. Bifidobacteria and probiotic effects: Action of Bifidobacterium species on conjugated bile salts. Curr. Microbiol. 31, 23–27 (1995).

54. Jia, W., Xie, G. & Jia, W. Bile acid–microbiota crosstalk in gastrointestinal inflammation and carcinogenesis. Nat. Rev. Gastro. Hepat. 15, 111–128 (2018).

55. Kiriyama, Y. & Nochi, H. The Biosynthesis, Signaling, and Neurological Functions of Bile Acids. Biomolecules 9, 232 (2019).

56. Russell, J. T. et al. Genetic risk for autoimmunity is associated with distinct changes in the human gut microbiome. Nat. Commun. 10, 1–12 (2019).

57. Callahan, B. J. et al. DADA2: High resolution sample inference from Illumina amplicon data. Nat. Methods. 13, 581–583 (2016).

58. Pruesse, E. et al. SILVA: a comprehensive online resource for quality checked and aligned ribosomal RNA sequence data compatible with ARB. Nucleic Acids Res. 35, 7188–7196 (2007).

59. Glöckner, F. O. et al. 25 years of serving the community with ribosomal RNA gene reference databases and tools. J. Biotechnol. 261, 169–176 (2017).

60. Yilmaz, P. et al. The SILVA and “All-species Living Tree Project (LTP)” taxonomic frameworks. Nucleic Acids Res. 42, D643–D648 (2014).

61. Quast, C. et al. The SILVA ribosomal RNA gene database project: improved data processing and web-based tools. Nucleic Acids Res. 41, D590–D596 (2013).

62. Stoddard, S. F., Smith, B. J., Hein, R., Roller, B. R. K. & Schmidt, T. M. rrnDB: improved tools for interpreting rRNA gene abundance in bacteria and archaea and a new foundation for future development. Nucleic Acids Res. 43, D593–D598 (2015).

63. Wang, Q., Garrity, G. M., Tiedje, J. M. & Cole, J. R. Naive Bayesian classifier for rapid assignment of rRNA sequences into the new bacterial taxonomy. Appl. Environ. Microbiol. 73, 5261–5267 (2007).

64. Chamberlain, C. A., Hatch, M. & Garrett, T. J. Metabolomic and lipidomic characterization of Oxalobacter formigenes strains HC1 and OxWR by UHPLC-HRMS. Anal. Bioanal. Chem. 411, 4807–4818 (2019).

65. Chambers, M. C. et al. A cross-platform toolkit for mass spectrometry and proteomics. Nat. Biotechnol. 30, 918–920 (2012).

66. Pluskal, T., Castillo, S., Villar-Briones, A. & Orešič, M. MZmine 2: Modular framework for processing, visualizing, and analyzing mass spectrometry-based molecular profile data. BMC Bioinform. 11, 395 (2010).

67. McMurdie, P. J. & Holmes, S. phyloseq: An R Package for Reproducible Interactive Analysis and Graphics of Microbiome Census Data. PLOS ONE 8, e61217 (2013).

68. Wickham, H. ggplot2: Elegant Graphics for Data Analysis. (Springer-Verlag, New York, 2009).

69. McDonald, D. et al. The Biological Observation Matrix (BIOM) Format Or: How I Learned to Stop Worrying and Love the Ome-Ome. Gigascience 1, 1–7 (2012).

70. Wilcoxon, F. Individual Comparisons by Ranking Methods. Biometrics Bulletin 1, 80–83 (1945).

71. Kruskal, W. H. & Wallis, W. A. Use of Ranks in One-Criterion Variance Analysis. J. Am. Stat. Assoc. 47, 583–621 (1952).

72. Benjamini, Y. & Hochberg, Y. Controlling the False Discovery Rate: A Practical and Powerful Approach to Multiple Testing. J. R. Statist. Soc. 57, 289–300 (1995).

73. Bakdash, J. Z. & Marusich, L. R. Repeated Measures Correlation. Front. Psychol. 8, 456 (2017).

