## Supplementary Figure S1 for "Antibiotics and the developing intestinal microbiome, metabolome and inflammatory environment: a randomized trial of preterm infants"

**Supplementary Figure S1. Integrated clinical and laboratory data charts for all infants.** Data included in each chart from top to bottom include: the infant ID, group assignment, antibiotic change status (bail), gestational age, any adverse clinical events, the type and duration of antibiotic use (if any), the copy-number corrected absolute composition of each weekly stool sample and it's log10-scale number of bacterial 16S rRNA copies, the type and duration of each feeding including administration of human milk fortifier, the relative levels of C-reactive protein measured from blood, and relative concentrations of measured stool immune markers (for infants where these measurements were performed). DBM: donor breast milk, MBM: mother's breast milk, NPO: no enteral nutrition, CRP: C-reactive protein, EGF: epidermal growth factor. Collected stool samples that did not amplify for 16S rRNA during PCR are indicated by a black dot. Samples that were collected on or close to the same day are stacked, with the first collected sample on top.

#### Bacterial Taxa

- Actinomyces
- Alloprevotella
- Asaccharobacter
- Bifidobacterium
- Chryseobacterium
- Clostridium XIV
- Deinococcus
- Enterococcus
- Finegoldia
- Fusobacterium
- Klebsiella
- Morganella
- Pantoea
- Prevotella
- Pseudomonas
- Romboutsia
- Selenomonas
- Streptococcus
- Veillonella
- Aerococcus
- Anaerococcus
- Atopobium
- Blautia
- Citrobacter
- Collinsella
- Dialister
- Escherichia/Shigella
- Franconibacter
- Gemella
- Lactobacillus
- Mycoplasma
- Parabacteroides
- Propionibacterium
- Raoultella
- Roseburia
- Serratia
- Corynebacterium
- Enterobacter
- Faecalibacterium
- Ureaplasma
- Alistipes
- Aquabacterium
- Bacteroides
- Clostridium sensu stricto
- Fusicatenibacter
- Haemophilus
- Megasphaera
- Negativicoccus
- Peptinophilus
- Proteus
- Robinsoniella
- Ruminococcus
- Staphylococcus
- Terrisporobacter
- Unclassified_Enteroboacteriaceae
- Unclassified_Ruminococcaceae
- Unclassified_Peptostreptococcaceae
- Unclassified_Coriobacteriaceae
- Unclasssifed_Bifidobacteriaceae
- Unclassified_Lachnospiraceae
- Unclassified_Veillonellaceae

#### Clinical Events

- A - Bacteremia
- B - Sepsis
- C - Fungemia
- D - NEC
- E - NEC rule out
- F - Sepsis rule out
- G - SIP
- H - Pneumonia
- I - Umbilical abscess or peritonitis
- J - Conjunctivitis
- K - Bowel stricture or resection
- L - OR central line placement
- M - Thrombophlebitis
- N - Sever IVH
- O - Death

- Candida - Candida infection *blood
- CoNS - Coagulase neg. Staph. *blood
- MRSA - MRSA infection *w=wound, s=swab, b=blood
- Entero - Enterobacteria infection *w=wound, s=swab, b=blood

- HMF - Human milk fortifier

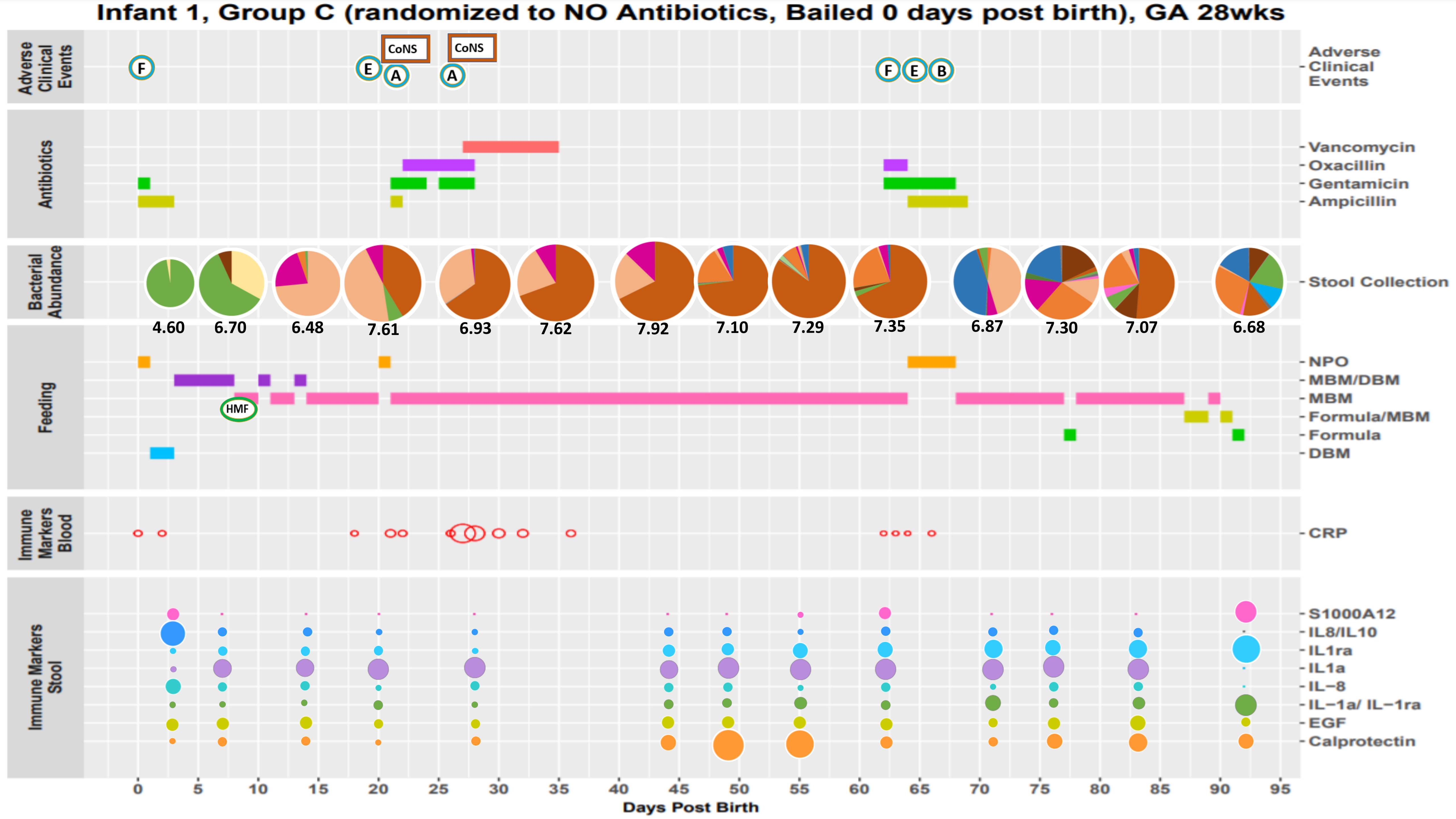

Infant 1, Group C (randomized to NO Antibiotics, Bailed 0 days post birth), GA 28wks
Adverse Clinical Events
F
E
A
CoNS
CoNS
A
F
E
B
Antibiotics
Vancomycin
Oxacillin
Gentamicin
Ampicillin
Bacterial Abundance
Stool Collection
4.60
6.70
6.48
7.61
6.93
7.62
7.92
7.10
7.29
7.35
6.87
7.30
7.07
6.68
Feeding
HMF
NPO
MBM/DBM
MBM
Formula/MBM
Formula
DBM
Immune Markers Blood
CRP
Immune Markers Stool
S1000A12
IL8/IL10
IL1ra
IL1a
IL−8
IL−1a/ IL−1ra
EGF
Calprotectin
0
5
10
15
20
25
30
35
40
45
50
55
60
65
70
75
80
85
90
95
Days Post Birth

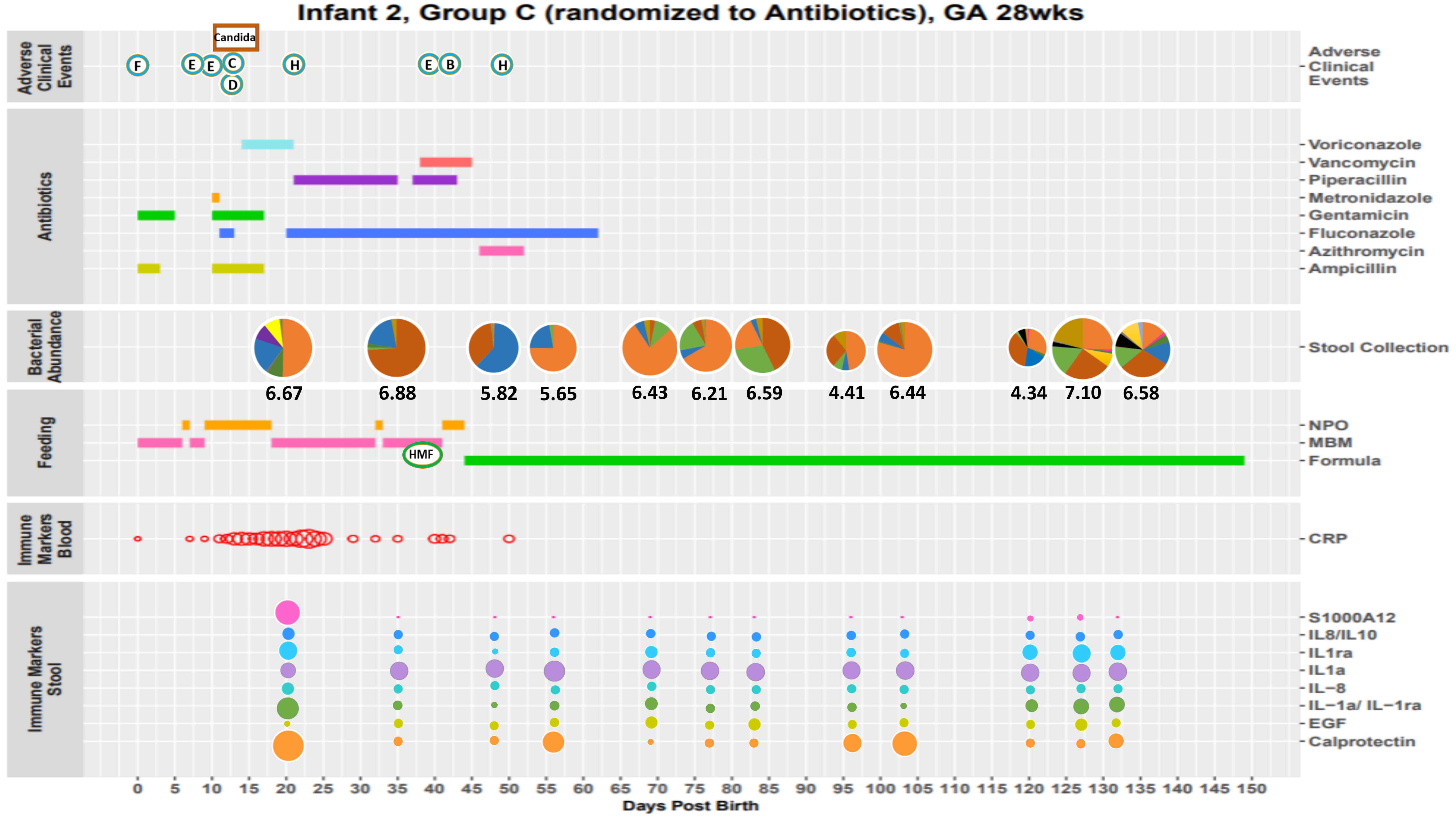

Infant 2, Group C (randomized to Antibiotics), GA 28wks
Adverse Clinical Events
Candida
F
E
E
C
D
H
E
B
H
Antibiotics
Voriconazole
Vancomycin
Piperacillin
Metronidazole
Gentamicin
Fluconazole
Azithromycin
Ampicillin
Bacterial Abundance
Stool Collection
6.67
6.88
5.82
5.65
6.43
6.21
6.59
4.41
6.44
4.34
7.10
6.58
Feeding
NPO
MBM
Formula
HMF
Immune Markers Blood
CRP
Immune Markers Stool
S1000A12
IL8/IL10
IL1ra
IL1a
IL−8
IL−1a/ IL−1ra
EGF
Calprotectin
0 5 10 15 20 25 30 35 40 45 50 55 60 65 70 75 80 85 90 95 100 105 110 115 120 125 130 135 140 145 150
Days Post Birth

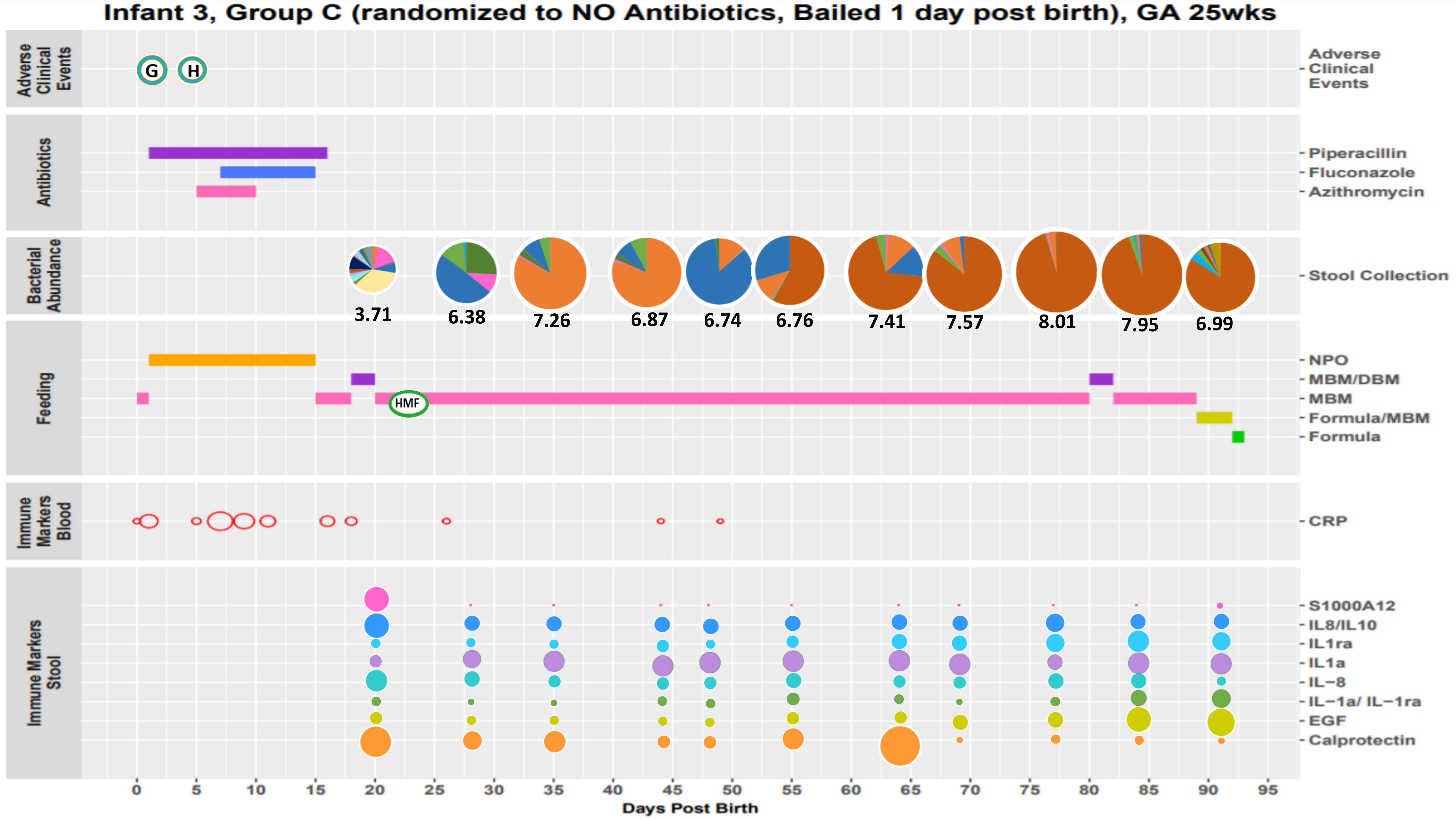

Infant 3, Group C (randomized to NO Antibiotics, Bailed 1 day post birth), GA 25wks
Adverse Clinical Events
G
H
Antibiotics
Piperacillin
Fluconazole
Azithromycin
Bacterial Abundance
Stool Collection
3.71
6.38
7.26
6.87
6.74
6.76
7.41
7.57
8.01
7.95
6.99
Feeding
NPO
MBM/DBM
MBM
Formula/MBM
Formula
HMF
Immune Markers Blood
CRP
Immune Markers Stool
S1000A12
IL8/IL10
IL1ra
IL1a
IL−8
IL−1a/ IL−1ra
EGF
Calprotectin
0
5
10
15
20
25
30
35
40
45
50
55
60
65
70
75
80
85
90
95
Days Post Birth

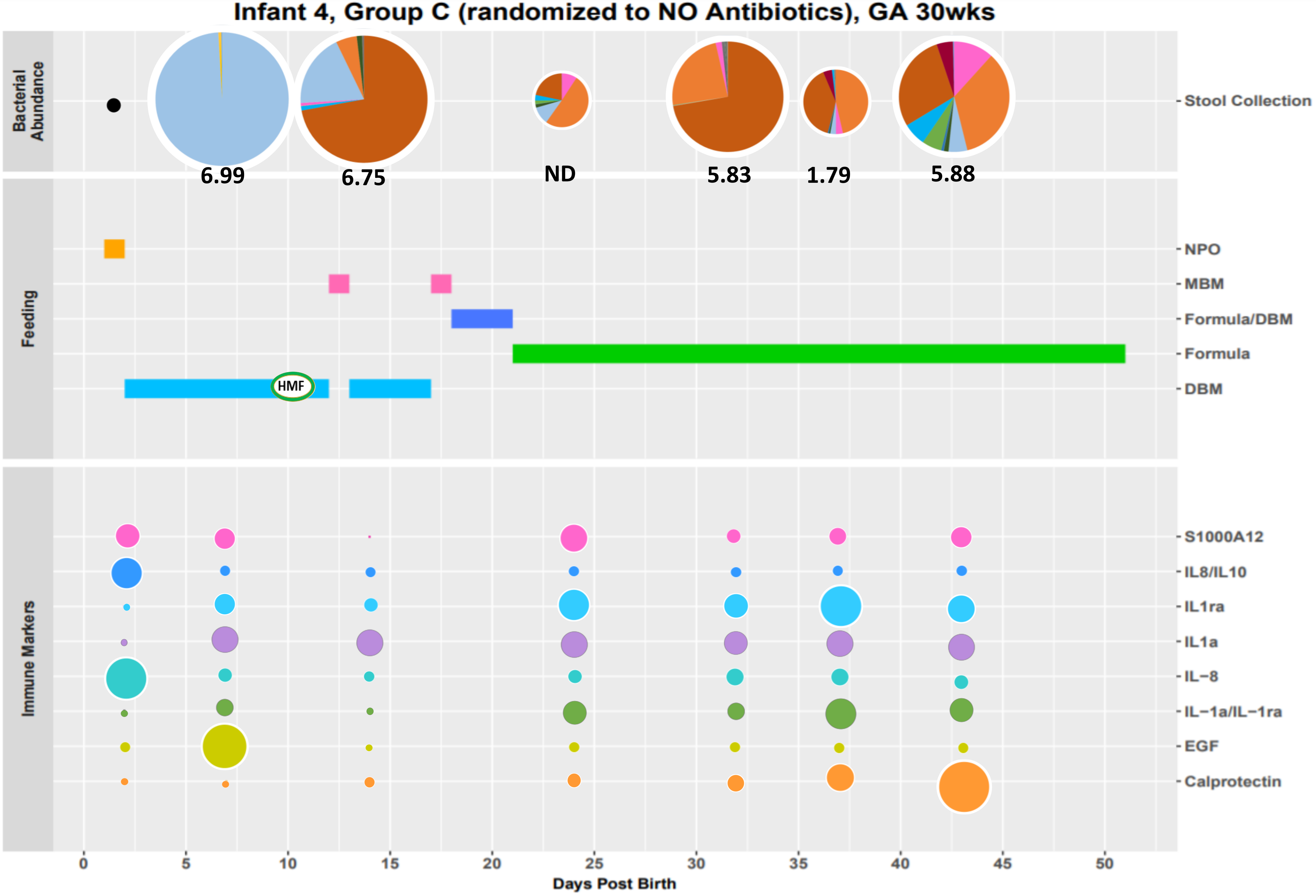

Infant 4, Group C (randomized to NO Antibiotics), GA 30wks
Bacterial Abundance
Stool Collection
6.99
6.75
ND
5.83
1.79
5.88
Feeding
NPO
MBM
Formula/DBM
Formula
DBM
HMF
Immune Markers
S1000A12
IL8/IL10
IL1ra
IL1a
IL−8
IL−1a/IL−1ra
EGF
Calprotectin
0
5
10
15
20
25
30
35
40
45
50
Days Post Birth

Infant 5, Group A (requires Antibiotics), GA 29wks

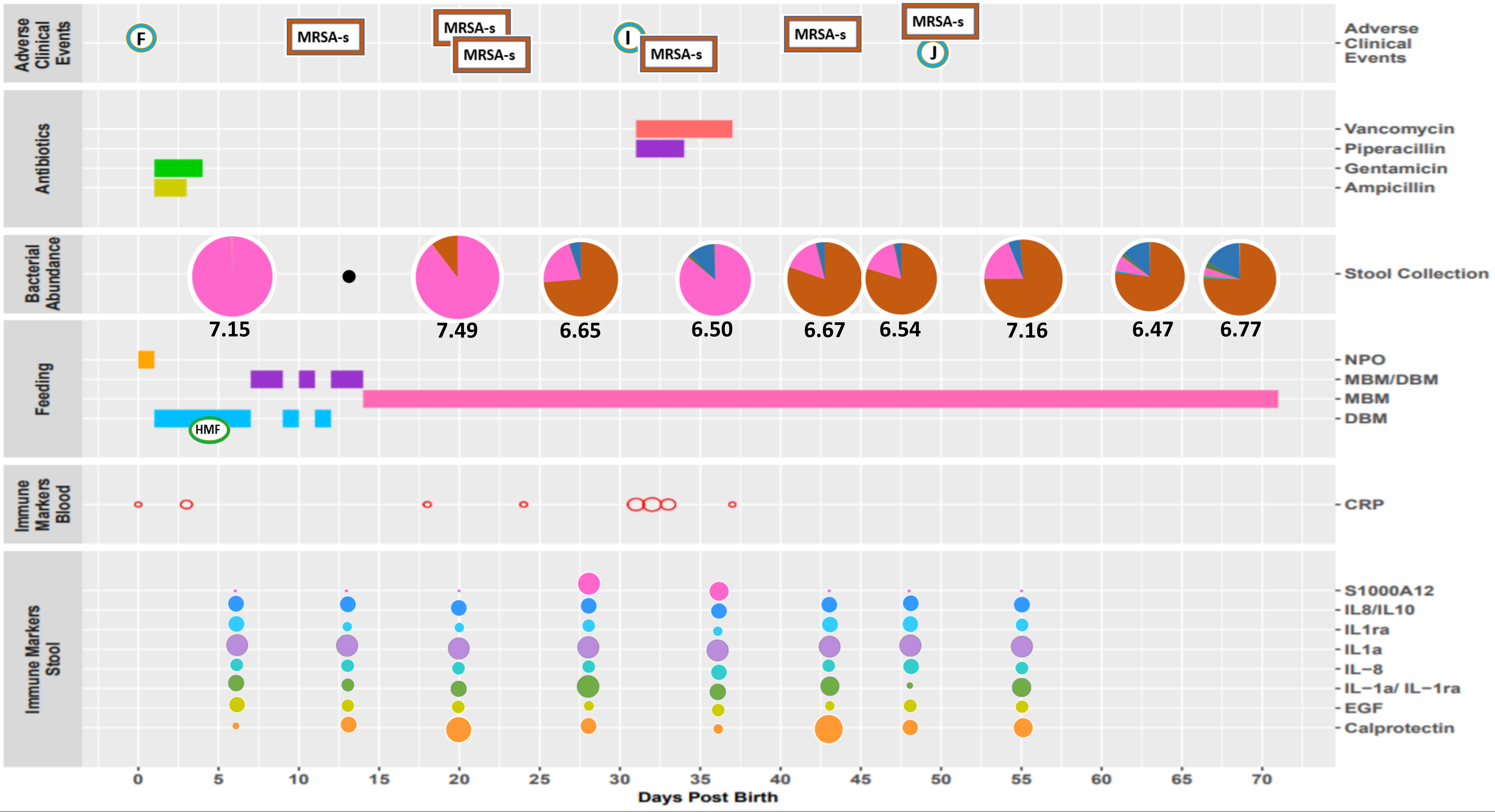

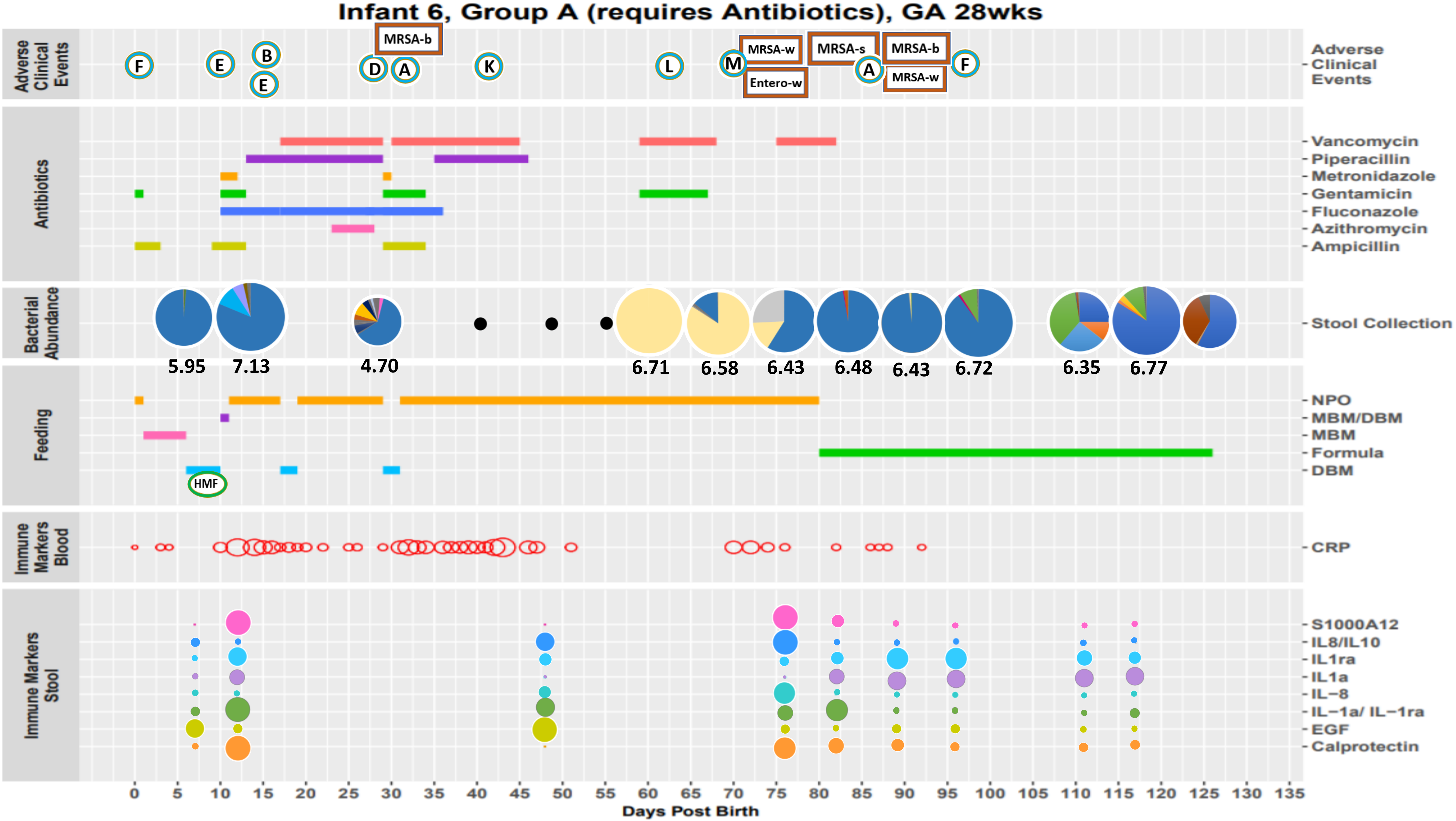
Infant 6, Group A (requires Antibiotics), GA 28wks
Adverse Clinical Events
F
E
B
E
D
A
MRSA-b
K
L
M
MRSA-w
Entero-w
MRSA-s
A
MRSA-b
MRSA-w
F
Antibiotics
Vancomycin
Piperacillin
Metronidazole
Gentamicin
Fluconazole
Azithromycin
Ampicillin
Bacterial Abundance
Stool Collection
5.95
7.13
4.70
6.71
6.58
6.43
6.48
6.43
6.72
6.35
6.77
Feeding
NPO
MBM/DBM
MBM
Formula
DBM
HMF
Immune Markers Blood
CRP
Immune Markers Stool
S1000A12
IL8/IL10
IL1ra
IL1a
IL−8
IL−1a/ IL−1ra
EGF
Calprotectin
0
5
10
15
20
25
30
35
40
45
50
55
60
65
70
75
80
85
90
95
100
105
110
115
120
125
130
135
Days Post Birth

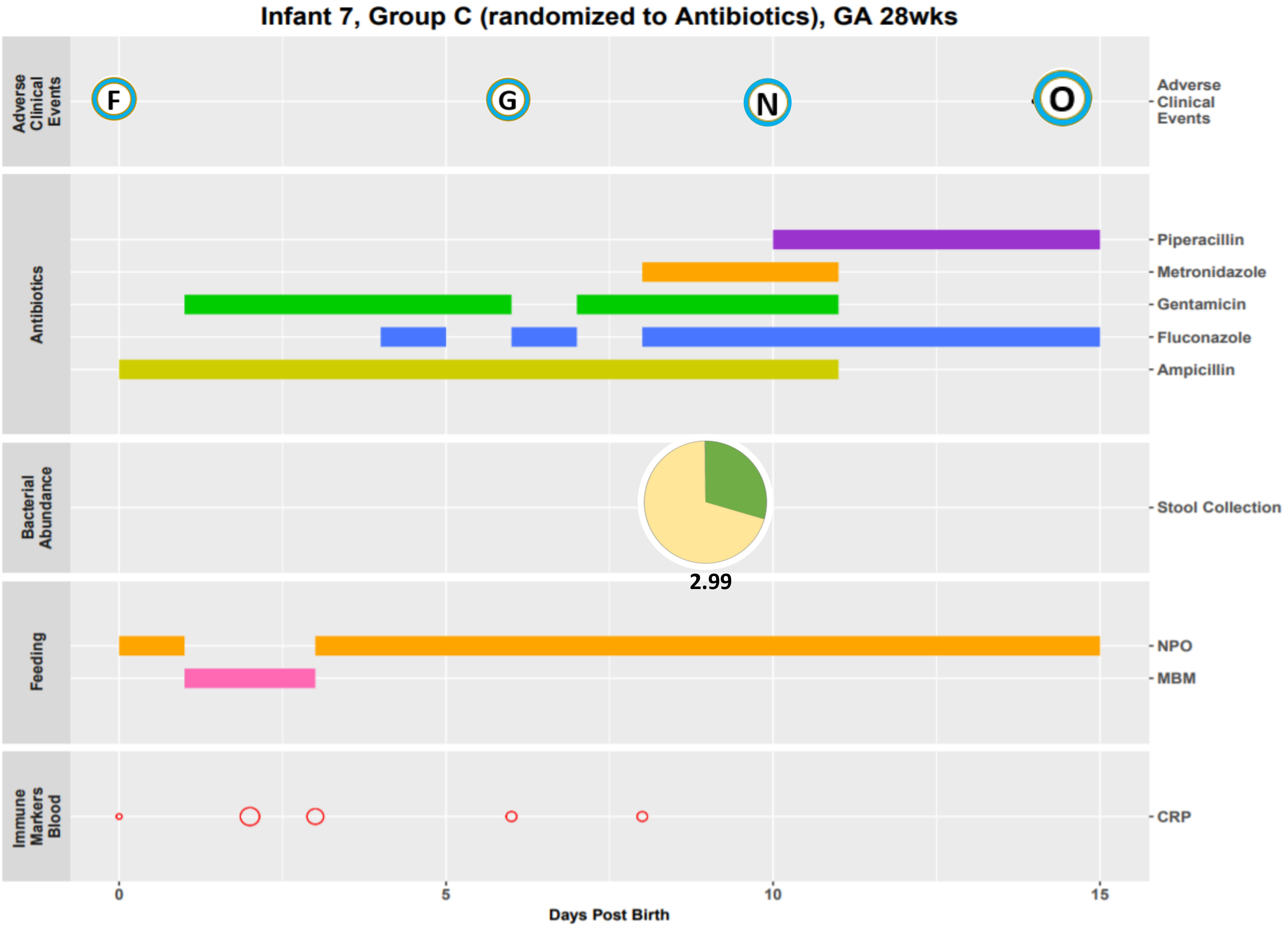
Infant 7, Group C (randomized to Antibiotics), GA 28wks
Adverse Clinical Events
F
G
N
O
Antibiotics
Piperacillin
Metronidazole
Gentamicin
Fluconazole
Ampicillin
Bacterial Abundance
Stool Collection
2.99
Feeding
NPO
MBM
Immune Markers Blood
CRP
0
5
10
15
Days Post Birth

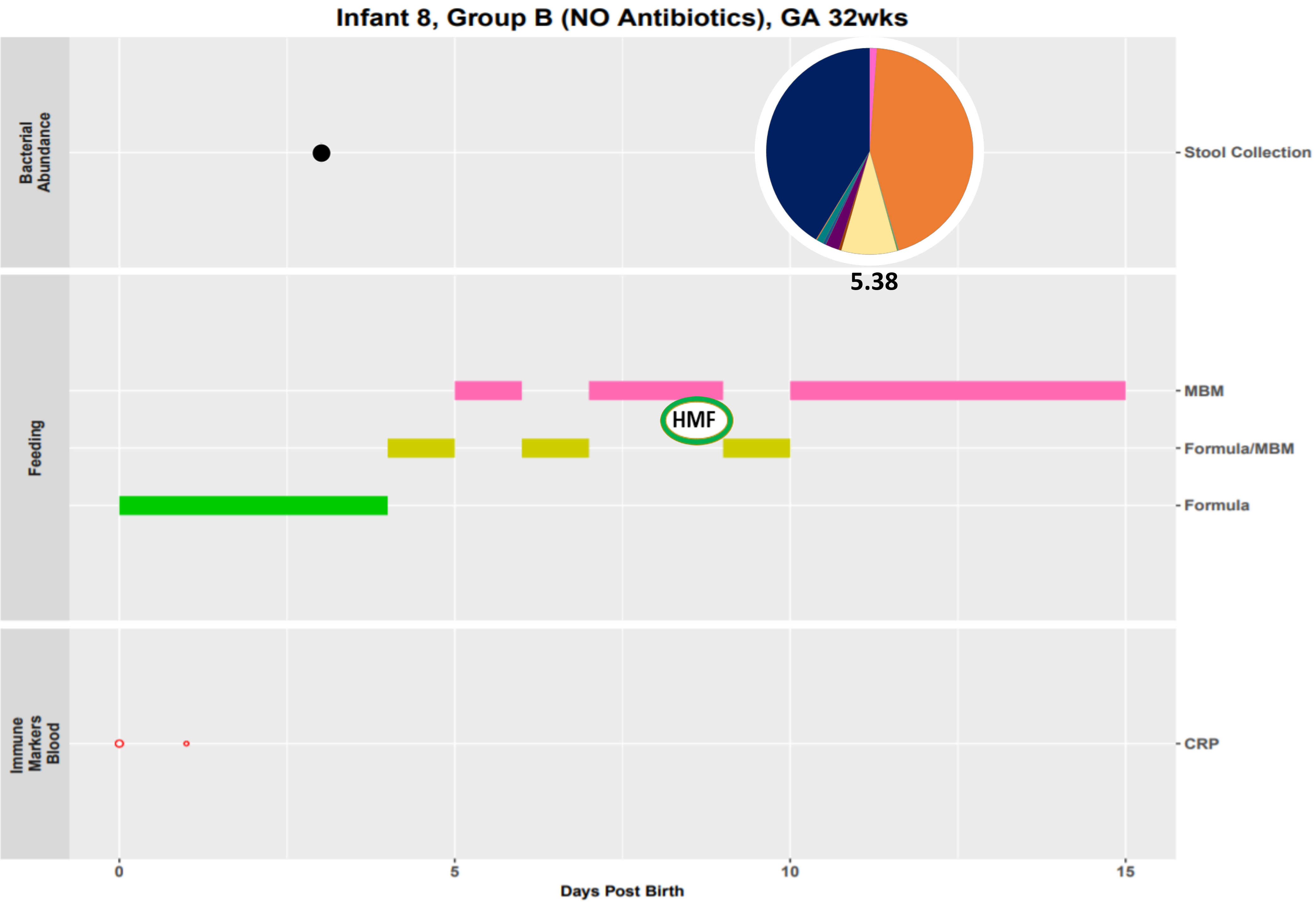

Infant 8, Group B (NO Antibiotics), GA 32wks
Bacterial Abundance
Feeding
Immune Markers Blood
Stool Collection
MBM
Formula/MBM
Formula
CRP
HMF
5.38
0
5
10
15
Days Post Birth

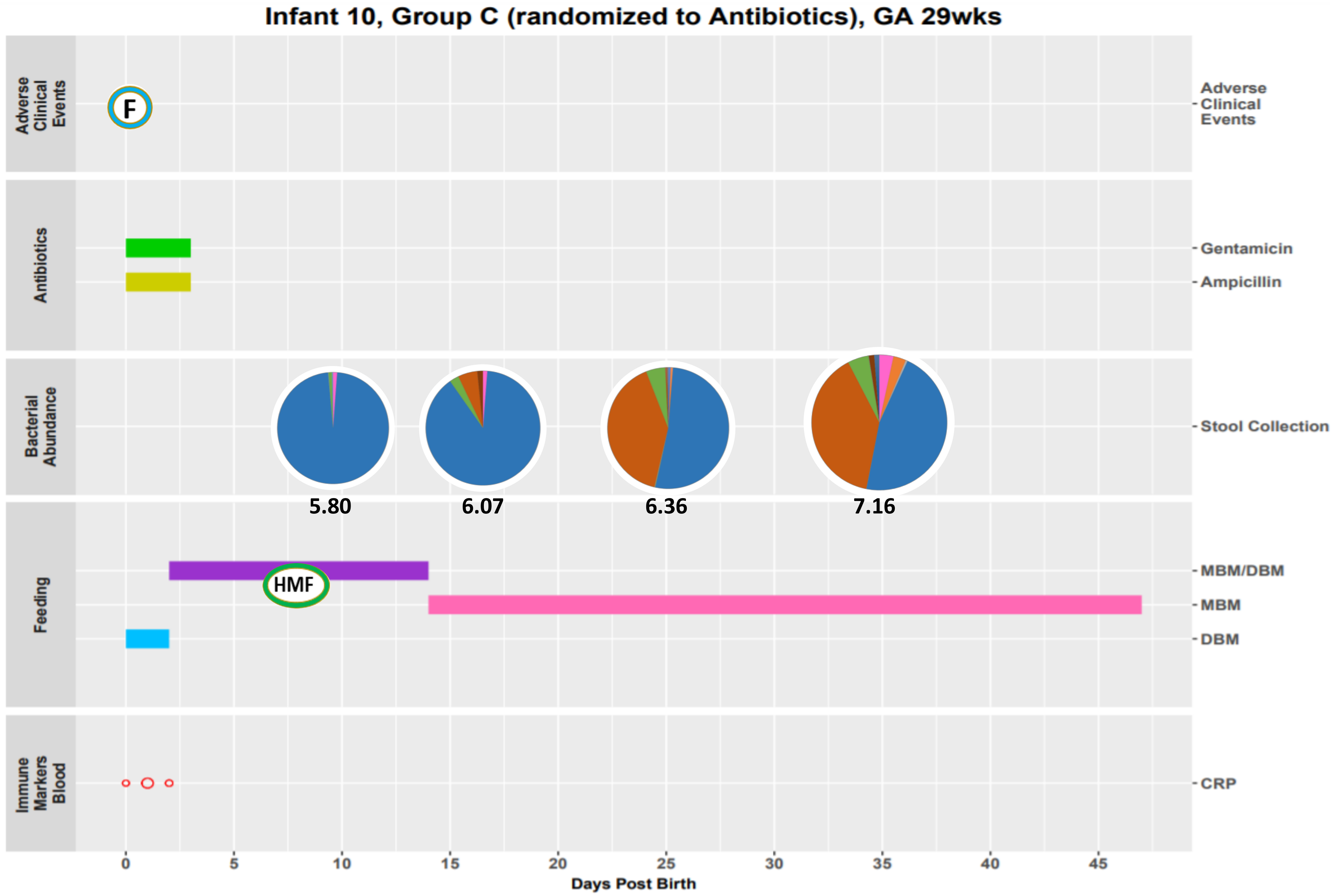
Infant 10, Group C (randomized to Antibiotics), GA 29wks
Adverse Clinical Events
F
Antibiotics
Gentamicin
Ampicillin
Bacterial Abundance
Stool Collection
5.80
6.07
6.36
7.16
Feeding
MBM/DBM
HMF
MBM
DBM
Immune Markers Blood
CRP
0
5
10
15
20
25
30
35
40
45
Days Post Birth

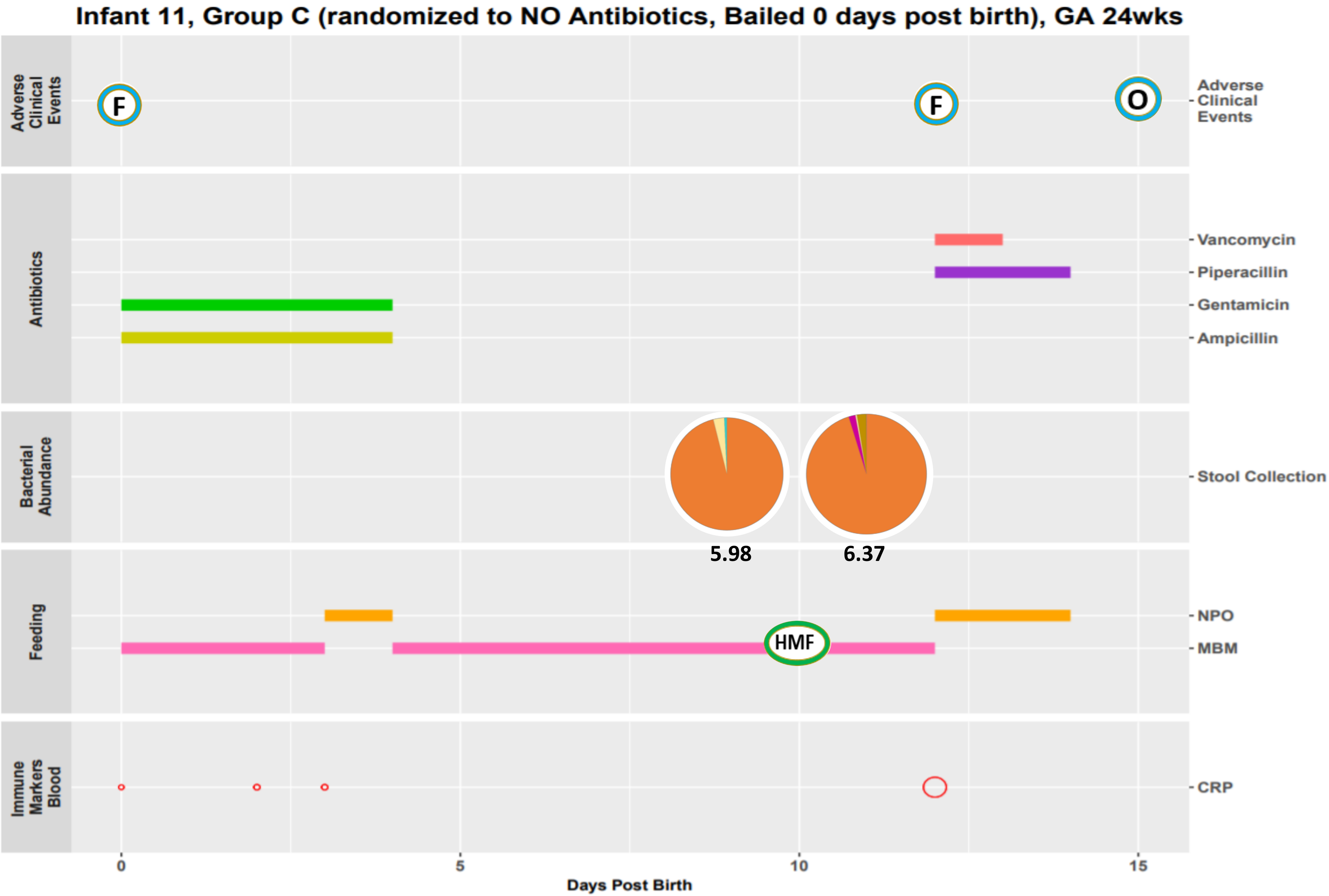
Infant 11, Group C (randomized to NO Antibiotics, Bailed 0 days post birth), GA 24wks
Adverse Clinical Events
F
F
O
Antibiotics
Vancomycin
Piperacillin
Gentamicin
Ampicillin
Bacterial Abundance
Stool Collection
5.98
6.37
Feeding
NPO
MBM
HMF
Immune Markers Blood
CRP
0
5
10
15
Days Post Birth

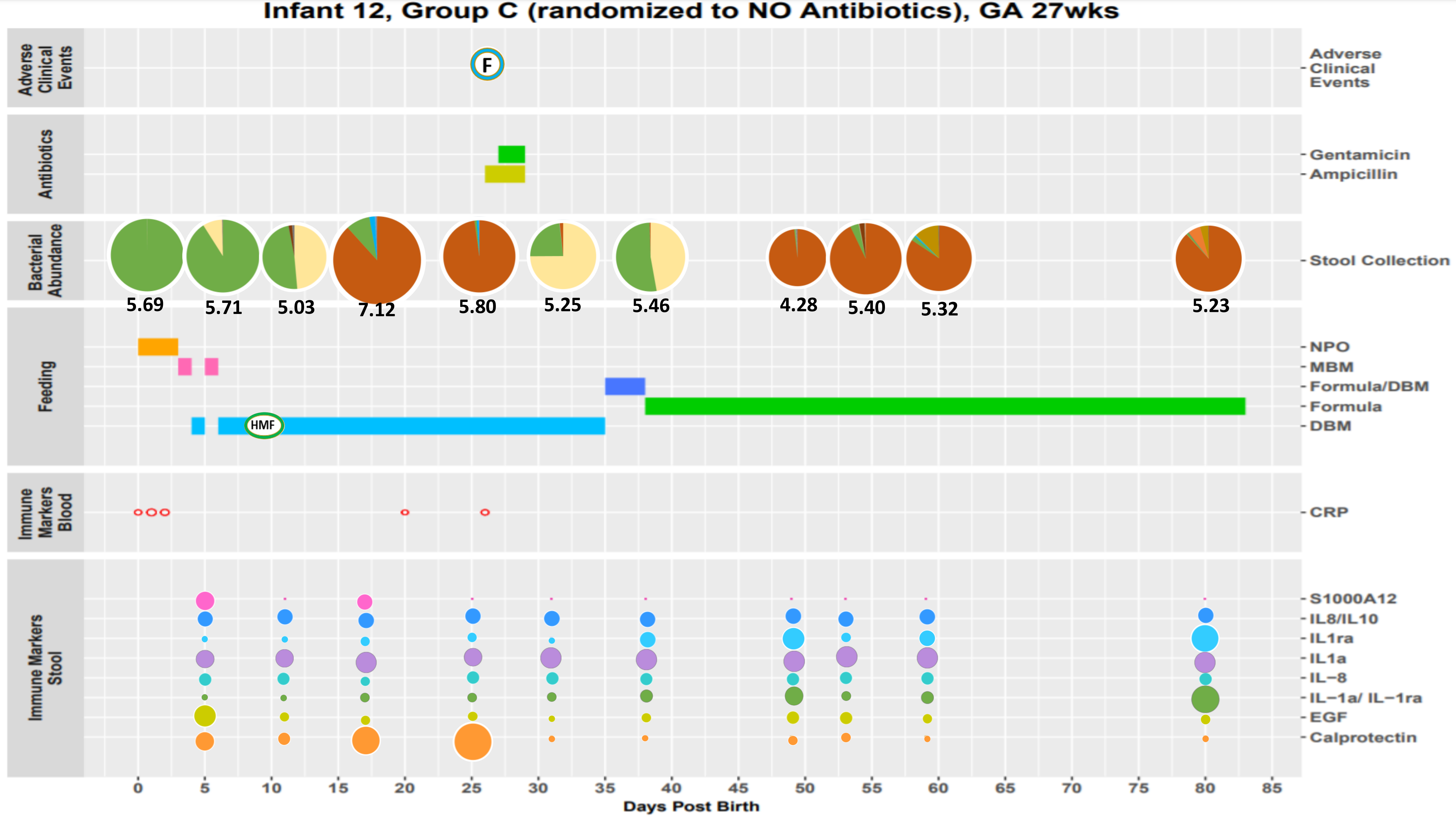
Infant 12, Group C (randomized to NO Antibiotics), GA 27wks
Adverse Clinical Events
F
Antibiotics
Gentamicin
Ampicillin
Bacterial Abundance
Stool Collection
5.69
5.71
5.03
7.12
5.80
5.25
5.46
4.28
5.40
5.32
5.23
Feeding
NPO
MBM
Formula/DBM
Formula
DBM
HMF
Immune Markers Blood
CRP
Immune Markers Stool
S1000A12
IL8/IL10
IL1ra
IL1a
IL−8
IL−1a/ IL−1ra
EGF
Calprotectin
0 5 10 15 20 25 30 35 40 45 50 55 60 65 70 75 80 85
Days Post Birth

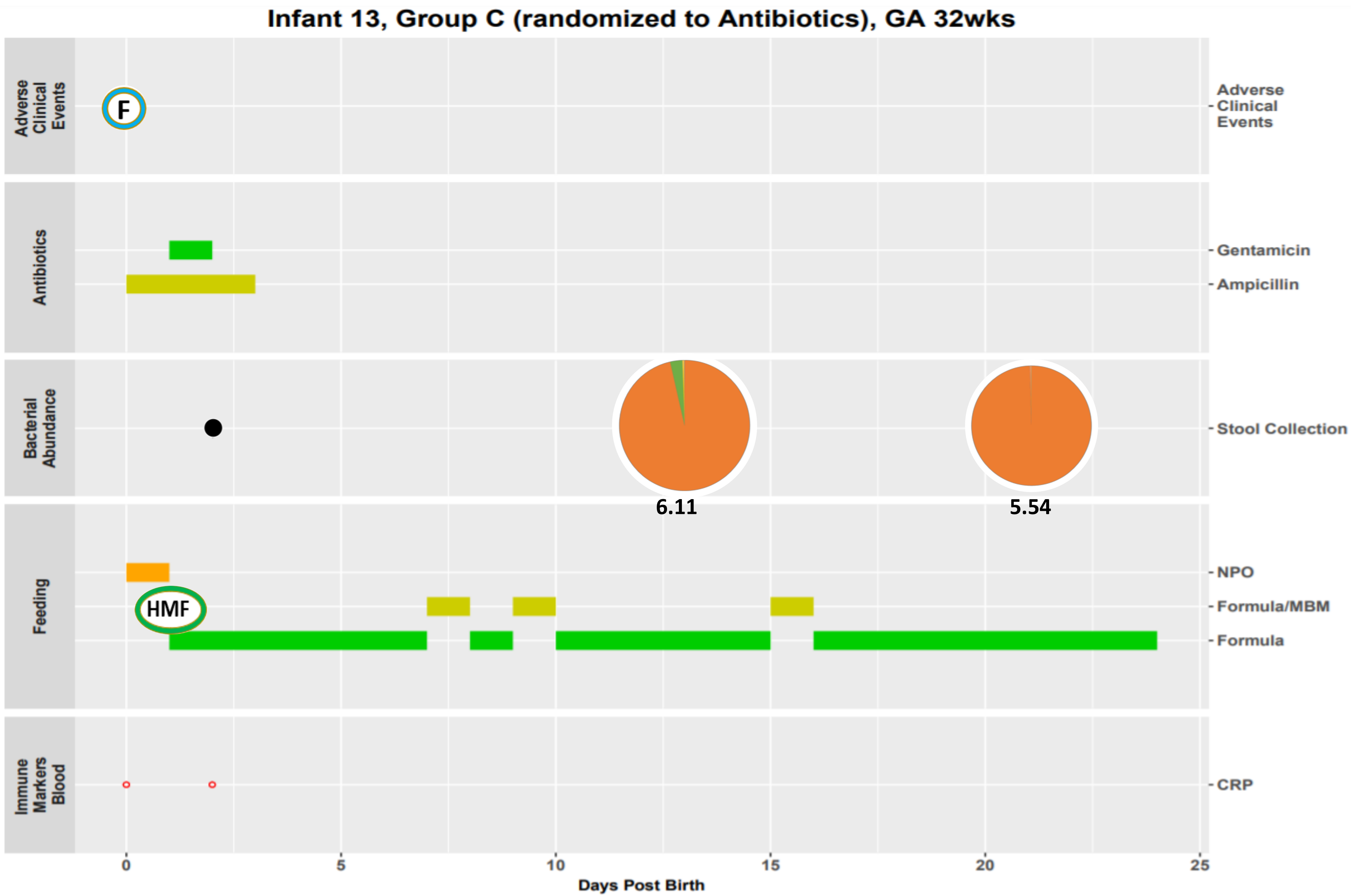

Infant 13, Group C (randomized to Antibiotics), GA 32wks
Adverse Clinical Events
F
Antibiotics
Gentamicin
Ampicillin
Bacterial Abundance
Stool Collection
6.11
5.54
Feeding
NPO
HMF
Formula/MBM
Formula
Immune Markers Blood
CRP
0
5
10
15
20
25
Days Post Birth

Infant 15, Group C (randomized to Antibiotics), GA 29wks

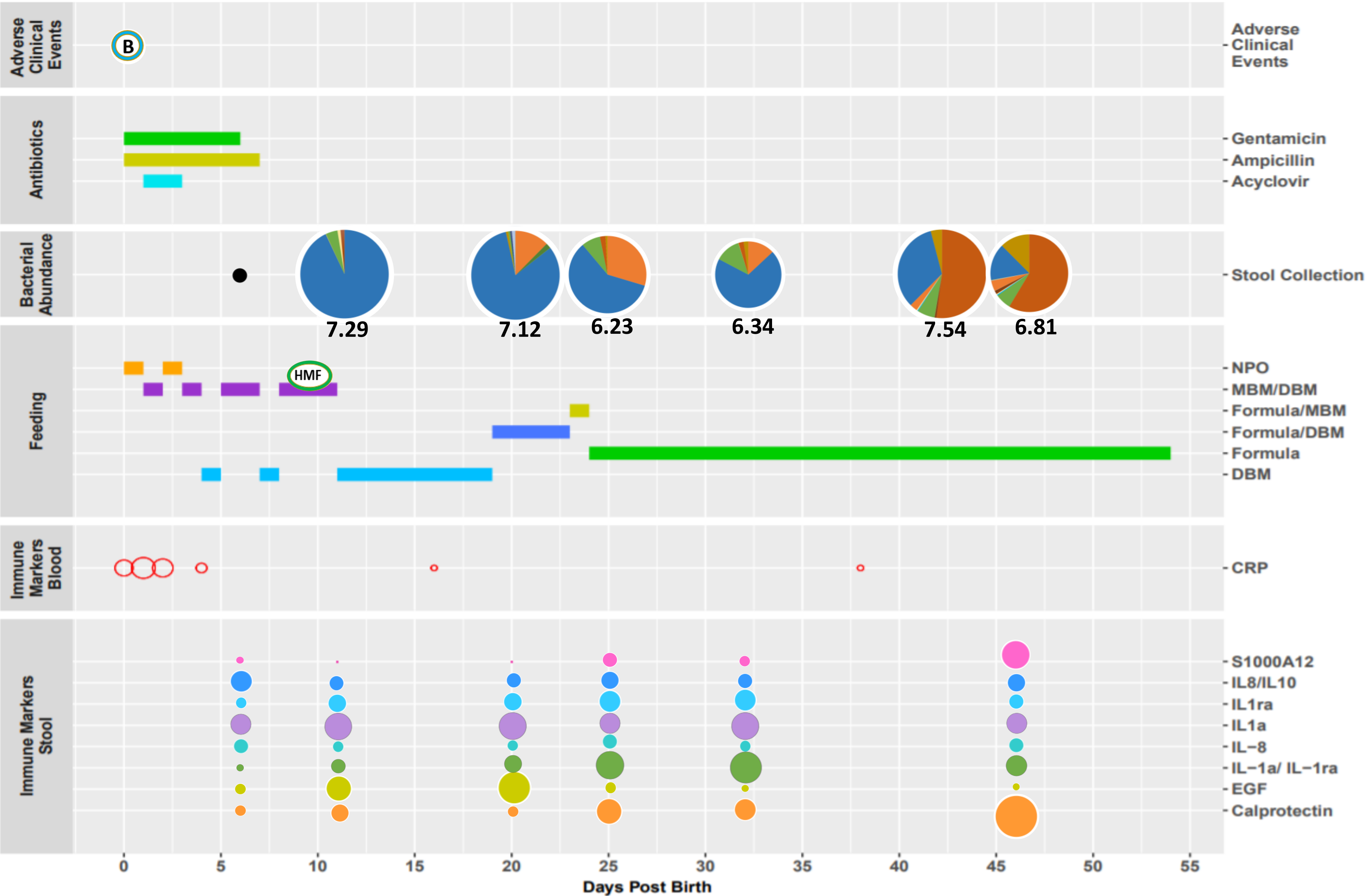

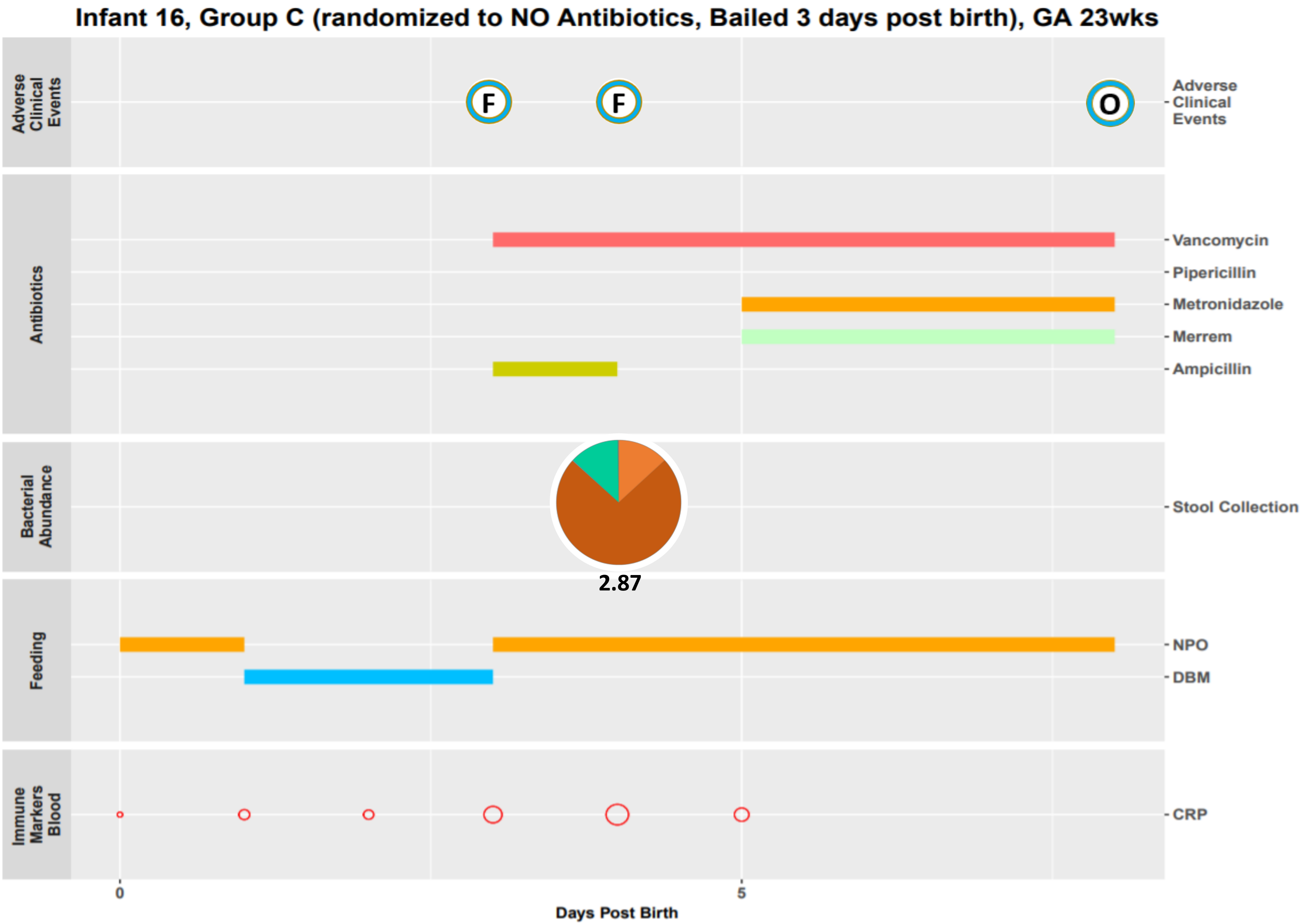

Infant 16, Group C (randomized to NO Antibiotics, Bailed 3 days post birth), GA 23wks
Adverse Clinical Events
F
F
O
Adverse Clinical Events
Antibiotics
Vancomycin
Pipericillin
Metronidazole
Merrem
Ampicillin
Bacterial Abundance
Stool Collection
2.87
Feeding
NPO
DBM
Immune Markers Blood
CRP
0
5
Days Post Birth

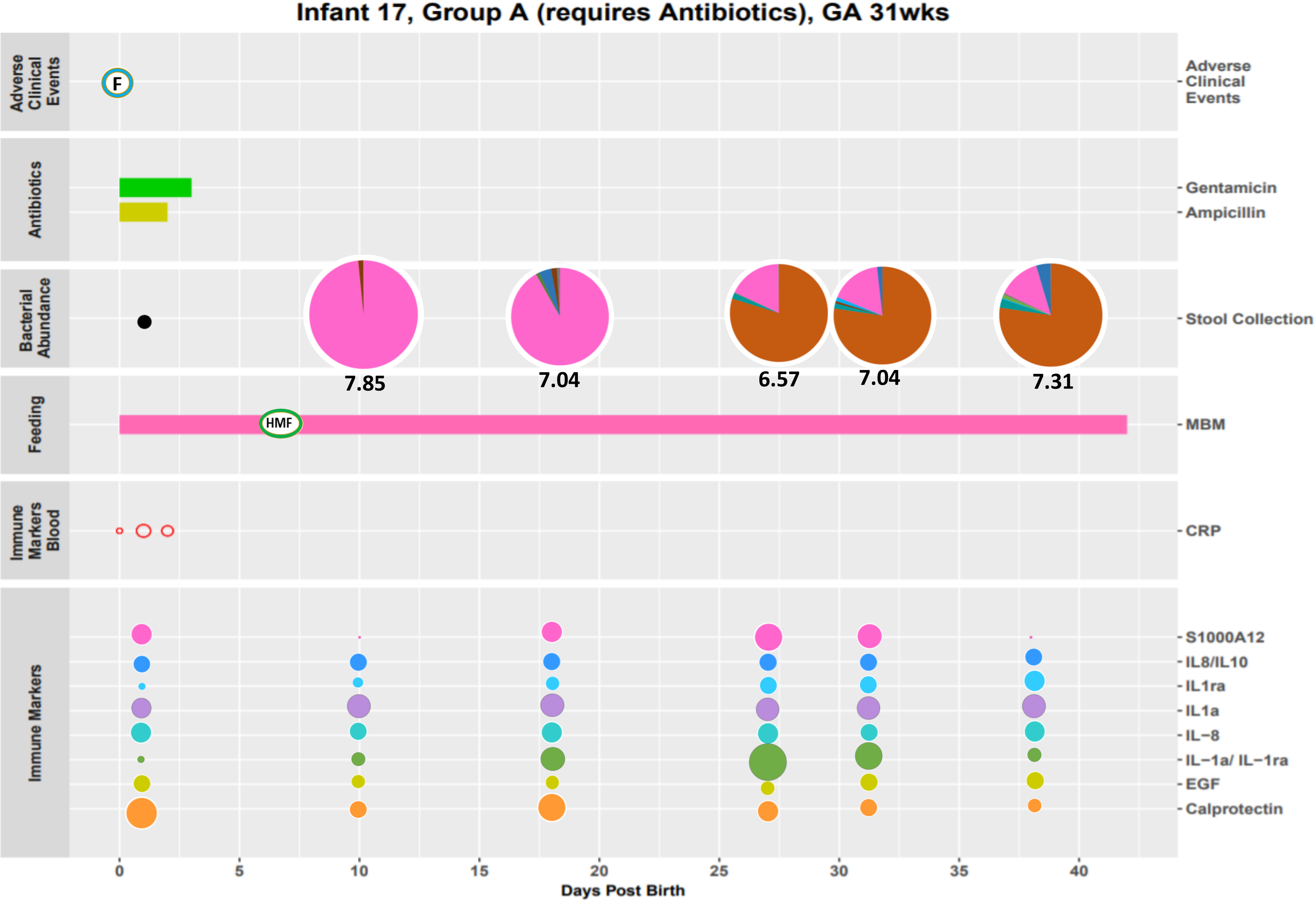

Infant 17, Group A (requires Antibiotics), GA 31wks
Adverse Clinical Events
F
Antibiotics
Gentamicin
Ampicillin
Bacterial Abundance
Stool Collection
7.85
7.04
6.57
7.04
7.31
Feeding
HMF
MBM
Immune Markers Blood
CRP
Immune Markers
S1000A12
IL8/IL10
IL1ra
IL1a
IL−8
IL−1a/ IL−1ra
EGF
Calprotectin
0
5
10
15
20
25
30
35
40
Days Post Birth

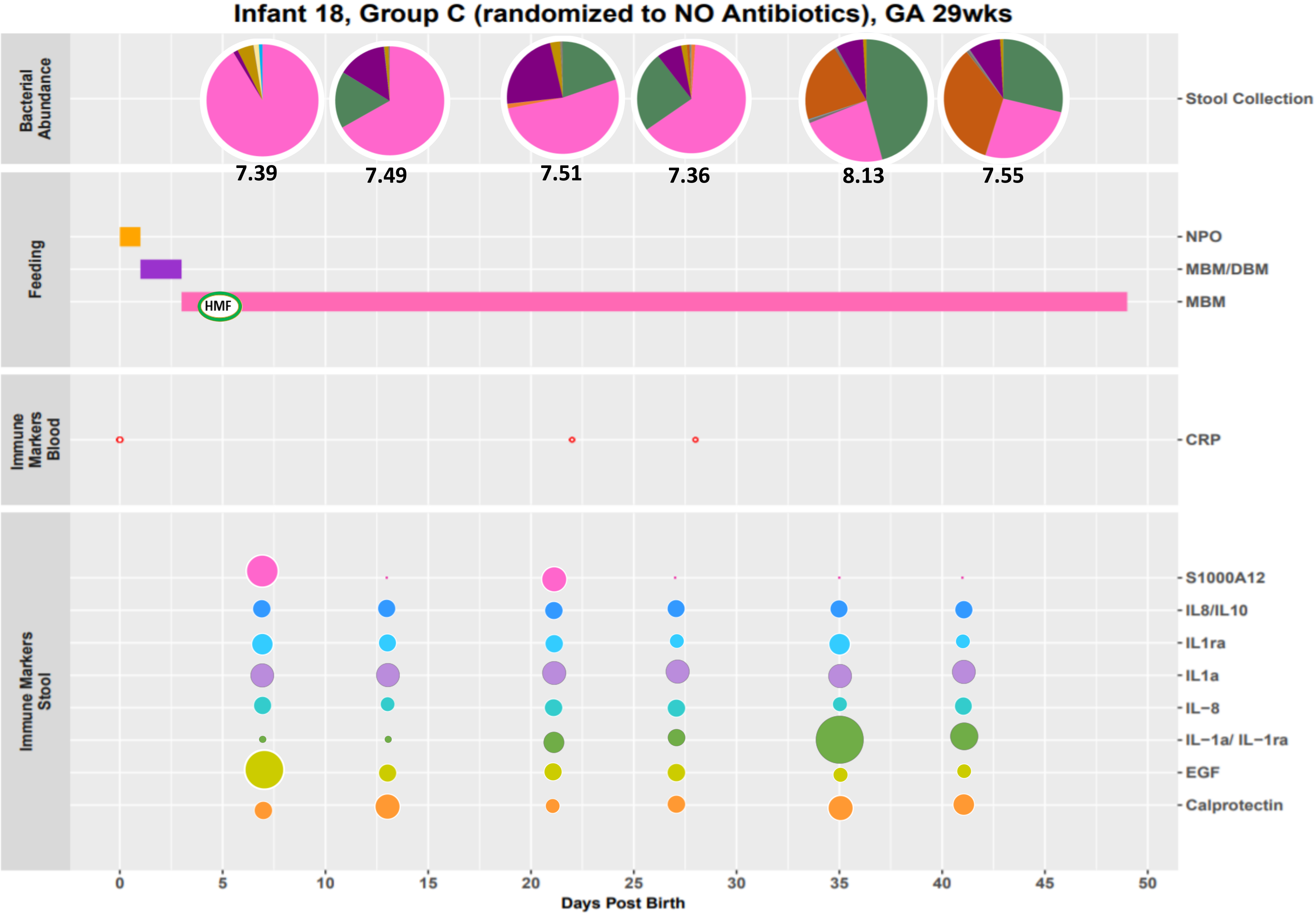
Infant 18, Group C (randomized to NO Antibiotics), GA 29wks
Bacterial Abundance
Stool Collection
7.39
7.49
7.51
7.36
8.13
7.55
Feeding
NPO
MBM/DBM
MBM
HMF
Immune Markers Blood
CRP
Immune Markers Stool
S1000A12
IL8/IL10
IL1ra
IL1a
IL−8
IL−1a/ IL−1ra
EGF
Calprotectin
0
5
10
15
20
25
30
35
40
45
50
Days Post Birth

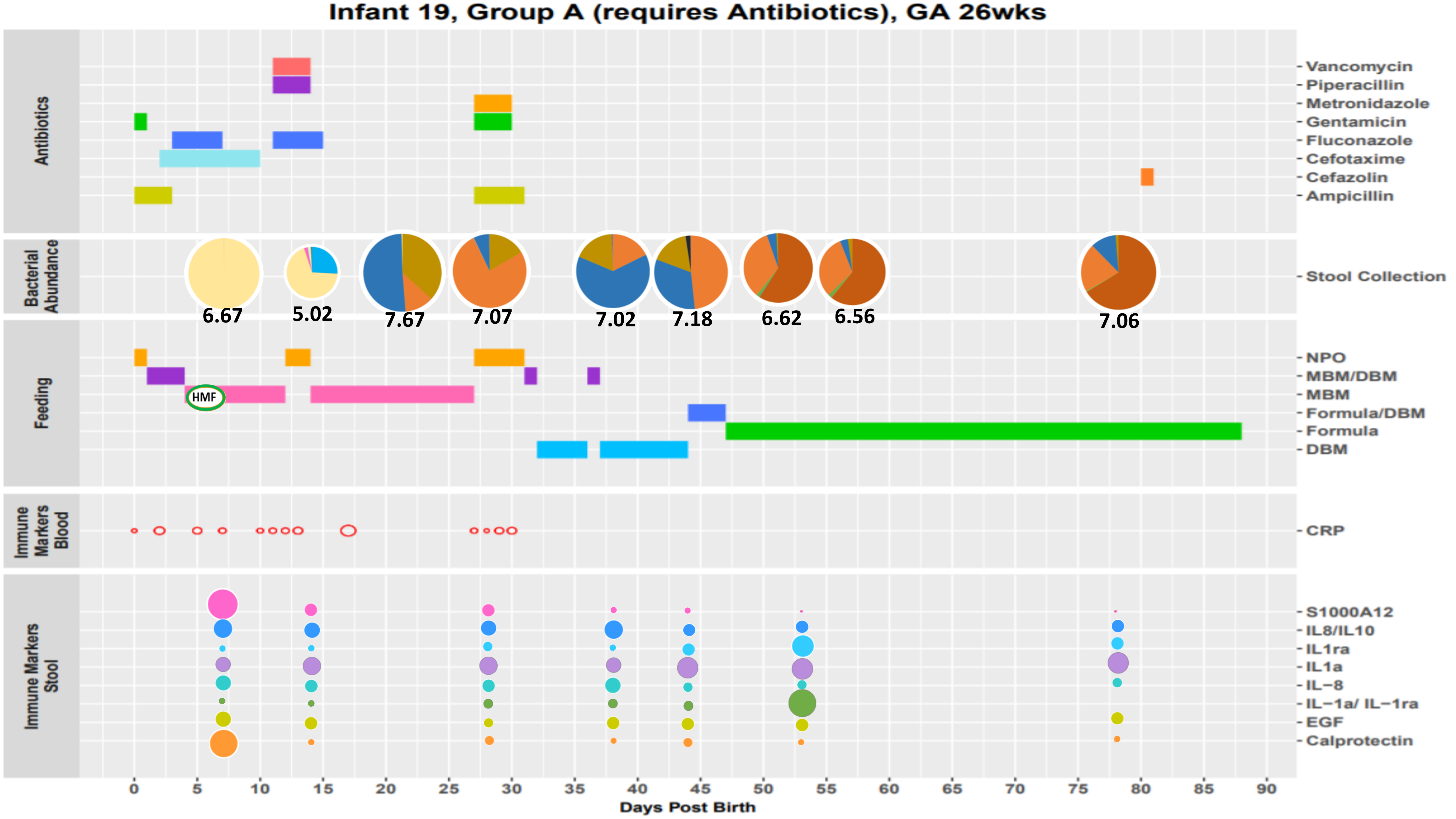
Infant 19, Group A (requires Antibiotics), GA 26wks
Antibiotics
Vancomycin
Piperacillin
Metronidazole
Gentamicin
Fluconazole
Cefotaxime
Cefazolin
Ampicillin
Bacterial Abundance
Stool Collection
6.67
5.02
7.67
7.07
7.02
7.18
6.62
6.56
7.06
Feeding
HMF
NPO
MBM/DBM
MBM
Formula/DBM
Formula
DBM
Immune Markers Blood
CRP
Immune Markers Stool
S1000A12
IL8/IL10
IL1ra
IL1a
IL−8
IL−1a/ IL−1ra
EGF
Calprotectin
0
5
10
15
20
25
30
35
40
45
50
55
60
65
70
75
80
85
90
Days Post Birth

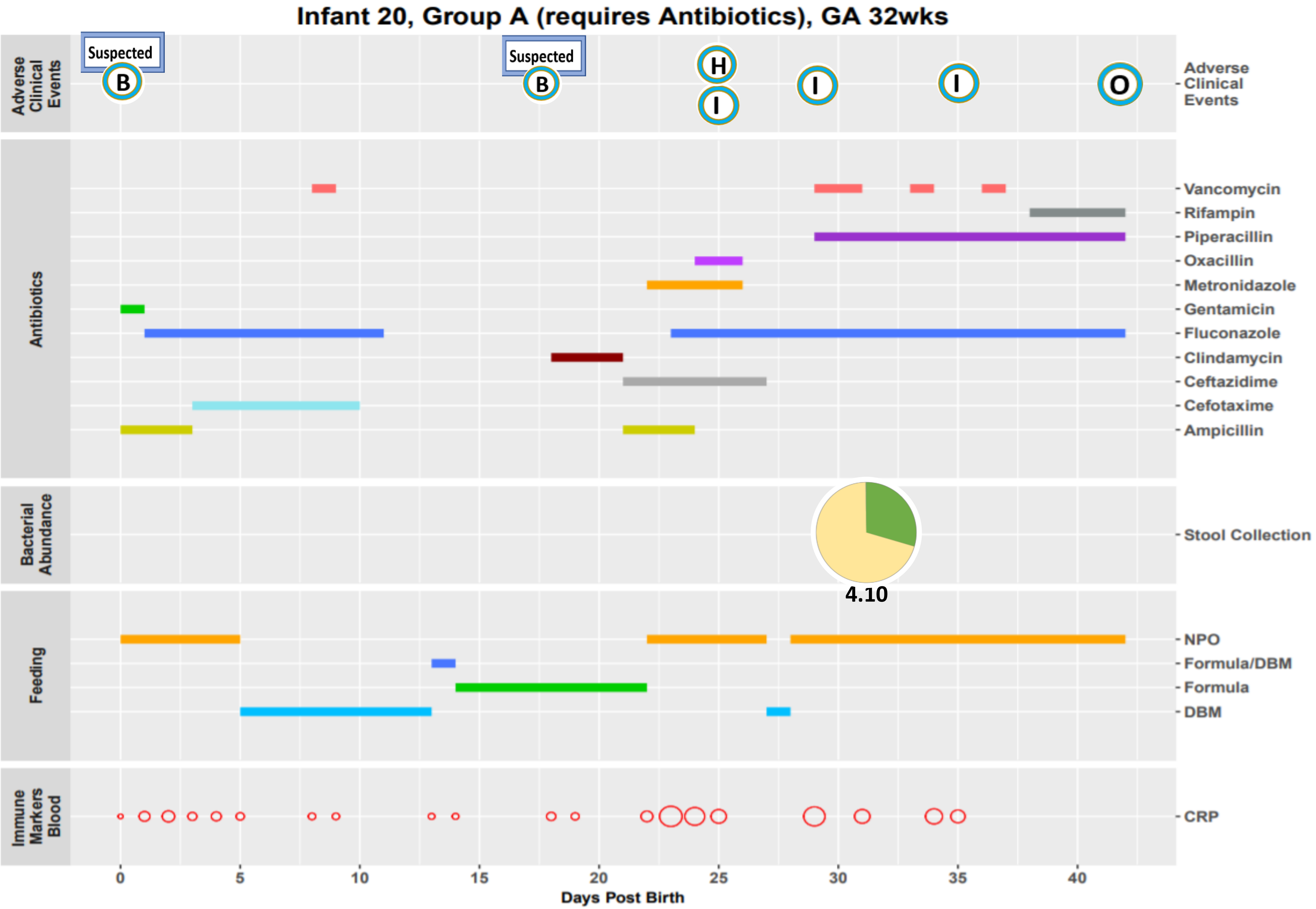
Infant 20, Group A (requires Antibiotics), GA 32wks
Adverse Clinical Events
Suspected
B
Suspected
B
H
I
I
I
O
Antibiotics
Vancomycin
Rifampin
Piperacillin
Oxacillin
Metronidazole
Gentamicin
Fluconazole
Clindamycin
Ceftazidime
Cefotaxime
Ampicillin
Bacterial Abundance
Stool Collection
4.10
Feeding
NPO
Formula/DBM
Formula
DBM
Immune Markers Blood
CRP
0
5
10
15
20
25
30
35
40
Days Post Birth

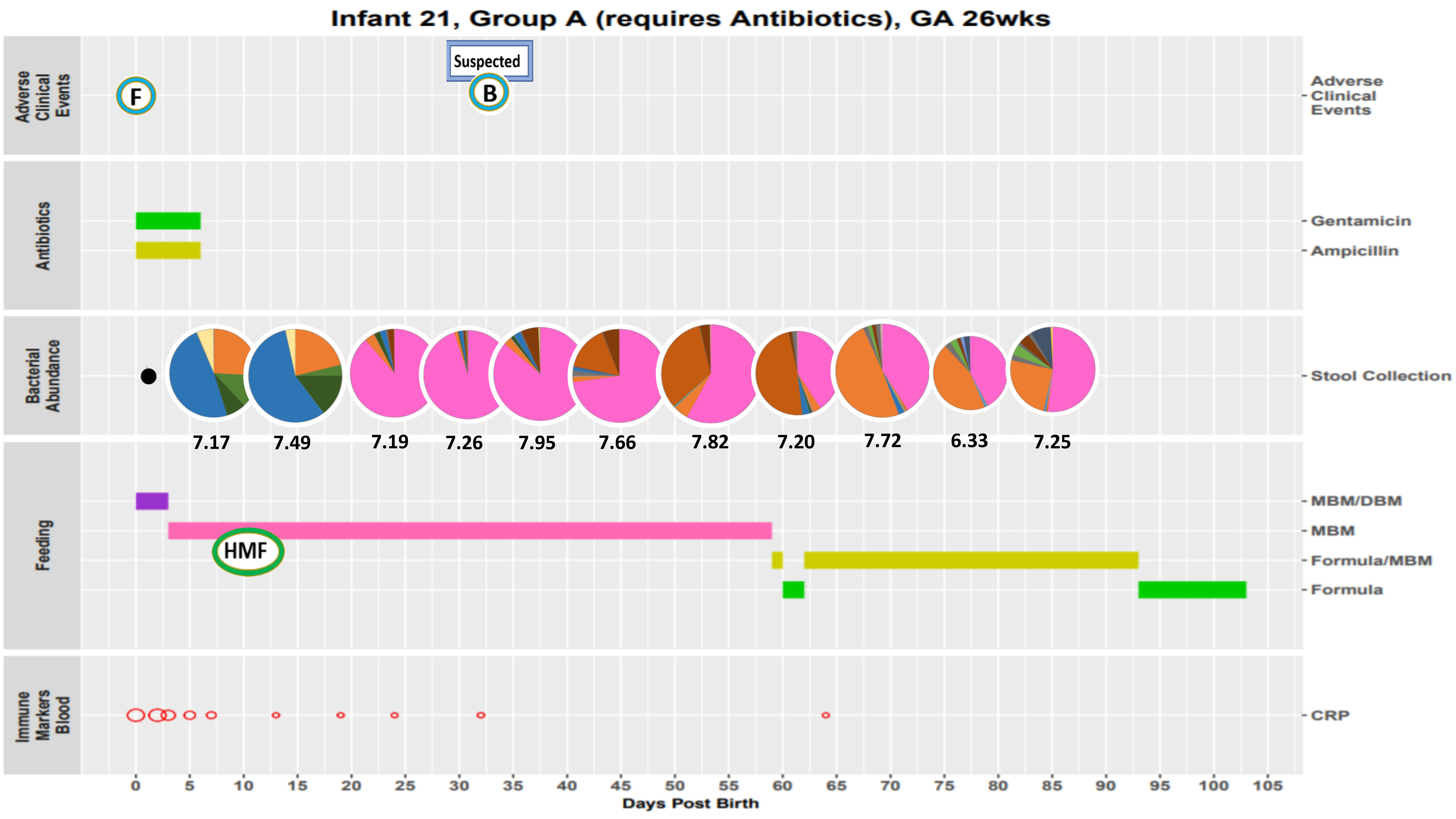
Infant 21, Group A (requires Antibiotics), GA 26wks
Adverse Clinical Events
Suspected
F
B
Antibiotics
Gentamicin
Ampicillin
Bacterial Abundance
Stool Collection
7.17
7.49
7.19
7.26
7.95
7.66
7.82
7.20
7.72
6.33
7.25
Feeding
MBM/DBM
MBM
HMF
Formula/MBM
Formula
Immune Markers Blood
CRP
0 5 10 15 20 25 30 35 40 45 50 55 60 65 70 75 80 85 90 95 100 105
Days Post Birth

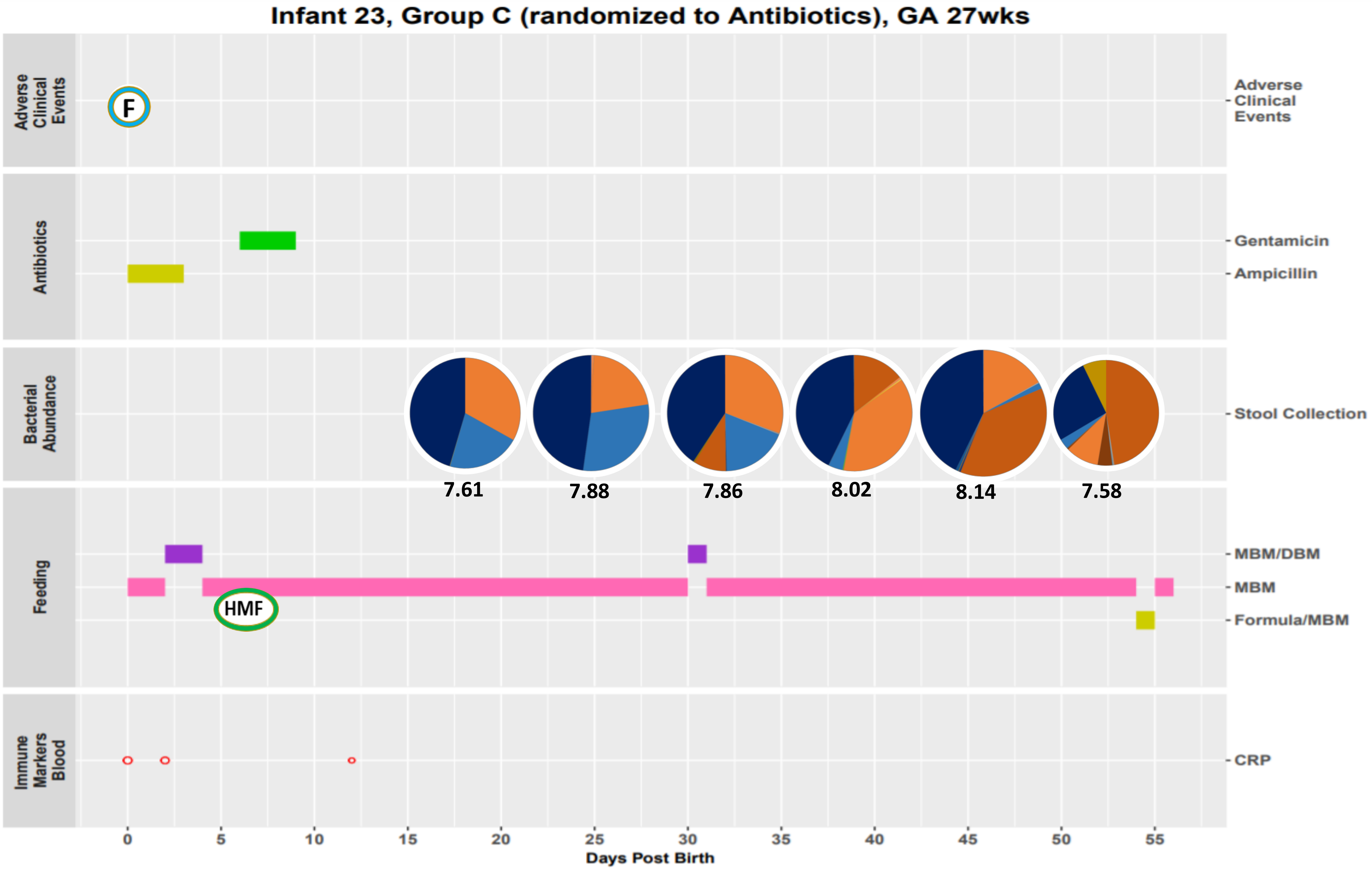

Infant 23, Group C (randomized to Antibiotics), GA 27wks
Adverse Clinical Events
F
Antibiotics
Gentamicin
Ampicillin
Bacterial Abundance
Stool Collection
7.61
7.88
7.86
8.02
8.14
7.58
Feeding
MBM/DBM
MBM
Formula/MBM
HMF
Immune Markers Blood
CRP
0
5
10
15
20
25
30
35
40
45
50
55
Days Post Birth

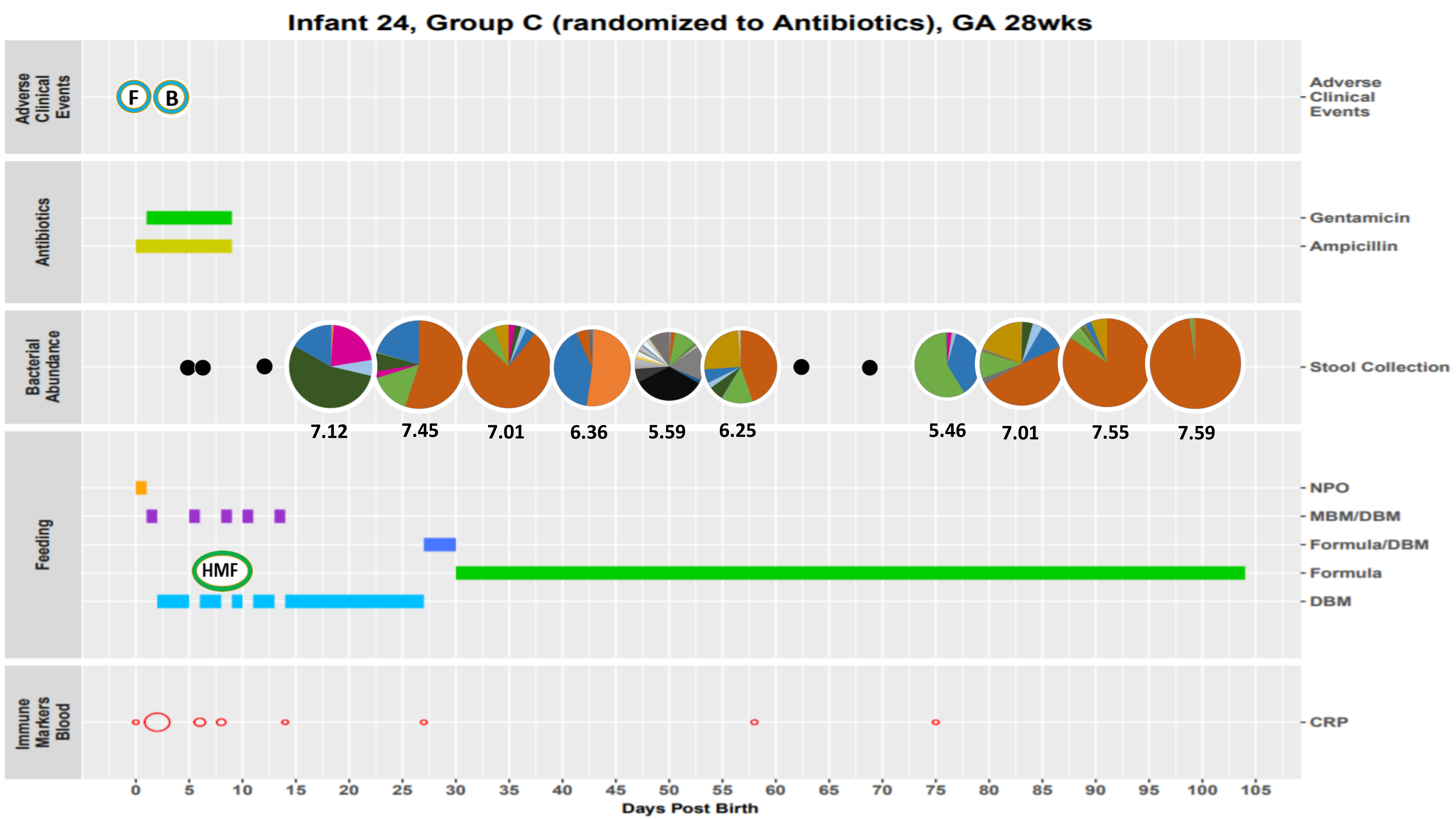
Infant 24, Group C (randomized to Antibiotics), GA 28wks
Adverse Clinical Events
F
B
Antibiotics
Gentamicin
Ampicillin
Bacterial Abundance
Stool Collection
7.12
7.45
7.01
6.36
5.59
6.25
5.46
7.01
7.55
7.59
Feeding
NPO
MBM/DBM
Formula/DBM
HMF
Formula
DBM
Immune Markers Blood
CRP
0
5
10
15
20
25
30
35
40
45
50
55
60
65
70
75
80
85
90
95
100
105
Days Post Birth

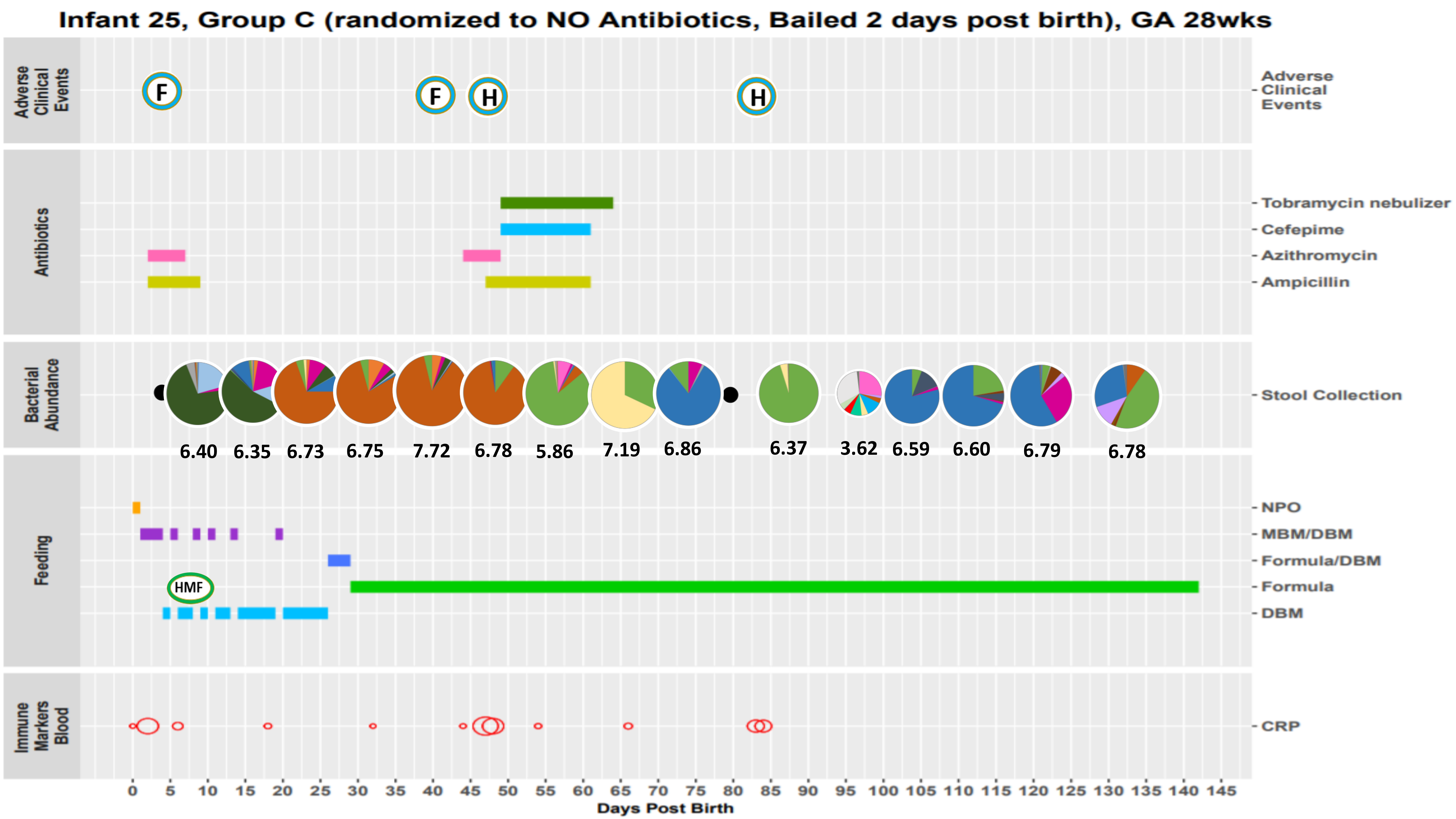
Infant 25, Group C (randomized to NO Antibiotics, Bailed 2 days post birth), GA 28wks
Adverse Clinical Events
F
F
H
H
Antibiotics
Tobramycin nebulizer
Cefepime
Azithromycin
Ampicillin
Bacterial Abundance
Stool Collection
6.40
6.35
6.73
6.75
7.72
6.78
5.86
7.19
6.86
6.37
3.62
6.59
6.60
6.79
6.78
Feeding
NPO
MBM/DBM
Formula/DBM
Formula
DBM
HMF
Immune Markers Blood
CRP
0 5 10 15 20 25 30 35 40 45 50 55 60 65 70 75 80 85 90 95 100 105 110 115 120 125 130 135 140 145
Days Post Birth

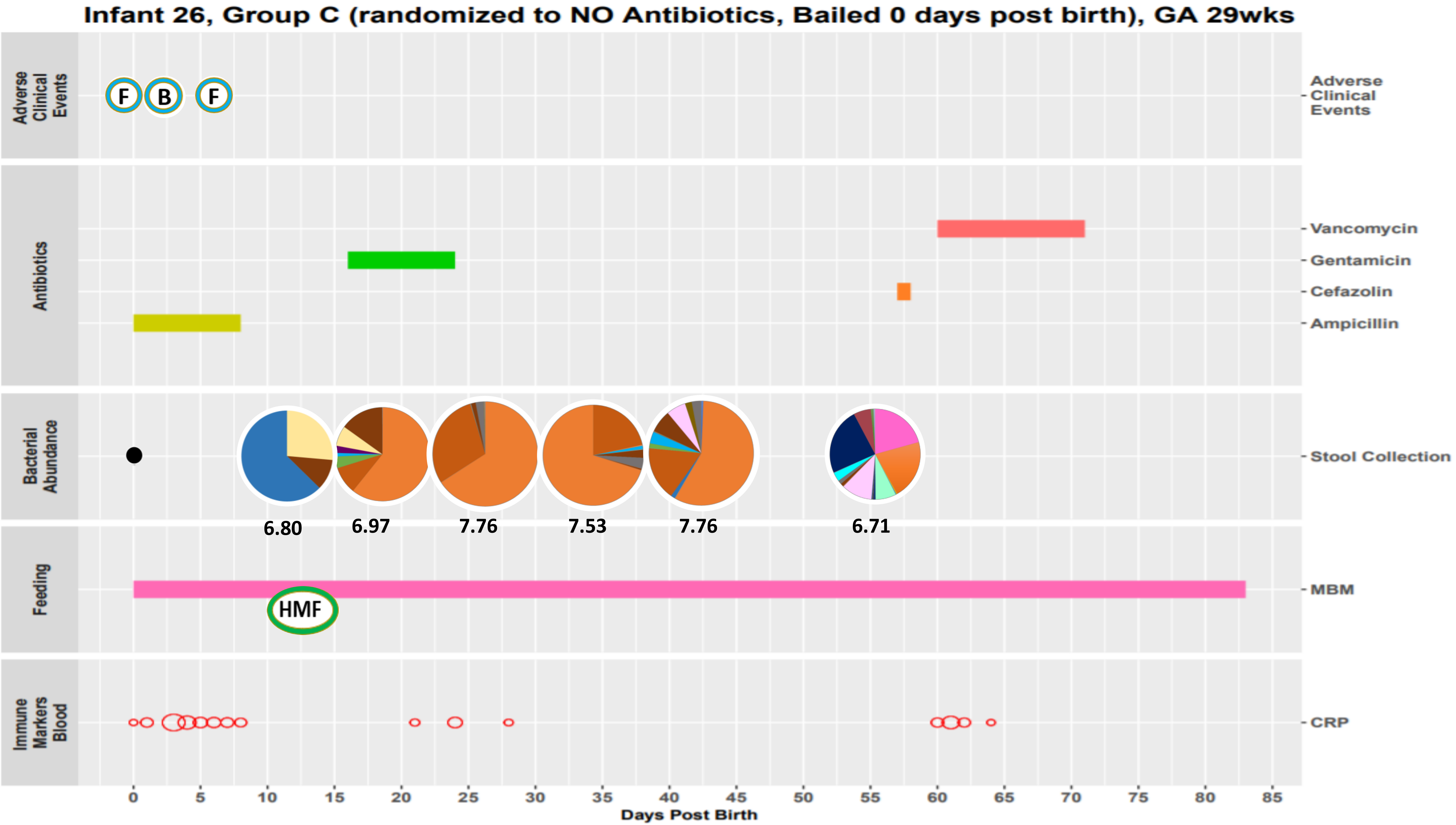

Infant 26, Group C (randomized to NO Antibiotics, Bailed 0 days post birth), GA 29wks
Adverse Clinical Events
F
B
F
Antibiotics
Vancomycin
Gentamicin
Cefazolin
Ampicillin
Bacterial Abundance
Stool Collection
6.80
6.97
7.76
7.53
7.76
6.71
Feeding
MBM
HMF
Immune Markers Blood
CRP
0
5
10
15
20
25
30
35
40
45
50
55
60
65
70
75
80
85
Days Post Birth

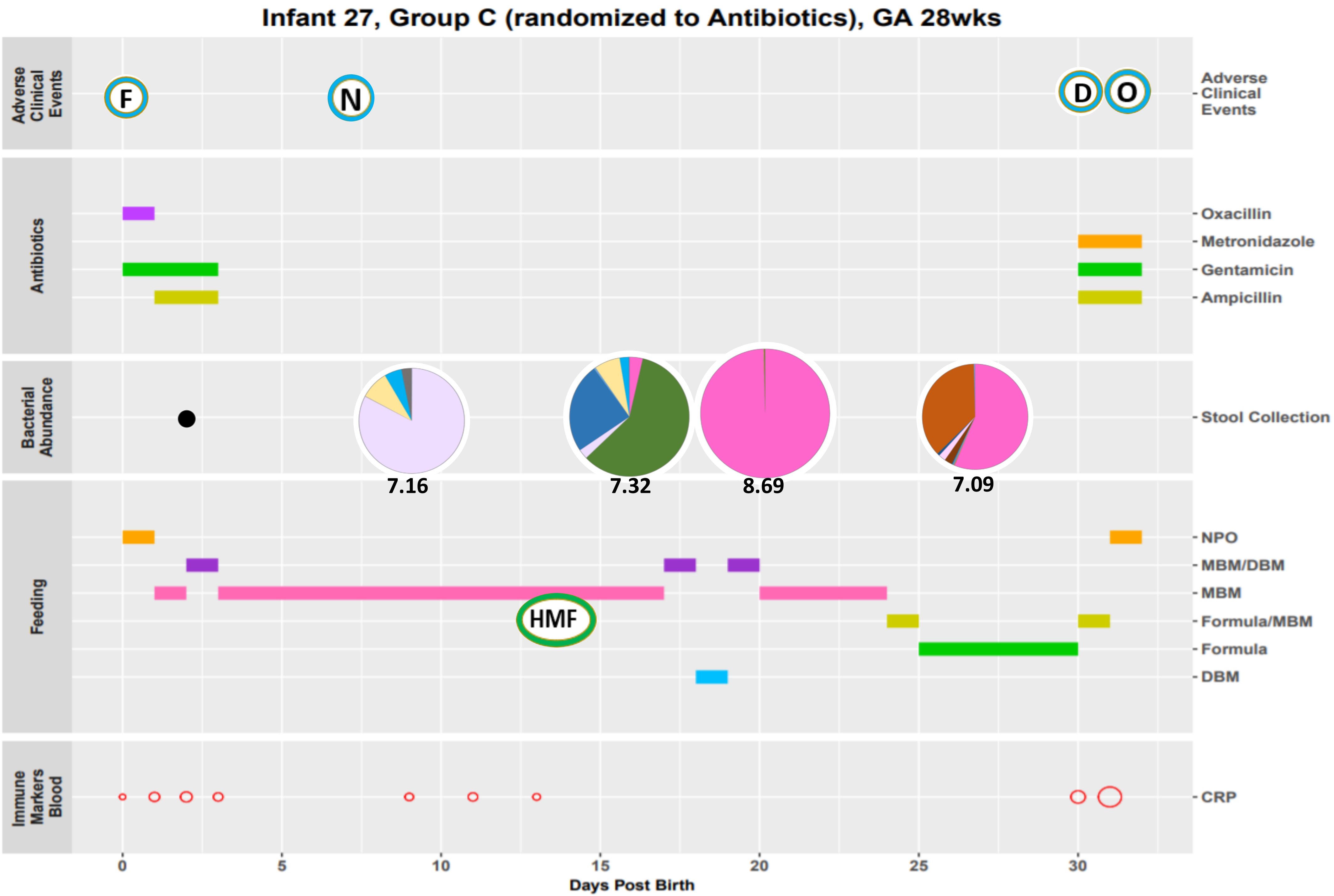
Infant 27, Group C (randomized to Antibiotics), GA 28wks
Adverse Clinical Events
F
N
D
O
Antibiotics
Oxacillin
Metronidazole
Gentamicin
Ampicillin
Bacterial Abundance
Stool Collection
7.16
7.32
8.69
7.09
Feeding
NPO
MBM/DBM
MBM
HMF
Formula/MBM
Formula
DBM
Immune Markers Blood
CRP
0
5
10
15
20
25
30
Days Post Birth

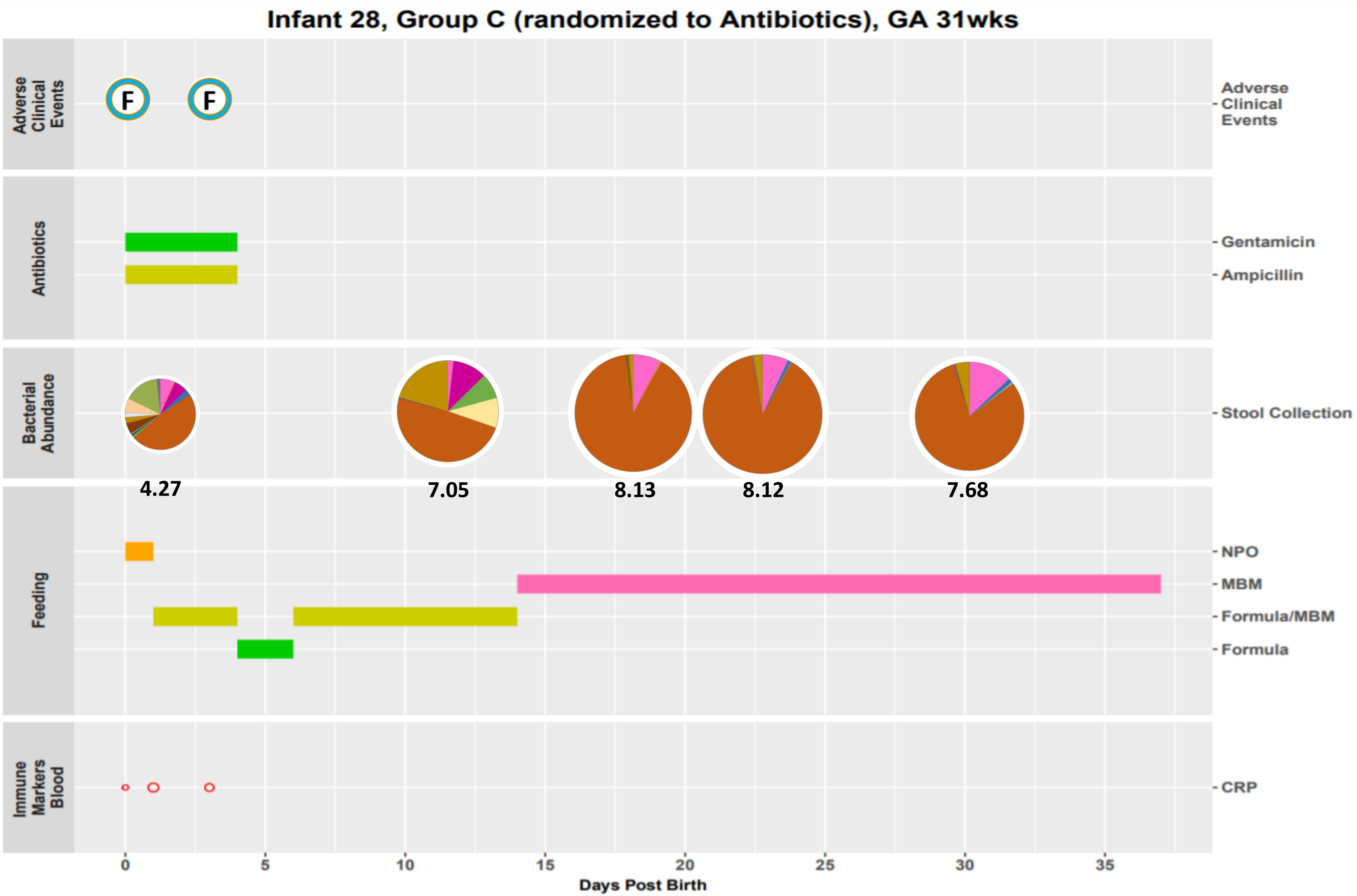
Infant 28, Group C (randomized to Antibiotics), GA 31wks
Adverse Clinical Events
F
F
Antibiotics
Gentamicin
Ampicillin
Bacterial Abundance
Stool Collection
4.27
7.05
8.13
8.12
7.68
Feeding
NPO
MBM
Formula/MBM
Formula
Immune Markers Blood
CRP
0
5
10
15
20
25
30
35
Days Post Birth

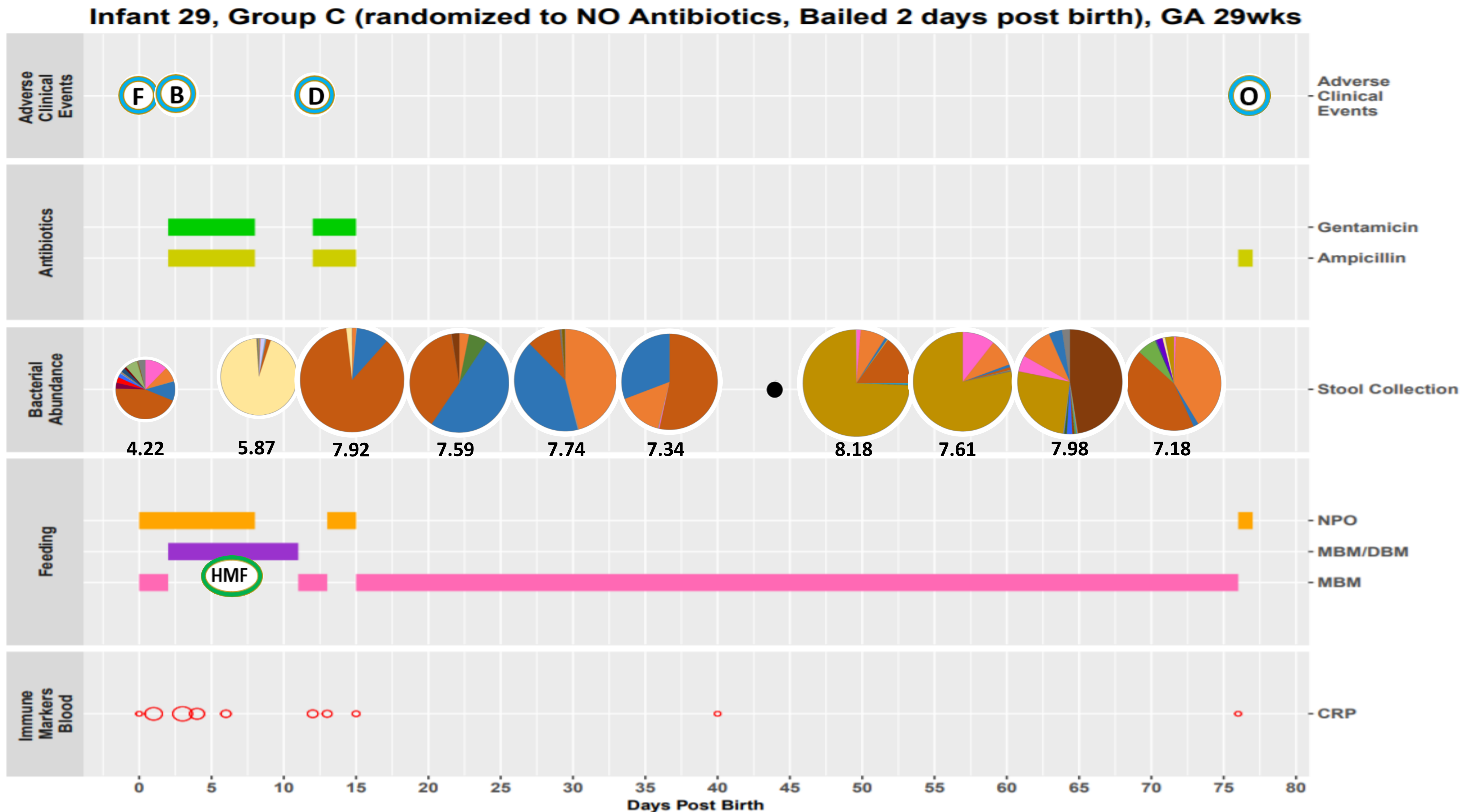
Infant 29, Group C (randomized to NO Antibiotics, Bailed 2 days post birth), GA 29wks
Adverse Clinical Events
F
B
D
O
Antibiotics
Gentamicin
Ampicillin
Bacterial Abundance
Stool Collection
4.22
5.87
7.92
7.59
7.74
7.34
8.18
7.61
7.98
7.18
Feeding
NPO
MBM/DBM
MBM
HMF
Immune Markers Blood
CRP
0
5
10
15
20
25
30
35
40
45
50
55
60
65
70
75
80
Days Post Birth

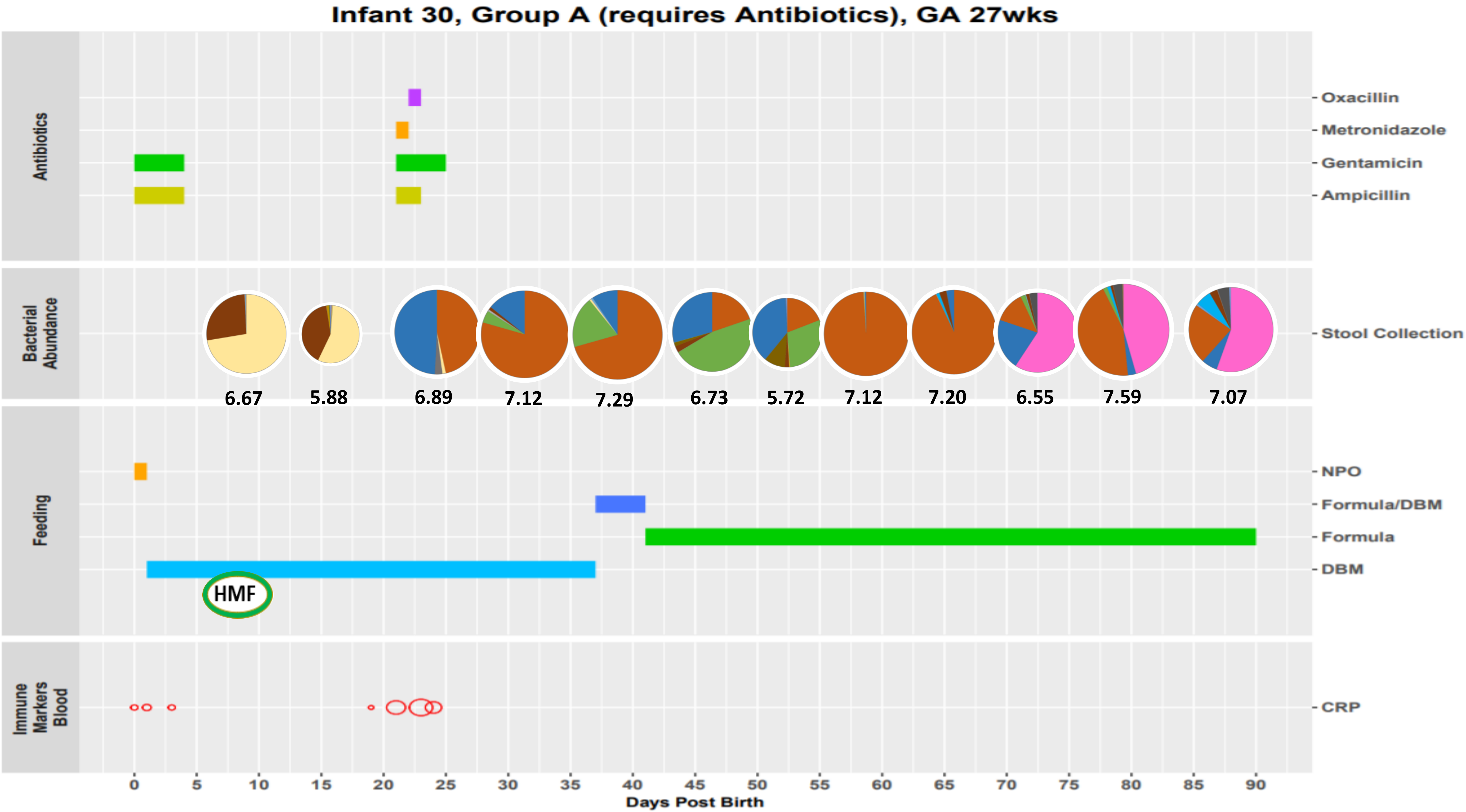

Infant 30, Group A (requires Antibiotics), GA 27wks
Antibiotics
Oxacillin
Metronidazole
Gentamicin
Ampicillin
Bacterial Abundance
Stool Collection
6.67
5.88
6.89
7.12
7.29
6.73
5.72
7.12
7.20
6.55
7.59
7.07
Feeding
NPO
Formula/DBM
Formula
DBM
HMF
Immune Markers Blood
CRP
0
5
10
15
20
25
30
35
40
45
50
55
60
65
70
75
80
85
90
Days Post Birth

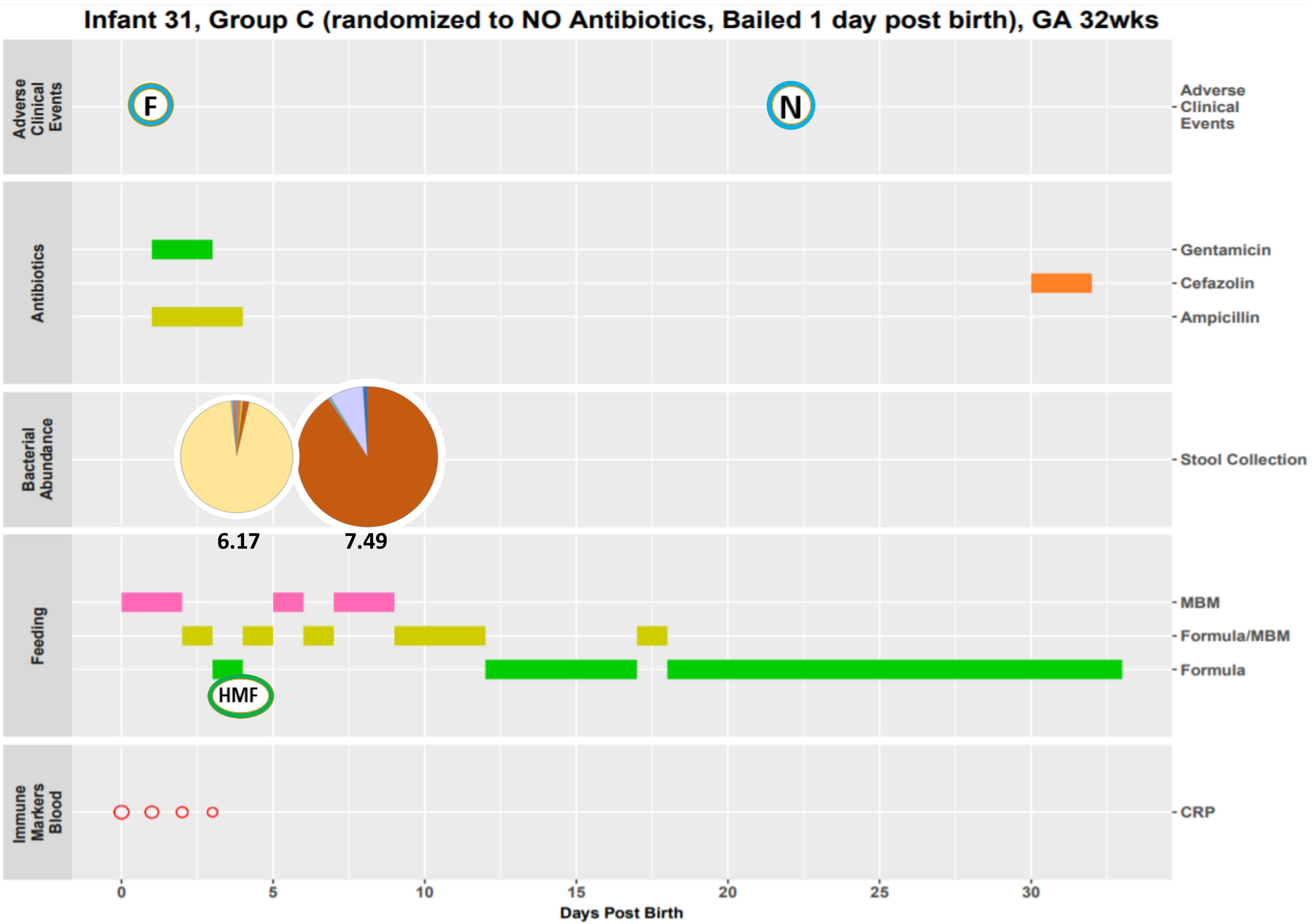
Infant 31, Group C (randomized to NO Antibiotics, Bailed 1 day post birth), GA 32wks
Adverse Clinical Events
F
N
Antibiotics
Gentamicin
Cefazolin
Ampicillin
Bacterial Abundance
Stool Collection
6.17
7.49
Feeding
MBM
Formula/MBM
Formula
HMF
Immune Markers Blood
CRP
0
5
10
15
20
25
30
Days Post Birth

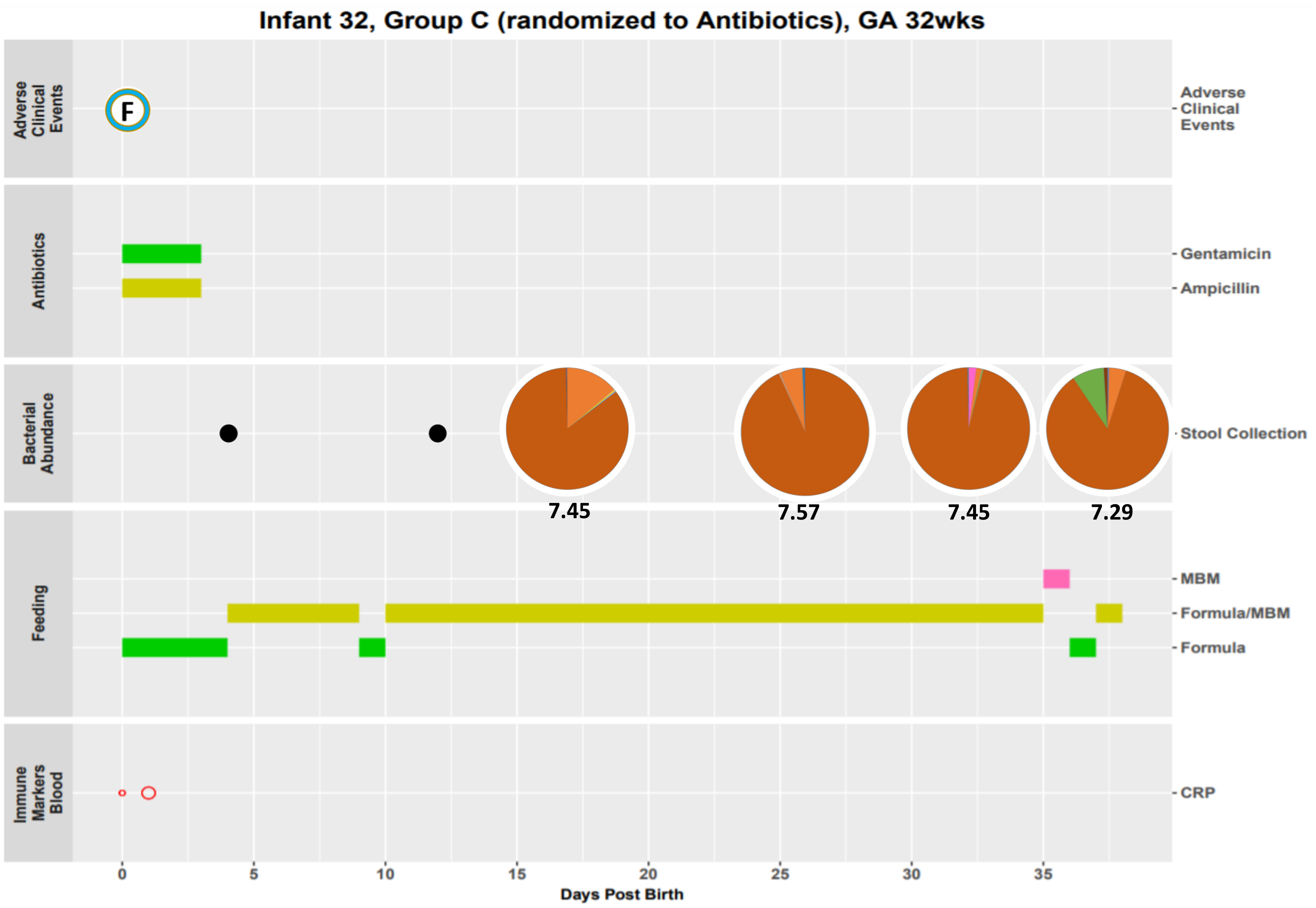

Infant 32, Group C (randomized to Antibiotics), GA 32wks
Adverse Clinical Events
F
Antibiotics
Gentamicin
Ampicillin
Bacterial Abundance
Stool Collection
7.45
7.57
7.45
7.29
Feeding
MBM
Formula/MBM
Formula
Immune Markers Blood
CRP
0
5
10
15
20
25
30
35
Days Post Birth

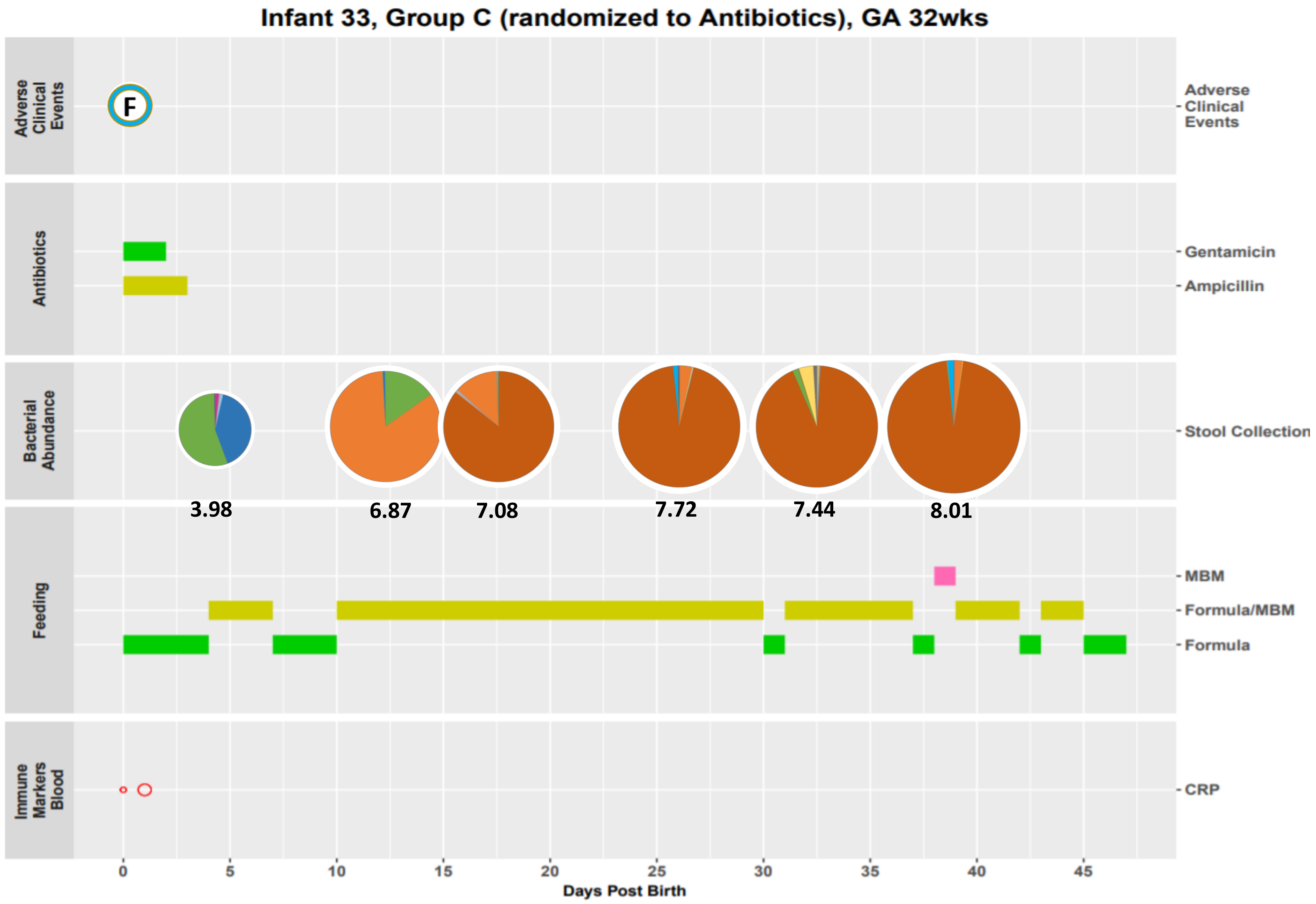

Infant 33, Group C (randomized to Antibiotics), GA 32wks
Adverse Clinical Events
F
Antibiotics
Gentamicin
Ampicillin
Bacterial Abundance
Stool Collection
3.98
6.87
7.08
7.72
7.44
8.01
Feeding
MBM
Formula/MBM
Formula
Immune Markers Blood
CRP
0
5
10
15
20
25
30
35
40
45
Days Post Birth

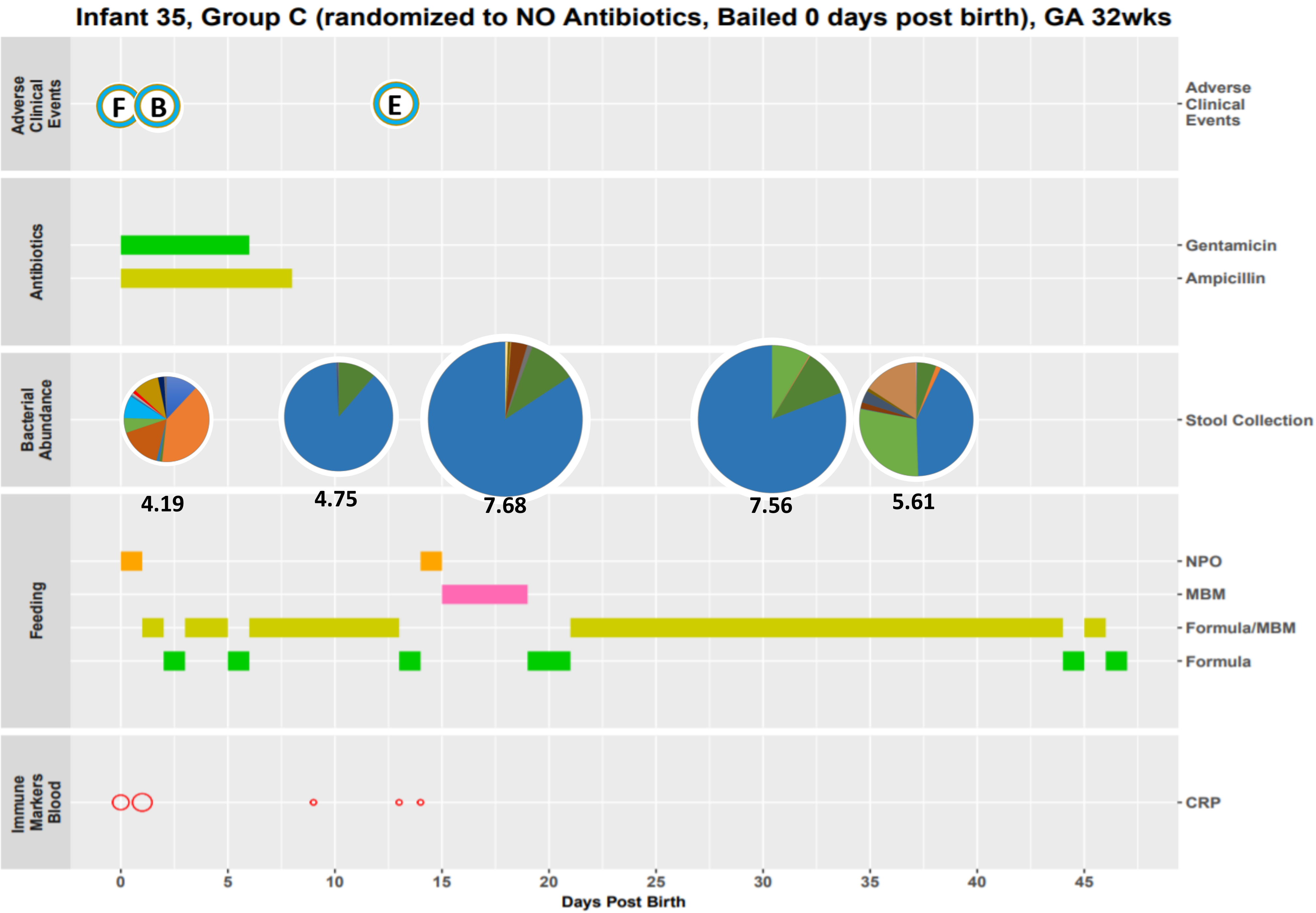
Infant 35, Group C (randomized to NO Antibiotics, Bailed 0 days post birth), GA 32wks
Adverse Clinical Events
F
B
E
Antibiotics
Gentamicin
Ampicillin
Bacterial Abundance
Stool Collection
4.19
4.75
7.68
7.56
5.61
Feeding
NPO
MBM
Formula/MBM
Formula
Immune Markers Blood
CRP
0
5
10
15
20
25
30
35
40
45
Days Post Birth

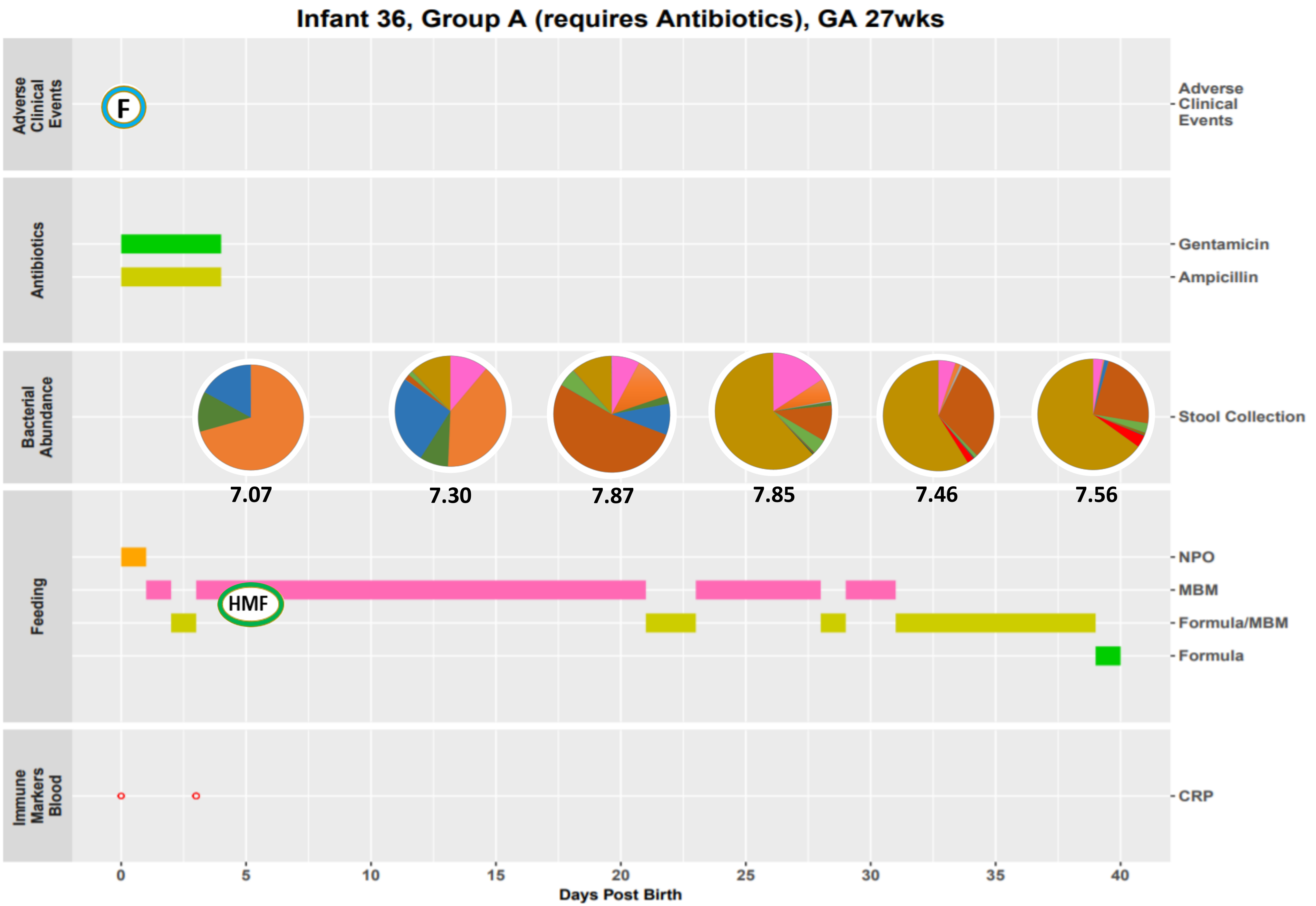
Infant 36, Group A (requires Antibiotics), GA 27wks
Adverse Clinical Events
F
Antibiotics
Gentamicin
Ampicillin
Bacterial Abundance
Stool Collection
7.07
7.30
7.87
7.85
7.46
7.56
Feeding
NPO
MBM
HMF
Formula/MBM
Formula
Immune Markers Blood
CRP
0
5
10
15
20
25
30
35
40
Days Post Birth

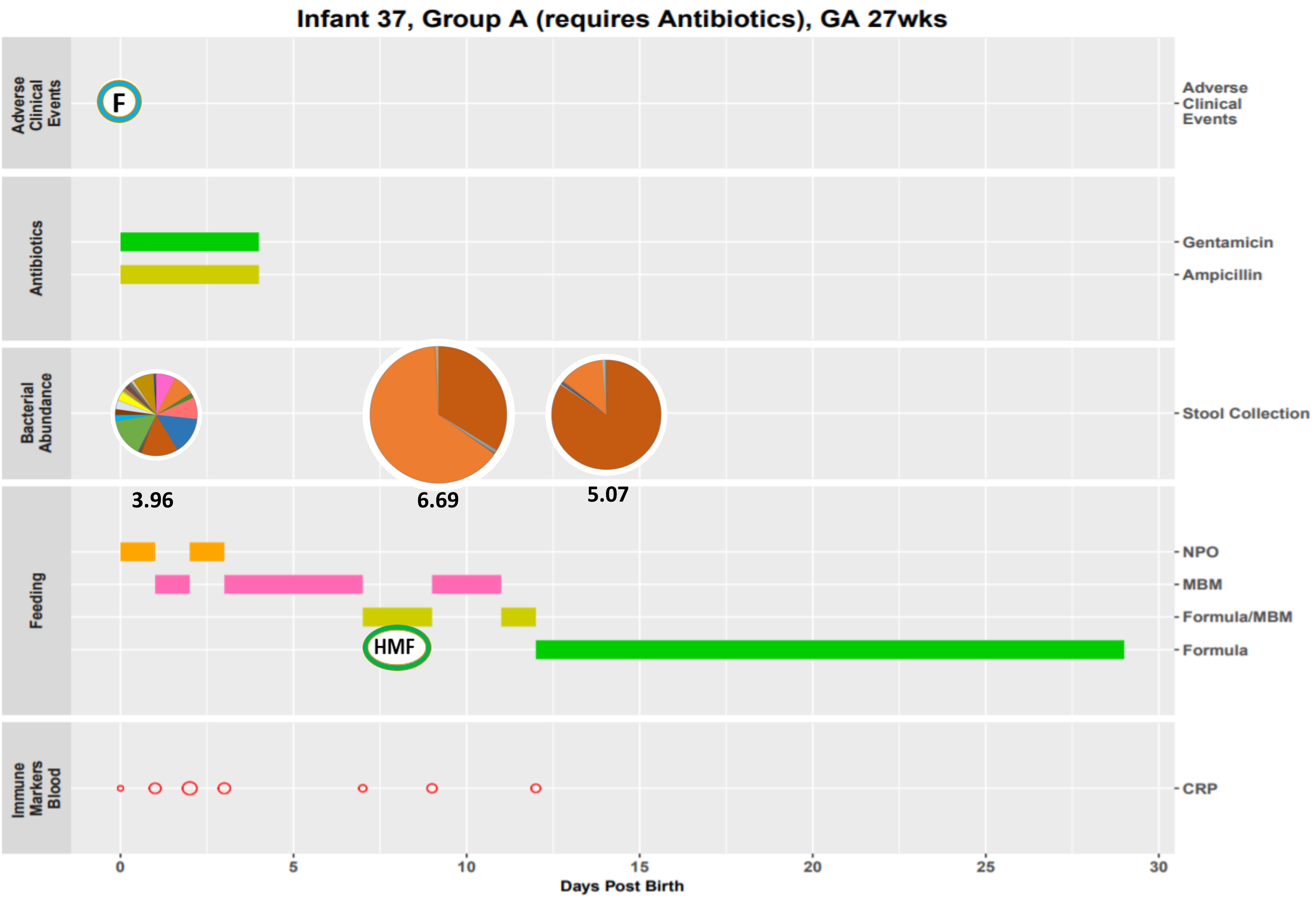

Infant 37, Group A (requires Antibiotics), GA 27wks
Adverse Clinical Events
F
Antibiotics
Gentamicin
Ampicillin
Bacterial Abundance
Stool Collection
3.96
6.69
5.07
Feeding
NPO
MBM
Formula/MBM
Formula
HMF
Immune Markers Blood
CRP
0
5
10
15
20
25
30
Days Post Birth

Infant 38, Group C (randomized to Antibiotics), GA 28wks

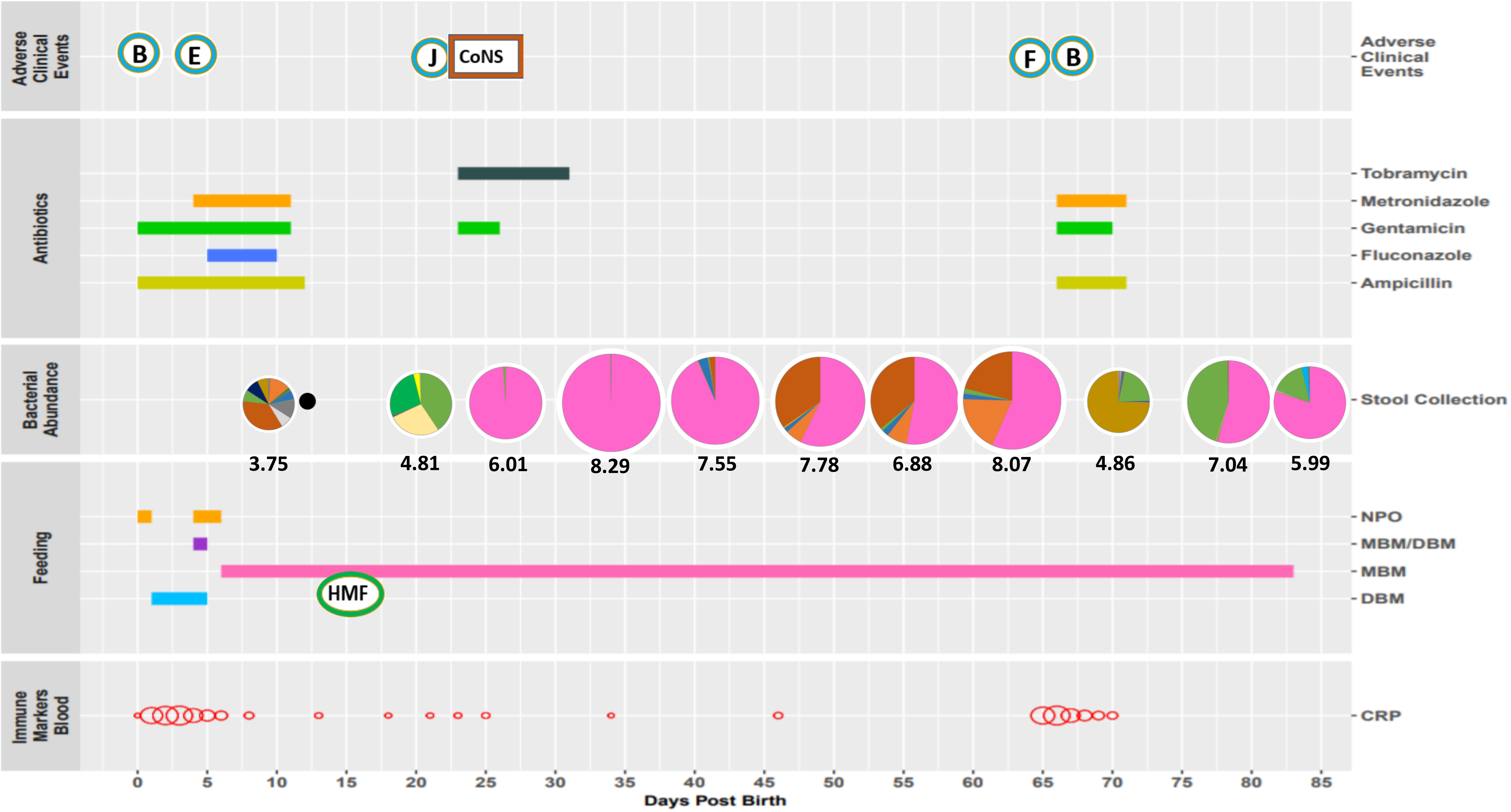

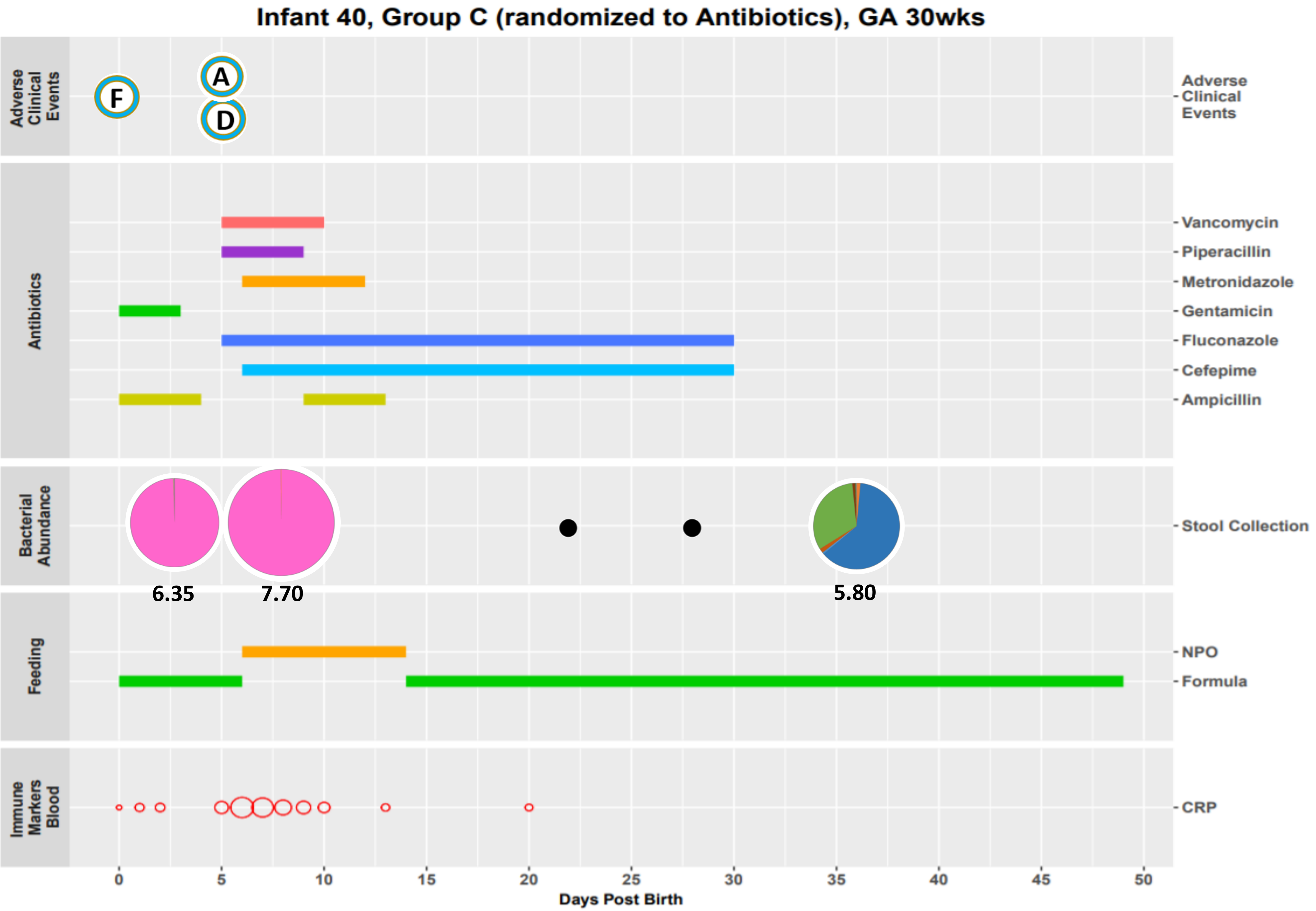
Infant 40, Group C (randomized to Antibiotics), GA 30wks
Adverse Clinical Events
F
A
D
Antibiotics
Vancomycin
Piperacillin
Metronidazole
Gentamicin
Fluconazole
Cefepime
Ampicillin
Bacterial Abundance
Stool Collection
6.35
7.70
5.80
Feeding
NPO
Formula
Immune Markers Blood
CRP
0
5
10
15
20
25
30
35
40
45
50
Days Post Birth

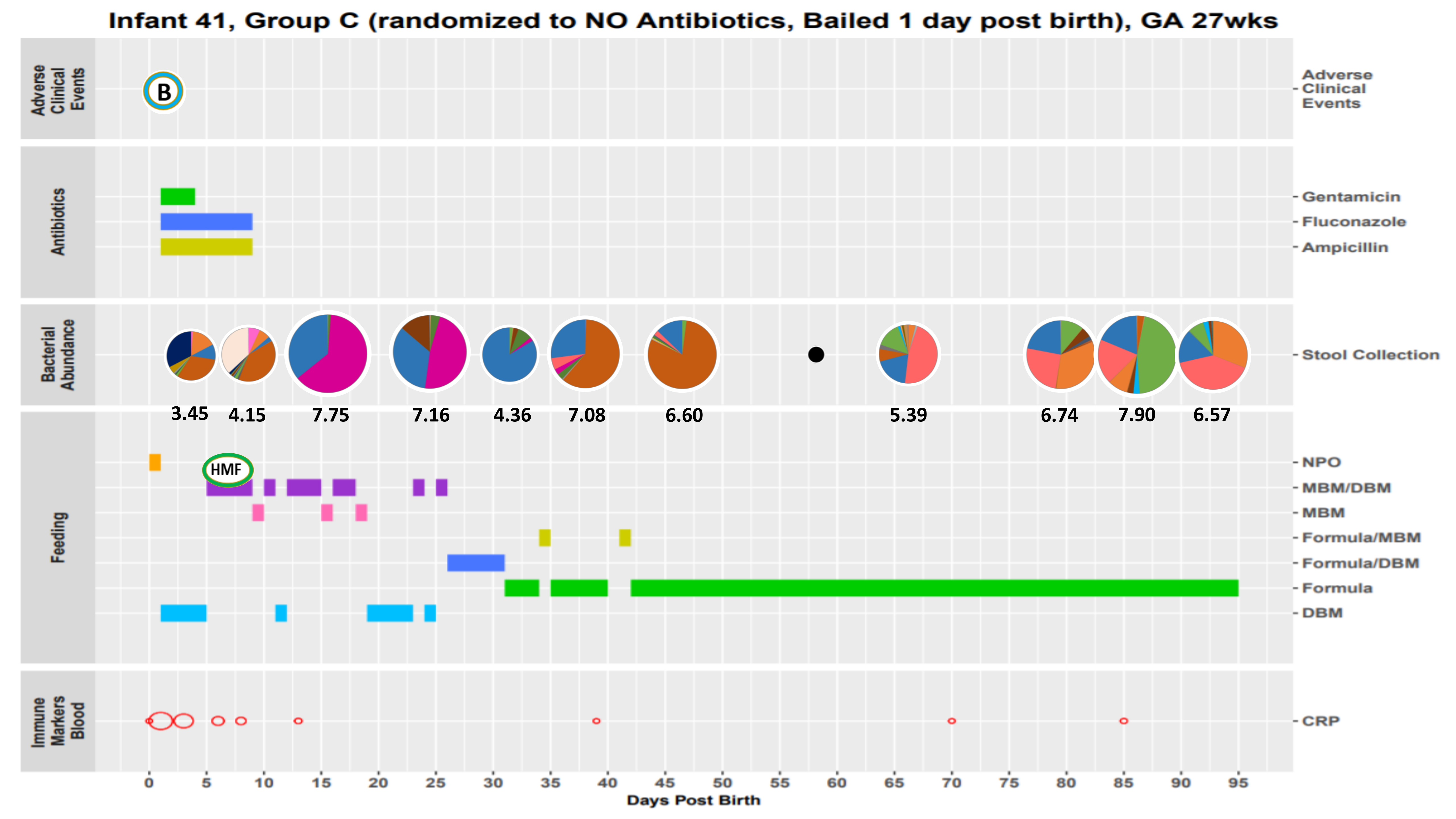
Infant 41, Group C (randomized to NO Antibiotics, Bailed 1 day post birth), GA 27wks
Adverse Clinical Events
B
Antibiotics
Gentamicin
Fluconazole
Ampicillin
Bacterial Abundance
Stool Collection
3.45
4.15
7.75
7.16
4.36
7.08
6.60
5.39
6.74
7.90
6.57
Feeding
HMF
NPO
MBM/DBM
MBM
Formula/MBM
Formula/DBM
Formula
DBM
Immune Markers Blood
CRP
0
5
10
15
20
25
30
35
40
45
50
55
60
65
70
75
80
85
90
95
Days Post Birth

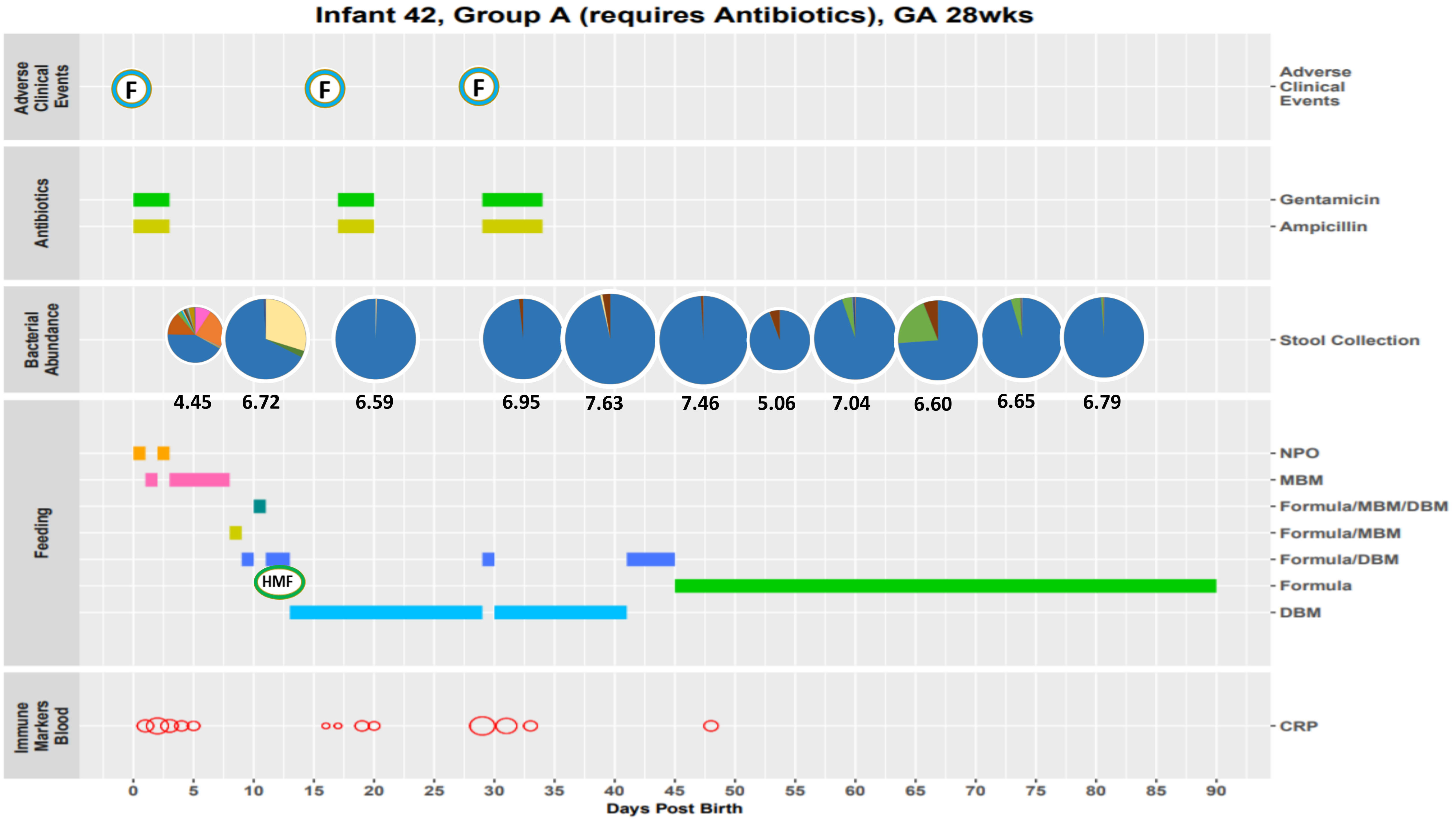

Infant 42, Group A (requires Antibiotics), GA 28wks
Adverse Clinical Events
F
F
F
Antibiotics
Gentamicin
Ampicillin
Bacterial Abundance
Stool Collection
4.45
6.72
6.59
6.95
7.63
7.46
5.06
7.04
6.60
6.65
6.79
Feeding
NPO
MBM
Formula/MBM/DBM
Formula/MBM
Formula/DBM
Formula
DBM
HMF
Immune Markers Blood
CRP
0
5
10
15
20
25
30
35
40
45
50
55
60
65
70
75
80
85
90
Days Post Birth

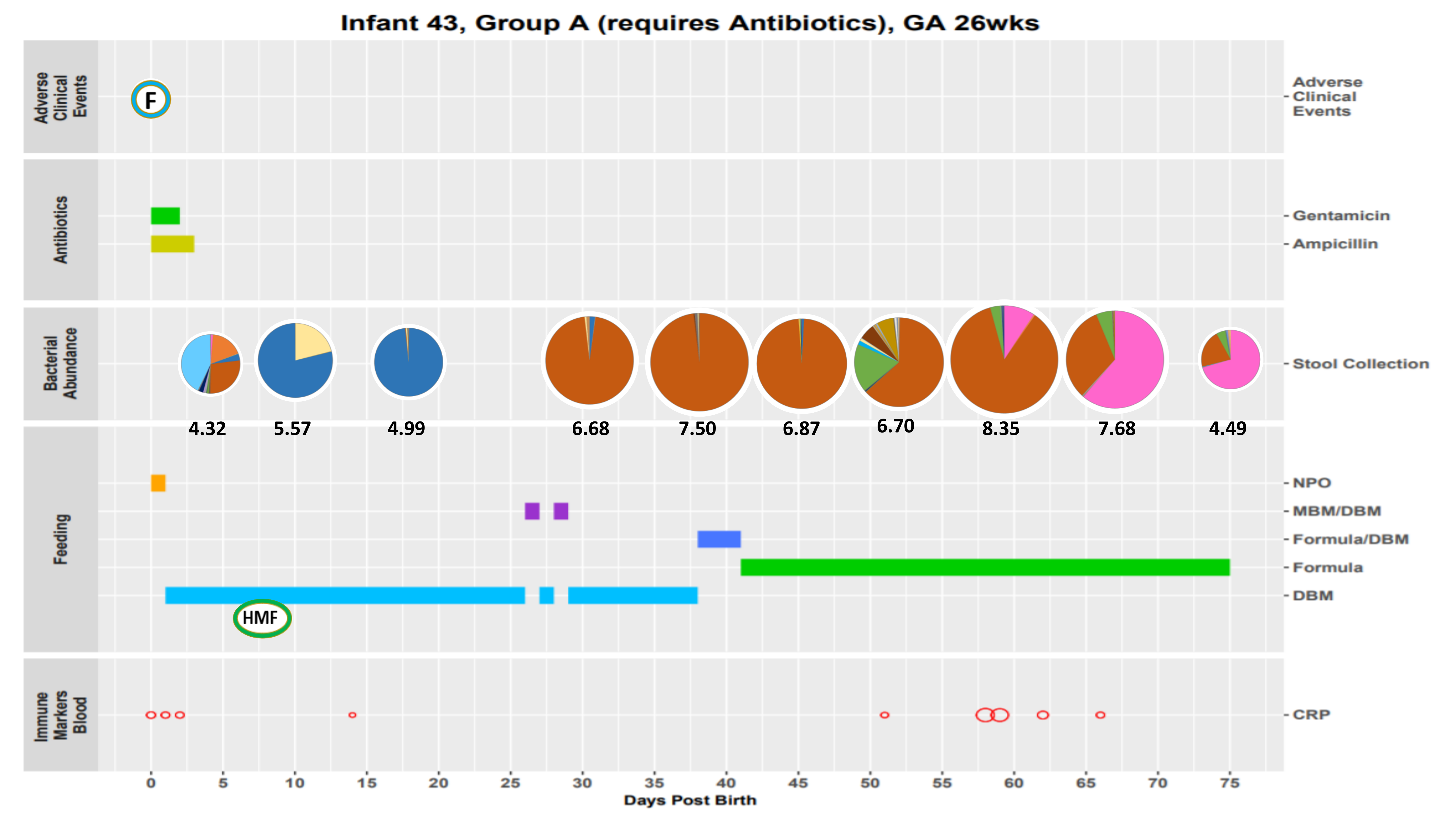
Infant 43, Group A (requires Antibiotics), GA 26wks
Adverse Clinical Events
F
Antibiotics
Gentamicin
Ampicillin
Bacterial Abundance
Stool Collection
4.32
5.57
4.99
6.68
7.50
6.87
6.70
8.35
7.68
4.49
Feeding
NPO
MBM/DBM
Formula/DBM
Formula
DBM
HMF
Immune Markers Blood
CRP
0
5
10
15
20
25
30
35
40
45
50
55
60
65
70
75
Days Post Birth

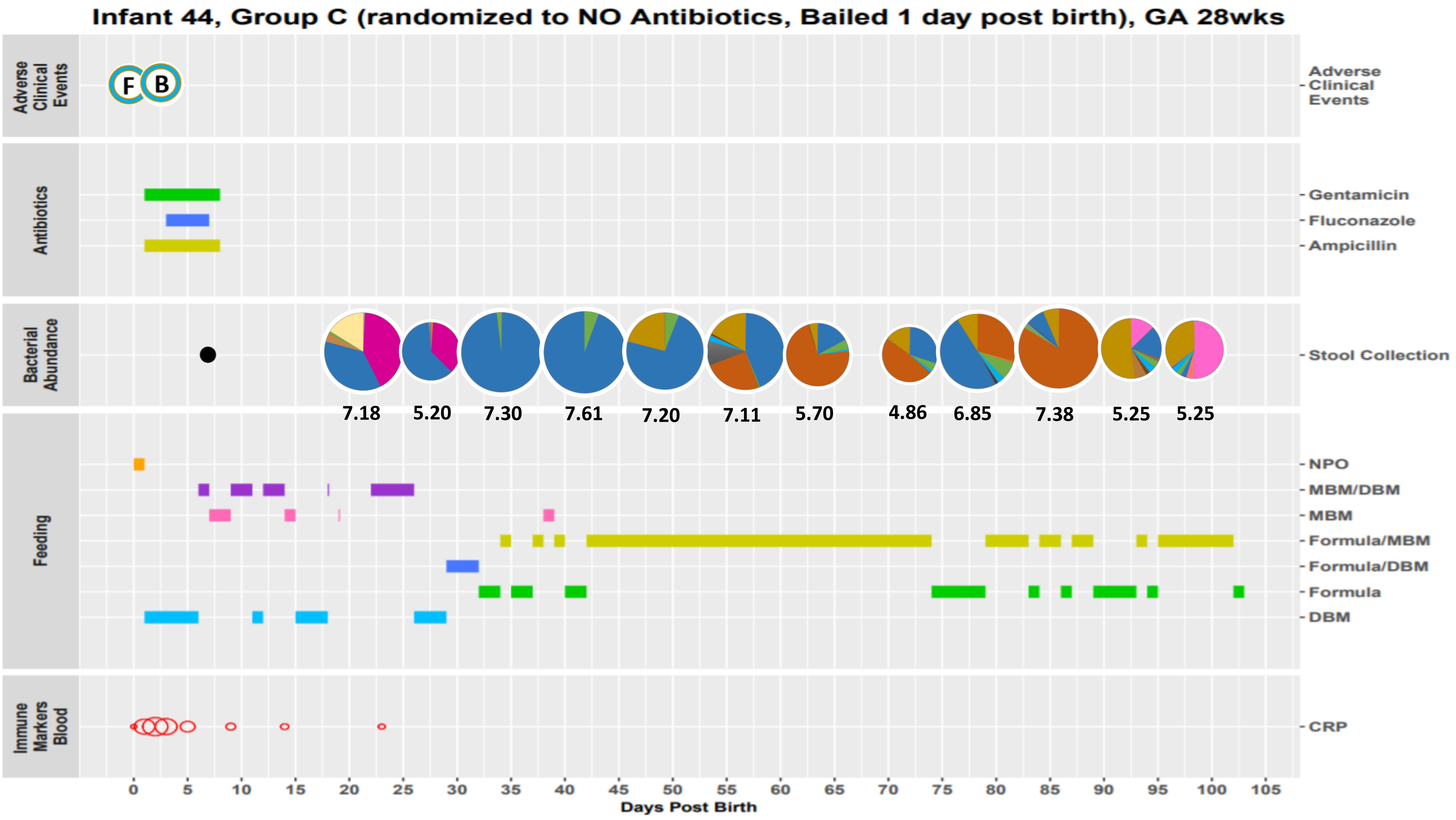

Infant 44, Group C (randomized to NO Antibiotics, Bailed 1 day post birth), GA 28wks
Adverse Clinical Events
F
B
Antibiotics
Gentamicin
Fluconazole
Ampicillin
Bacterial Abundance
Stool Collection
7.18
5.20
7.30
7.61
7.20
7.11
5.70
4.86
6.85
7.38
5.25
5.25
Feeding
NPO
MBM/DBM
MBM
Formula/MBM
Formula/DBM
Formula
DBM
Immune Markers Blood
CRP
0 5 10 15 20 25 30 35 40 45 50 55 60 65 70 75 80 85 90 95 100 105
Days Post Birth

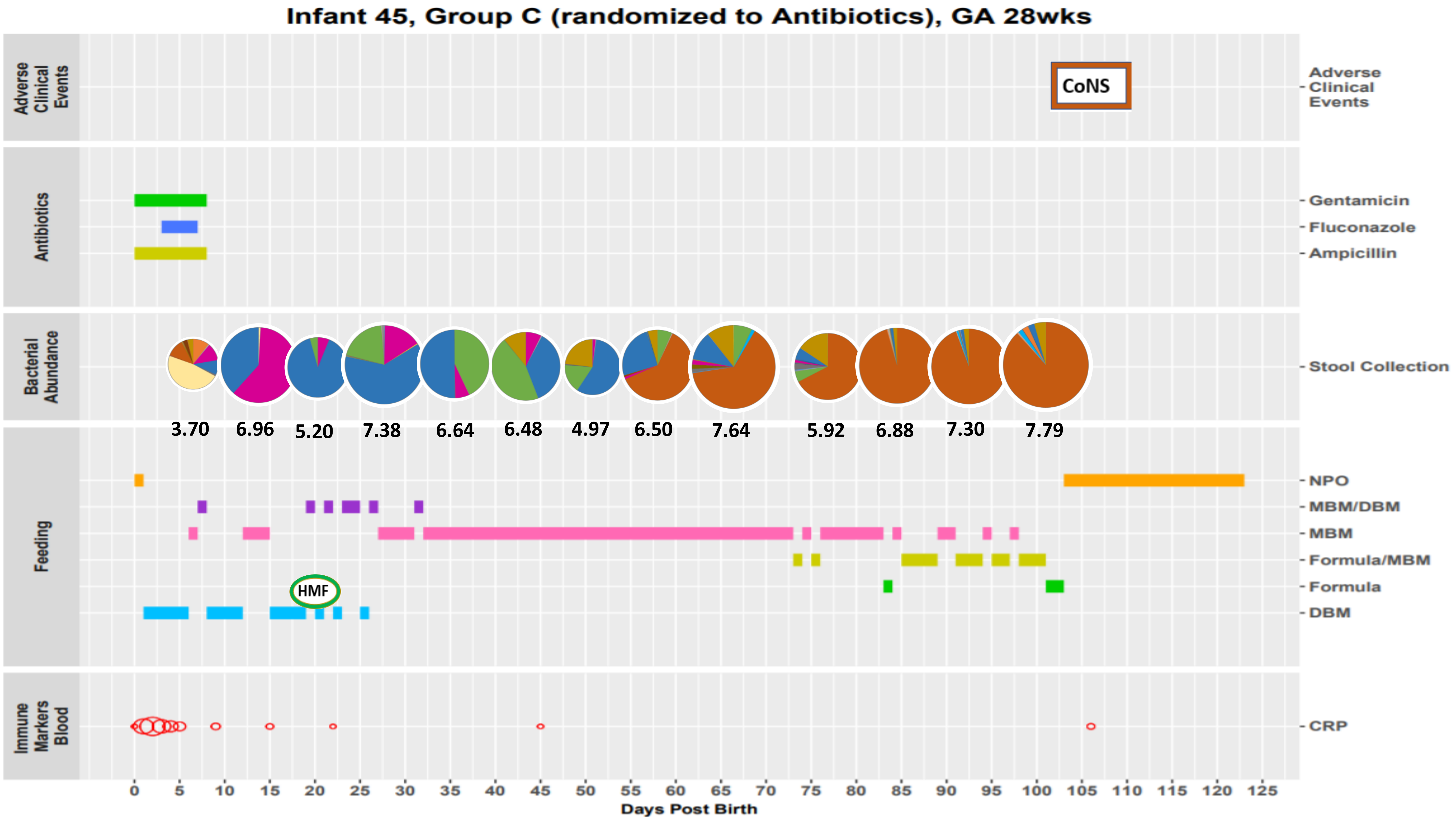

Infant 45, Group C (randomized to Antibiotics), GA 28wks
Adverse Clinical Events
CoNS
Antibiotics
Gentamicin
Fluconazole
Ampicillin
Bacterial Abundance
Stool Collection
3.70
6.96
5.20
7.38
6.64
6.48
4.97
6.50
7.64
5.92
6.88
7.30
7.79
Feeding
NPO
MBM/DBM
MBM
Formula/MBM
Formula
DBM
HMF
Immune Markers Blood
CRP
0 5 10 15 20 25 30 35 40 45 50 55 60 65 70 75 80 85 90 95 100 105 110 115 120 125
Days Post Birth

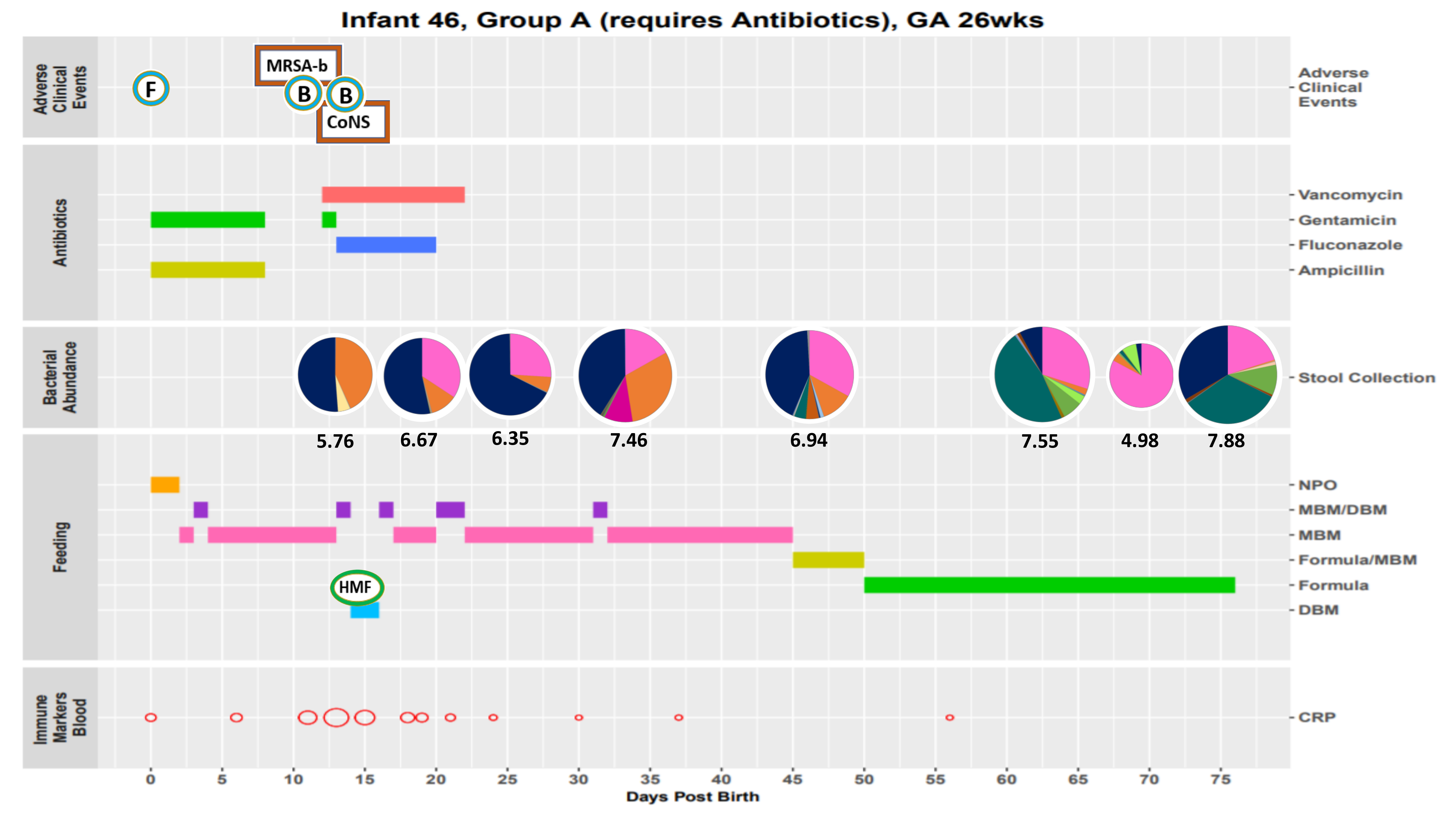

Infant 46, Group A (requires Antibiotics), GA 26wks
Adverse Clinical Events
F
MRSA-b
B
B
CoNS
Antibiotics
Vancomycin
Gentamicin
Fluconazole
Ampicillin
Bacterial Abundance
Stool Collection
5.76
6.67
6.35
7.46
6.94
7.55
4.98
7.88
Feeding
NPO
MBM/DBM
MBM
Formula/MBM
Formula
DBM
HMF
Immune Markers Blood
CRP
0
5
10
15
20
25
30
35
40
45
50
55
60
65
70
75
Days Post Birth

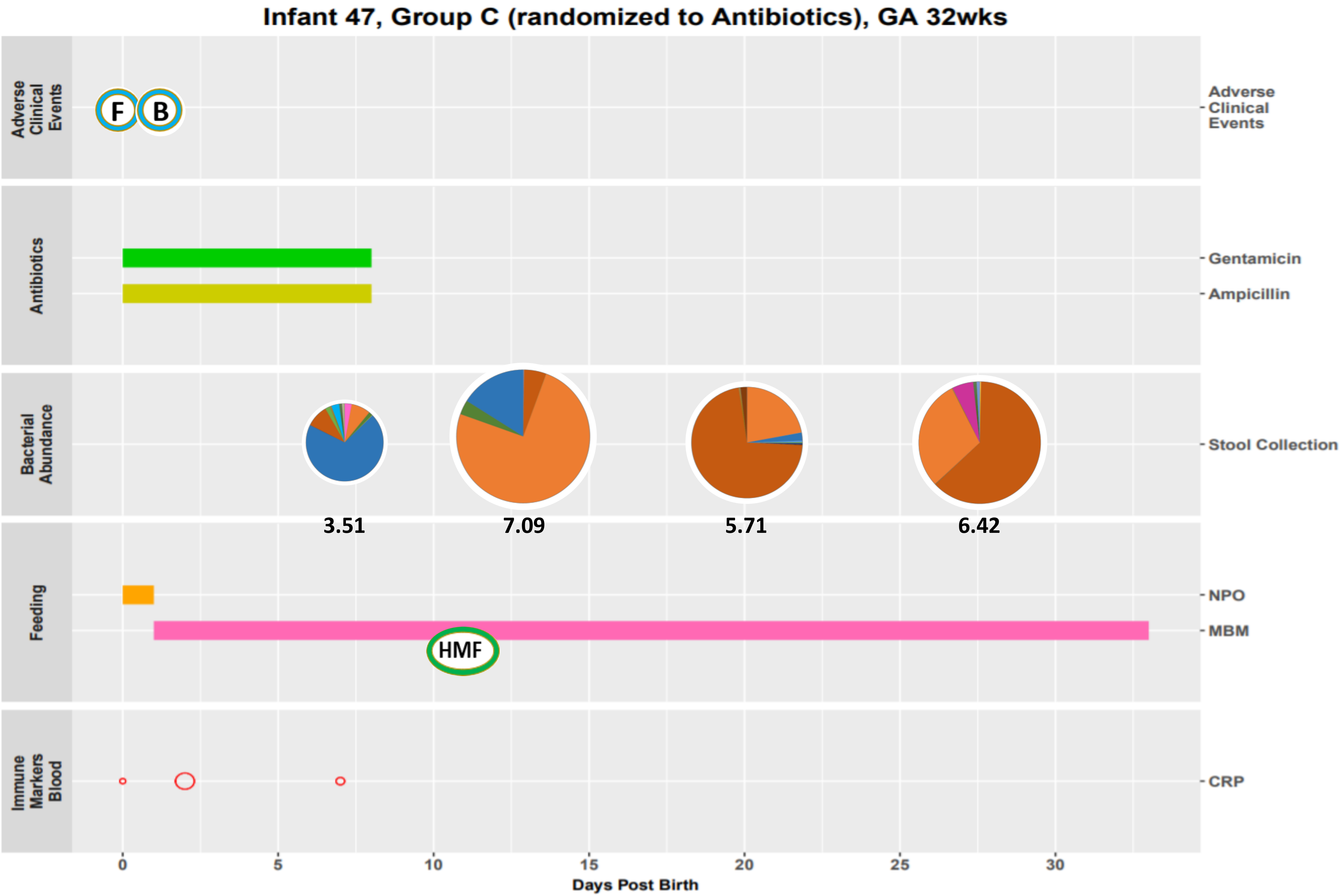
Infant 47, Group C (randomized to Antibiotics), GA 32wks
Adverse Clinical Events
F
B
Antibiotics
Gentamicin
Ampicillin
Bacterial Abundance
Stool Collection
3.51
7.09
5.71
6.42
Feeding
NPO
MBM
HMF
Immune Markers Blood
CRP
0
5
10
15
20
25
30
Days Post Birth

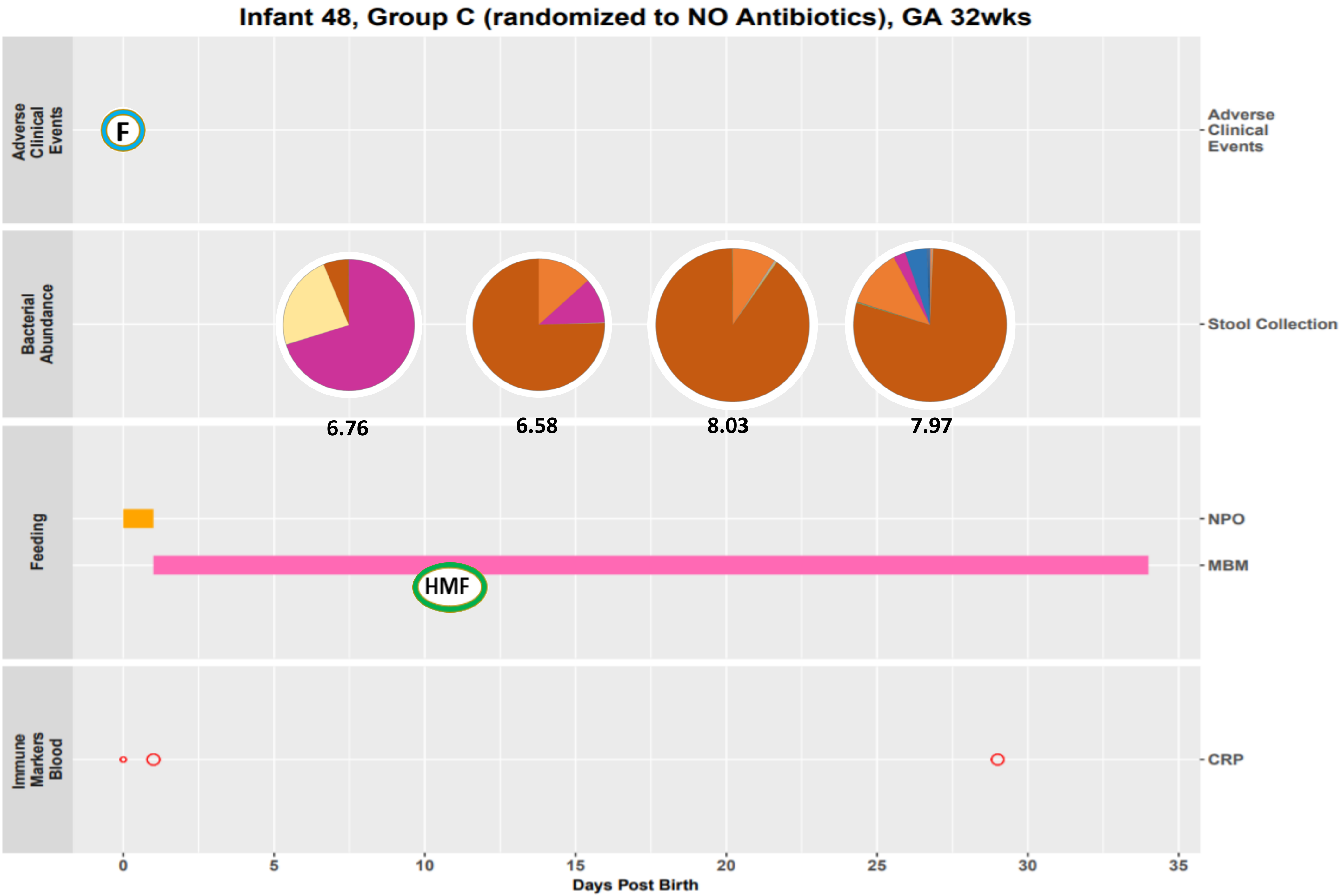
Infant 48, Group C (randomized to NO Antibiotics), GA 32wks
Adverse Clinical Events
F
Bacterial Abundance
Stool Collection
6.76
6.58
8.03
7.97
Feeding
NPO
MBM
HMF
Immune Markers Blood
CRP
0
5
10
15
20
25
30
35
Days Post Birth

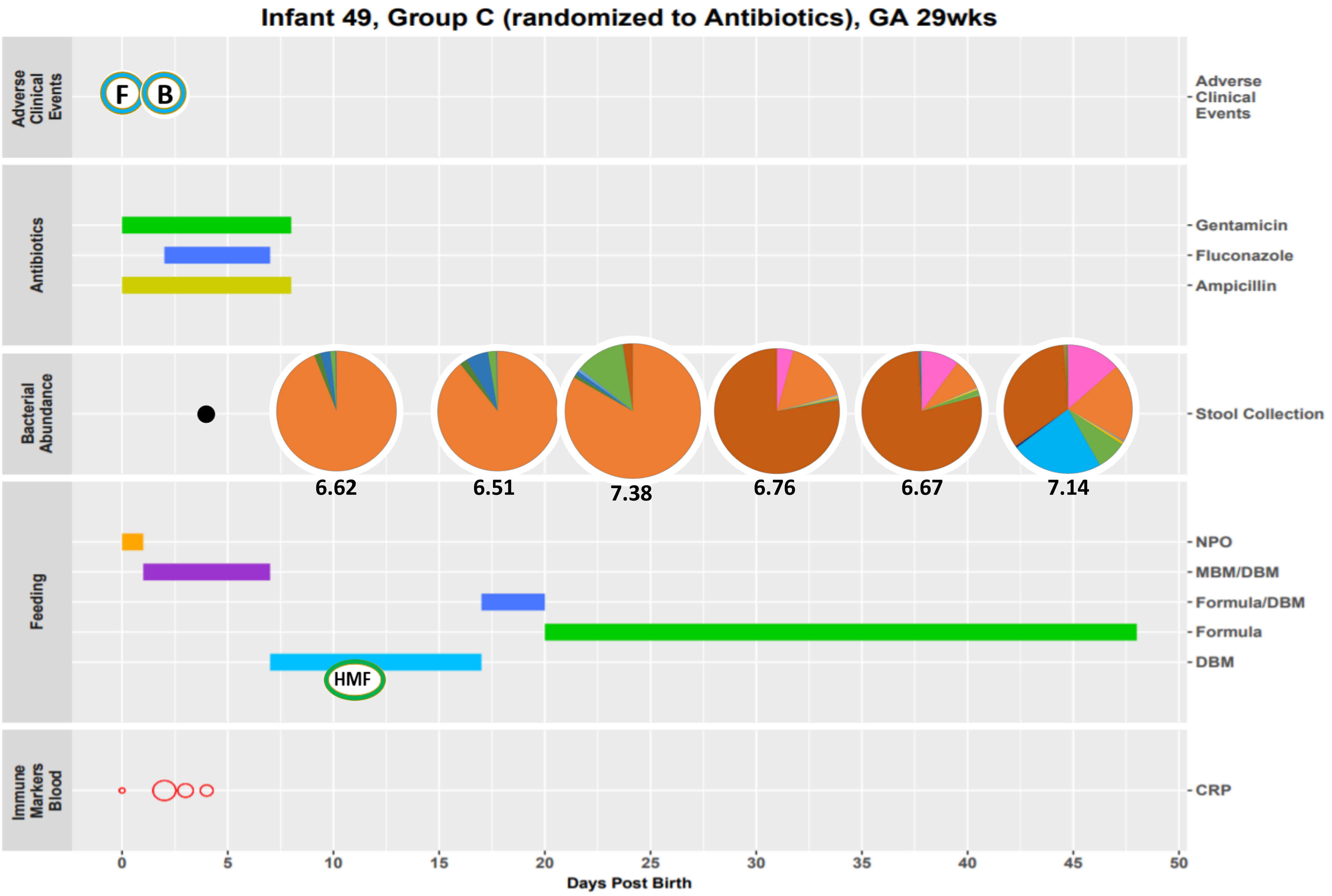

Infant 49, Group C (randomized to Antibiotics), GA 29wks
Adverse Clinical Events
F
B
Antibiotics
Gentamicin
Fluconazole
Ampicillin
Bacterial Abundance
Stool Collection
6.62
6.51
7.38
6.76
6.67
7.14
Feeding
NPO
MBM/DBM
Formula/DBM
Formula
DBM
HMF
Immune Markers Blood
CRP
0
5
10
15
20
25
30
35
40
45
50
Days Post Birth

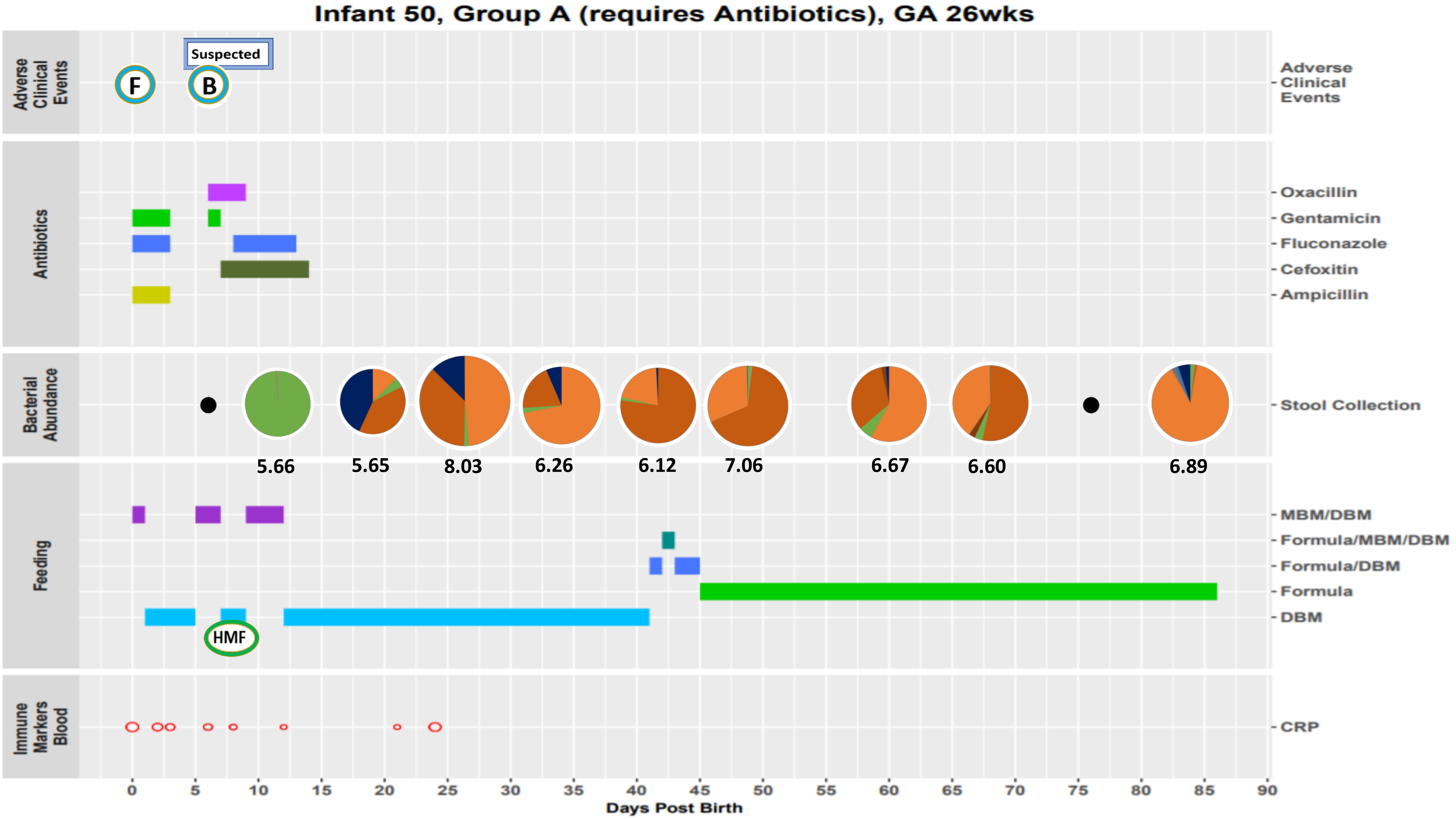
Infant 50, Group A (requires Antibiotics), GA 26wks
Adverse Clinical Events
Suspected
F
B
Antibiotics
Oxacillin
Gentamicin
Fluconazole
Cefoxitin
Ampicillin
Bacterial Abundance
Stool Collection
5.66
5.65
8.03
6.26
6.12
7.06
6.67
6.60
6.89
Feeding
MBM/DBM
Formula/MBM/DBM
Formula/DBM
Formula
DBM
HMF
Immune Markers Blood
CRP
0
5
10
15
20
25
30
35
40
45
50
55
60
65
70
75
80
85
90
Days Post Birth

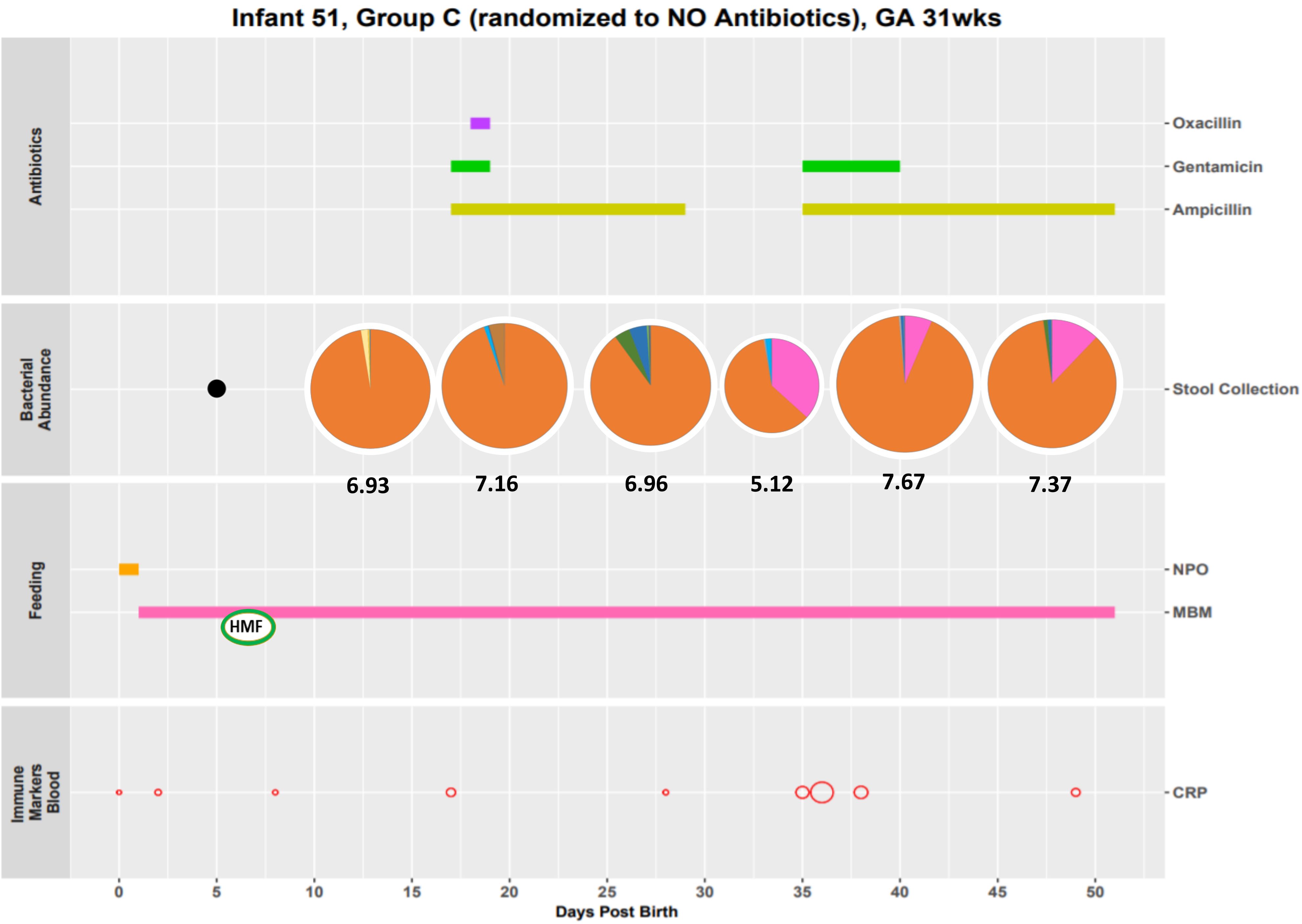
Infant 51, Group C (randomized to NO Antibiotics), GA 31wks
Antibiotics
Oxacillin
Gentamicin
Ampicillin
Bacterial Abundance
Stool Collection
6.93
7.16
6.96
5.12
7.67
7.37
Feeding
NPO
MBM
HMF
Immune Markers Blood
CRP
0
5
10
15
20
25
30
35
40
45
50
Days Post Birth

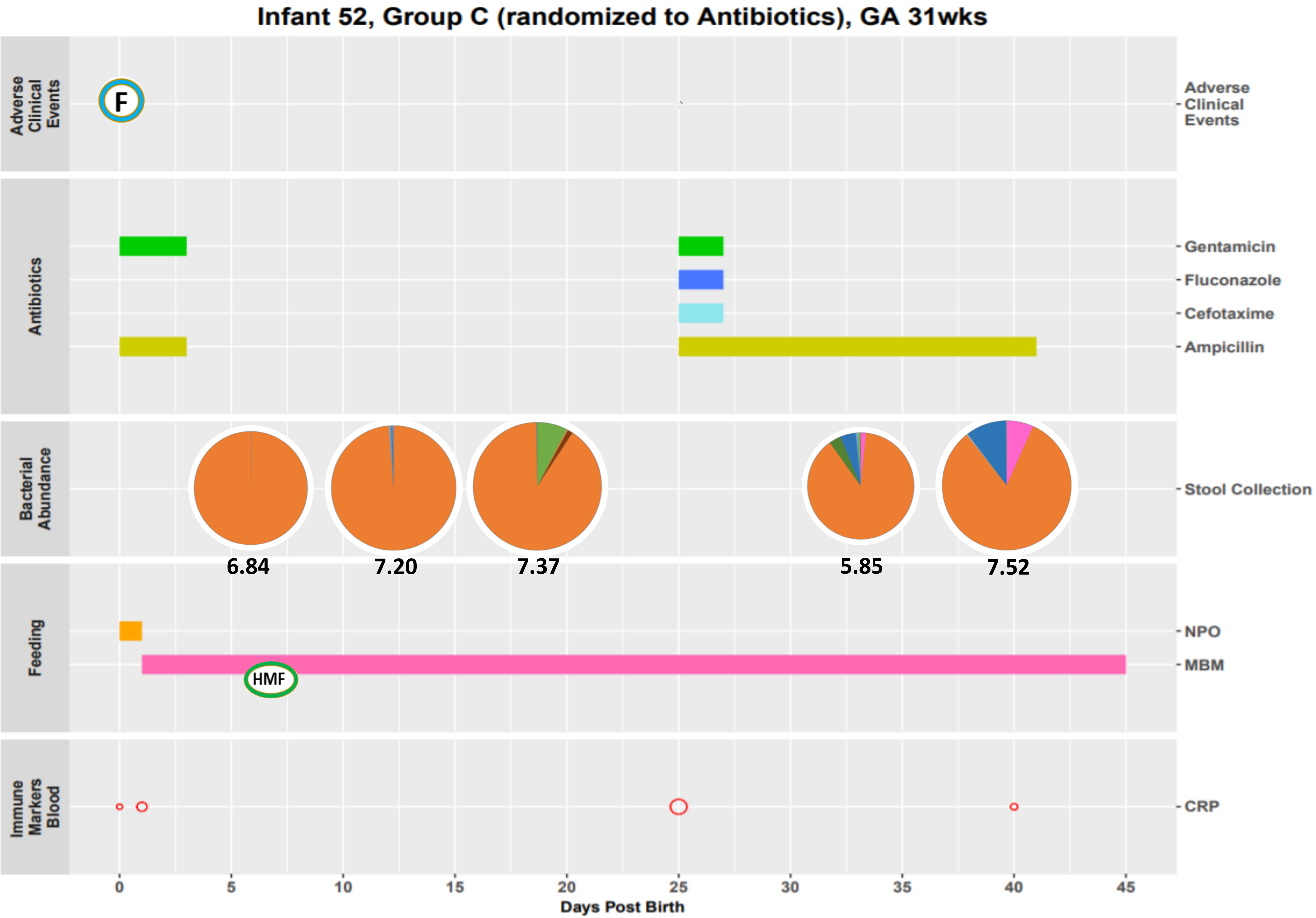
Infant 52, Group C (randomized to Antibiotics), GA 31wks
Adverse Clinical Events
F
Antibiotics
Gentamicin
Fluconazole
Cefotaxime
Ampicillin
Bacterial Abundance
Stool Collection
6.84
7.20
7.37
5.85
7.52
Feeding
NPO
MBM
HMF
Immune Markers Blood
CRP
0
5
10
15
20
25
30
35
40
45
Days Post Birth

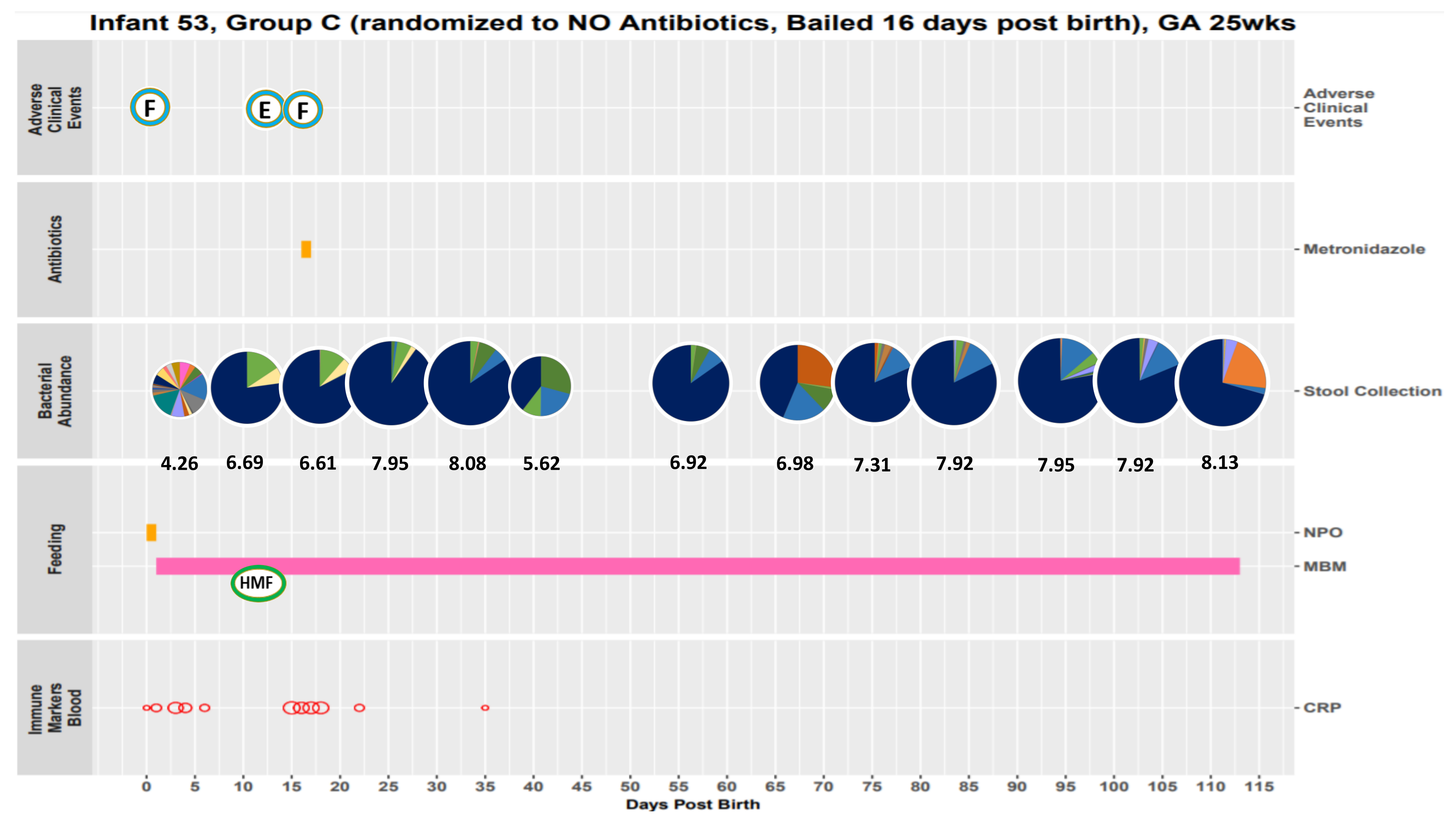
Infant 53, Group C (randomized to NO Antibiotics, Bailed 16 days post birth), GA 25wks
Adverse Clinical Events
F
E
F
Antibiotics
Metronidazole
Bacterial Abundance
Stool Collection
4.26
6.69
6.61
7.95
8.08
5.62
6.92
6.98
7.31
7.92
7.95
7.92
8.13
Feeding
NPO
MBM
HMF
Immune Markers Blood
CRP
0 5 10 15 20 25 30 35 40 45 50 55 60 65 70 75 80 85 90 95 100 105 110 115
Days Post Birth

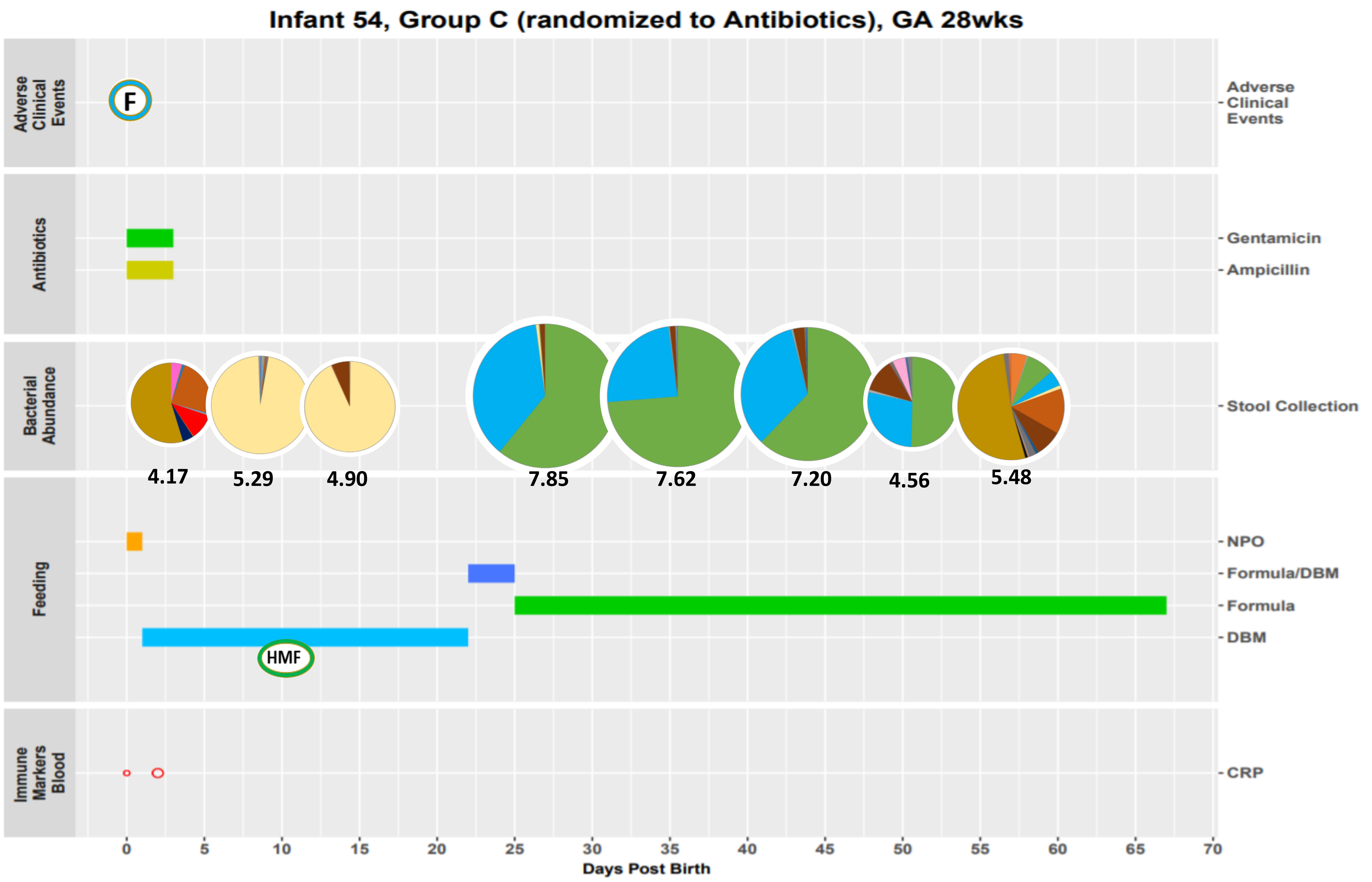

Infant 54, Group C (randomized to Antibiotics), GA 28wks
Adverse Clinical Events
F
Antibiotics
Gentamicin
Ampicillin
Bacterial Abundance
Stool Collection
4.17
5.29
4.90
7.85
7.62
7.20
4.56
5.48
Feeding
NPO
Formula/DBM
Formula
DBM
HMF
Immune Markers Blood
CRP
0
5
10
15
20
25
30
35
40
45
50
55
60
65
70
Days Post Birth

Infant 55, Group C (randomized to NO Antibiotics), GA 30wks

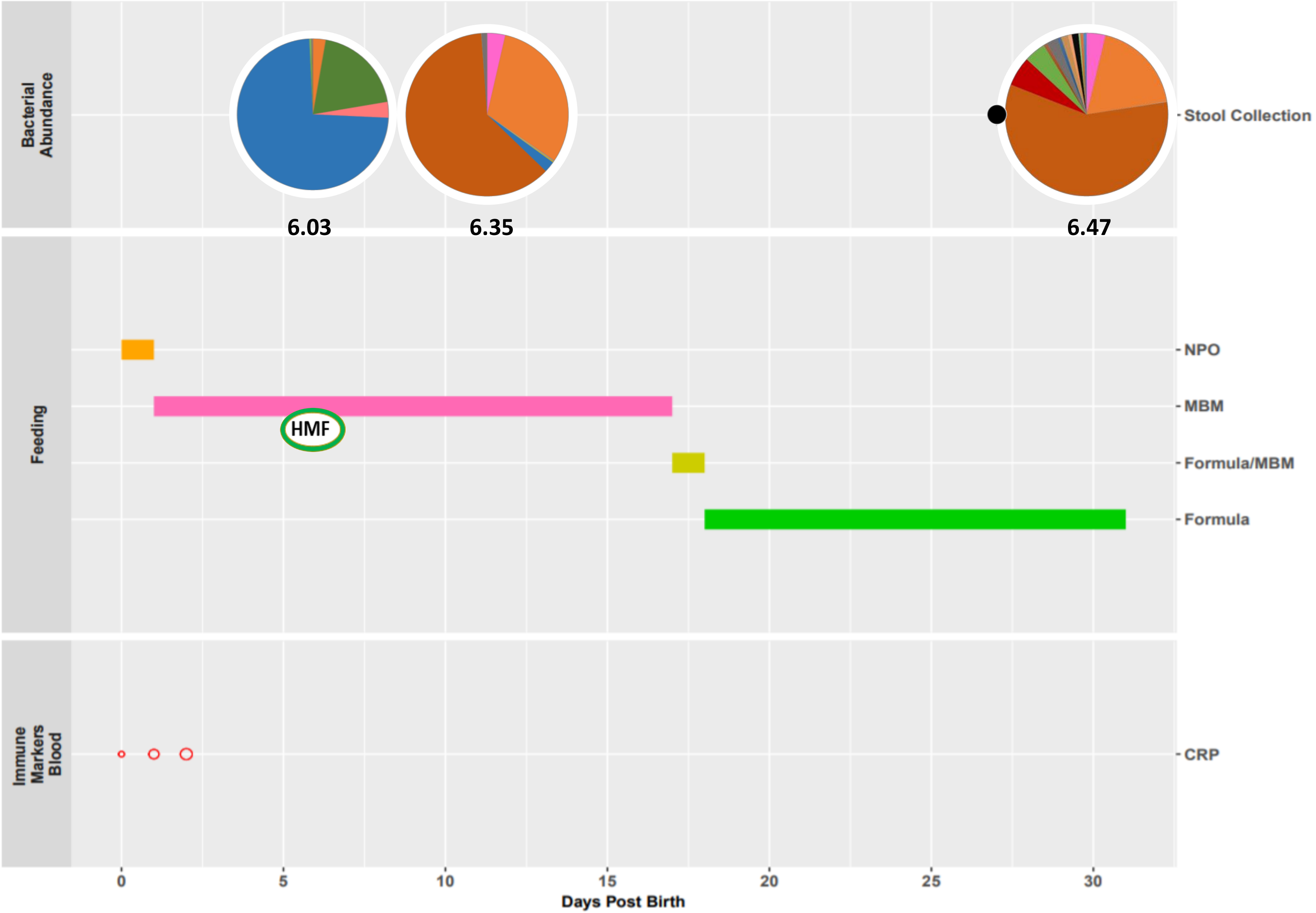

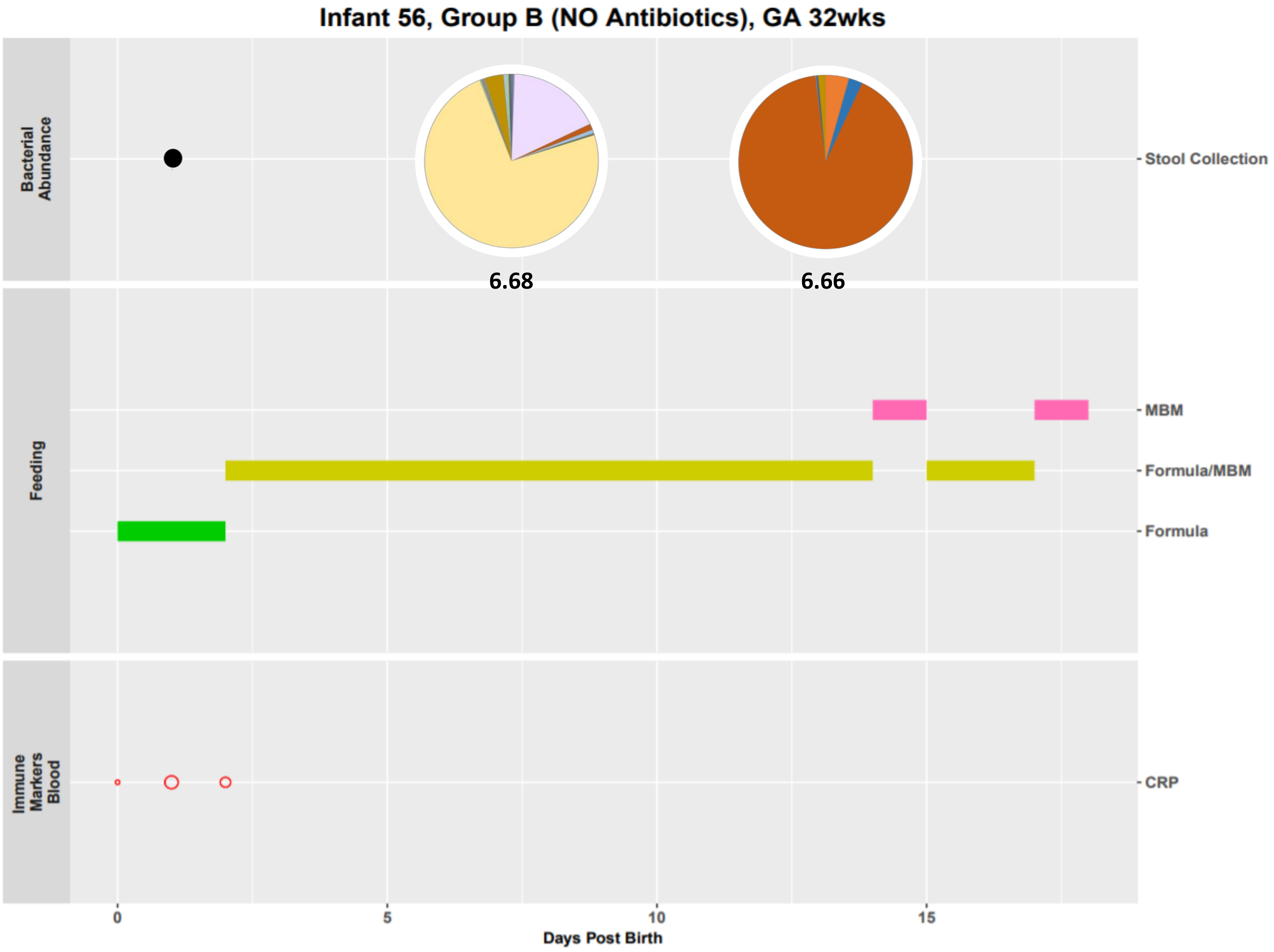
Infant 56, Group B (NO Antibiotics), GA 32wks
Bacterial Abundance
Feeding
Immune Markers Blood
Stool Collection
MBM
Formula/MBM
Formula
CRP
6.68
6.66
0
5
10
15
Days Post Birth

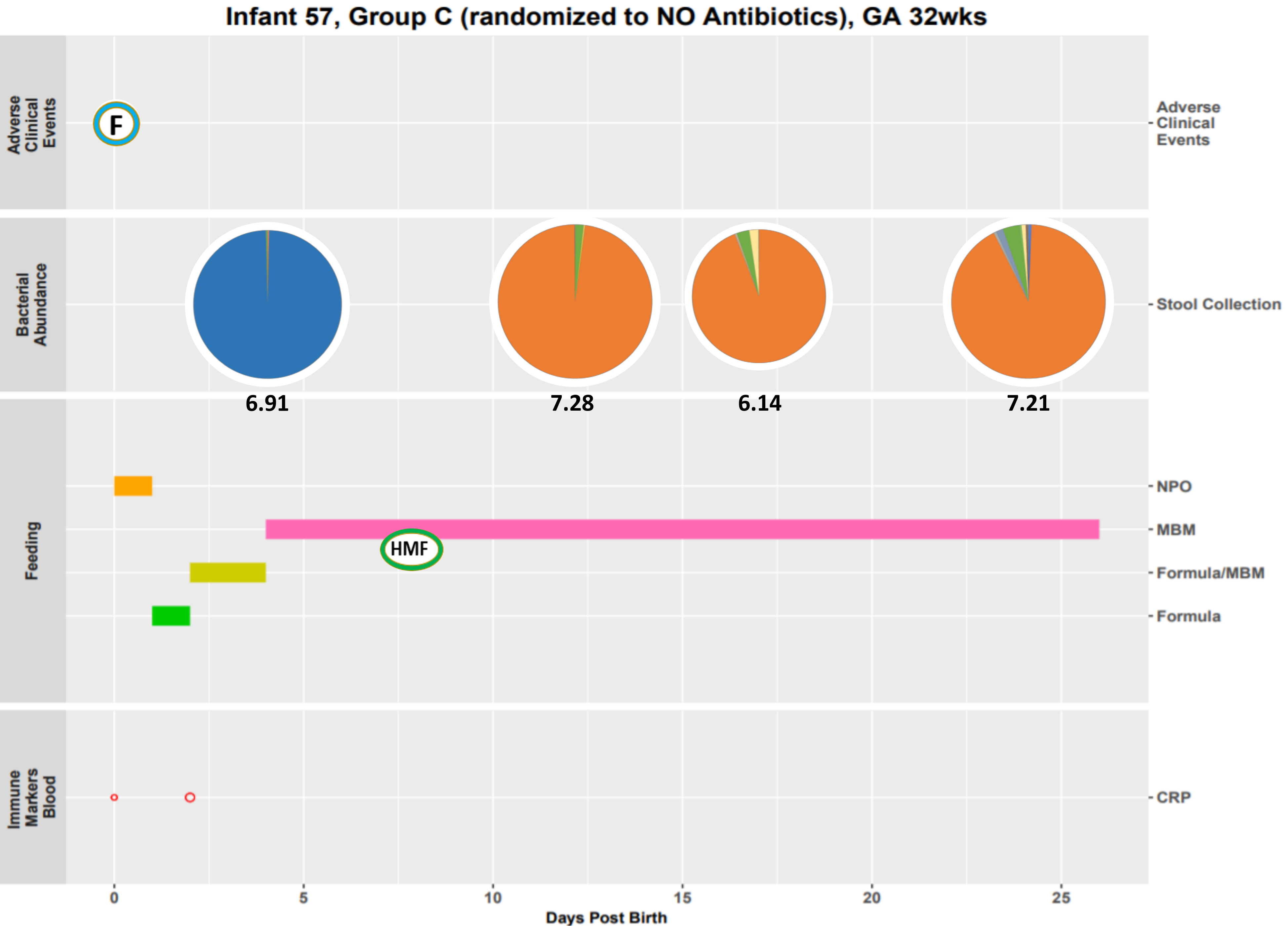

Infant 57, Group C (randomized to NO Antibiotics), GA 32wks
Adverse Clinical Events
F
Adverse Clinical Events
Bacterial Abundance
Stool Collection
6.91
7.28
6.14
7.21
Feeding
NPO
MBM
HMF
Formula/MBM
Formula
Immune Markers Blood
CRP
0
5
10
15
20
25
Days Post Birth

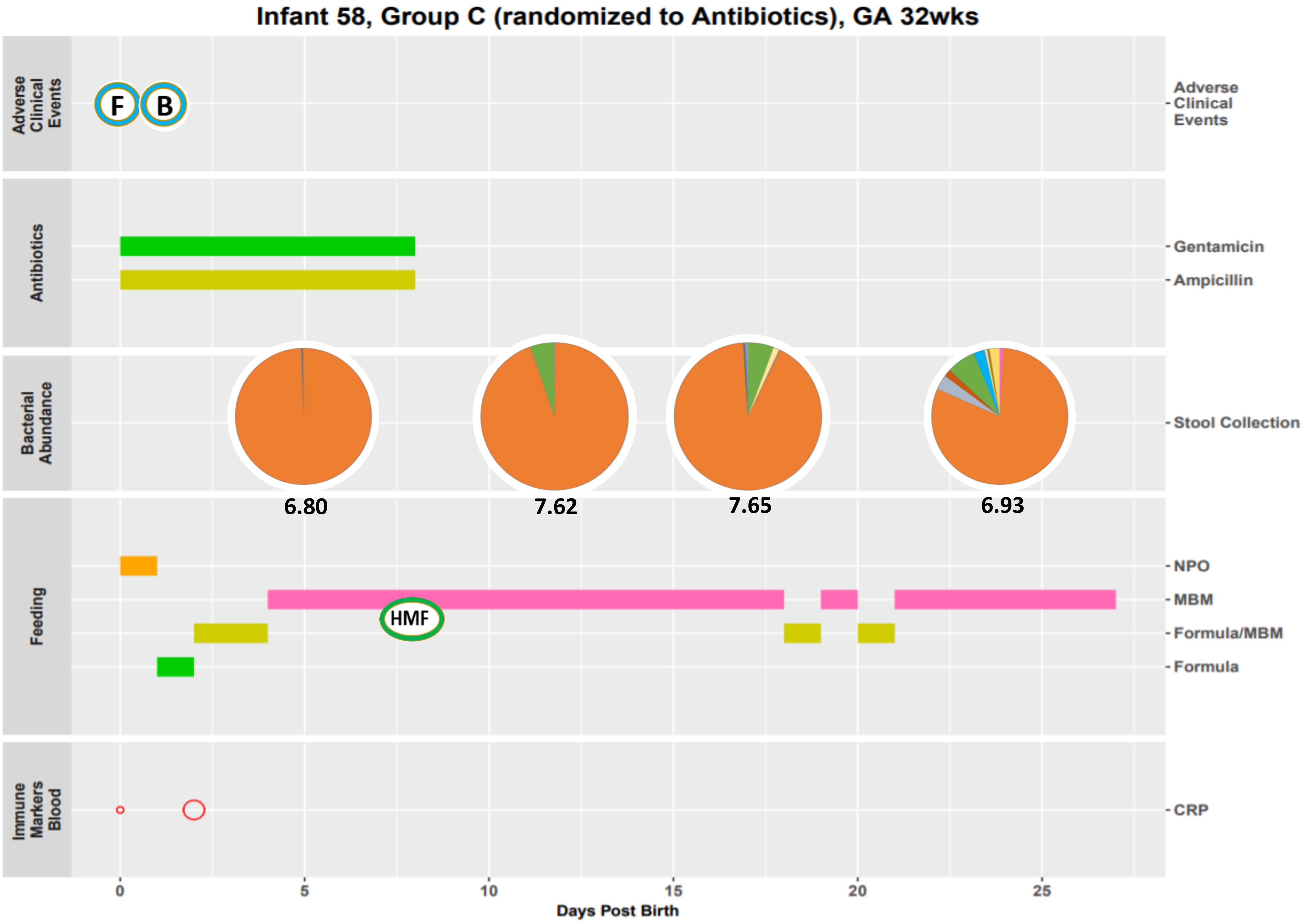
Infant 58, Group C (randomized to Antibiotics), GA 32wks
Adverse Clinical Events
F
B
Antibiotics
Gentamicin
Ampicillin
Bacterial Abundance
Stool Collection
6.80
7.62
7.65
6.93
Feeding
NPO
MBM
HMF
Formula/MBM
Formula
Immune Markers Blood
CRP
0
5
10
15
20
25
Days Post Birth

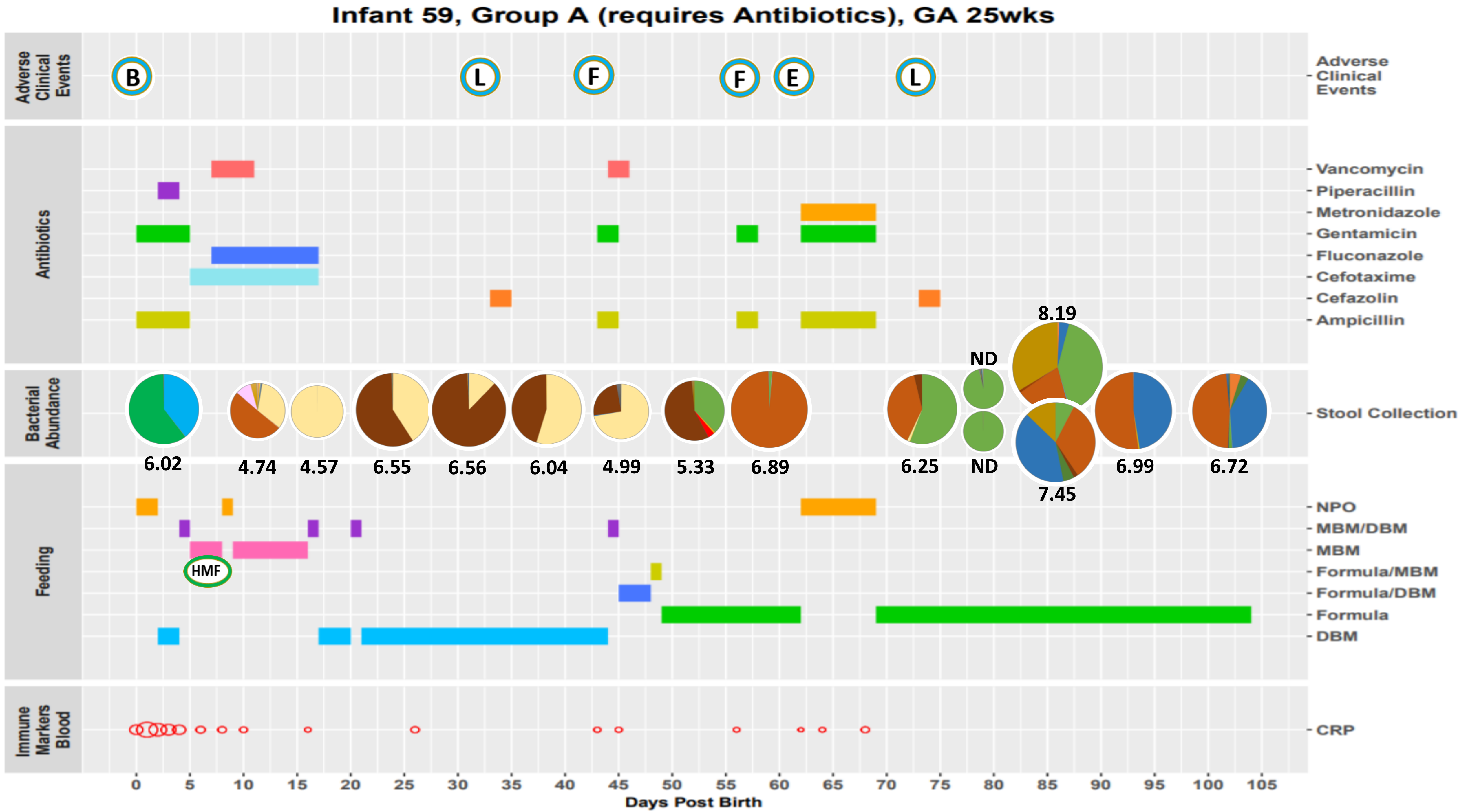

Infant 59, Group A (requires Antibiotics), GA 25wks
Adverse Clinical Events
B
L
F
F
E
L
Antibiotics
Vancomycin
Piperacillin
Metronidazole
Gentamicin
Fluconazole
Cefotaxime
Cefazolin
Ampicillin
Bacterial Abundance
Stool Collection
6.02
4.74
4.57
6.55
6.56
6.04
4.99
5.33
6.89
6.25
ND
ND
8.19
7.45
6.99
6.72
Feeding
NPO
MBM/DBM
MBM
HMF
Formula/MBM
Formula/DBM
Formula
DBM
Immune Markers Blood
CRP
0 5 10 15 20 25 30 35 40 45 50 55 60 65 70 75 80 85 90 95 100 105
Days Post Birth

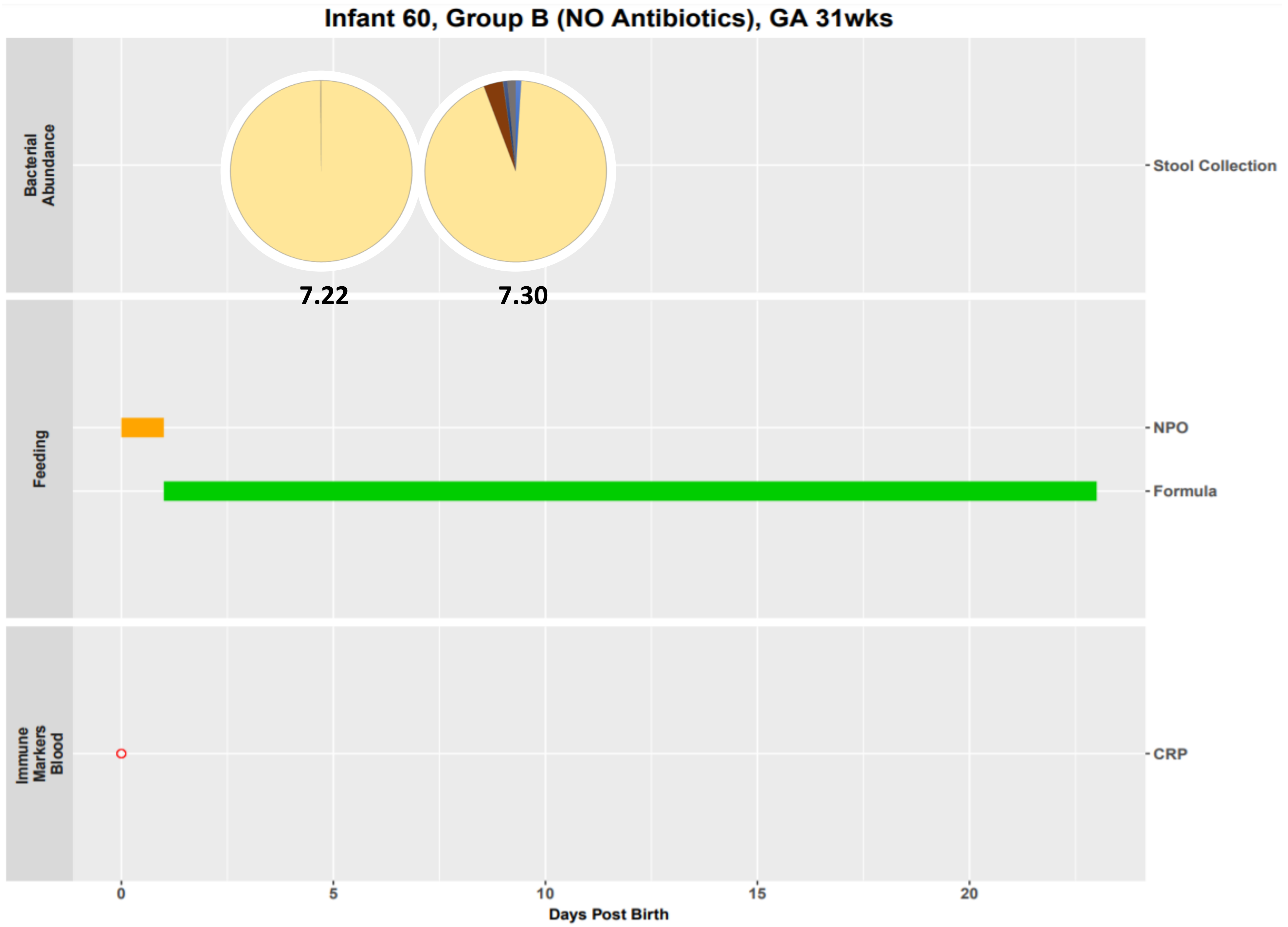

Infant 60, Group B (NO Antibiotics), GA 31wks
Bacterial Abundance
7.22
7.30
Stool Collection
Feeding
NPO
Formula
Immune Markers Blood
CRP
0
5
10
15
20
Days Post Birth

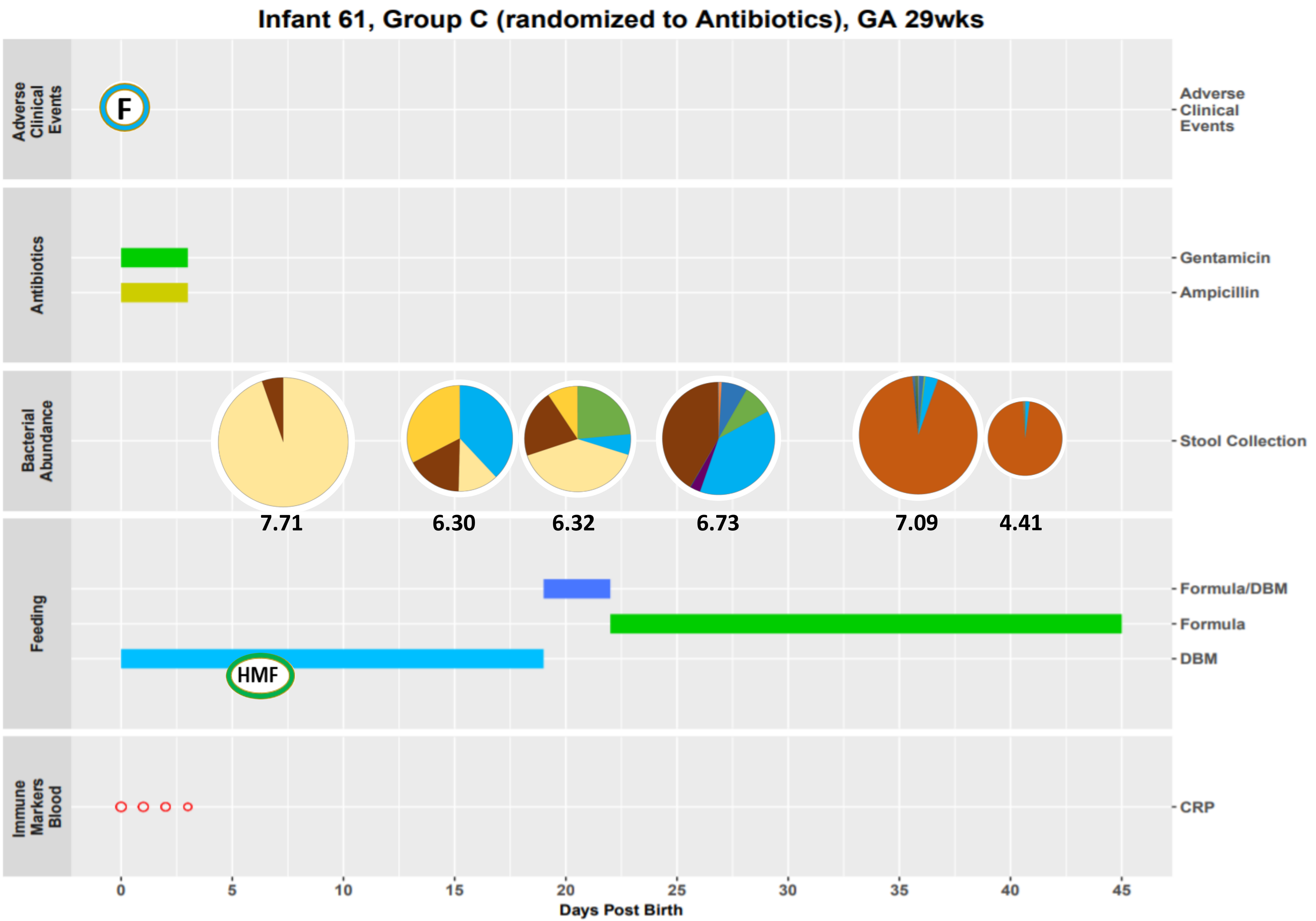

Infant 61, Group C (randomized to Antibiotics), GA 29wks
Adverse Clinical Events
F
Antibiotics
Gentamicin
Ampicillin
Bacterial Abundance
Stool Collection
7.71
6.30
6.32
6.73
7.09
4.41
Feeding
Formula/DBM
Formula
DBM
HMF
Immune Markers Blood
CRP
0
5
10
15
20
25
30
35
40
45
Days Post Birth

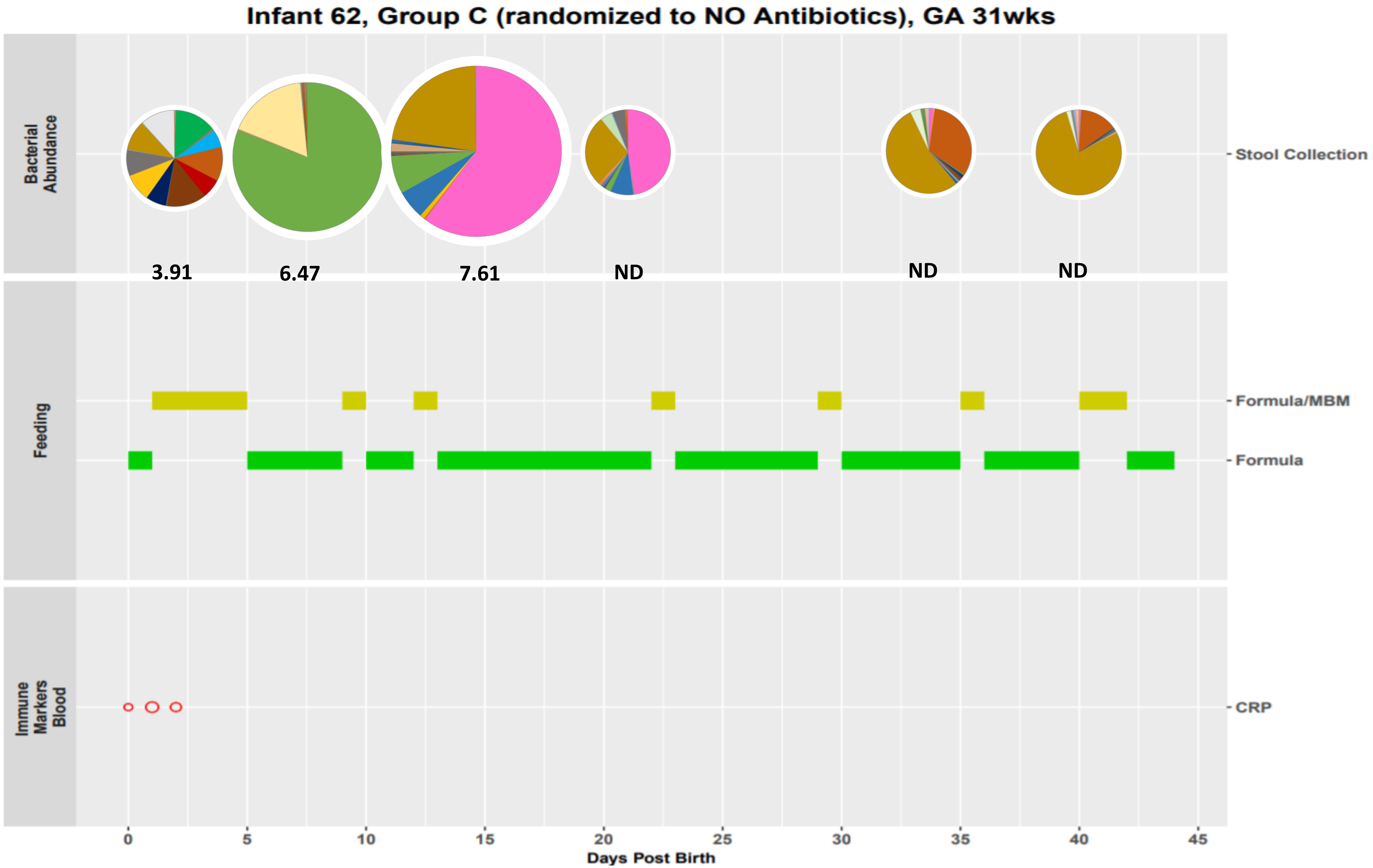
Infant 62, Group C (randomized to NO Antibiotics), GA 31wks
Bacterial Abundance
3.91
6.47
7.61
ND
ND
ND
Stool Collection
Feeding
Formula/MBM
Formula
Immune Markers Blood
CRP
0
5
10
15
20
25
30
35
40
45
Days Post Birth

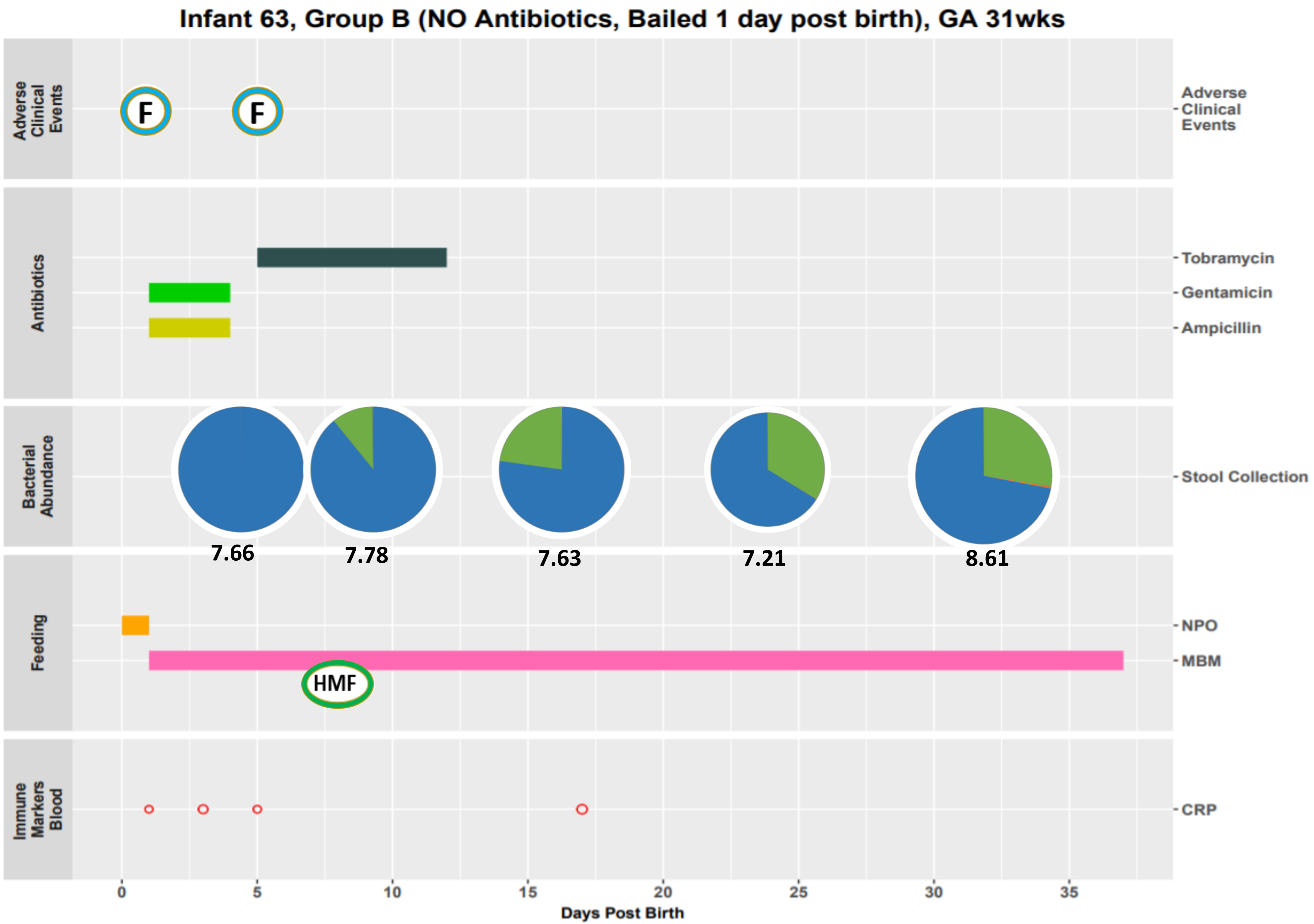

Infant 63, Group B (NO Antibiotics, Bailed 1 day post birth), GA 31wks
Adverse Clinical Events
F
F
Antibiotics
Tobramycin
Gentamicin
Ampicillin
Bacterial Abundance
Stool Collection
7.66
7.78
7.63
7.21
8.61
Feeding
NPO
MBM
HMF
Immune Markers Blood
CRP
0
5
10
15
20
25
30
35
Days Post Birth

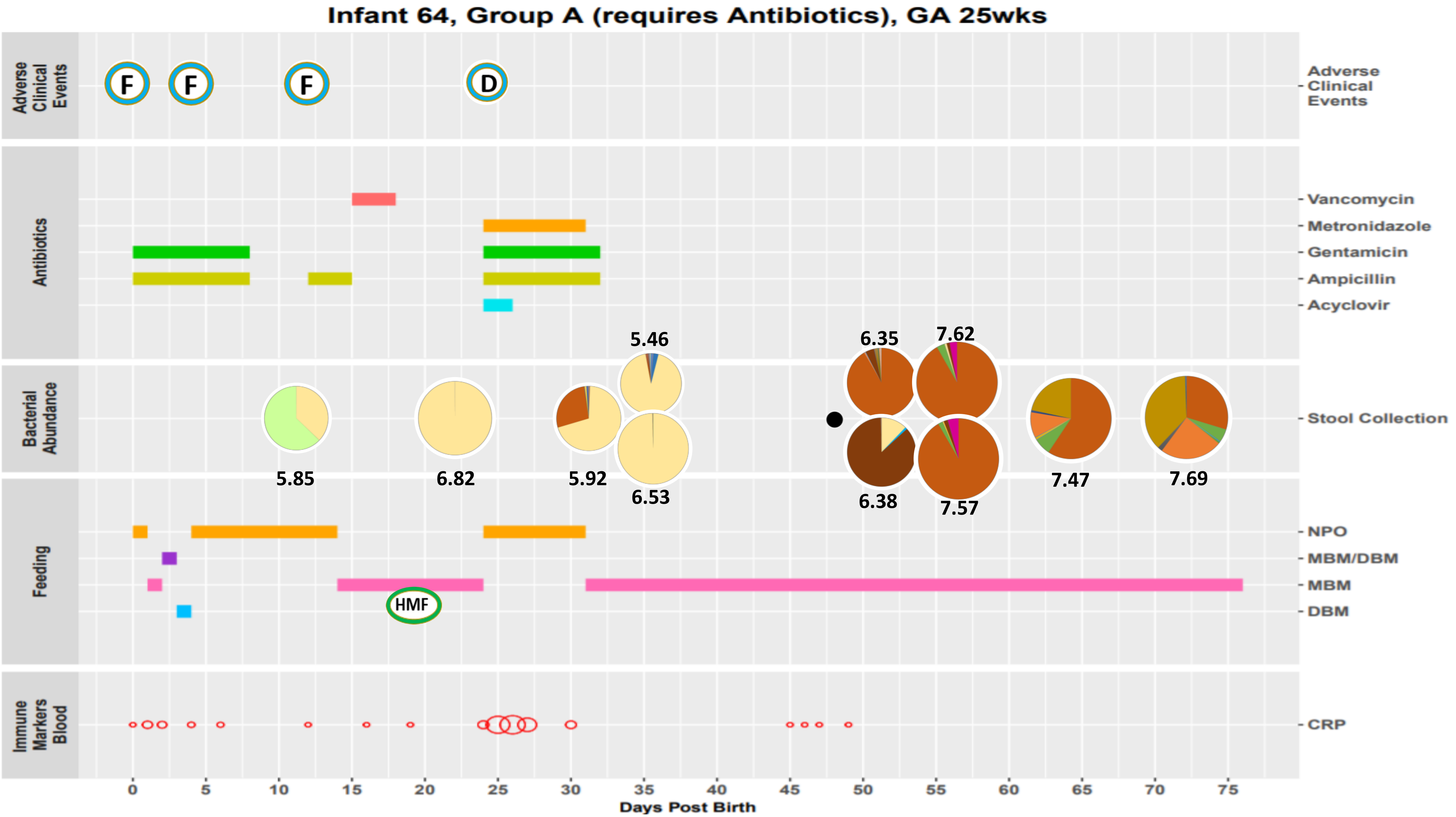
Infant 64, Group A (requires Antibiotics), GA 25wks
Adverse Clinical Events
F
F
F
D
Antibiotics
Vancomycin
Metronidazole
Gentamicin
Ampicillin
Acyclovir
Bacterial Abundance
Stool Collection
5.85
6.82
5.92
5.46
6.53
6.35
6.38
7.62
7.57
7.47
7.69
Feeding
NPO
MBM/DBM
MBM
DBM
HMF
Immune Markers Blood
CRP
0
5
10
15
20
25
30
35
40
45
50
55
60
65
70
75
Days Post Birth

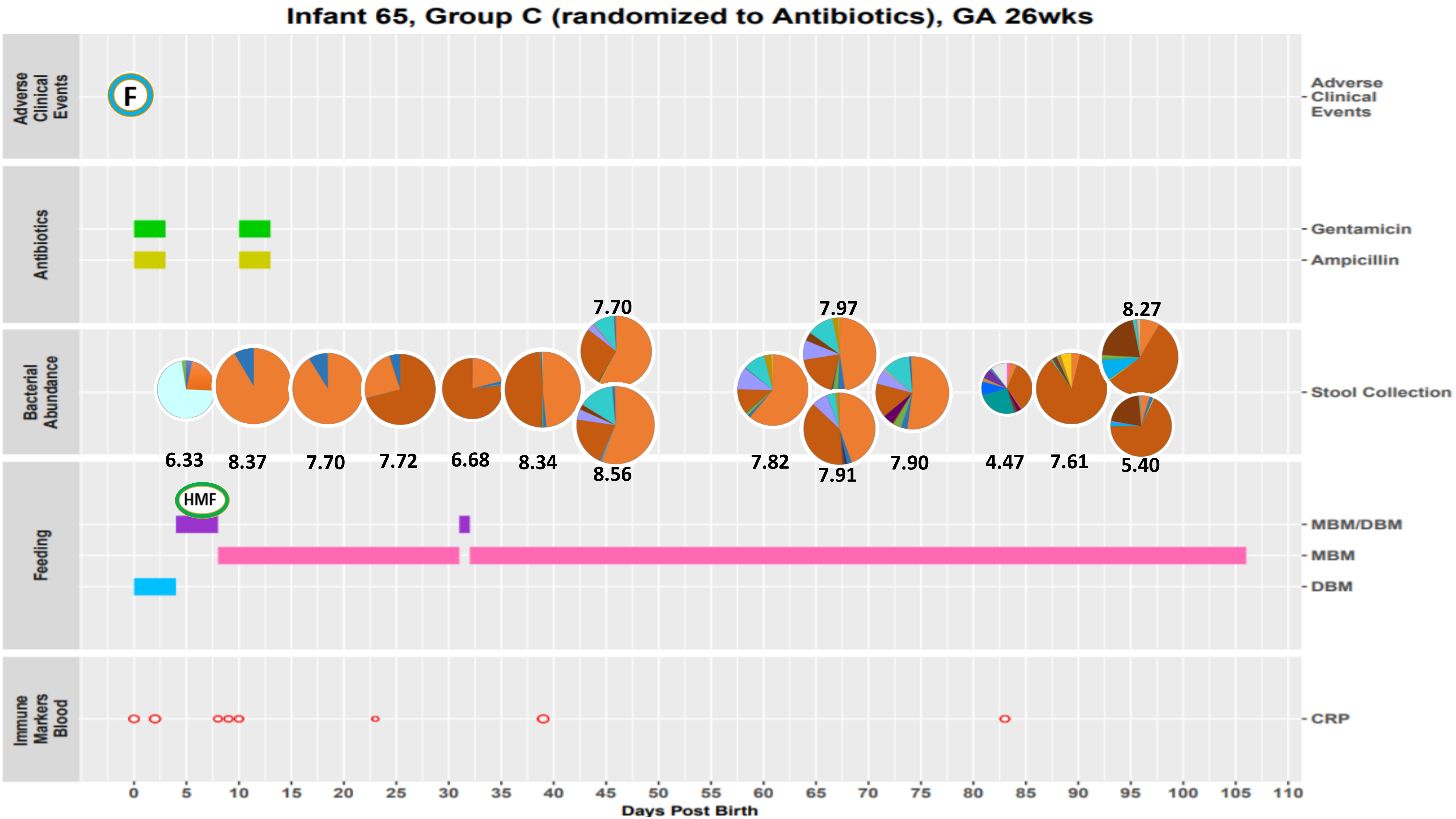
Infant 65, Group C (randomized to Antibiotics), GA 26wks
Adverse Clinical Events
F
Antibiotics
Gentamicin
Ampicillin
Bacterial Abundance
Stool Collection
6.33
8.37
7.70
7.72
6.68
8.34
7.70
8.56
7.82
7.97
7.91
7.90
4.47
7.61
8.27
5.40
Feeding
HMF
MBM/DBM
MBM
DBM
Immune Markers Blood
CRP
0 5 10 15 20 25 30 35 40 45 50 55 60 65 70 75 80 85 90 95 100 105 110
Days Post Birth

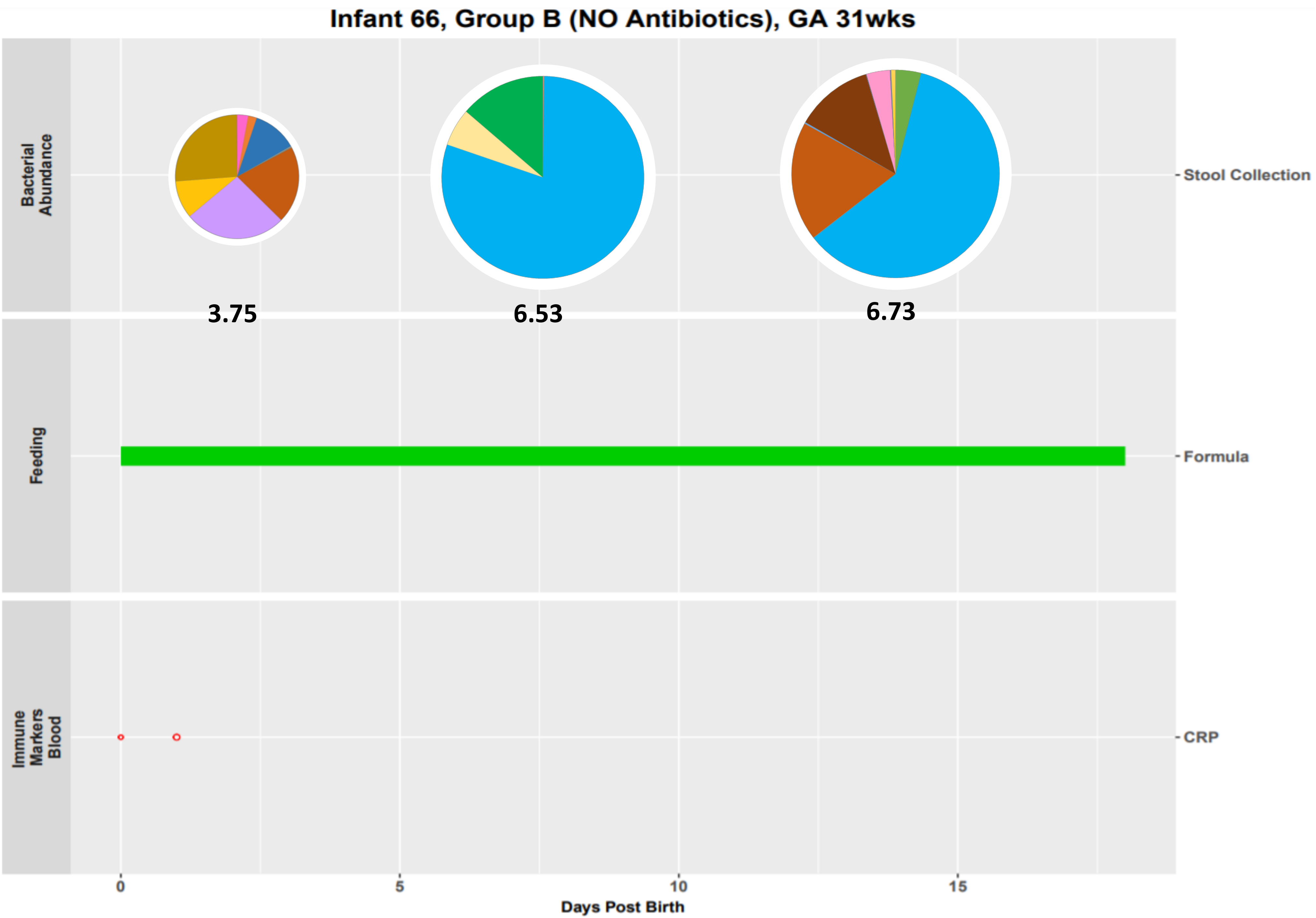
Infant 66, Group B (NO Antibiotics), GA 31wks
Bacterial Abundance
Stool Collection
3.75
6.53
6.73
Feeding
Formula
Immune Markers Blood
CRP
0
5
10
15
Days Post Birth

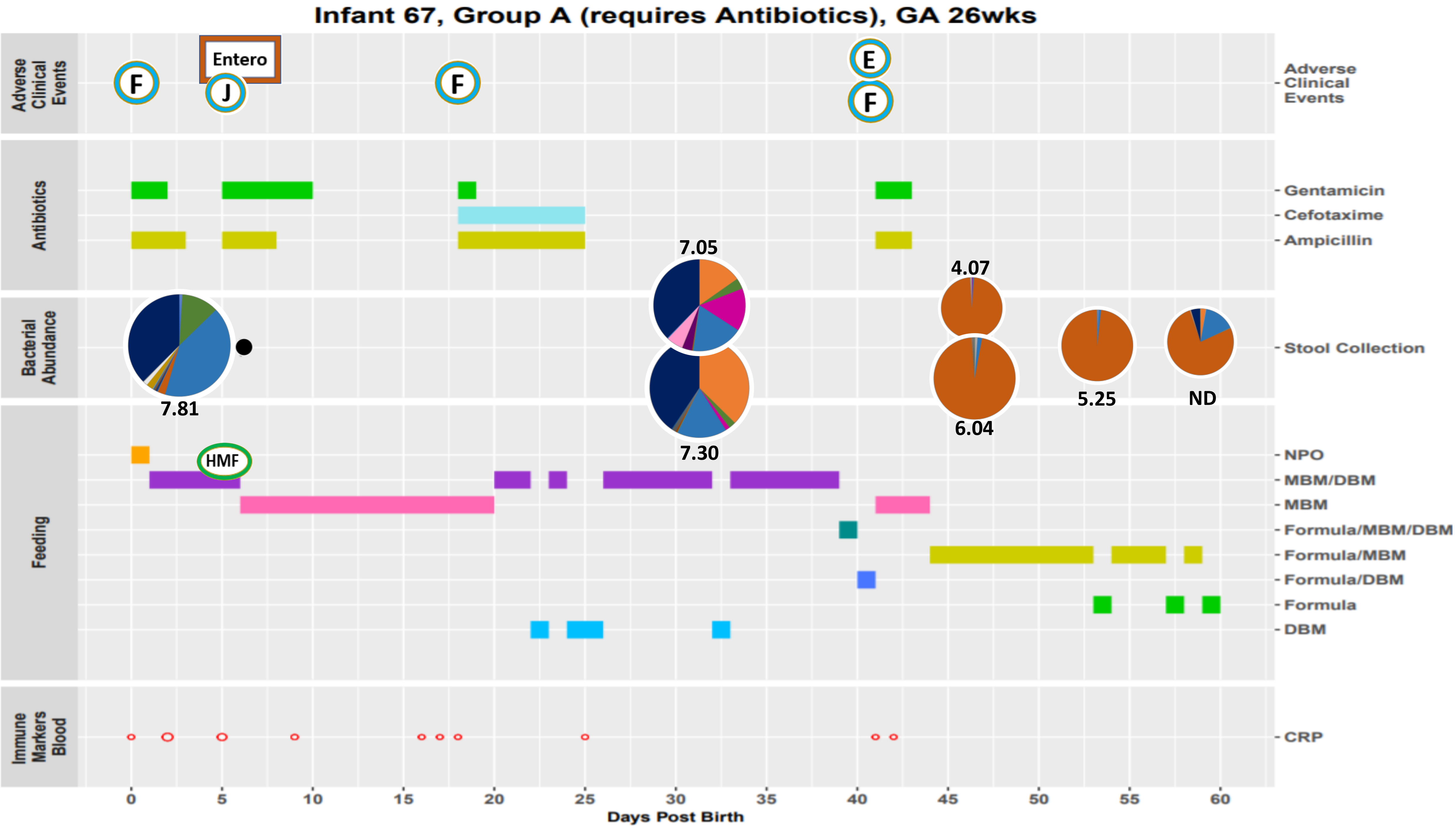
Infant 67, Group A (requires Antibiotics), GA 26wks
Adverse Clinical Events
Entero
F
J
F
E
F
Antibiotics
Gentamicin
Cefotaxime
Ampicillin
Bacterial Abundance
Stool Collection
7.81
7.05
7.30
4.07
6.04
5.25
ND
Feeding
HMF
NPO
MBM/DBM
MBM
Formula/MBM/DBM
Formula/MBM
Formula/DBM
Formula
DBM
Immune Markers Blood
CRP
0
5
10
15
20
25
30
35
40
45
50
55
60
Days Post Birth

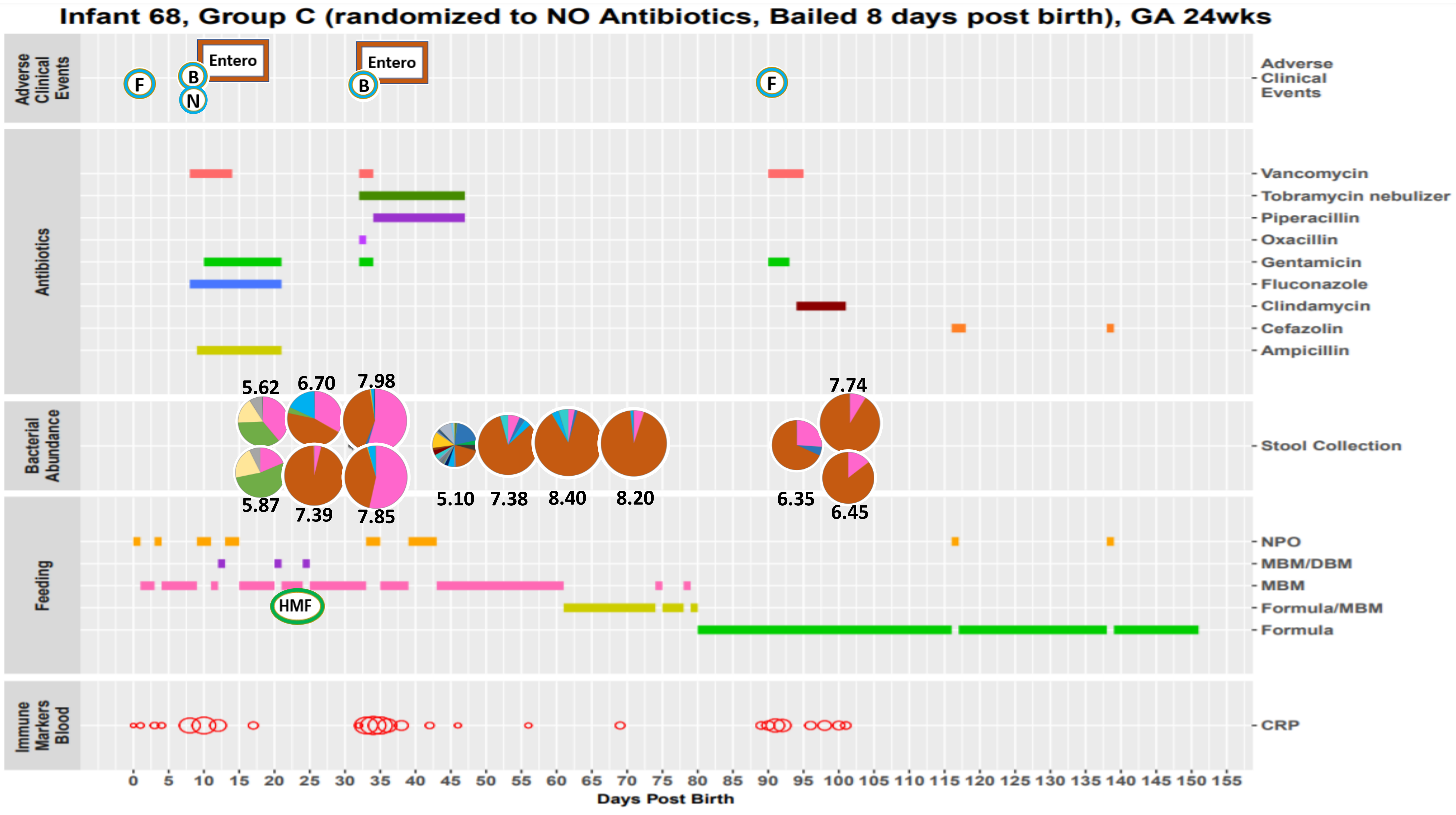
Infant 68, Group C (randomized to NO Antibiotics, Bailed 8 days post birth), GA 24wks
Adverse Clinical Events
Entero
Entero
F
B
N
B
F
Antibiotics
Vancomycin
Tobramycin nebulizer
Piperacillin
Oxacillin
Gentamicin
Fluconazole
Clindamycin
Cefazolin
Ampicillin
Bacterial Abundance
Stool Collection
5.62
6.70
7.98
7.74
5.87
7.39
7.85
5.10
7.38
8.40
8.20
6.35
6.45
Feeding
NPO
MBM/DBM
MBM
Formula/MBM
Formula
HMF
Immune Markers Blood
CRP
0 5 10 15 20 25 30 35 40 45 50 55 60 65 70 75 80 85 90 95 100 105 110 115 120 125 130 135 140 145 150 155
Days Post Birth

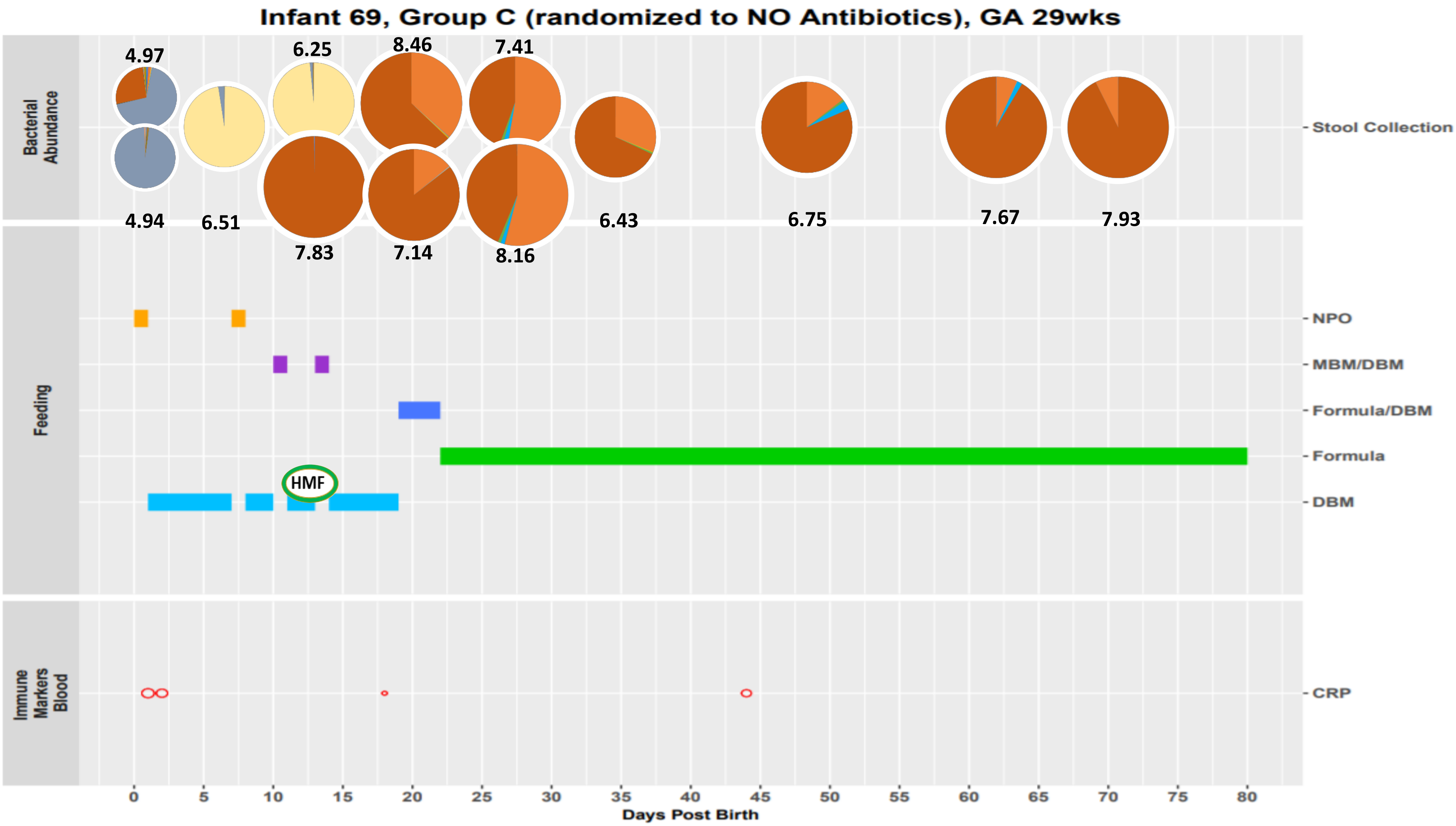

Infant 69, Group C (randomized to NO Antibiotics), GA 29wks
Bacterial Abundance
Feeding
Immune Markers Blood
4.97
6.25
8.46
7.41
4.94
6.51
7.83
7.14
8.16
6.43
6.75
7.67
7.93
HMF
Stool Collection
NPO
MBM/DBM
Formula/DBM
Formula
DBM
CRP
0
5
10
15
20
25
30
35
40
45
50
55
60
65
70
75
80
Days Post Birth

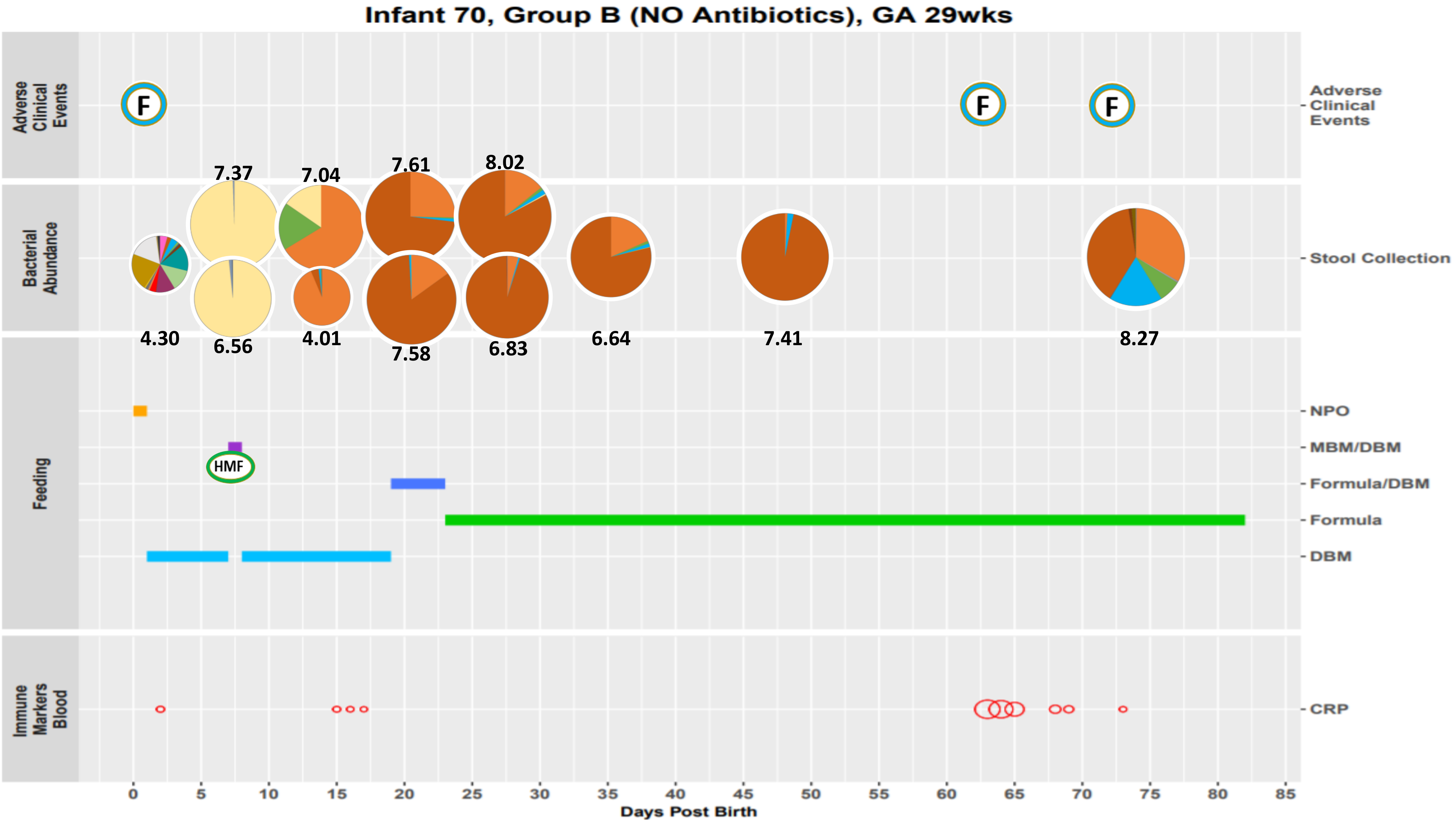

Infant 70, Group B (NO Antibiotics), GA 29wks
Adverse Clinical Events
Bacterial Abundance
Feeding
Immune Markers Blood
F
F
F
7.37
7.04
7.61
8.02
4.30
6.56
4.01
7.58
6.83
6.64
7.41
8.27
HMF
Stool Collection
NPO
MBM/DBM
Formula/DBM
Formula
DBM
CRP
0 5 10 15 20 25 30 35 40 45 50 55 60 65 70 75 80 85
Days Post Birth

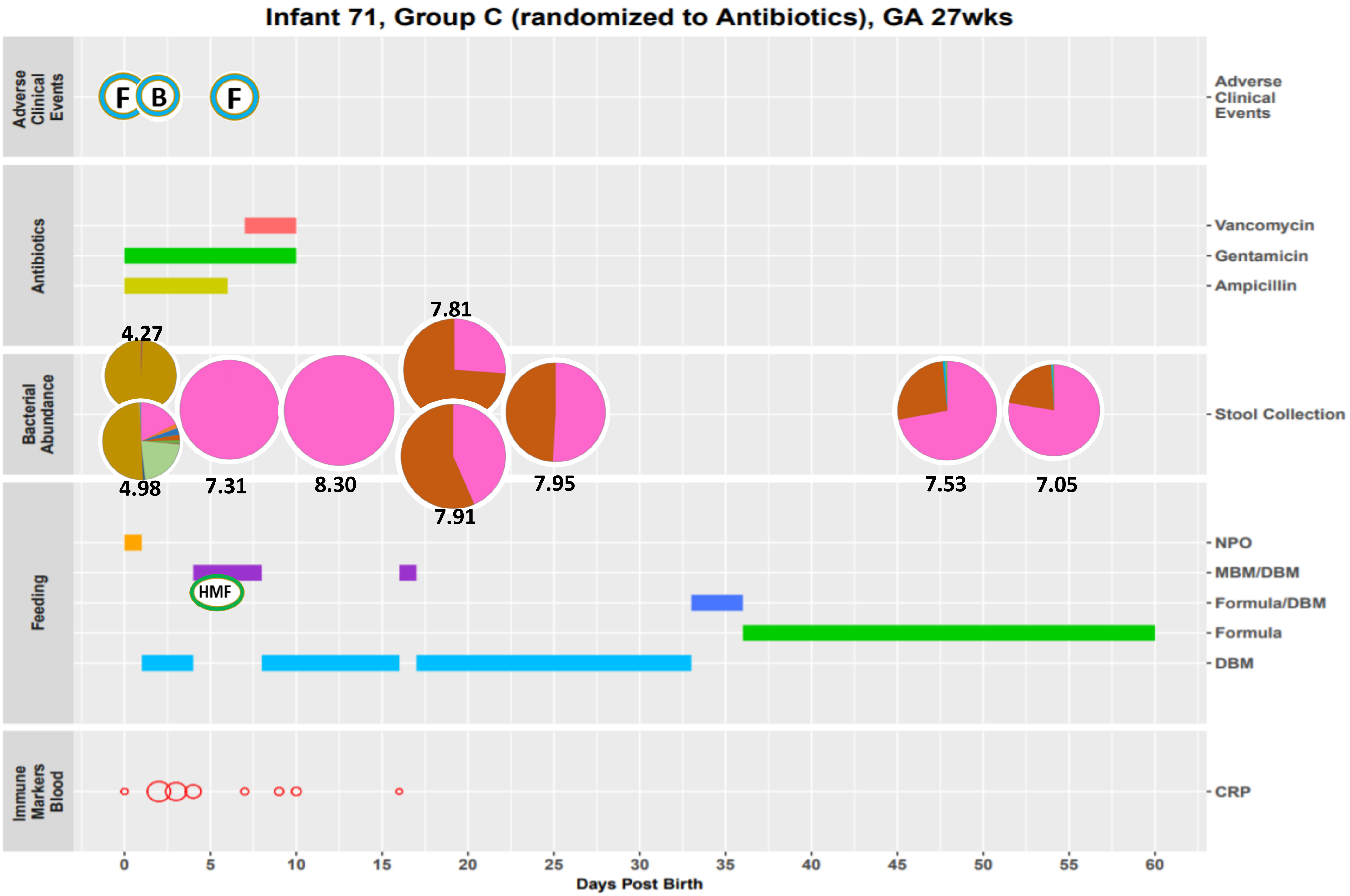
Infant 71, Group C (randomized to Antibiotics), GA 27wks
Adverse Clinical Events
F
B
F
Antibiotics
Vancomycin
Gentamicin
Ampicillin
Bacterial Abundance
Stool Collection
4.27
4.98
7.31
8.30
7.81
7.91
7.95
7.53
7.05
Feeding
NPO
MBM/DBM
HMF
Formula/DBM
Formula
DBM
Immune Markers Blood
CRP
0
5
10
15
20
25
30
35
40
45
50
55
60
Days Post Birth

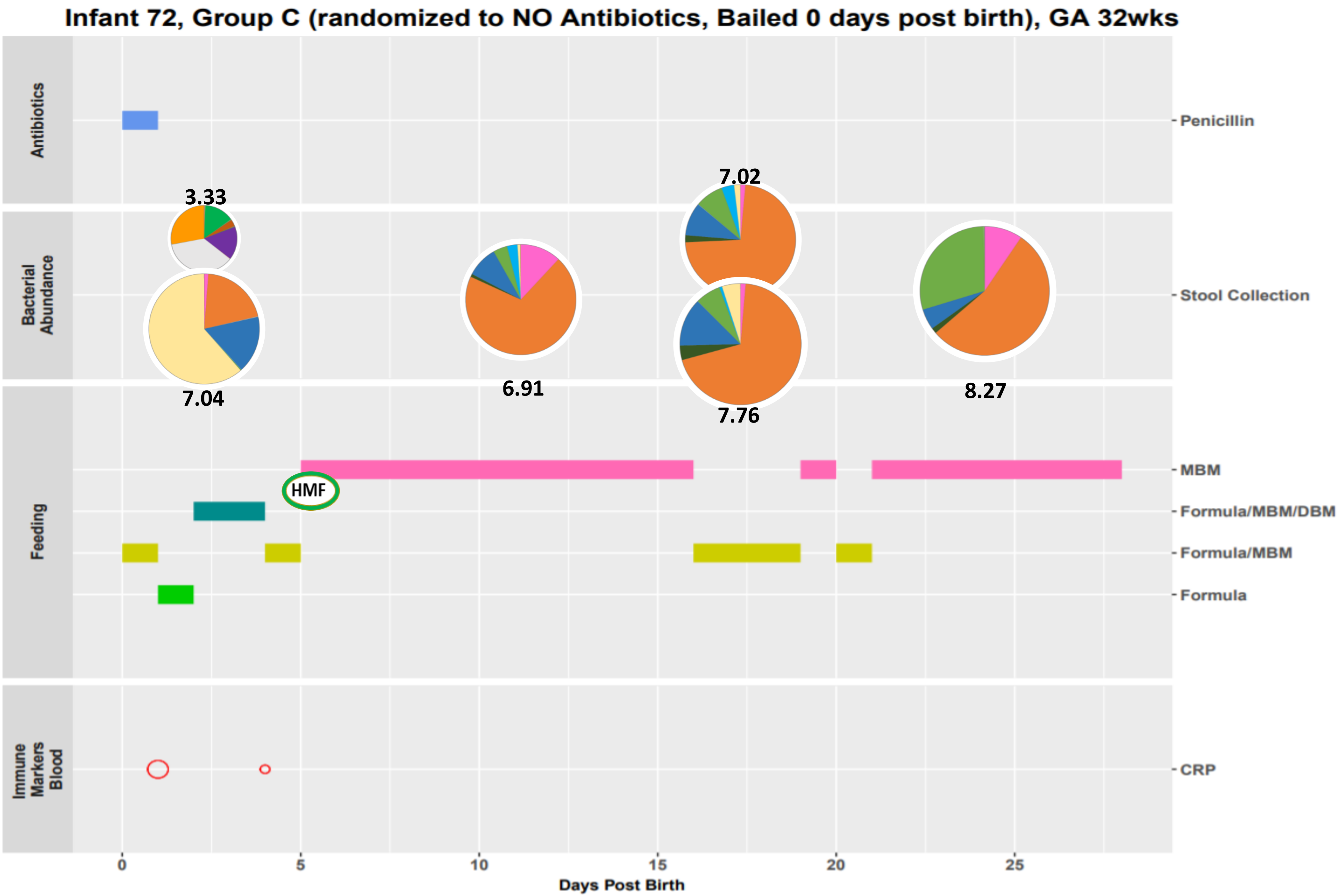
Infant 72, Group C (randomized to NO Antibiotics, Bailed 0 days post birth), GA 32wks
Antibiotics
Penicillin
Bacterial Abundance
Stool Collection
3.33
7.04
6.91
7.02
7.76
8.27
Feeding
MBM
Formula/MBM/DBM
Formula/MBM
Formula
HMF
Immune Markers Blood
CRP
0
5
10
15
20
25
Days Post Birth

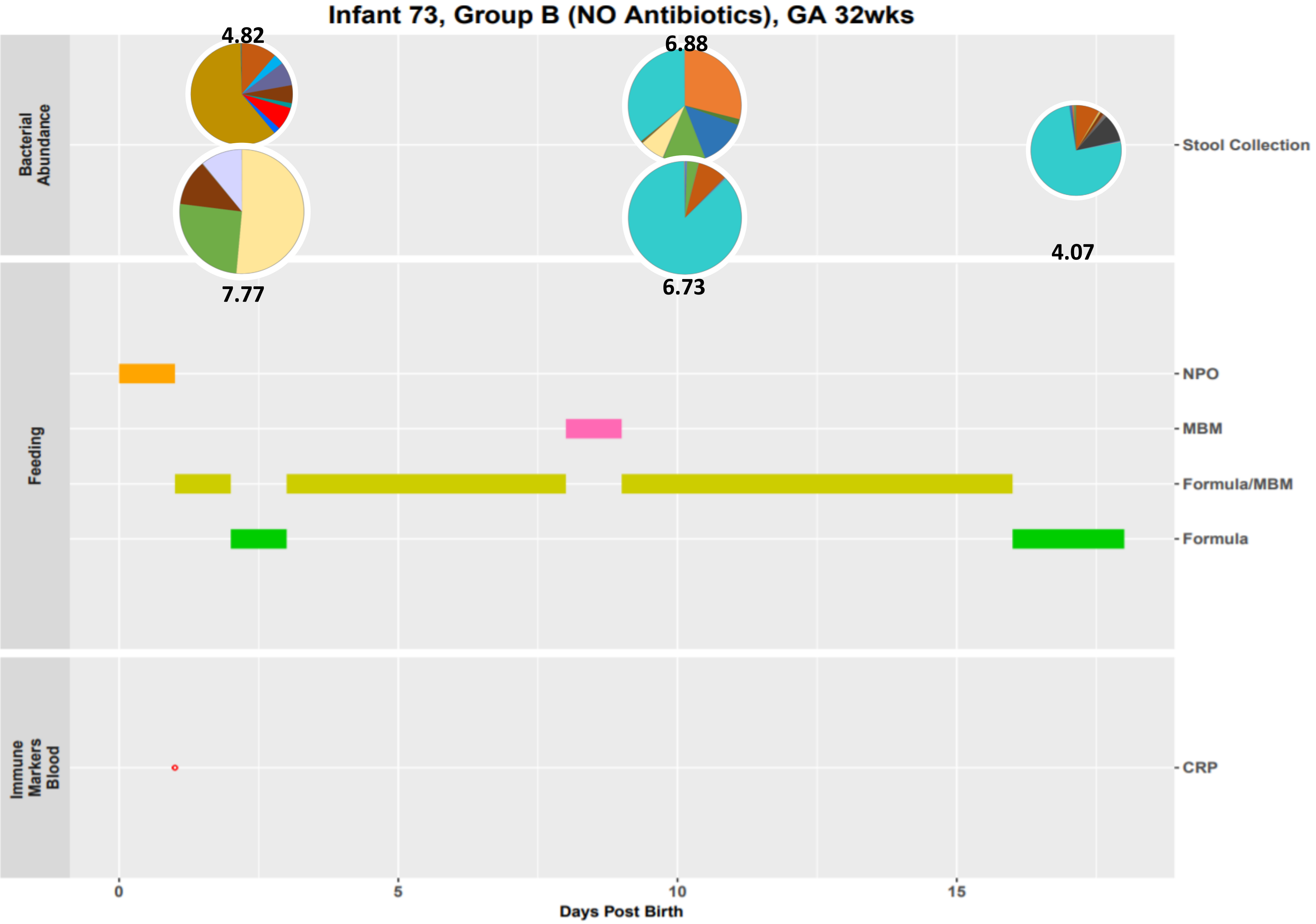
Infant 73, Group B (NO Antibiotics), GA 32wks
Bacterial Abundance
Feeding
Immune Markers Blood
4.82
7.77
6.88
6.73
4.07
Stool Collection
NPO
MBM
Formula/MBM
Formula
CRP
0
5
10
15
Days Post Birth

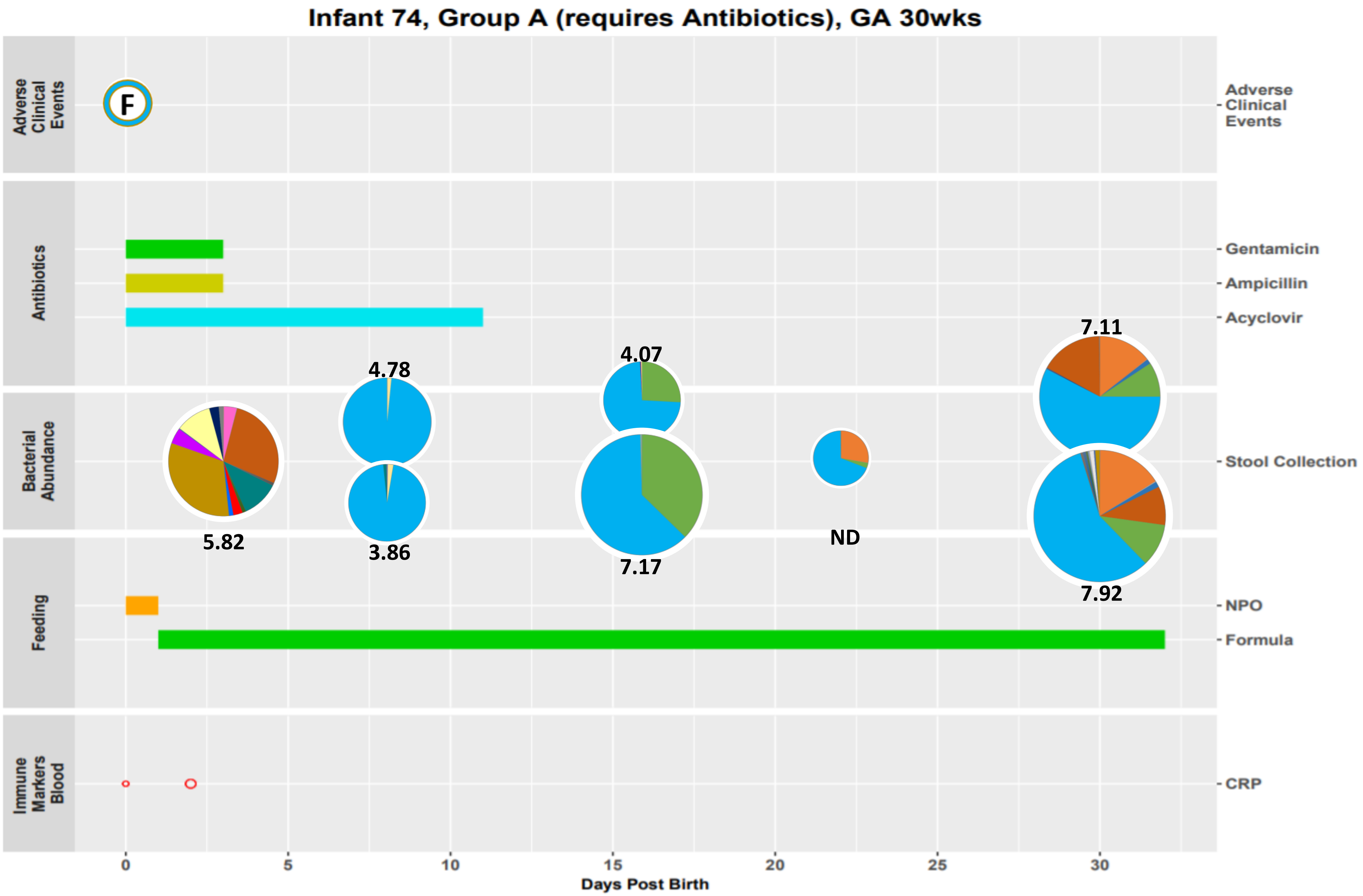
Infant 74, Group A (requires Antibiotics), GA 30wks
Adverse Clinical Events
F
Antibiotics
Gentamicin
Ampicillin
Acyclovir
Bacterial Abundance
Stool Collection
4.78
3.86
4.07
7.17
5.82
ND
7.11
7.92
Feeding
NPO
Formula
Immune Markers Blood
CRP
0
5
10
15
20
25
30
Days Post Birth

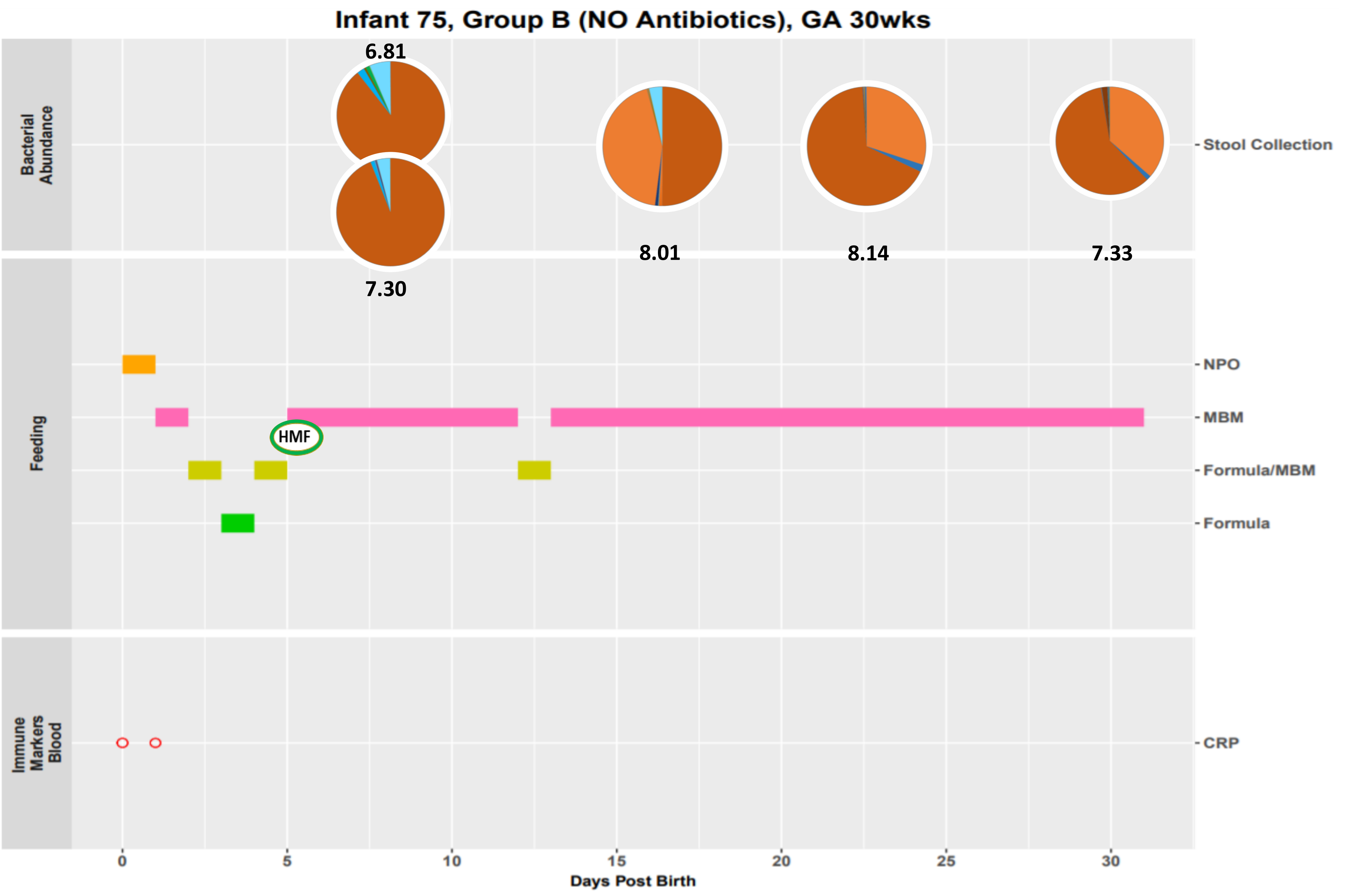

Infant 75, Group B (NO Antibiotics), GA 30wks
Bacterial Abundance
6.81
7.30
8.01
8.14
7.33
Stool Collection
Feeding
NPO
MBM
HMF
Formula/MBM
Formula
Immune Markers Blood
CRP
0
5
10
15
20
25
30
Days Post Birth

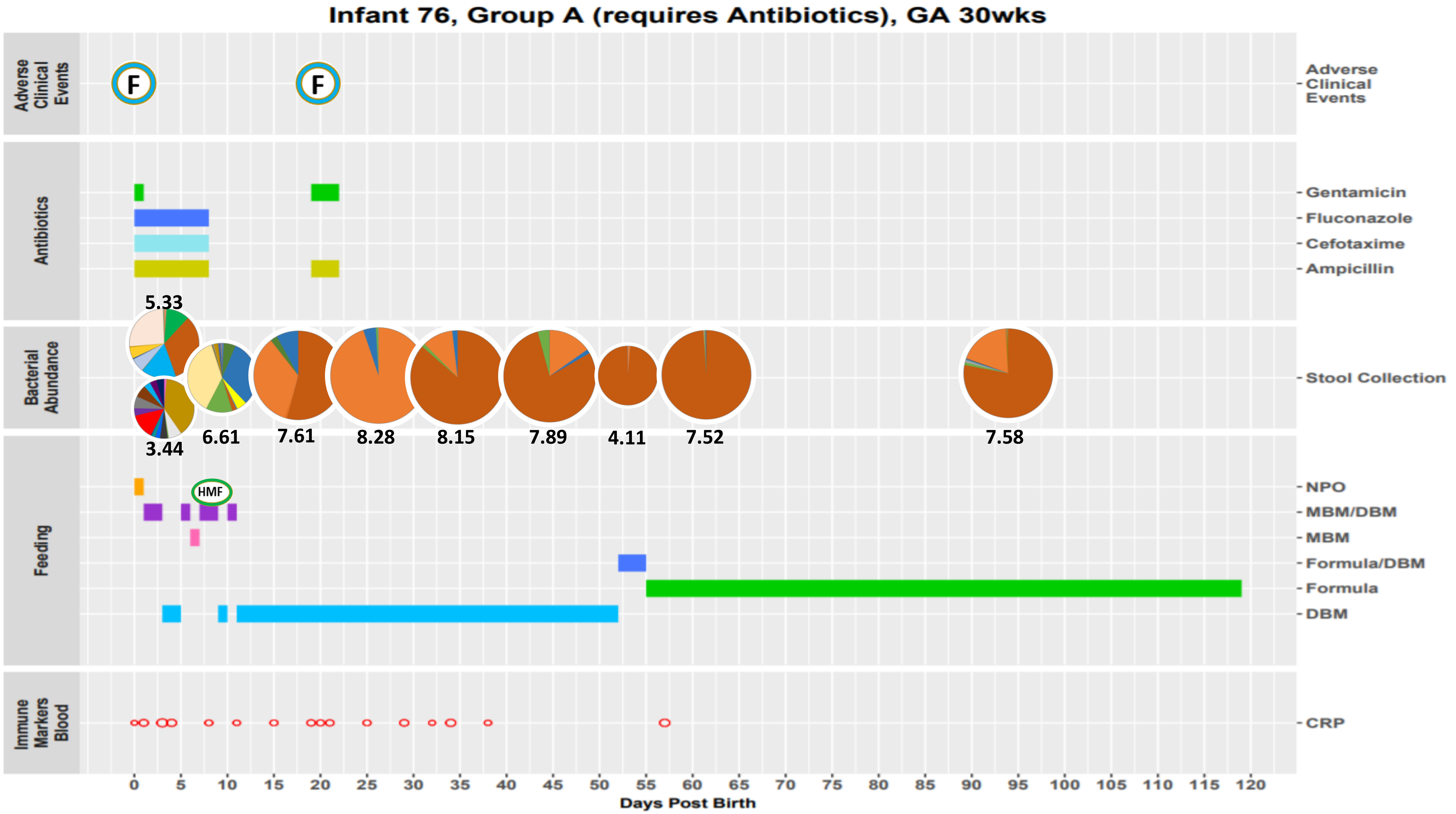
Infant 76, Group A (requires Antibiotics), GA 30wks
Adverse Clinical Events
F
F
Antibiotics
Gentamicin
Fluconazole
Cefotaxime
Ampicillin
Bacterial Abundance
Stool Collection
5.33
3.44
6.61
7.61
8.28
8.15
7.89
4.11
7.52
7.58
Feeding
HMF
NPO
MBM/DBM
MBM
Formula/DBM
Formula
DBM
Immune Markers Blood
CRP
0 5 10 15 20 25 30 35 40 45 50 55 60 65 70 75 80 85 90 95 100 105 110 115 120
Days Post Birth

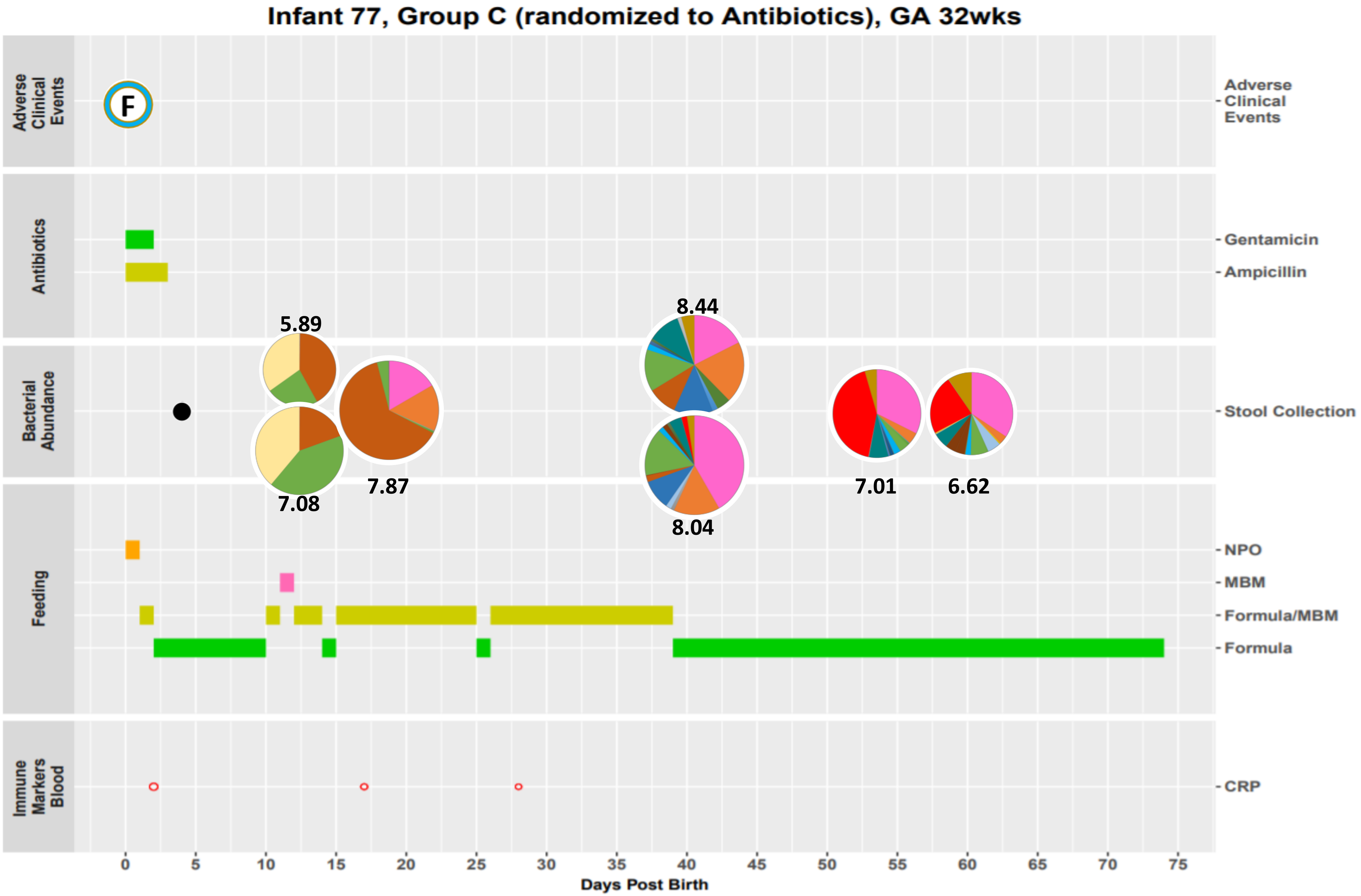
Infant 77, Group C (randomized to Antibiotics), GA 32wks
Adverse Clinical Events
F
Antibiotics
Gentamicin
Ampicillin
Bacterial Abundance
Stool Collection
5.89
7.08
7.87
8.44
8.04
7.01
6.62
Feeding
NPO
MBM
Formula/MBM
Formula
Immune Markers Blood
CRP
0
5
10
15
20
25
30
35
40
45
50
55
60
65
70
75
Days Post Birth

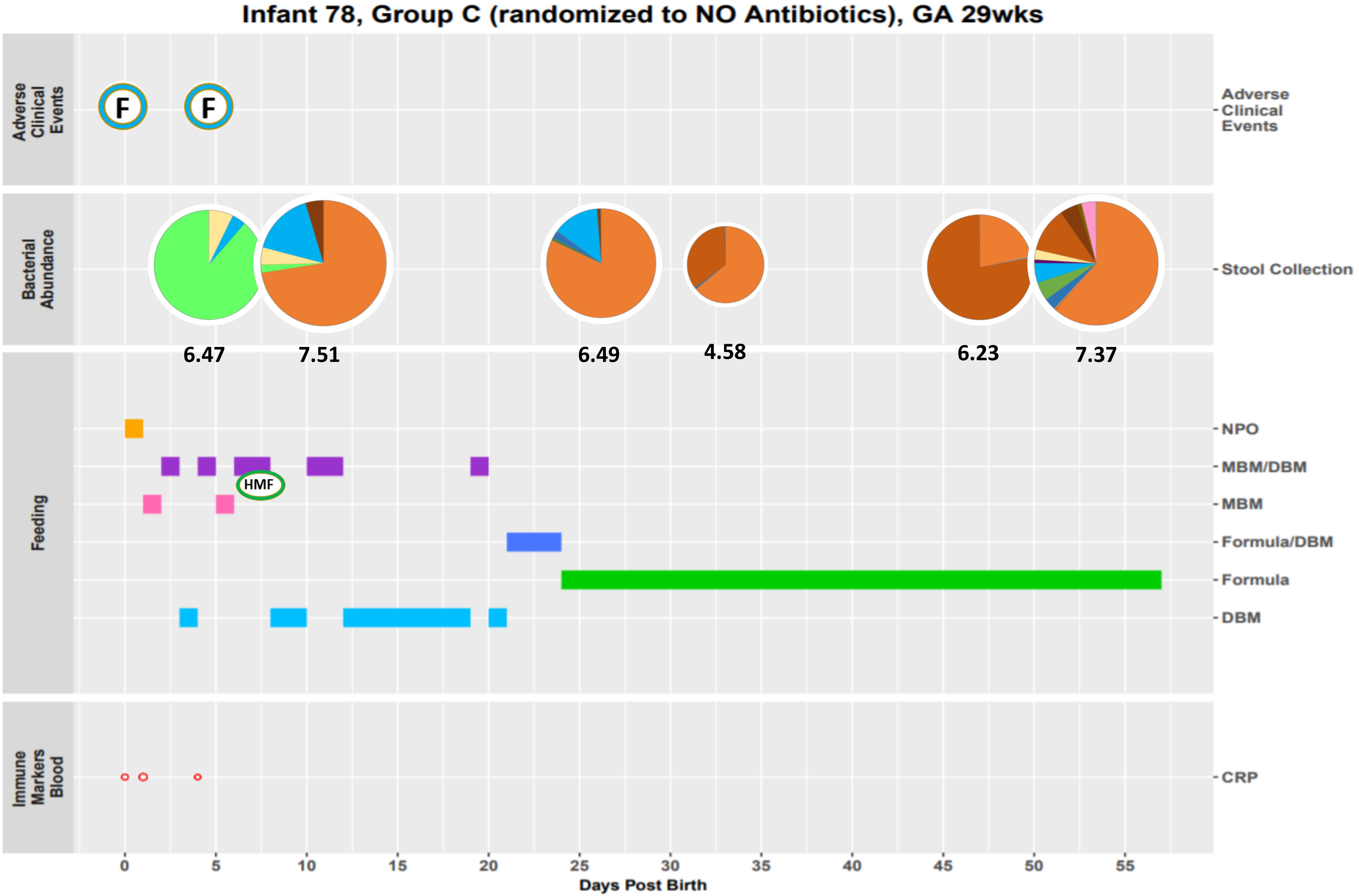

Infant 78, Group C (randomized to NO Antibiotics), GA 29wks
Adverse Clinical Events
Bacterial Abundance
Feeding
Immune Markers Blood
F
F
6.47
7.51
6.49
4.58
6.23
7.37
HMF
Adverse Clinical Events
Stool Collection
NPO
MBM/DBM
MBM
Formula/DBM
Formula
DBM
CRP
0
5
10
15
20
25
30
35
40
45
50
55
Days Post Birth

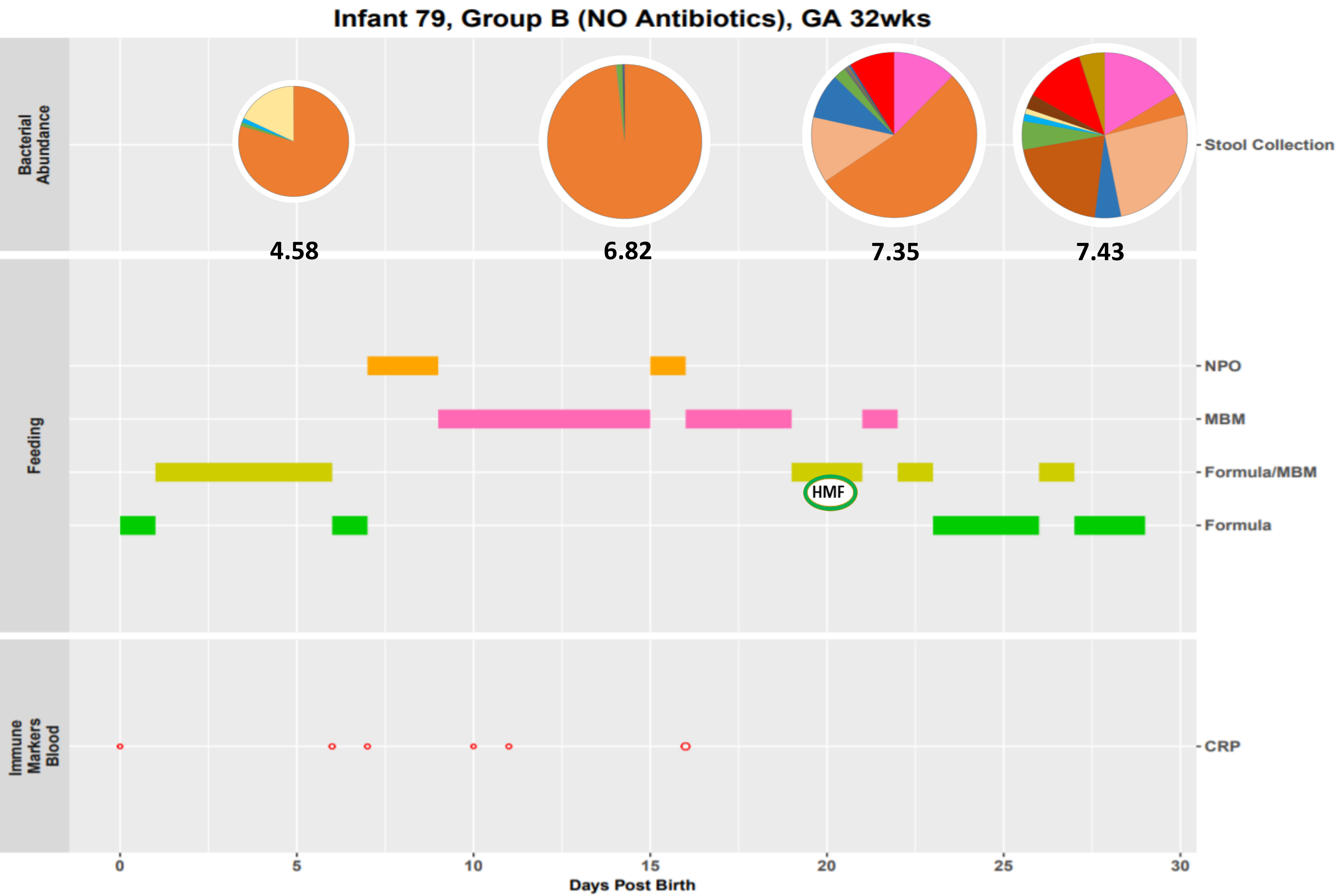

Infant 79, Group B (NO Antibiotics), GA 32wks
Bacterial Abundance
4.58
6.82
7.35
7.43
Stool Collection
Feeding
NPO
MBM
Formula/MBM
HMF
Formula
Immune Markers Blood
CRP
0
5
10
15
20
25
30
Days Post Birth

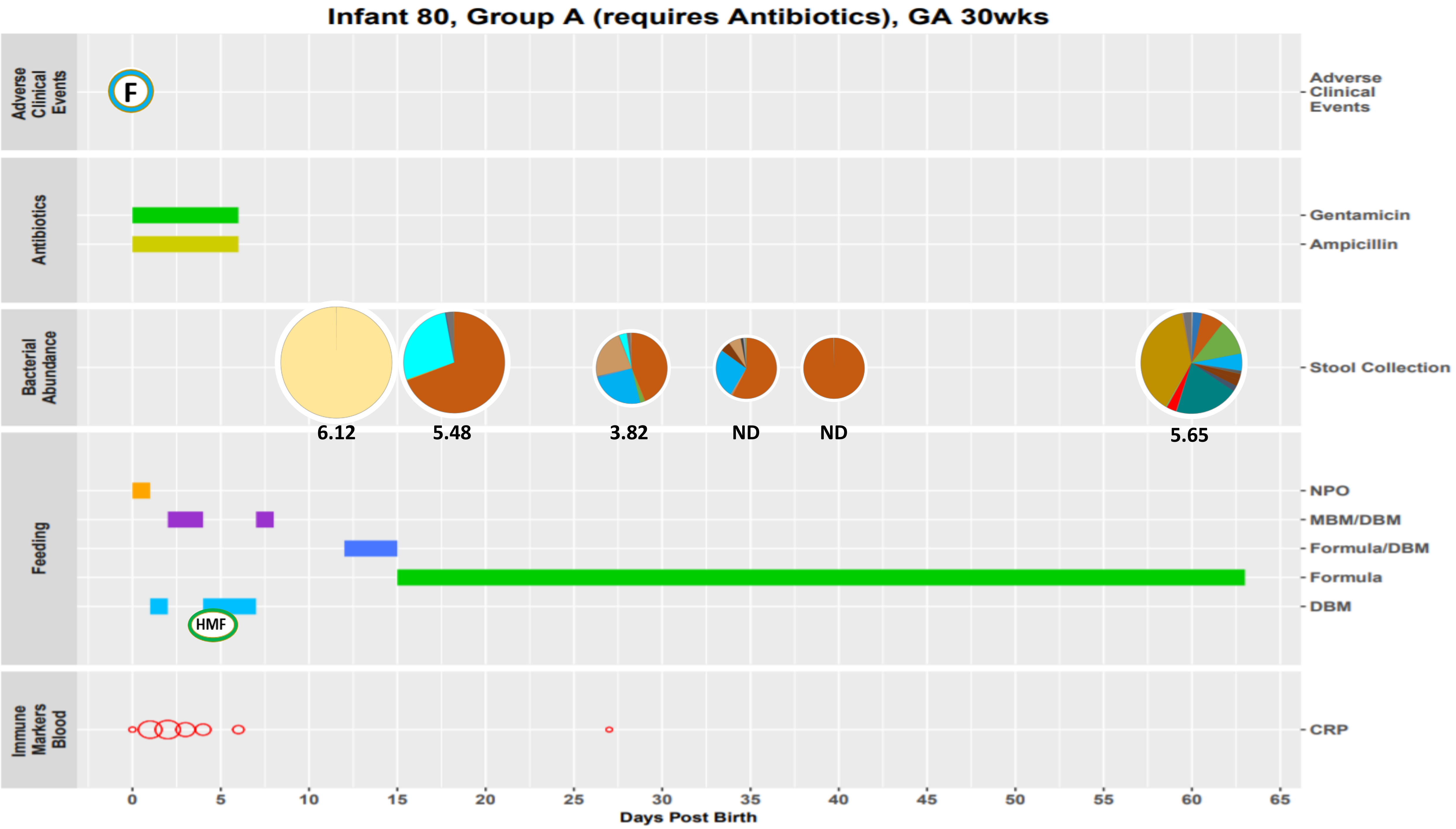

Infant 80, Group A (requires Antibiotics), GA 30wks
Adverse Clinical Events
F
Antibiotics
Gentamicin
Ampicillin
Bacterial Abundance
Stool Collection
6.12
5.48
3.82
ND
ND
5.65
Feeding
NPO
MBM/DBM
Formula/DBM
Formula
DBM
HMF
Immune Markers Blood
CRP
0
5
10
15
20
25
30
35
40
45
50
55
60
65
Days Post Birth

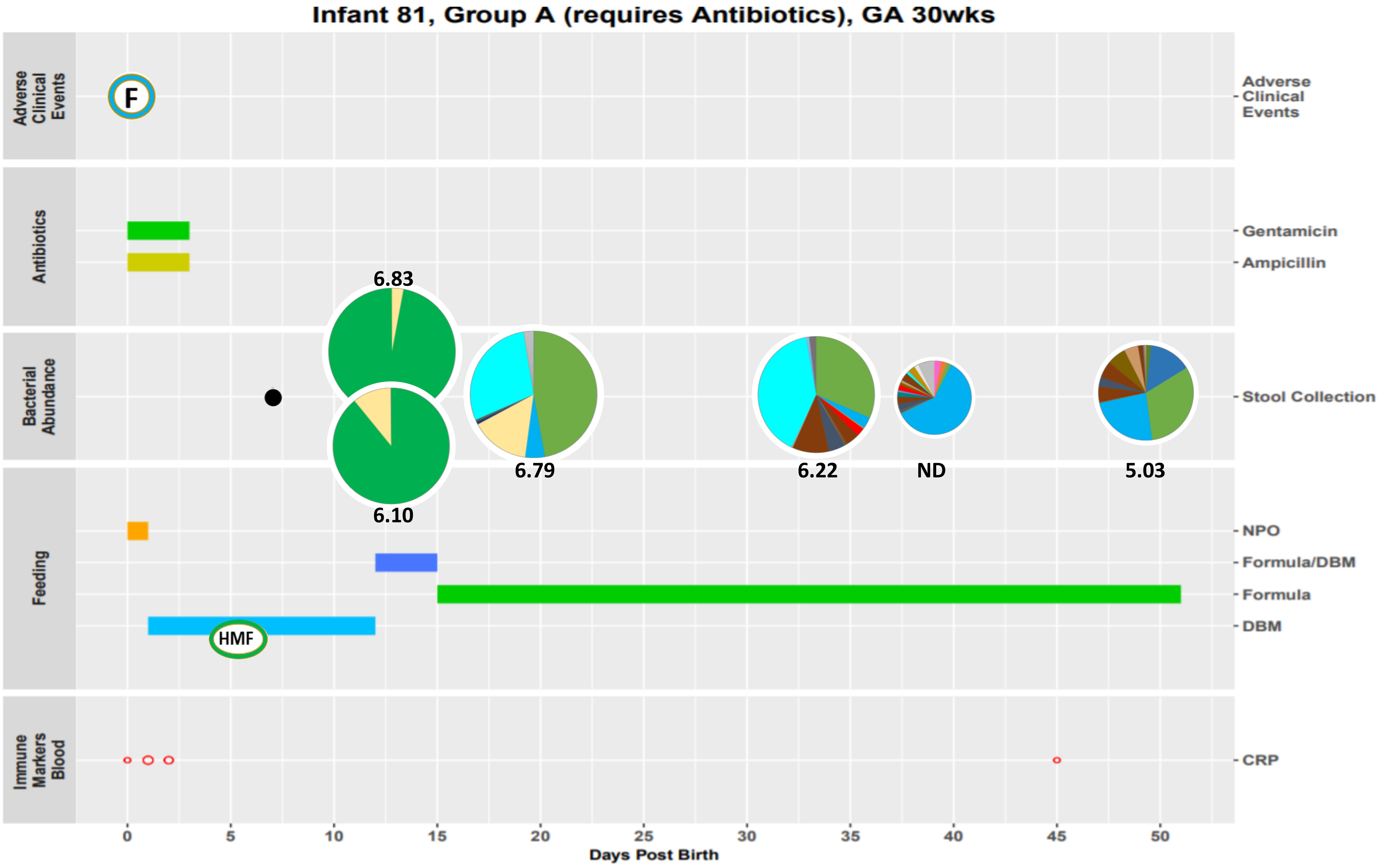
Infant 81, Group A (requires Antibiotics), GA 30wks
Adverse Clinical Events
F
Antibiotics
Gentamicin
Ampicillin
Bacterial Abundance
6.83
6.10
6.79
6.22
ND
5.03
Stool Collection
Feeding
NPO
Formula/DBM
Formula
DBM
HMF
Immune Markers Blood
CRP
0
5
10
15
20
25
30
35
40
45
50
Days Post Birth

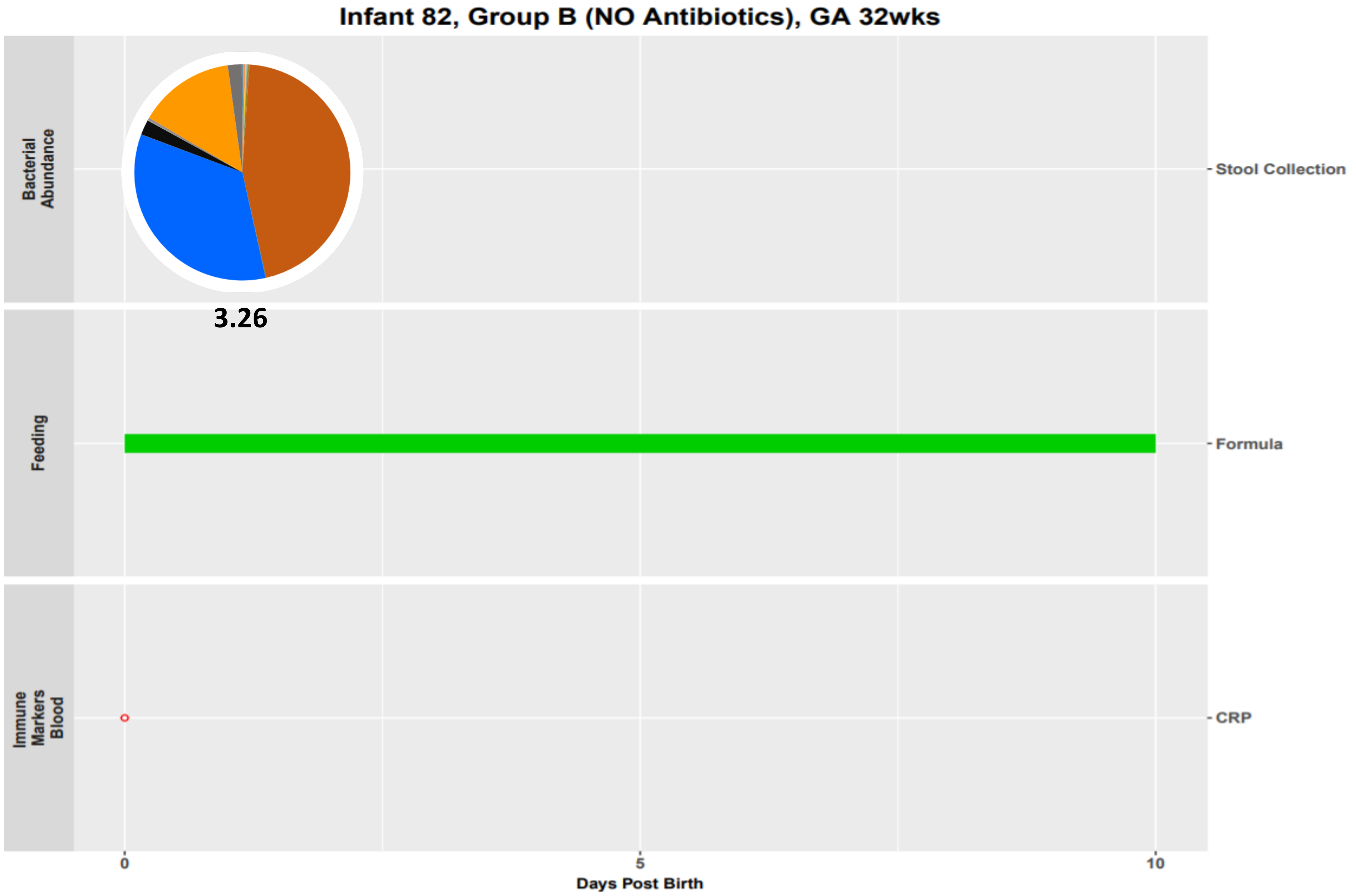
Infant 82, Group B (NO Antibiotics), GA 32wks
Bacterial Abundance
Stool Collection
3.26
Feeding
Formula
Immune Markers Blood
CRP
0
5
10
Days Post Birth

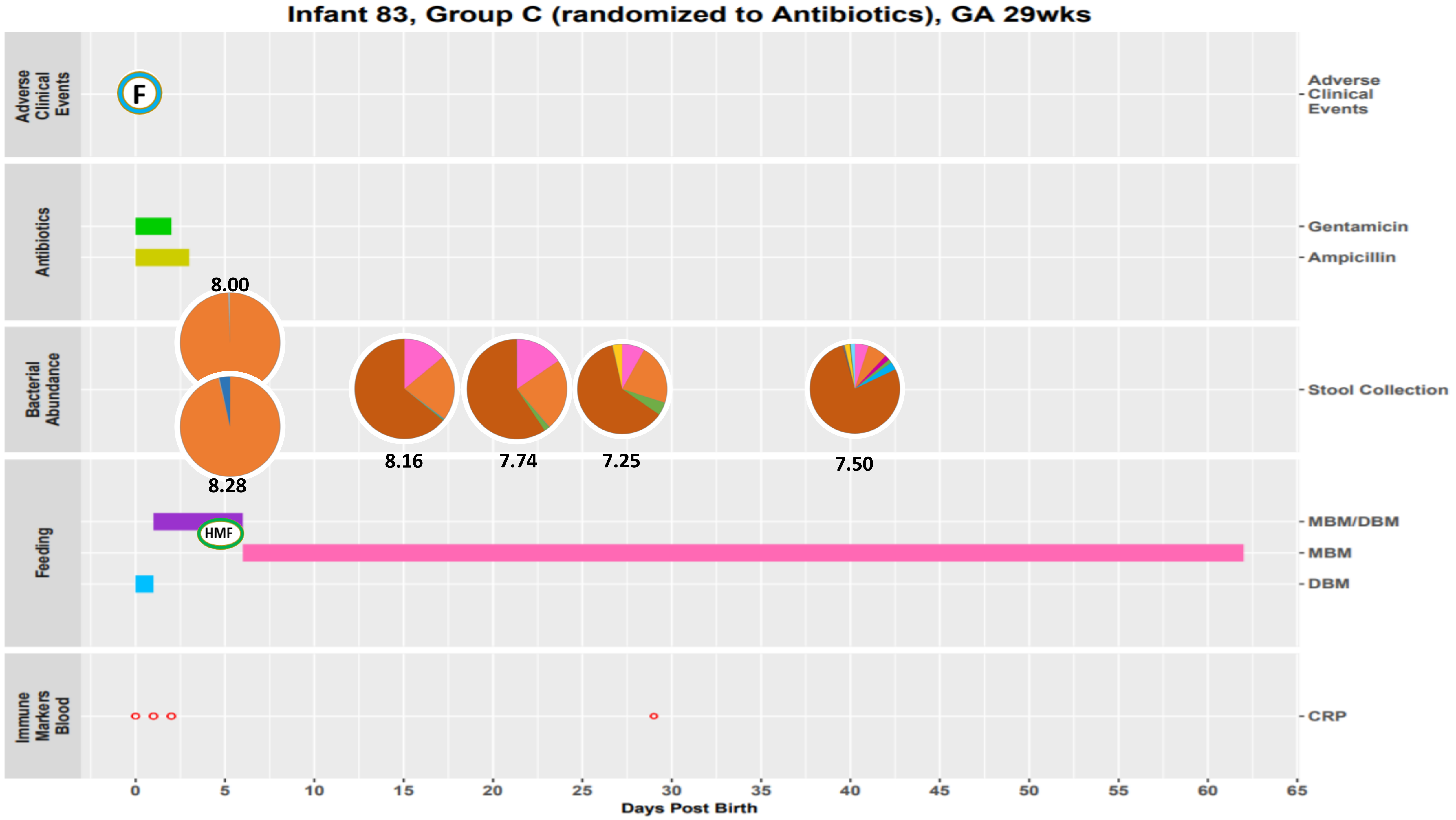
Infant 83, Group C (randomized to Antibiotics), GA 29wks
Adverse Clinical Events
Antibiotics
Bacterial Abundance
Feeding
Immune Markers Blood
F
Gentamicin
Ampicillin
Stool Collection
8.00
8.28
8.16
7.74
7.25
7.50
MBM/DBM
HMF
MBM
DBM
CRP
0
5
10
15
20
25
30
35
40
45
50
55
60
65
Days Post Birth

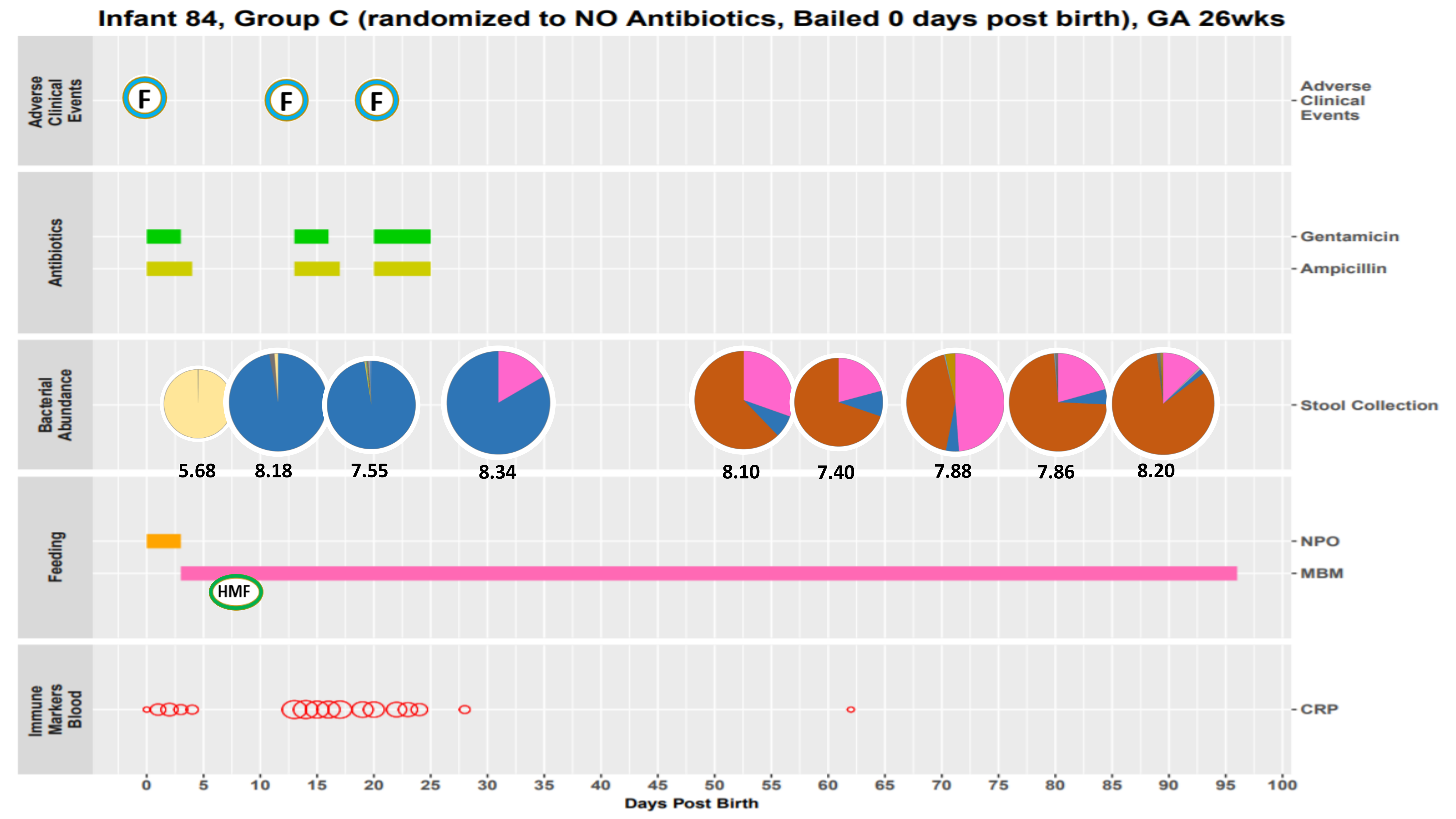
Infant 84, Group C (randomized to NO Antibiotics, Bailed 0 days post birth), GA 26wks
Adverse Clinical Events
F
F
F
Antibiotics
Gentamicin
Ampicillin
Bacterial Abundance
Stool Collection
5.68
8.18
7.55
8.34
8.10
7.40
7.88
7.86
8.20
Feeding
NPO
MBM
HMF
Immune Markers Blood
CRP
0
5
10
15
20
25
30
35
40
45
50
55
60
65
70
75
80
85
90
95
100
Days Post Birth

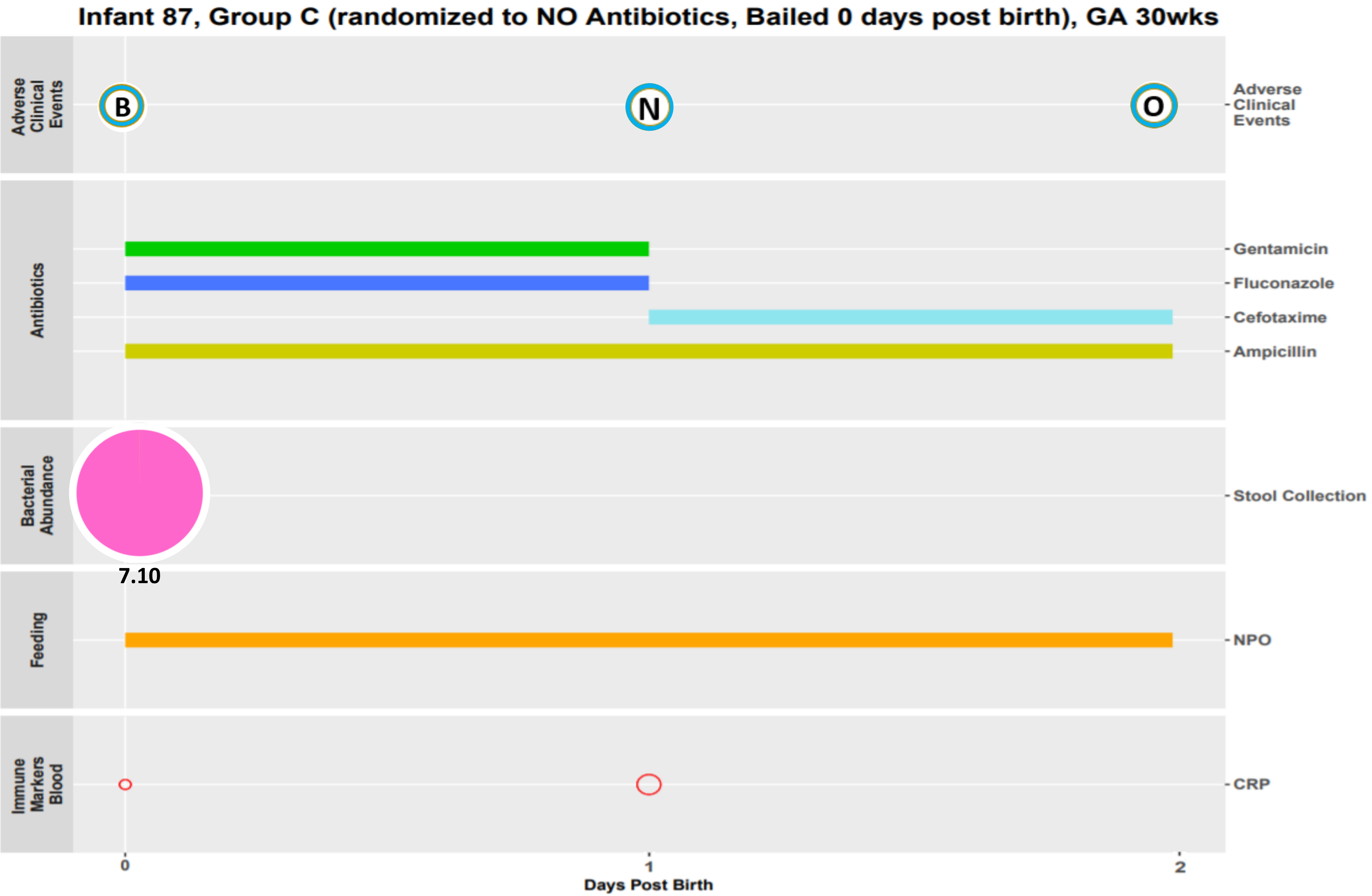
Infant 87, Group C (randomized to NO Antibiotics, Bailed 0 days post birth), GA 30wks
Adverse Clinical Events
B
N
O
Antibiotics
Gentamicin
Fluconazole
Cefotaxime
Ampicillin
Bacterial Abundance
Stool Collection
7.10
Feeding
NPO
Immune Markers Blood
CRP
0
1
2
Days Post Birth

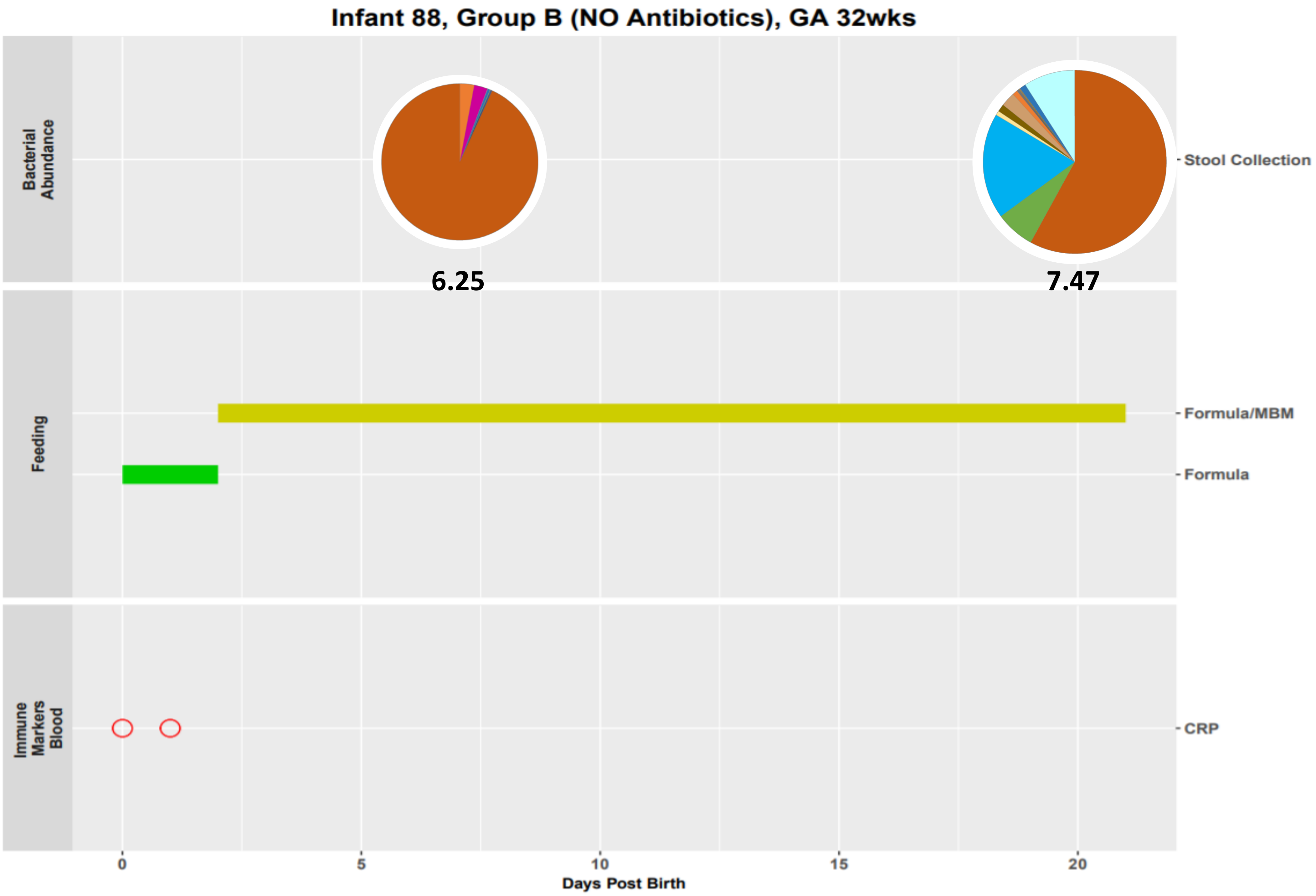

Infant 88, Group B (NO Antibiotics), GA 32wks
Bacterial Abundance
6.25
7.47
Stool Collection
Feeding
Formula/MBM
Formula
Immune Markers Blood
CRP
0
5
10
15
20
Days Post Birth

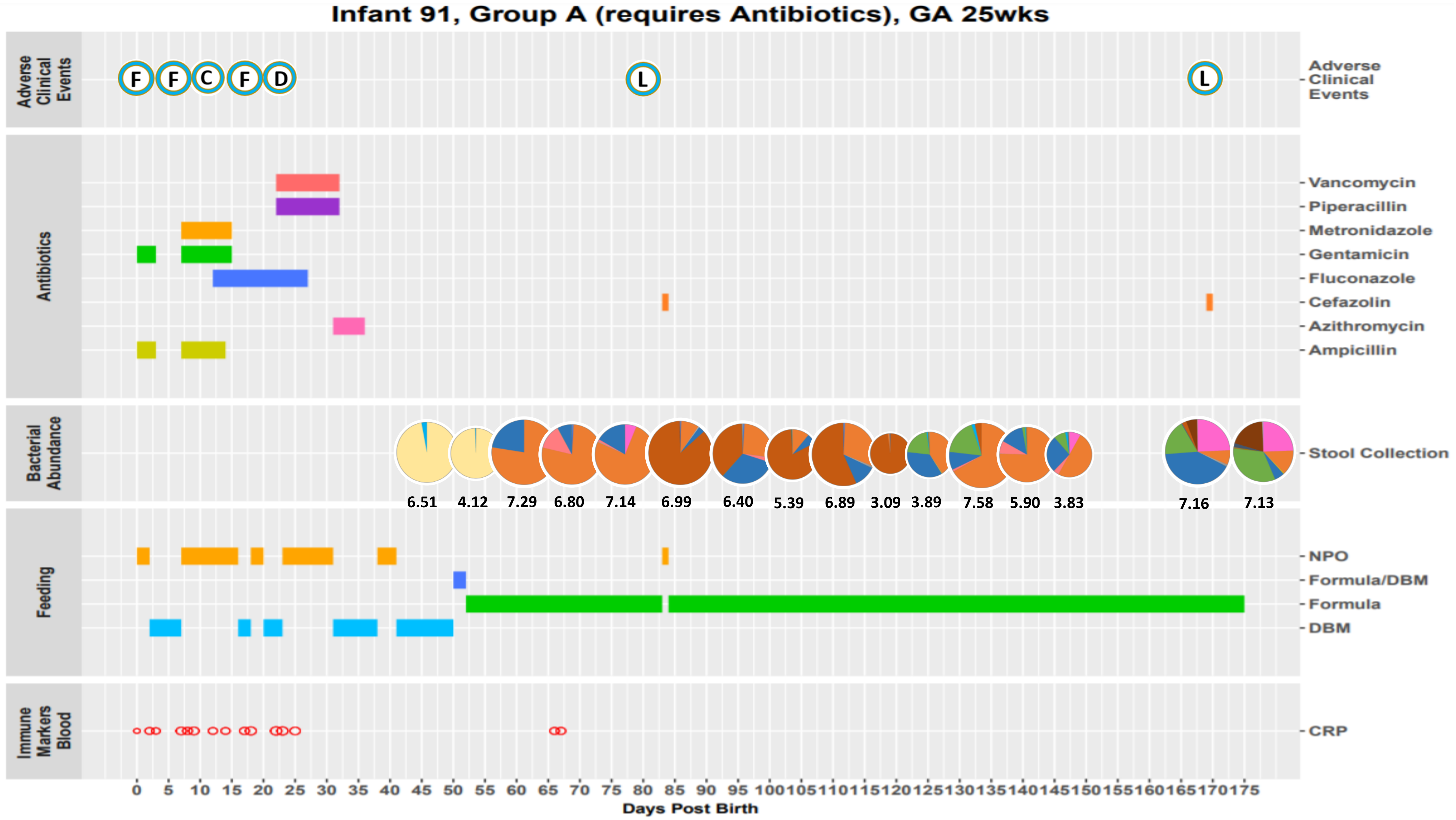

Infant 91, Group A (requires Antibiotics), GA 25wks
Adverse Clinical Events
F
F
C
F
D
L
L
Antibiotics
Vancomycin
Piperacillin
Metronidazole
Gentamicin
Fluconazole
Cefazolin
Azithromycin
Ampicillin
Bacterial Abundance
Stool Collection
6.51
4.12
7.29
6.80
7.14
6.99
6.40
5.39
6.89
3.09
3.89
7.58
5.90
3.83
7.16
7.13
Feeding
NPO
Formula/DBM
Formula
DBM
Immune Markers Blood
CRP
0 5 10 15 20 25 30 35 40 45 50 55 60 65 70 75 80 85 90 95 100 105 110 115 120 125 130 135 140 145 150 155 160 165 170 175
Days Post Birth

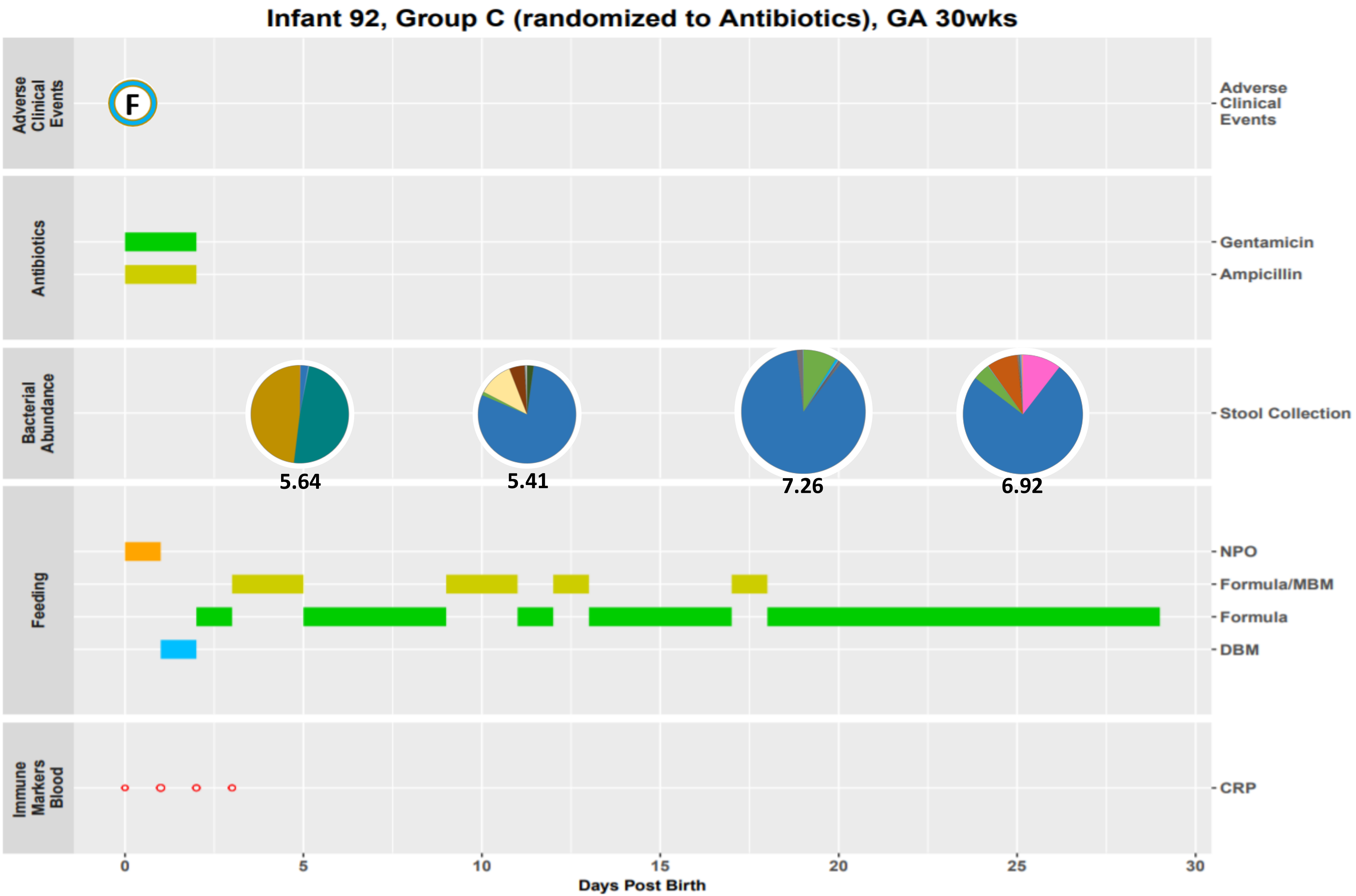

Infant 92, Group C (randomized to Antibiotics), GA 30wks
Adverse Clinical Events
F
Antibiotics
Gentamicin
Ampicillin
Bacterial Abundance
Stool Collection
5.64
5.41
7.26
6.92
Feeding
NPO
Formula/MBM
Formula
DBM
Immune Markers Blood
CRP
0
5
10
15
20
25
30
Days Post Birth

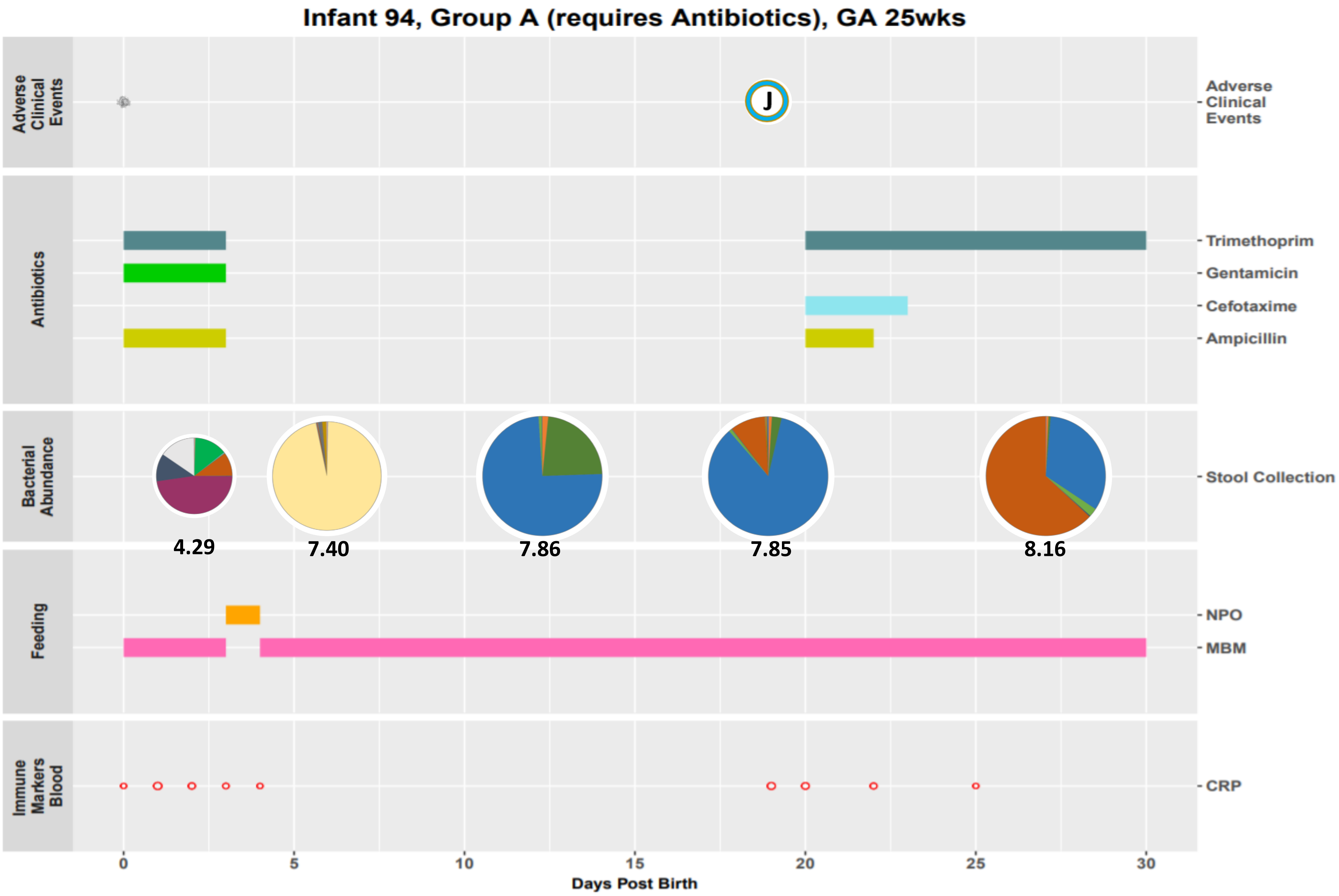

Infant 94, Group A (requires Antibiotics), GA 25wks
Adverse Clinical Events
J
Antibiotics
Trimethoprim
Gentamicin
Cefotaxime
Ampicillin
Bacterial Abundance
Stool Collection
4.29
7.40
7.86
7.85
8.16
Feeding
NPO
MBM
Immune Markers Blood
CRP
0
5
10
15
20
25
30
Days Post Birth

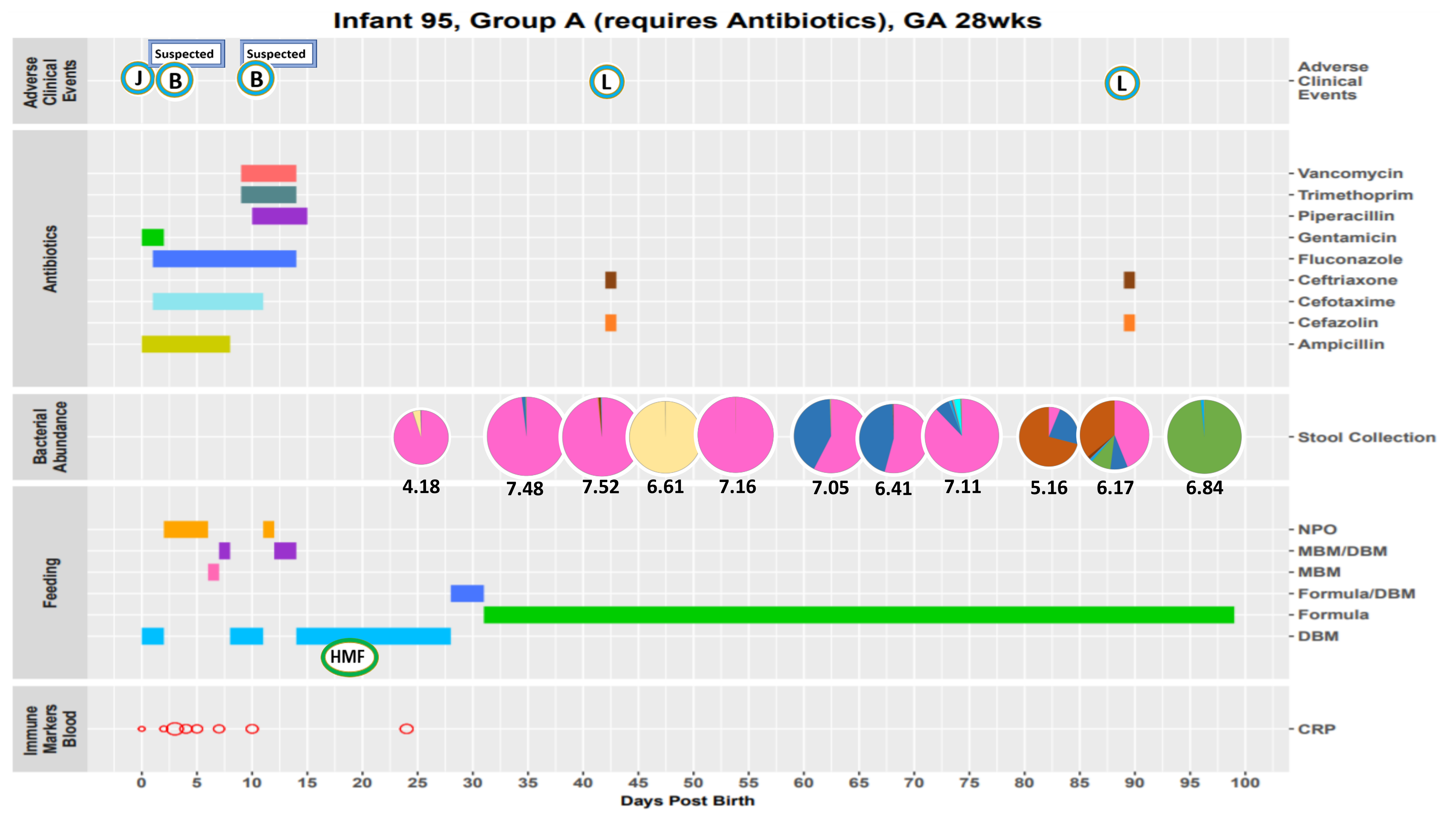
Infant 95, Group A (requires Antibiotics), GA 28wks
Adverse Clinical Events
Suspected
Suspected
J
B
B
L
L
Antibiotics
Vancomycin
Trimethoprim
Piperacillin
Gentamicin
Fluconazole
Ceftriaxone
Cefotaxime
Cefazolin
Ampicillin
Bacterial Abundance
Stool Collection
4.18
7.48
7.52
6.61
7.16
7.05
6.41
7.11
5.16
6.17
6.84
Feeding
NPO
MBM/DBM
MBM
Formula/DBM
Formula
DBM
HMF
Immune Markers Blood
CRP
0
5
10
15
20
25
30
35
40
45
50
55
60
65
70
75
80
85
90
95
100
Days Post Birth

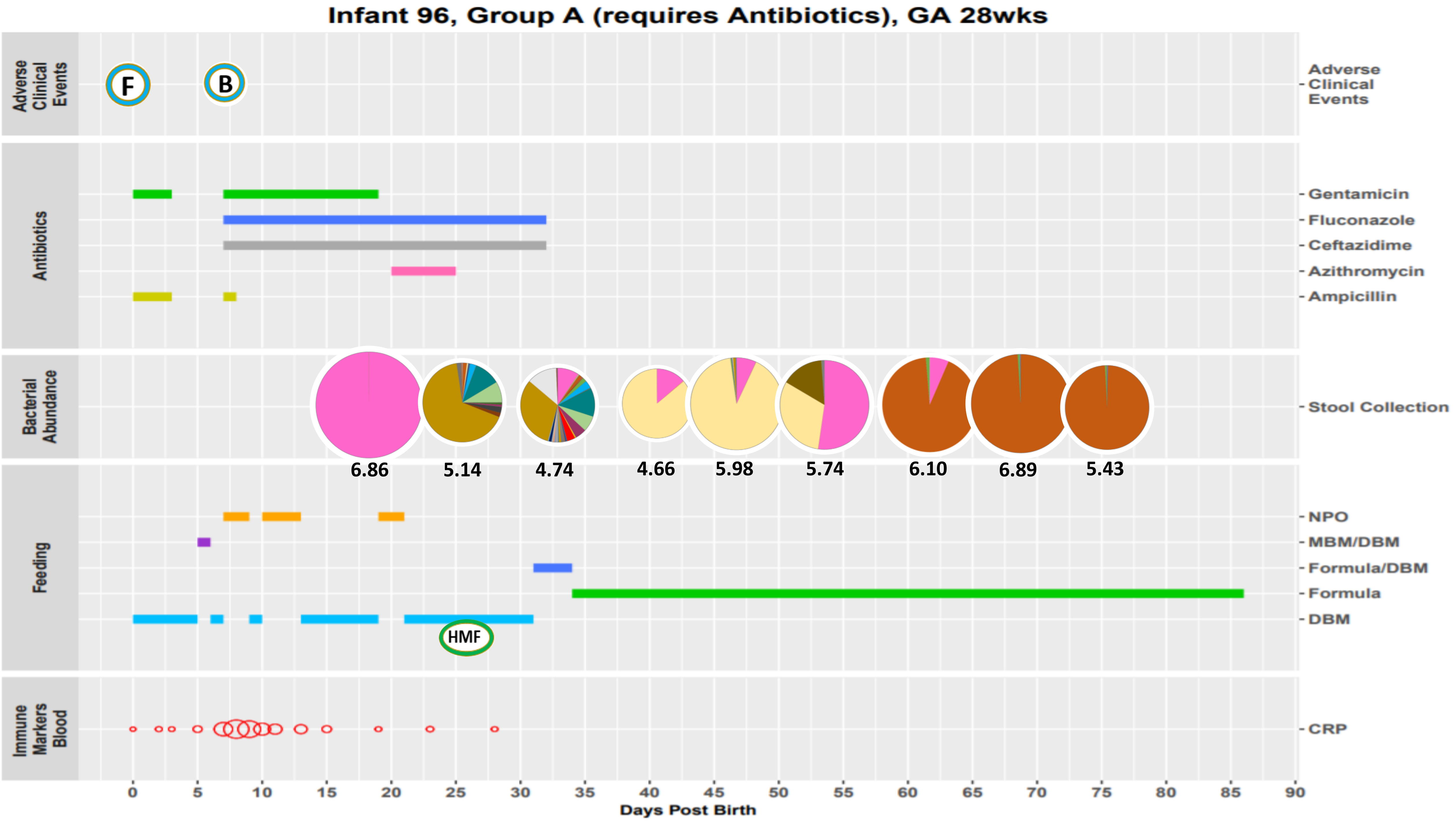

Infant 96, Group A (requires Antibiotics), GA 28wks
Adverse Clinical Events
F
B
Antibiotics
Gentamicin
Fluconazole
Ceftazidime
Azithromycin
Ampicillin
Bacterial Abundance
Stool Collection
6.86
5.14
4.74
4.66
5.98
5.74
6.10
6.89
5.43
Feeding
NPO
MBM/DBM
Formula/DBM
Formula
DBM
HMF
Immune Markers Blood
CRP
0 5 10 15 20 25 30 35 40 45 50 55 60 65 70 75 80 85 90
Days Post Birth

### Infant 97, Group C (randomized to Antibiotics), GA 25wks

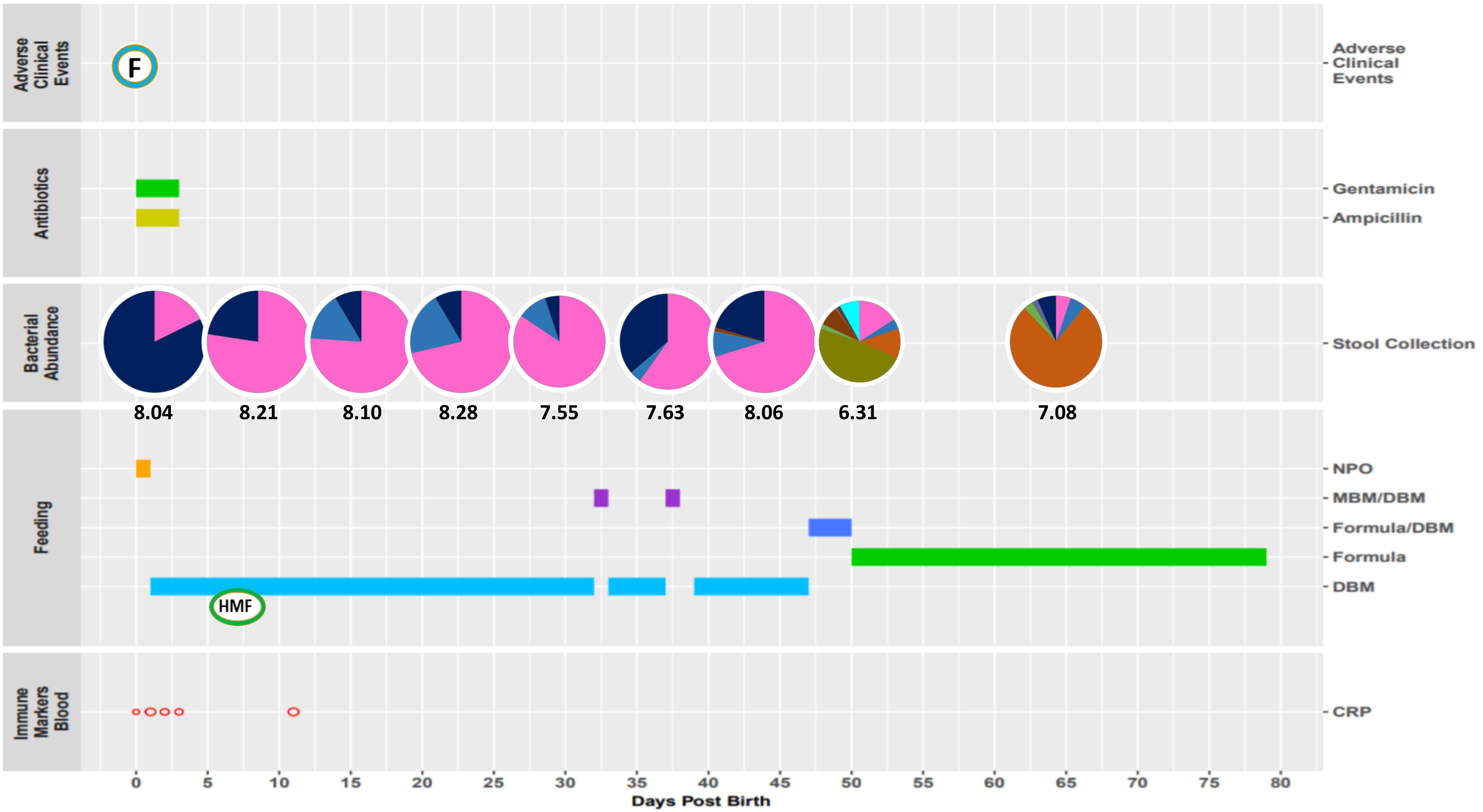

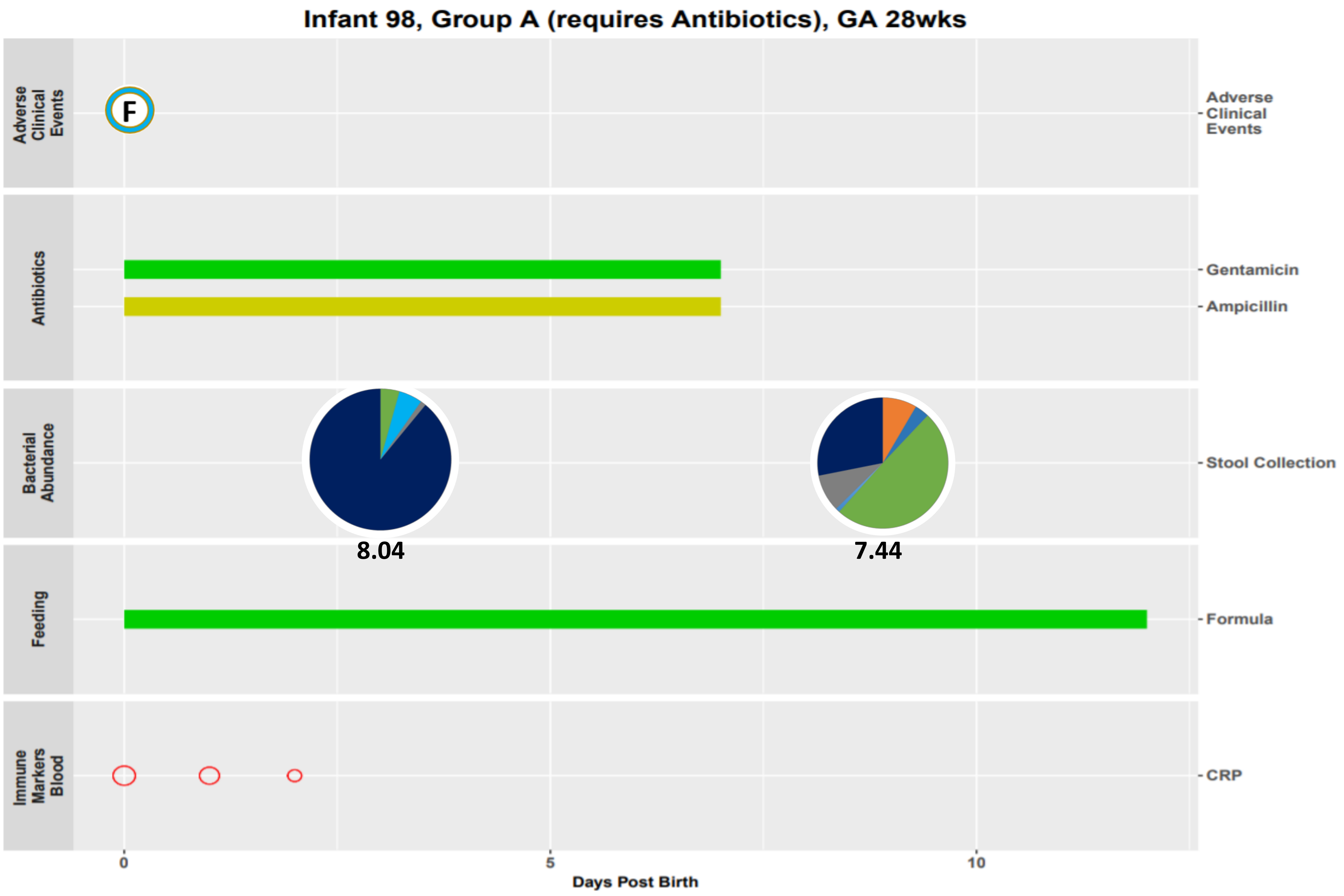
Infant 98, Group A (requires Antibiotics), GA 28wks
Adverse Clinical Events
F
Antibiotics
Gentamicin
Ampicillin
Bacterial Abundance
Stool Collection
8.04
7.44
Feeding
Formula
Immune Markers Blood
CRP
0
5
10
Days Post Birth

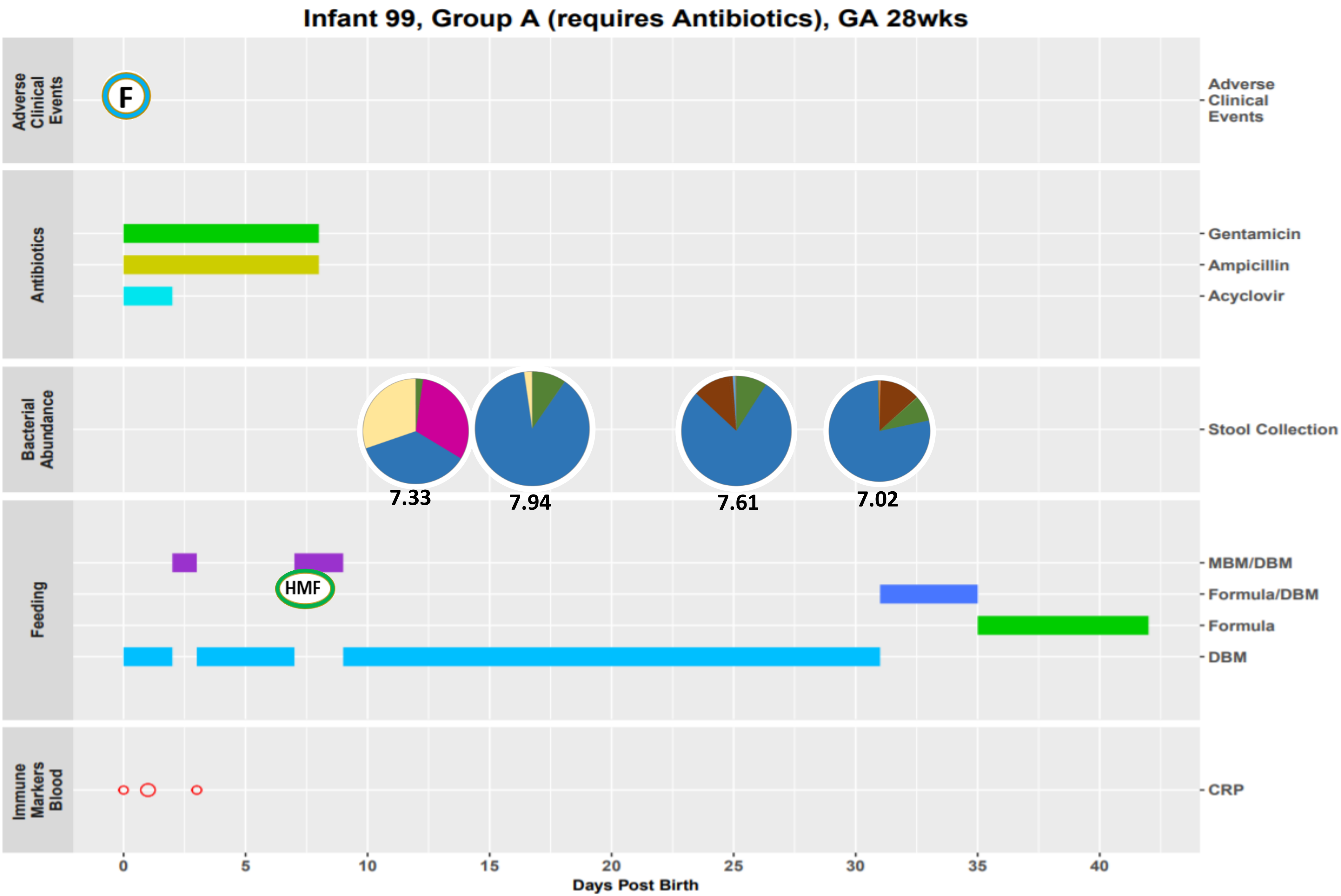

Infant 99, Group A (requires Antibiotics), GA 28wks
Adverse Clinical Events
F
Antibiotics
Gentamicin
Ampicillin
Acyclovir
Bacterial Abundance
Stool Collection
7.33
7.94
7.61
7.02
Feeding
MBM/DBM
HMF
Formula/DBM
Formula
DBM
Immune Markers Blood
CRP
0
5
10
15
20
25
30
35
40
Days Post Birth

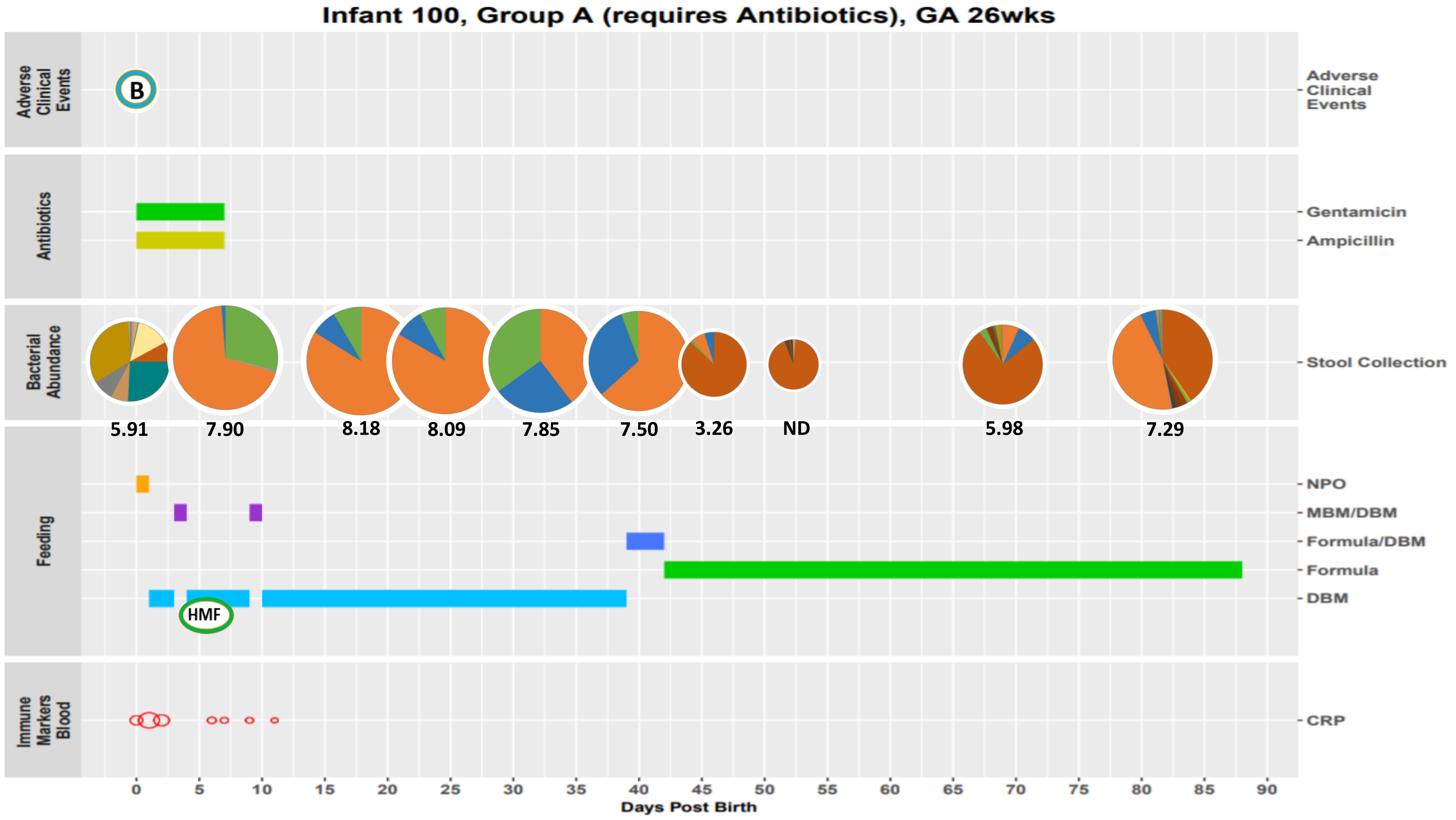

Infant 100, Group A (requires Antibiotics), GA 26wks
Adverse Clinical Events
B
Antibiotics
Gentamicin
Ampicillin
Bacterial Abundance
Stool Collection
5.91
7.90
8.18
8.09
7.85
7.50
3.26
ND
5.98
7.29
Feeding
NPO
MBM/DBM
Formula/DBM
Formula
DBM
HMF
Immune Markers Blood
CRP
0 5 10 15 20 25 30 35 40 45 50 55 60 65 70 75 80 85 90
Days Post Birth
