## Supplementary Figure S2 for "Antibiotics and the developing intestinal microbiome, metabolome and inflammatory environment: a randomized trial of preterm infants"

**Supplemental Figure S2. Abundance of top bacterial genera correlate with abundances of inferred metabolic pathways.** Heatmap of repeated measures correlation values between counts of the 10 most highly abundant bacterial genera and abundance of inferred metabolic pathways using PICRUSt2. Only bacterial genera with at least one significant correlation are depicted (p-values < 0.05). n=90 stool samples.

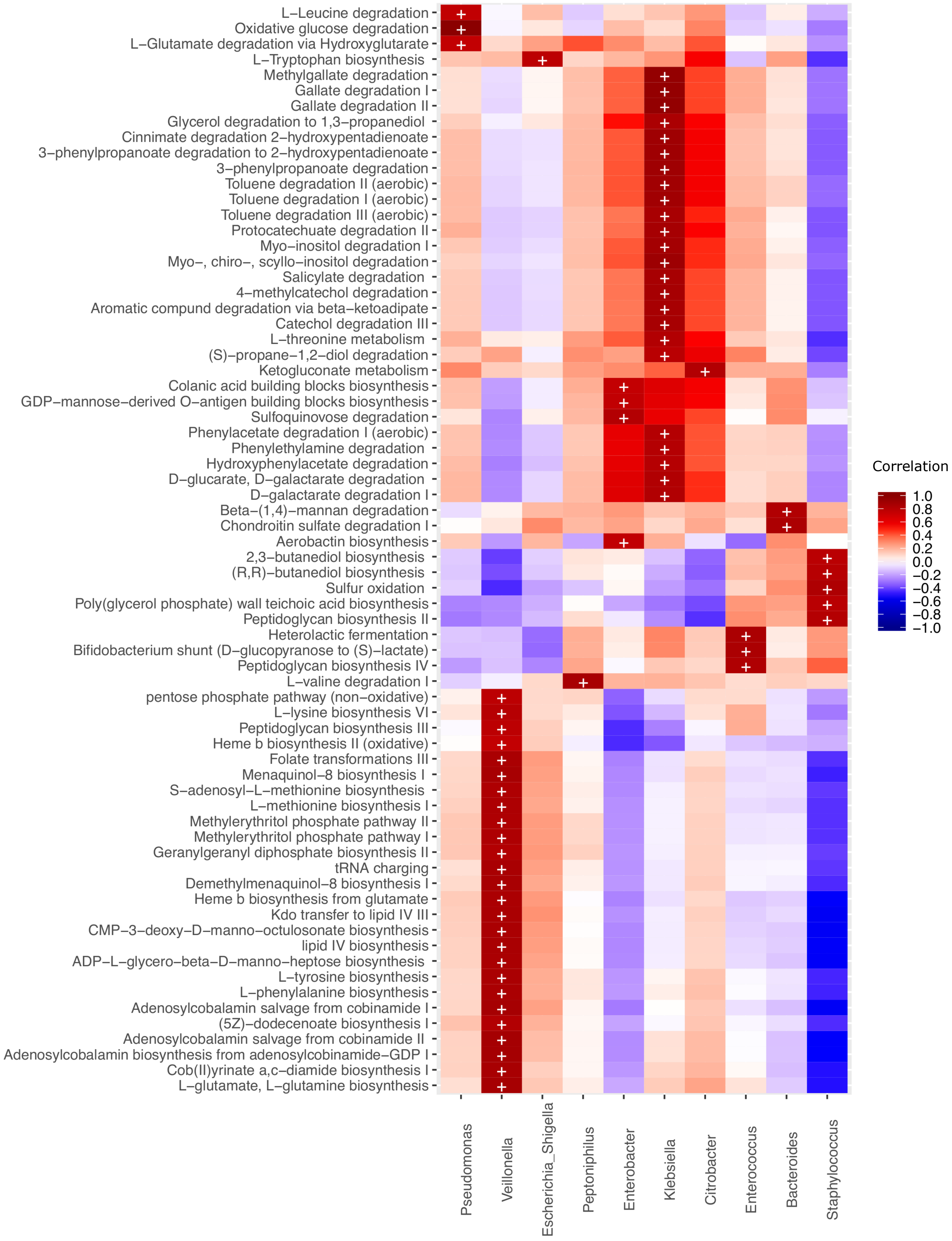

L−Leucine degradation
Oxidative glucose degradation
L−Glutamate degradation via Hydroxyglutarate
L−Tryptophan biosynthesis
Methylgallate degradation
Gallate degradation I
Gallate degradation II
Glycerol degradation to 1,3−propanediol
Cinnimate degradation 2−hydroxypentadienoate
3−phenylpropanoate degradation to 2−hydroxypentadienoate
3−phenylpropanoate degradation
Toluene degradation II (aerobic)
Toluene degradation I (aerobic)
Toluene degradation III (aerobic)
Protocatechuate degradation II
Myo−inositol degradation I
Myo−, chiro−, scyllo−inositol degradation
Salicylate degradation
4−methylcatechol degradation
Aromatic compund degradation via beta−ketoadipate
Catechol degradation III
L−threonine metabolism
(S)−propane−1,2−diol degradation
Ketogluconate metabolism
Colanic acid building blocks biosynthesis
GDP−mannose−derived O−antigen building blocks biosynthesis
Sulfoquinovose degradation
Phenylacetate degradation I (aerobic)
Phenylethylamine degradation
Hydroxyphenylacetate degradation
D−glucarate, D−galactarate degradation
D−galactarate degradation I
Beta−(1,4)−mannan degradation
Chondroitin sulfate degradation I
Aerobactin biosynthesis
2,3−butanediol biosynthesis
(R,R)−butanediol biosynthesis
Sulfur oxidation
Poly(glycerol phosphate) wall teichoic acid biosynthesis
Peptidoglycan biosynthesis II
Heterolactic fermentation
Bifidobacterium shunt (D−glucopyranose to (S)−lactate)
Peptidoglycan biosynthesis IV
L−valine degradation I
pentose phosphate pathway (non−oxidative)
L−lysine biosynthesis VI
Peptidoglycan biosynthesis III
Heme b biosynthesis II (oxidative)
Folate transformations III
Menaquinol−8 biosynthesis I
S−adenosyl−L−methionine biosynthesis
L−methionine biosynthesis I
Methylerythritol phosphate pathway II
Methylerythritol phosphate pathway I
Geranylgeranyl diphosphate biosynthesis II
tRNA charging
Demethylmenaquinol−8 biosynthesis I
Heme b biosynthesis from glutamate
Kdo transfer to lipid IV III
CMP−3−deoxy−D−manno−octulosonate biosynthesis
lipid IV biosynthesis
ADP−L−glycero−beta−D−manno−heptose biosynthesis
L−tyrosine biosynthesis
L−phenylalanine biosynthesis
Adenosylcobalamin salvage from cobinamide I
(5Z)−dodecenoate biosynthesis I
Adenosylcobalamin salvage from cobinamide II
Adenosylcobalamin biosynthesis from adenosylcobinamide−GDP I
Cob(II)yrinate a,c−diamide biosynthesis I
L−glutamate, L−glutamine biosynthesis
Pseudomonas
Veillonella
Escherichia_Shigella
Peptoniphilus
Enterobacter
Klebsiella
Citrobacter
Enterococcus
Bacteroides
Staphylococcus
Correlation
1.0
0.8
0.6
0.4
0.2
0.0
−0.2
−0.4
−0.6
−0.8
−1.0
