## Supplementary Table S1 for "Antibiotics and the developing intestinal microbiome, metabolome and inflammatory environment: a randomized trial of preterm infants"

Number and type of total antibiotics used by group and within the specified corrected GA analysis window.

**Antibiotics given within 48 hours after birth

|  | Group A** | Group B  (1 bailed infant) | Group C1** | Group C2 | Group C2Bailed** |
| --- | --- | --- | --- | --- | --- |
| # of days on antibiotics or antifungal (all routes) | 984 | 88 | 590 | 445 | 232 |
| # days of enteral/parenteral antibiotics total | 771 | 59 | 503 | 368 | 212 |
| # days of enteral/parenteral antibiotics x 28-39 weeks corrected GA | 476 | 14 | 479 | 272 | 179 |
| Antibiotics used | cefazolin; ampicillin; metronidazole; clindamycin; ceftriaxone; gentamicin; piperacillin-tazobactam; vancomycin; oxacillin; cefotaxime; azithromycin; ceftazidime; cefoxitin; meropenem; fluconazole | ampicillin; gentamicin; vancomycin; metronidazole | flagyl; oxacillin; cefotaxime; acyclovir; gentamicin; metronidazole; ampicillin; vancomycin; zosyn; azithromycin; tobramicyn; cefepime; fluconazole | vancomycin; ampicillin; oxacillin; metronidazole; penicillin; gentamicin; zosyn; cefotaxime; merem; flagyl; cefazolin; azithromycin; clindamycin; cefepime | vancomycin; cafazolin; zosyn; gentamicin; ampicillin; azithromycin; oxacillin; cefepime |
