## Supplementary Table S2 for "Antibiotics and the developing intestinal microbiome, metabolome and inflammatory environment: a randomized trial of preterm infants"

**Supplementary Table S2 – Samples for figure 1 by group and corrected gestational age**

The number of stool samples used in Figure 1 summarized by enrollment group and corrected gestational age.

|  | Group A | Group B | Group C1 | Group C2 | Group C2Bailed |
| --- | --- | --- | --- | --- | --- |
| Corrected GA 28 | 12 | 0 | 6 | 3 | 4 |
| Corrected GA 29 | 9 | 0 | 9 | 4 | 5 |
| Corrected GA 30 | 17 | 0 | 14 | 6 | 8 |
| Corrected GA 31 | 19 | 3 | 18 | 7 | 7 |
| Corrected GA 32 | 24 | 7 | 19 | 12 | 9 |
| Corrected GA 33 | 20 | 9 | 16 | 10 | 10 |
| Corrected GA 34 | 19 | 7 | 18 | 11 | 8 |
| Corrected GA 35 | 18 | 4 | 17 | 10 | 5 |
| Corrected GA 36 | 16 | 2 | 10 | 8 | 8 |
| Corrected GA 37 | 16 | 0 | 4 | 4 | 8 |
| Corrected GA 38 | 9 | 0 | 5 | 2 | 7 |
| Corrected GA 39 | 7 | 0 | 6 | 2 | 4 |
