## Supplementary Table S3 for "Antibiotics and the developing intestinal microbiome, metabolome and inflammatory environment: a randomized trial of preterm infants"

**Supplementary Table S3 - Samples for figures 2 and 3 by group, corrected gestational age and feeding type**

The number of stool samples used in Figure 2 summarized by enrollment group, corrected gestational age and feeding type. DBM: Donor Breast Milk; MBM: Mother’s Breast Milk; NPO: Nil Per Os (nothing by mouth)

|  | Group A | Group B | Group C1 | Group C2 | Group C2Bailed |
| --- | --- | --- | --- | --- | --- |
| Corrected GA 28 | 12 DBM: 8 MBM: 4 | 0 | 5 DBM: 3 MBM: 2 | 3 MBM: 3 | 3 MBM: 3 |
| Corrected GA 29 | 8 DBM: 4 MBM: 2 NPO: 2 | 0 | 7 DBM: 4 MBM: 3 | 4 DBM: 2 MBM: 2 | 4 MBM: 2 NPO: 2 |
| Corrected GA 30 | 15 DBM: 9 MBM: 4 NPO: 2 | 0 | 14 DBM: 4 Formula: 2 MBM: 5 MBM_DBM: 3 | 5 DBM: 2 MBM: 3 | 7 DBM: 2 MBM: 3 MBM_DBM: 2 |
| Corrected GA 31 | 18 DBM: 7 DBM_Formula: 5 Formula: 2 MBM: 4 | 0 | 15 DBM: 4 DBM_Formula: 2 MBM: 9 | 6 DBM: 3 MBM: 3 | 3 MBM: 3 |
| Corrected GA 32 | 24 DBM: 3 DBM_Formula: 3 Formula: 10 MBM: 6 MBM_Formula: 2 | 6 Formula: 4 MBM: 2 | 17 Formula: 6 MBM: 9 MBM_Formula: 2 | 9 Formula: 9 MBM: 7 | 7 Formula: 3 MBM: 4 |
| Corrected GA 33 | 19 DBM: 2 Formula: 11 MBM: 6 | 9 Formula: 3 MBM: 3 MBM_Formula: 3 | 16 Formula: 6 MBM: 8 MBM_Formula: 2 | 9 Formula: 5 MBM: 4 | 10 Formula: 2 MBM: 6 MBM_Formula: 2 |
| Corrected GA 34 | 19 Formula: 15 MBM: 4 | 6 Formula: 3 MBM: 3 | 18 Formula: 7 MBM: 8 MBM_Formula: 3 | 10 Formula: 5 MBM: 5 | 6 MBM: 6 |
| Corrected GA 35 | 15 Formula: 11 MBM: 4 | 2 MBM: 2 | 17 Formula: 8 MBM: 7 MBM_Formula: 2 | 10 Formula: 4 MBM: 6 | 3 MBM: 3 |
| Corrected GA 36 | 15 Formula: 9 MBM: 4 NPO: 2 | 2 MBM: 2 | 10 Formula: 4 MBM: 4 MBM_Formula: 2 | 8 Formula: 5 MBM: 3 | 8 Formula: 2 MBM: 4 MBM_Formula: 2 |
| Corrected GA 37 | 13 Formula: 10 MBM: 3 | 0 | 4 MBM: 2 MBM_Formula: 2 | 2 Formula: 2 | 7 MBM: 5 MBM_Formula: 2 |
| Corrected GA 38 | 6  Formula: 6 | 0 | 5  Formula: 2  MBM: 3 | 0 | 6  Formula: 3  MBM: 3 |
| Corrected GA 39 | 7 Formula: 7 | 0 | 5 Formula: 3 MBM: 2 | 0 | 2 Formula:2 |
